# Exploring Leishmania linear motifs directing protein secretion, sorting and autophagy

**DOI:** 10.1101/2023.07.05.547820

**Authors:** Andras Zeke, Toby J. Gibson, Laszlo Dobson

## Abstract

The pathogenic tropical flagellates *Leishmania* belong to an early-branching eukaryotic lineage (Kinetoplastida) with several unique features. Here, we explore three ancient protein targeting linear motif systems and their receptors and demonstrate how they resemble or differ from other eukaryotic organisms, including their hosts. Secretory signal peptides, endoplasmic reticulum (ER) retention motifs (KDEL motifs), and autophagy signals (motifs interacting with ATG8 family members) are essential components of cellular life. Although expected to be conserved, we observe that all three systems show a varying degree of divergence from the eukaryotic version observed in animals, plants, or fungi. We not only describe their behavior but also build predictive models that allow the prediction of localization or function for several proteins in *Leishmania* species for the first time. Several of these critical protein-protein interactions could serve as targets of selective antimicrobial agents against Leishmaniasis due to their divergence from the host.

## INTRODUCTION

Proper protein trafficking and localization within cells are crucial for maintaining cellular integrity and homeostasis. Eukaryotic organisms carry a largely conserved set of core signaling and sorting systems ensuring the proper execution of critical cellular processes. These not only include the signals for protein secretion (through the Sec61 translocon), but also instructions for the subsequent targeting of secreted proteins into appropriate compartments. Cytoplasmic proteins might also be subject to a broad variety of sorting phenomena, from nuclear import and export to secondary cytoplasm-to-vesicle targeting through autophagy. The receptors involved in these sorting systems typically utilize short, disordered protein motifs, highly conserved among Eukaryotic crown groups despite their divergence at ∼ 1 billion years ago^1,2^. Secretory signal recognition particles (SRPs), endoplasmic reticulum (ER) retaining receptors (KDEL receptors), and autophagy signaling ATG8 proteins are essential components of these processes, contributing to the regulation of protein localization, retrieval, and degradation. While each process has distinct mechanisms and functions, they collectively ensure efficient protein trafficking and cellular quality control.

The signal recognition particle (SRP) is a ribonucleoprotein complex in Eukaryota (as well as Archaea) that plays a vital role in protein targeting and translocation across cellular membranes. By recognizing specific signal sequences on the N-terminus of newly synthesized proteins, the SRP guides them to the SEC61 translocon on the endoplasmic reticulum (ER), allowing the biogenesis of both secreted and transmembrane proteins. Within this complex, the SRP54 protein is responsible for the recognition of N-terminal signal peptides or signal-anchors protruding from the translating ribosome. While signal peptides are co-translationally removed by a dedicated signal peptidase after targeting, the similarly SRP-dependently recognized signal-anchors are inserted to the membrane. Eukaryotic ER-resident proteins with a signal peptide but no transmembrane segment are retained in the lumen by a specific transport system involving the so-called KDEL receptors. These receptors (that obtained their namesake from the sequence they recognize) orchestrate the retrograde transport of target proteins, retrieving proteins containing the C-terminal KDEL motif from the Golgi apparatus back to the ER^3^. By maintaining the proper localization of ER-resident chaperones, KDEL receptors are also critical components of cellular quality control and homeostasis.

Autophagy is a cellular process responsible for the degradation and recycling of cytoplasmic components. It is regulated by an intricate signaling network involving autophagy-related proteins (ATGs) and key signaling pathways. Through cargo recognition mechanisms, autophagy selectively degrades damaged or unwanted proteins and organelles, promoting cellular renewal and adapting to nutrient deprivation. ATG8, a small ubiquitin-like modifier, is perhaps the most critical core component of this machinery. ATG8 proteins are conjugated to phosphatidylethanolamine lipids by the ubiquitin protease and ligase-like ATG4 proteins. Lipidated ATG8 not only directs phagophore membrane formation, it also serves as a recruiting agent to the autophagosome through multiple linear motifs. Among these motifs, the so-called LIR (LC3 interacting region) motifs are the best known autophagy regulators in animals, fungi and plants^4^.

Leishmaniasis is a neglected tropical disease with cutaneous, or systemic forms, caused by unicellular flagellates of the genus *Leishmania*, with a complex life cycle involving insect and mammalian hosts. These protozoan parasites are related to the *Trypanosoma* species, causing the African sleeping sickness as well as the South American Chagas disease. Classified as members of the basal eukaryotic group Kinetoplastida, it is now recognized that these dangerous pathogens are part of a much broader group also including free-living species (e.g. *Bodo saltans*) alongside the obligate pathogenic Trypanosomatids. Among Trypanosomatids, it is now clear that the ability to parasitize mammalian hosts evolved multiple times independently. Hence, the closest genetically studied relatives of *Leishmania* species are the insect parasite *Leptomonas* flagellates (forming the *Leishmaniinae* family or subfamily), and they are only very distantly related to the *Trypanosoma* genus ^5,6^. While the heterotrophic Kinetoplastids are connected to the secondarily photosynthetic *Euglena* species, they represent an early branch on the eukaryotic tree of life: Only the anaerobic metamonads (e.g. *Giardia intestinalis* from the group Fornicata) are thought to have diverged earlier^7^. Kinetoplastids present a whole array of unusual molecular biology features, including a polycistronic genome, with transcripts processed by trans-splicing, as well as a highly complex mitochondrial mRNA editing apparatus^6^. While Kinetoplastids appear to possess secretory signal peptides resembling other eukaryotes, as well as a fairly canonical ER physiology and autophagy, we sought to explore if these systems do match with those of higher eukaryotes at a molecular level. During our analysis, we made extensive use of the LeishMANIAdb^8^ database developed by our group. Our studies suggest that while these systems are of an ancient eukaryotic heritage, they heavily diverged in Kinetoplastids until becoming difficult to recognize in several cases.

## RESULTS AND DISCUSSION

### Kinetoplastid signal peptides differ considerably from the usual eukaryotic pattern

*Leishmania* species are known to have an unusual way to secrete proteins by utilizing exosomes^9^, often associated as the primary way to export proteins into the host cells. However, as in other Kinetoplastids, the canonical eukaryotic Sec61-dependent secretion is also present. The latter is indispensable for soluble lysosomal or ER-resident as well as transmembrane protein biogenesis. While the known *Leishmania* signal peptides (SPs) do have a broad similarity to generic eukaryotic signals, they more closely resemble bacterial type II signal peptides in several aspects, such as their amino acid content. For a concise analysis, we collected SPs predicted by SignalP6^10^ in LeishMANIAdb^8^ and in SwissProt^11^. When examining their hydrophobic region, we noticed that SPs in *Leishmania* strongly prefer small hydrophobic amino acids, resulting in higher alanine and valine and lower leucine and phenylalanine content, compared to other eukaryotes (Figure 1/A, Supplementary Table 1). Although the difference is not significant in purely statistical terms (using proteome-level statistics), it is still strikingly apparent in many SP examples.

**Figure 1:**
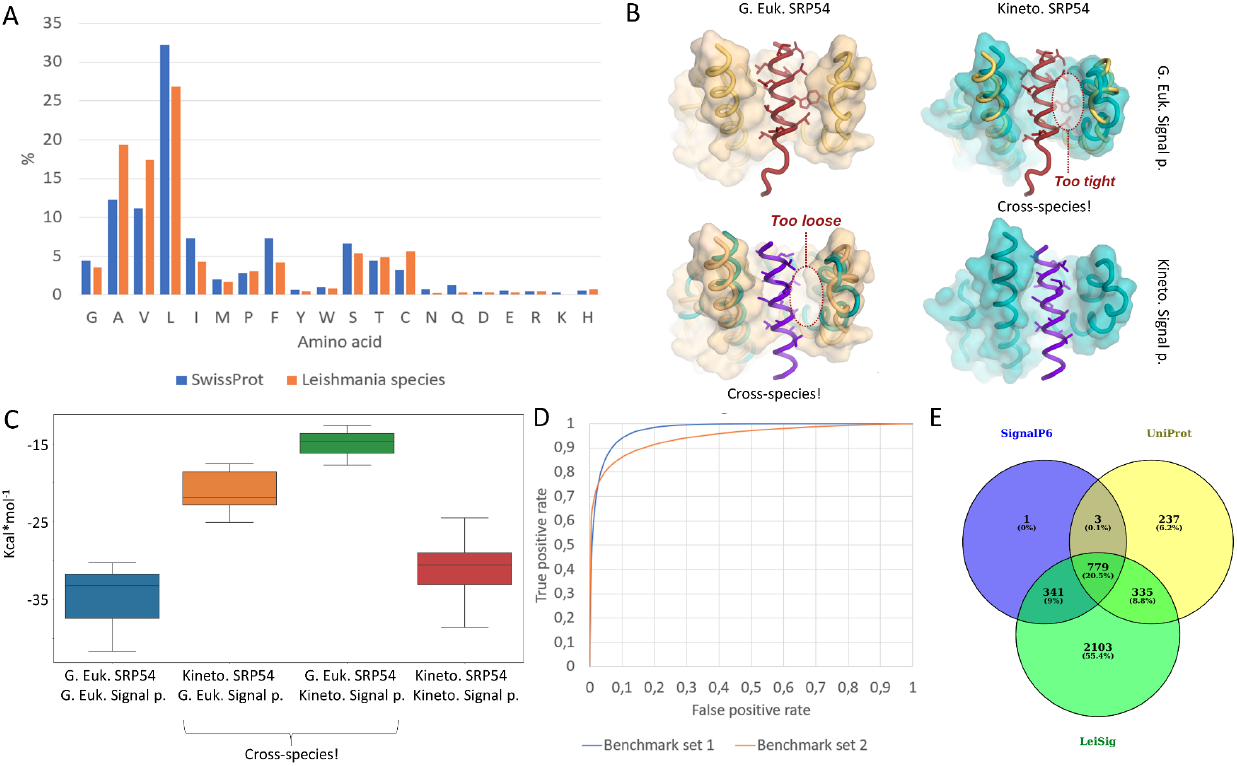
A: Amino acid distribution in the hydrophobic regions of signal peptides. B: left: AF2 predicted structures with different combinations between SRP54 proteins and signal peptides from L. infantum and S. cerevisiae. C: Energy (free enthalpy) calculation results from different combinations of SRP54 and signal peptides D: Receiver Operating Characteristics (ROC) of the Leishmania specific signal peptide prediction method (LeiSig) E: Signal peptide predictions on 5 Leishmania species with different approaches.

To study the structural background of this unusual finding, we resorted to structural modeling of SRP54 M domains, responsible for the recognition of signal peptides. As the only experimentally available structure was of an archaeal origin, we had to model a crown group eukaryotic complex as well (we chose the yeast *Saccharomyces cerevisiae*) with a cognate signal peptide. In the original archaeal complex, three helices (αM1, αM2, αM4) in the M domain recognize and form contacts with a signal peptide. Additionally, αM1b and αMF might also help the recognition by closing on the signal peptide ^12^ (Supplementary Figure 1). Although no eukaryotic complex has been determined yet, homology modeling by Phyre 2^13^ and AlphaFold2^14^ (AF2) could readily be utilized to assess structural differences. Despite the similar main topology, Phyre2 struggles capturing secondary structure elements around the SP, likely because we could only model the monomer and not the bound structure (Supplementary Figure 2). We compared the AF2 predicted structure of SRP54 M domain together with a SP in different combinations from *L. infantum* and *S. cerevisiae*. While cognate complexes including proteins from the same species seem to be compact and do not contain empty volumes or clashes, predicted structures of cross-species combinations are markedly problematic: *S. cerevisiae* SRP54 *and L. infantum* SP form a too loose structure, while *L. infantum* SRP54 and *S. cerevisiae* SP contains visible clashes (Figure 1/B).

To further support our observation, we generated all combinations between SRP54 M domains from select species (*A. thaliana, H. sapiens, S. cerevisiae, S. solfataricus, B. saltans, T. cruzi, T. brucei, L. seymouri, L. infantum*) and SP complexes (*L. infantum* and *S. cerevisiae)* using AF2 and calculated the overall stability of these structures. Considering a “generic” eukaryotic SRP54 and *S. cerevisiae* SP, their mean stability suggested a reasonable energy threshold for the system (−35.07 kcal/mol). On the other hand, when we used cross-species combinations (“generic” eukaryotic SRP54 and *L. infantum* SP, or *L. infantum* SRP54 and generic eukaryotic SP) their calculated binding energy was much lower (−14.79 kcal/mol and -21.20 kcal/mol, respectively) in absolute terms. Importantly, with cognate kinetoplastid SRP54 and *L. infantum* SP the calculated mean energy is similar (−31.17 kcal/mol) to other eukaryotes (Figure 1/C, Supplementary Table 2).

These calculations suggest that the Leishmania receptor is unnaturally strained, with a weaker energy estimate when modelled with general eukaryotic SPs, thus the complex between a “usual eukaryotic” signal peptide and the kinetoplastid receptor was unlikely to form in vivo. Note that the yeast peptide also serves as a decent surrogate for animal or plant signals as well. This means that the recognition surface of kinetoplastid SRP45 is unusually narrow, explaining the Ala/Val preference. We believe that this also means that the latter might be unique enough to serve as a target for some novel antibiotics.

If the kinetoplastid signal peptides are so different from other, better studied eukaryotic organisms, we wondered if some SP examples might be divergent enough to be missed by classical prediction algorithms. To this aim, we extended the set of known *Leishmania* signal peptides using the homologous information stored in LeishMANIAdb: when SignalP6 predicted a signal peptide (Supplementary Table 3), we checked I) the number of gaps in the signal peptide alignment II) the sequence identity of the signal peptide within the homology group (Supplementary Table 4). Using this approach, we collected hundreds of proteins where SignalP6 predicts a signal peptide on some sequences, but not on other closely related ones, even when the sequence identity was strikingly high and there were no gaps in the segment. Altogether we assembled 1027 such candidate sequences, from which 465 had lower than 50% gap content and more than 30% sequence identity.

Homology mapping is only helpful when we can assign related proteins, however in LeishMANIAdb we have thousands of proteins without many homologs. To see whether these proteins have a signal peptide, we developed a machine learning-based prediction method. First we created a training set using positive examples from SignalP6 predicted signal peptides and the 465 protein where homology could be used to assign signal peptides. For negative predictions we included membrane proteins (where the first membrane region predicted by CCTOP^15^ is at the N-terminal 50 residues of the proteins) and proteins without predicted membrane region or signal peptide. The dataset was divided into training and independent test sets (benchmark set 1, Supplementary Table 5,6). We also prepared an additional dataset of 89 proteins (by randomly selecting proteins from *Leishmania* proteomes), where we manually checked the presence of signal peptides using deep homology searches (benchmark set 2, Supplementary Table 7). Next we used transfer learning to fine-tune a pre-trained protein language model on the training dataset^16^. On benchmark set 1 we achieved 93% balanced accuracy and 95% Area Under Curve, while on benchmark set 2 we achieved 81% balanced accuracy and 92% Area Under Curve (Figure 1/D, Supplementary Table 8). By setting the cutoff to a higher value (from default 0.5) specificity can be further raised. The developed method (LeiSig) is available for download at https://leishmaniadb.ttk.hu/files/LeiSig.zip. Despite the many shortcomings of our method (see Methods), this approach (that confers high specificity at the cost of sensitivity) revealed 3558 candidate proteins with SPs in 5 Leishmania species, from which 2103 were not predicted by any other method (Figure 1/E, Supplementary Table 9). This suggests that a good number of potentially novel secreted or transmembrane proteins are present in *Leishmania* proteomes.

### The endoplasmic reticulum retaining KDEL motif is unusually divergent in parasitic kinetoplastids

After the signal peptides, we turned our attention to another unusual feature of *Leishmania*. The group of constitutively endoplasmic reticulum (ER) lumenal proteins encompasses many highly conserved proteins with roles in secreted protein folding, glycosylation and quality control. In all known Eukaryota, these proteins are constitutively sorted to the ER with the help of a retrograde transport system from the Golgi apparatus, utilizing a single receptor (also known as the KDEL receptor). However, the usual, highly conserved (H/K)DEL consensus at the C-termini of these proteins, that is conserved in animals, plants and fungi (Figure 2/A) cannot be found in *Leishmania* species. Instead, careful examination shows that the conserved, known or predicted ER resident *Leishmania* proteins carry a carboxy-terminal DL motif (Figure 2/B). This consensus differs from all other known eukaryotic organisms outside Kinetoplastida, including the early-branching flagellate *Giardia intestinalis*, but matches with other kinetoplastid species.

**Figure 2:**
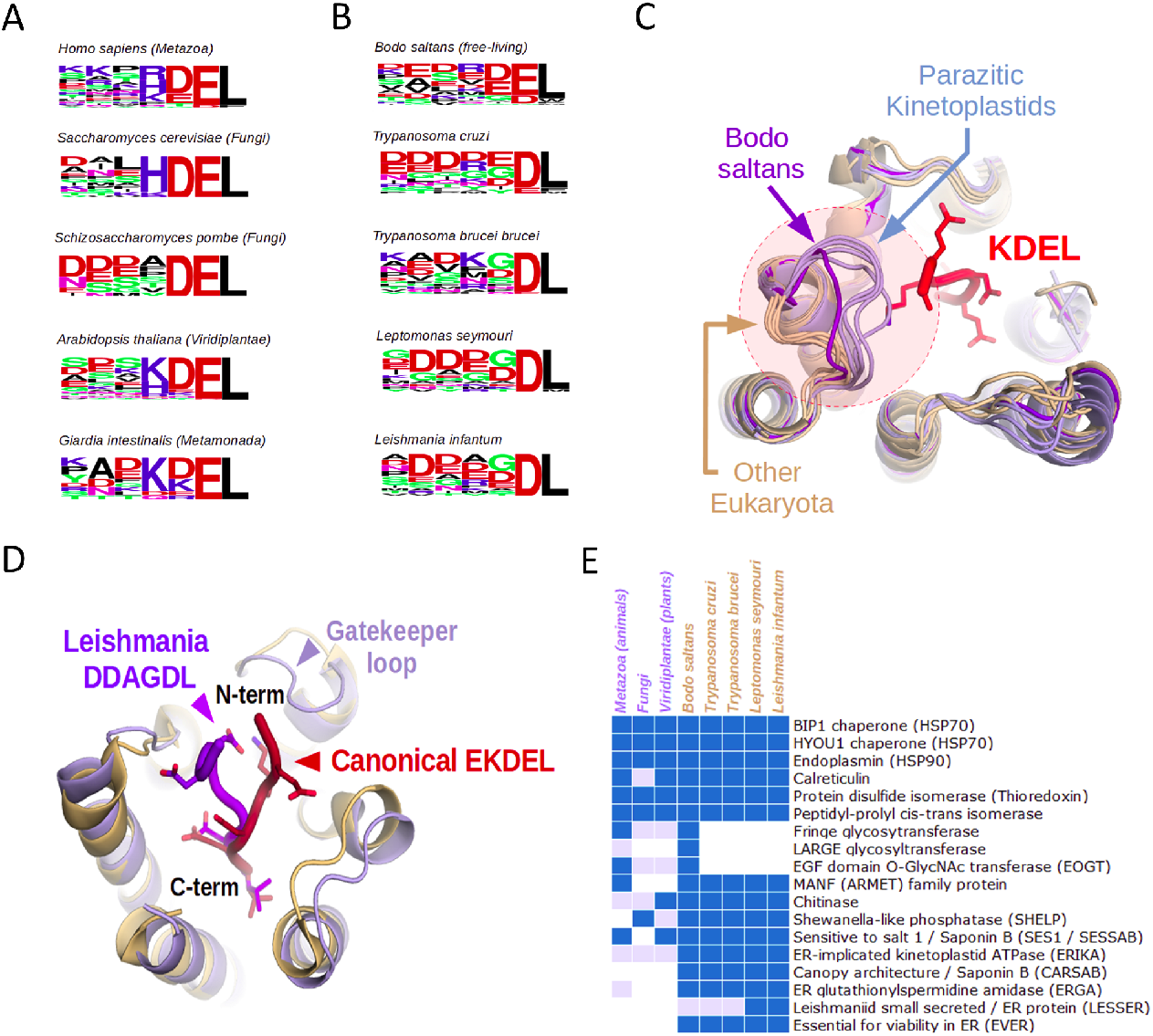
A: Comparison of the sequence logos of all known or predicted KDEL-like motifs of five eukaryotes from different lineages show its conservation. B: This original K/H/RDEL consensus is altered in kinetoplastids, mostly yielding peptides ending in DL. C: Superimposed protein models show that the gatekeeper loop of the ER retrieval receptor (KDEL receptor) is different in kinetoplastids, clashing with the main chain of an incoming, canonical KDEL peptide (pdb:6I6H) in case of all examined parasitic species, but not in the free-living Bodo saltans. D: HADDOCK models show that the long gatekeeper loop forces the ligands of the Leishmania receptor to take different main chain conformation than in the mammalian KDEL receptor. E: Recognizing the altered consensus of receptors allows identification of conserved families of kinetoplastid ER-resident proteins including newly-identified protein groups (Blue bars show the presence of KDEL-like peptide-containing orthologs, while light blue bars indicate homologous proteins without KDEL.)

A close examination of the AlphaFold-predicted ER-retaining KDEL receptor structures helps to explain this discrepancy. Although the bottom of the receptor has almost 100% identity in terms of the amino acids lining the receptor cavity, the entrance to the KDEL receptor in *Leishmania* is narrowed by an elongated loop, leading to a clash when superimposed on animal KDEL receptor-ligand crystal structures (Figure 2/C). This forces the peptide chain in *Leishmania* species to adopt a substantially different geometry when entering the receptor. To demonstrate this phenomenon, we built a model of *Leishmania infantum* KDEL receptor based on the AlphaFold2 structure with a peptide ligand (best possible consensus sequence), docked and optimized its geometry with HADDOCK^17^ (see Methods). Comparison with the crystal structure of an animal KDEL receptor complex clearly shows the different main chain geometry (Figure 2/D).

Interestingly, this deviation from the classical patterns is not restricted to *Leishmania* species only. The same “gatekeeper” loop features in the KDEL receptor are also seen in other Kinetoplastid species, such as *Trypanosoma brucei brucei*. However, with differently charged amino acids, the resulting KxDL-like motif is closer to the canonical one than in *Leishmania sp*. Curiously, we found that the aberrant DL-preference at the C-terminus is restricted to parasitic lineages only. Detailed analysis of motifs in the free-living kinetoplastid *Bodo saltans* shows that it still prefers the canonical KDEL motif, although with a relatively broad variation of individual positions. In this early-branching species, the elongated loop consists of 5 amino acids (compared to 6 in parasitic kinetoplastids and 3 with other eukaryotes), but it is directed outwards, leading to a broadly open cavity, presumably with relatively permissive ligand binding properties. It is the further elongation of this “gatekeeper” loop that led to a restricted, but non-canonical KDEL-like DL motif in parasites. While this motif differs considerably from their host organisms, it is unclear if it is truly orthogonal to the ER receptor of their (animal or plant) host. Therefore the biological role of this unusual divergence (if any) is currently unclear.

The redefined motif regular expression together with the developed signal prediction method enabled us to scan species from LeishMANIAdb for candidate ER proteins (Supplementary Table 10). To further limit possible false positive hits, we also supplied information about protein disorder and localization. We also supplied the position-specific scoring matrix calculated by PSSMSearch^18^ to perform a more sensitive scan (Supplementary Table 11). Recognizing the correct KDEL-equivalent motif in Leishmania species, we not only managed to confirm conservation of well-known ER-resident proteins: Hsp70 (both the BIP and HYOU1 families) and Hsp90 (endoplasmin) chaperones, calreticulin, protein-disulfide isomerases and peptidyl-prolyl isomerases in *Leishmania* species, but also a number of other, unusual proteins (Figure 2/E, Supplementary Material). We located an ER-resident chitinase in *Leishmaniinae*, with a conserved KDEL-like signal also found in other eukaryotes, though not in animals. The relatively poorly known MANF (Mesencephalic astrocyte-derived neurotrophic factor, also known as ARMET) proteins that have previously only been identified in animals, might be ancient eukaryotic chaperones^19^. We uncovered more than one family of Saponin B domain containing proteins, one distinctly homologous to the Arabidopsis sensitive to salt (SES1) protein (SESSAB)^20,21^, and another, different one showing some architectural similarity to Canopy proteins found in animals and plants (CARSAB)^22^. We also confidently identified a kinetoplastid Shewanella-like phosphatase (SHELP) subfamily, with homologous ER retention signals already found in *Euglena*, hinting at an ancient origin, even if their function is currently unclear^23^.

Notably, kinetoplastids also seem to have paralogous pairs of some enzymes, with one lineage consistently bearing a KDEL-equivalent signal (in addition to a signal peptide), and another pair that does not. This is the case with torsin family ATPases (with a KDEL-bearing paralog, that we termed ERIKA dissimilar to animal torsins)^24^, as well as with the better-known trypanothione synthases. In the latter case, the ER-resident glutathionylspermidine amidase (ERGA) might have a different role in trypanothione metabolism, than the better-known, membrane-anchored paralog^25^.

Finally, there are examples with protein families consistently bearing an ER retention signal (in addition to the signal peptide) in *Leishmania* species as well as their relatives, but without any known homology to proteins in other eukaryotes. One such protein (LESSER), with a very small domain, has an ER retention signal in *Leishmaniinae*, but appears to be secreted in other kinetoplastids. Another, even more intriguing kinetoplastid-specific protein (EVER) has recently been identified to be essential for viability in *L. donovani*^*26*^. Although originally described as a secreted protein (due to the signal peptide), we now confidently predict it to be an ER component. The fact that *Leishmania* species (as well as other parasitic trypanosomatids) seem to have an overall reduced inventory of ER-resident enzymes compared to *Bodo saltans* is significant. This gene loss suggests that many retained ER-resident proteins might absolutely be vital for *Leishmania*, possibly providing decent, specific drug targets due to their uniqueness or divergence.

### The identification of LIR-like motifs reveals a fast-evolving autophagy network in Leishmaniids

ATG8 proteins are small ubiquitin-like modifiers essential for autophagy in all known eukaryotic organisms^27^. Unlike ubiquitin, they are covalently attached to the membrane lipid phosphatidylethanolamine (by ATG4 enzymes), thereby directing the formation of the phagophore membrane. The location, extent and cellular contents engulfed by the autophagosome are dictated by ATG8-interacting proteins that typically bind ATG8 through the so-called LIR motifs in both animals, fungi and plants. Although minor variations exist (e.g. in animals, ATG8 proteins have split into LC3 and GABARAP subfamilies, with different LIR motif ligands)^28^, this essential linear motif is largely conserved across known eukaryotes with only mild alterations.

As already noted by previous studies, *Leishmania* and *Leptomonas* species have split their ancestral ATG8 / ATG12 proteins into a multitude of paralogs^29^, with a complicated nomenclature, but major differences were shown in other autophagy related systems too^30^. Our careful, detailed analysis of kinetoplastid ATG8 evolutionary trees shows that there are five different, major families of ATG8s in *Leishmania* species^31^. The previously identified ATG8A, ATG8B and ATG8C subgroups are highly divergent and atypical in terms of sequence and structure. We observed that the previously annotated *Leishmania* ATG12 gene has a sibling evolutionary relationship to divergent Leishmania ATG8A, ATG8B and ATG8C proteins, and these groups all seem to have evolved from a single ATG12-like gene also found in other trypanosomatids (Supplementary Figure 3). Unfortunately, it cannot be decided from the available scarce data the kinetoplastid ATG12 would biochemically more resemble canonical eukaryotic ATG8 or ATG12 in function. ATG8 and ATG12 proteins share strong structural and sequence homology, having evolved from the same ancestral ubiquitin-like modifier^32^. Notably, the kinetoplastid ATG12 proteins require cleavage by proteases, similarly to ATG8s (processed by ATG4 enzymes) and unlike other eukaryotic ATG12s^31^. Finally, the

ATG8 gene and protein of *Leishmania* species incorporates fairly canonical members, with a surface highly similar to other eukaryotic ATG8 proteins. A careful comparison of surfaces shows that the LIR binding surface has been severely altered in ATG8A, B, C and ATG12 proteins (Figure 3/A). Therefore we assume that these proteins are incapable of binding the canonical LIR motifs, these motifs can only target ATG8.

**Figure 3:**
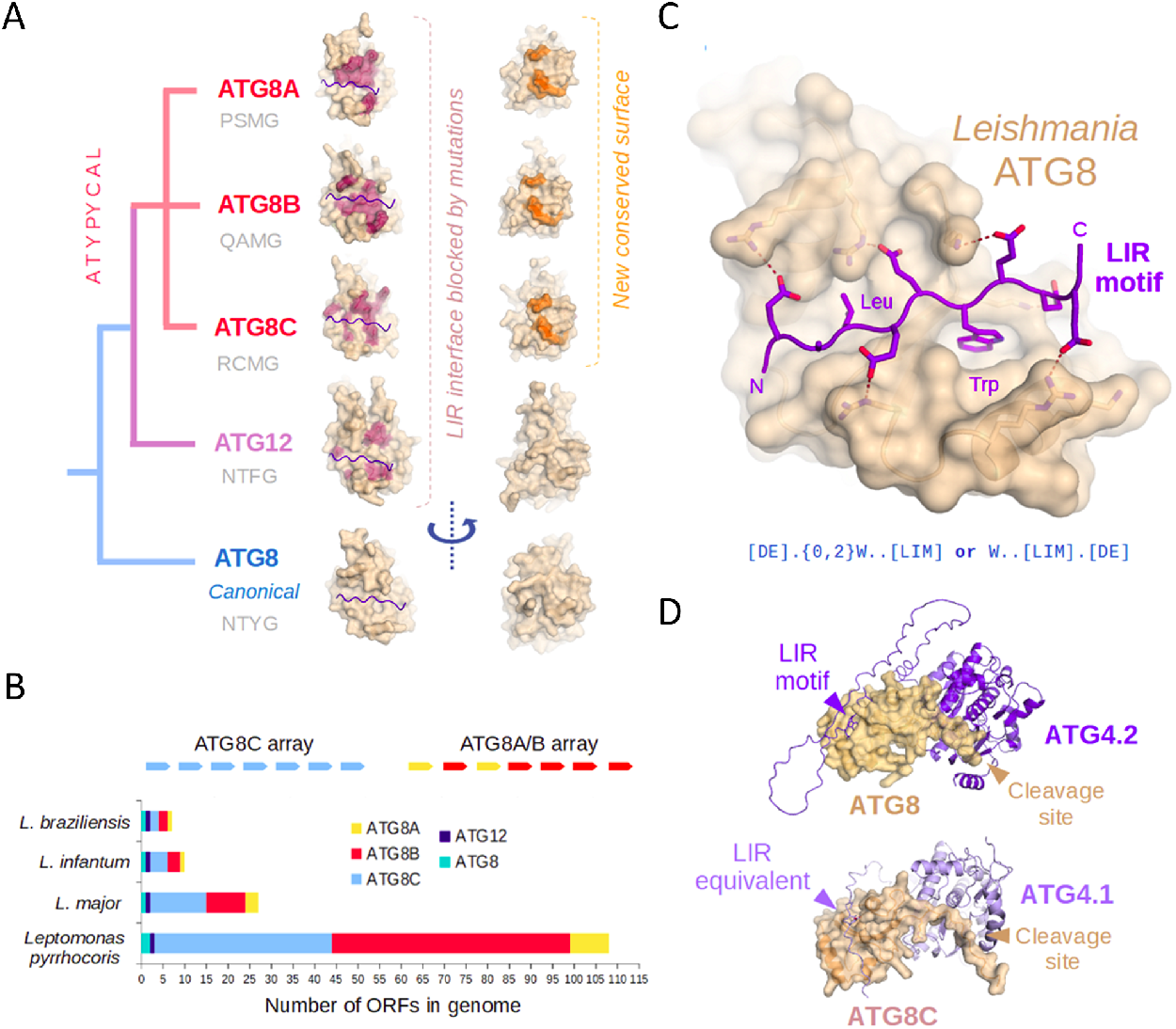
A: Our evolutionary model of ATG8 protein groups in Leishmaniinae. The surface mutations expected to block LIR binding are shown in red, while the novel invariant surface is in orange. The small structural figures show matching protein surfaces from the same angle. Predicted ATG4 cleavage sites (P-4 to P-1) for each protein group are exemplified by a sequence written in gray below each group name. B: The ATG8A, ATG8B and ATG8C genes are typically found in two tandem arrays within leishmanial genomes but their copy numbers vary highly across species. C: The preserved LIR peptide-binding ability of ATG8 proteins is illustrated by a simulated model between this protein and an idealized Leishmania LIR motif (satisfying both the canonical as well as an alternative consensus, written below as a regular expression). D: The simulated complexes between ATG8 and the enzyme ATG4.2 and between ATG8C and ATG4.2 are highly similar and betray a potential existence of a LIR-equivalent motif for atypical ATG8s.

While neither the ATG8A, ATG8B or ATG8C proteins appear to be capable of LIR motif binding (as that interface is not conserved), they do have another putative protein-protein interaction surface (Figure 3/A). The latter region that is invariant across these atypical ATG8s falls fairly close to the so-called UIM (ubiquitin interacting motif) binding surface of ATG8 proteins found in other eukaryotes^33^. Thus it is quite likely that this interface is functional, although its partners are currently unknown. Notably, most ATG8A, ATG8B or ATG8C *Leishmania* proteins present two cysteine amino acids adjacent each other at this region, suggesting a possible redox-dependent regulation.

As evidence of recent diversification, most atypical ATG8 genes are found as part of large gene arrays in *Leishmaniinae*. ATG8C genes form an array of their own, while ATG8A/B genes form joint arrays at a different genomic location, with a lot of identical or highly similar genes (Figure 3/B, Supplementary Table 12-13). Most of these arrays also contain fragmentary, incomplete ATG8-like ORFs, that nevertheless might contribute to recombination between related genes. This rampant gene amplification is clearly not due to adaptation to mammalian hosts, as matching arrays are already seen in the insect parasite *Leptomonas pyrrhocoris*. Still, the ATG8-like gene copy number increases at each separate cluster are remarkably consistent across species. Interestingly, ATG12 genes do not seem to be subject to gene amplification, nor are the canonical ATG8 genes (Figure 3/B).

Encouraged by the conservation of the ATG8 surface, we set out to identify potential LIR motif-containing, autophagy-related proteins in *Leishmania* genomes (Table 1, also see Supplementary Table 14-15 for more details). We found a number of plausible hits (Supplementary Table 14) using a conservative, Trp-containing consensus [DE].{0,2}W..[LIM], including the *Leishmania* ATG4.2 enzyme itself (we uncovered no such biologically relevant hits with a Phe-containing motif). In some proteins, these canonical eukaryotic LIR motifs were found as multiplets, suggesting strong biochemical connection to the autophagosome. However, the conservation pattern of motif singlets and multiplets also suggested that a variation of this Trp-containing LIR motif might also be functional in *Leishmania*. Here, the requirement for the first negatively charged amino acid is moved from the N-terminus to the C-terminus, yielding a consensus W..[LIM].[DE] (Supplementary Table 15). The latter motif was fully supported by our structural modeling of the Leishmania ATG8-LIR complex (Figure 3/C).

Using our *Leishmania*-specific LIR motif definitions, we uncovered a considerable number of proteins carrying either the original LIR or its variant motif, with a subset of manually selected, most likely targets shown on Table 1. Both LIR subtypes tended to occur in functionally similar proteins. Calpain-like proteases are a large and diverse, but enigmatic protein family in *Leishmania*, where only a portion of proteins preserve the proteolytic functions^34^. Intriguingly, we found that calpain-like proteins often carry LIR motifs, including motif multiplets, evolved independently from each other. For the first time, this suggests an important physiological function of *Leishmania* calpain-like proteins in autophagy. We also identified LIR motifs in *Leishmania* Neurobeachin/ALFY and FIS1 orthologs (both protein families are involved in autophagy in animals)^35,36^. Other LIR instances also show some intrinsic logic (such as the presence of such motifs together with secreted protein quality control associated PUB domains or the bacterial glycan-sensing LysM domains involved in clearance of invading bacteria^37^). Intriguingly, most of the LIR motif instances we identified had a very shallow ancestry. These motifs were rarely conserved beyond the *Leishmania + Leptomonas* (*Leishmaniinae*) group, despite the overall good surface conservation of ATG8. Thus we must assume the *Leishmania* autophagy network as having been rewired extensively and relatively recently in terms of evolution.

**Table 1:**
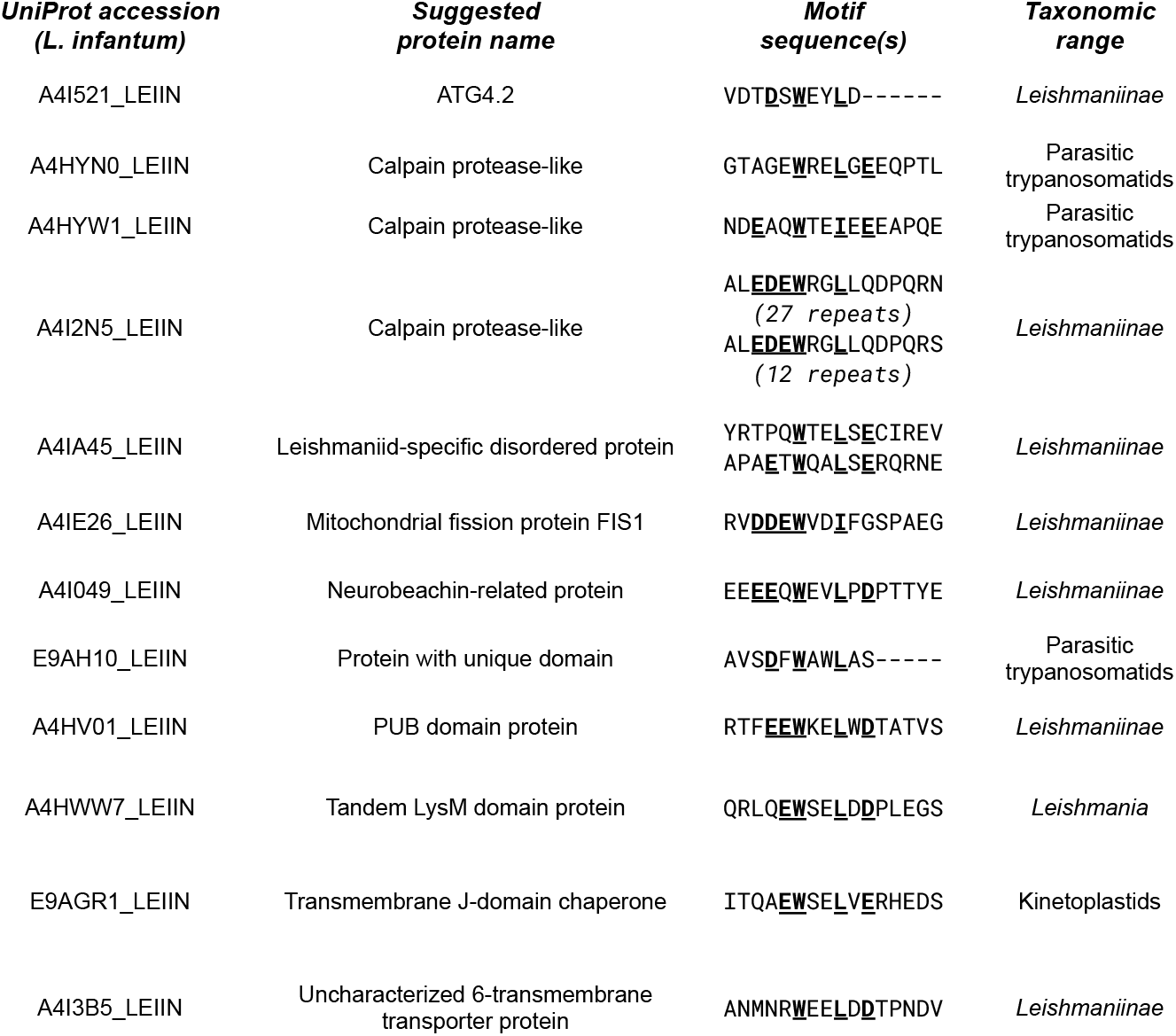
select Leishmania infantum proteins with likely functional LIR motifs and the evolutionary conservation of the motifs

Since AlphaFold2 multimer has proven rather successful in predicting the correct geometry of LIR motifs^38^ and other ATG8 partners, we applied it to *Leishmania* complexes as well. Our models (Figure 3/B) suggest that the *Leishmania* ATG8 is more charged than the mammalian ATG8s (LC3 / GABARAP), with a deep hydrophobic pocket binding Trp. Accordingly, *Leishmania* LIR motifs can likely engage in strong, charged interactions internally as well as at both ends of the motif, explaining why these motifs can have Asp/Glu amino acids facultatively at either end.

Out of our suggested autophagy regulators, ATG4.2 is by far the most solid. The presence of a LIR motif on the extreme C-terminus of ATG4 is almost warranted, as it is also found there in many other eukaryotes. Interestingly, all *Leishmania* relatives possess not one, but two clearly distinguishable ATG4 paralogs (ATG4.1 and ATG4.2), where only ATG4.2 has a LIR motif. *Leishmania* ATG4.1 and ATG4.2 were suggested to selectively process different ATG8-like proteins in the past. However, no evolutionarily or structurally consistent pattern has been reported^31^. Based on structural and evolutionary arguments alone, we speculate that the canonical *Leishmania* ATG8 is probably processed by the matching LIR-motif containing ATG4.2 in cells. This would leave ATG4.1 to be the main protease responsible for the activation of ATG8B, ATG8C and possibly ATG8A, that all have a similar cleavage site, but different from that of ATG8. To corroborate this theory, we used AlphaFold2 multimer to build a complete structure of the hypothetical complex between ATG4.2 and ATG8. This complex displays (Figure 3/D) excellent structural compatibility between partners, a perfectly fit cleavage site, complete with a LIR motif from the C-terminus docked into ATG8. We also had the potential complex of ATG4.1 and ATG8C modeled, and this gave a surprising suggestion: The C-terminus of ATG4.1 has a sequence very different from canonical LIR motifs, and yet it has the potential to interact with the atypical ATG8 relatives in a similar manner. Unfortunately, we cannot predict these LIR-equivalent, beta-sheet forming linear motifs yet, unless at least one example is experimentally validated first.

Out of the three biochemical systems studied in the current article, the ATG8 system is the most complex, yet least amenable to antimicrobial development. *Leishmania* LIR motifs are too similar to human ones, and we still do not know enough of the novel, kinetoplastid-specific protein-protein interactions and linear motifs to pinpoint an exact molecular target.

## METHODS

### Signal peptide predictions

We downloaded Leishmania reference proteomes (*L. major, L. infantum, L. mexicana, L. braziliensis, L. donovani)* from LeishMANIAdb^8^, while SwissProt^11^ was used as a reference. Signal peptides were predicted using SignalP6.0^10^. Amino acid distribution was calculated using these results (Figure 1/A). SRP54 protein (*A. thaliana, H. sapiens, S. solfataricus, B. saltans, T. cruzi, T. brucei, L. seymouri, L. infantum*) combined with signal peptide (*S. cerevisiae, L. infantum*) were predicted using ColabFold^39^ using the crystal structure as a template^12^. SRP54 was also predicted using Phyre2^13^. We used BLAST^40^ to search for the closest hit for *L. infantum* signal peptide (UniProt: A4HY78/A4HY79) considering the *S. cerevisiae* sequence from the crystal structure (notably the first hit was also predicted to have SP by SignalP6, Figure 1/B). FoldX^41^ was used to relax the structures and to calculate their stability (Figure 1/C).

For prediction, we prepared positive (signal peptide) and negative datasets (other) using the following approach: we selected proteins with predicted signal peptide, then extended these predictions to homologs using LeishMANIAdb (max 50% gap content and more than 30% sequence identity were required). Negative set contains proteins with membrane region predicted in the first 50 residues by CCTOP^15^, and other proteins (without any kind of prediction). Positive data were used multiple times so the two dataset had equal size. The positive and the negative dataset were splitted into training and test (benchmark set 1) sets (90% and 10%). In parallel, we randomly selected 100 proteins and manually annotated whether they have a signal peptide, or not (from among 11 proteins we couldn’t make a strong decision, these were omitted). The language model used the pretrained data (esm2_t12_35M_UR50D) fine-tuned on the constructed training dataset^16^ (Figure 1/D). After predicting sigal peptides on the 5 Leishmania reference proteomes, we compared our result with other methods (Figure 1/E).

As for the limitation of our prediction algorithm, the lack of experimental data is a key problem. In addition, we could not ensure that the test set is fully independent: reducing redundancy to 70% would remove 80% of the sequences (as these are short, N-terminal segments only), therefore this approach would not have left enough data to train and test the network. Therefore our results should better be interpreted as an extension to already recognized signal peptide prediction methods. There is also a high ambiguity whether membrane regions from the negative set are truly membrane segments, and not signal peptides. Although CCTOP is equipped to discriminate against these regions, it also relies on earlier versions of SignalP that we meant to amend here. Notably, this rather strict approach will less likely yield false positive hits.

### Modelling of KDEL receptors and identification of ER-resident proteins

To explore KDEL-equivalent motifs in kinetoplastid proteomes, we first observed the C-termini of proteins from universal eukaryotic ER-resident protein families (BIP1, HYOU1, Endoplasmin, Calreticulin, Thioredoxins, Cyclophilins, etc., Figure 2/A). Then we used the observation to search for similar, signal peptide-containing proteins in the *Leishmania infantum* and *Leishmania donovani* proteomes with either C-terminal El or DL consensus sequence (Figure 2/B). Then we assessed conservation of hits to increase our confidence in predictions. To compare them to the ER-resident proteome of the free-living *Bodo saltans*, we applied evolutionary conservation inferences (all protein families that have a KDEL-equivalent motif in any proteomes). Alignments were prepared using ClustalOmega^42^. We used IUPred3^43^ and AlphaFold2^44^ to find disordered regions, and DeepLoc^45^ to predict protein localizations.

Kinetoplastid KDEL-like motif logos presented on Figure 2/B include both known and novel (predicted ER-resident) families. For non-kinetoplastid eukaryotic organisms, we used annotated ER-resident examples for logo generation, except for *Giardia intestinalis*, where we had to resort to the same EL / DL C-terminus search in conjunction with the signal peptide requirement to predict ER-retained proteins. To see our complete motif collection with evolutionary comparisons across these organisms, see supplementary. Novel kinetoplastid families were analyzed in evolutionary terms using UniProt BLAST searches and alignments (Supplementary Material). Families that either had no previously known homologs, or possessed multiple paralogs with only different branches studied were given new, human-readable names and suggested abbreviations (Figure 2/C).

For comparative analysis, we used the AlphaFold2 (AF2) predicted models of orthologous KDEL receptors from the following species: *Homo sapiens* (UniProt: P24390), *Saccharomyces cerevisiae* (UniProt: P18414), *Schizosaccharomyces pombe* (UniProt: O94270), *Arabidopsis thaliana* (UniProt: P35402), *Giardia intestinalis* (UniProt: A8B6P2), *Bodo saltans* (UniProt: A0A0S4JT51), *Trypanosoma cruzi* (UniProt: Q4DMI8), *Trypanosoma brucei brucei* (UniProt: Q384K5), *Leptomonas seymouri* (UniProt: A0A0N1I068), and *Leishmania infantum* (UniProt: A4I3D7). Visual analysis of the structures was done by superimposing them over an experimentally determined KDEL-receptor complex (Figure 2/D, pdb: 6I6H). To obtain a working model for the Leishmania DL motif recognition, we built a model of an optimal Leishmania ligand (using the peptide in pdb: 6I6H as a guide), and docked this peptide into the AlphaFold2 predicted receptor using HADDOCK^17^ (Figure 2/E). Models from the best energy cluster were merged into a multistate PDB (Supplementary Material) and a representative instance was used to generate Figure 2/D, by superimposing it over the animal receptor-ligand structure (pdb: 6I6H)

### ATG8 structural analysis and LIR motif-containing partner prediction

To build a tree of ATG8/ATG12 family proteins in kinetoplastids, we retrieved individual sample proteins from UniProt, and performed BLAST searches to obtain more representatives of the same family. For reference, we included better-annotated ATG8 and ATG12 protein sequences from various animal, fungal and plant proteomes. The alignment was done using the UniProt interface with default settings (see Supplementary for the resulting tree). These multiple alignments suggested that rooting of the tree problematic, and while leishmanial ATG8A, ATG8B, ATG8C and ATG12 clearly belong to the same family and diverged only in *Leishmaniinae*, deeper relationships are unclear. AlphaFold2 modelled structures of the *Leishmania infantum* ATG8 (A4HYJ2), ATG12 (E9AH00), ATG8C.x (A4HTT6), ATG8B.2 (A4HYA4) and ATG8A.1 (E9AGS4) proteins were superimposed in Pymol and critical amino acids were manually colored (Figure 3/A). The LIR-containing *L. infantum* ATG8 complex model structure was generated by AlphaFold2, and manually optimized in PyMol (Figure 3/B). The same peptide outline was overlain over the first row of structures on Figure 3/A to indicate the LIR binding interface.

Genetic architecture of ATG8/ATG12 family ORFs was analyzed by re-mapping UniProt annotated proteins to the chromosomal models at NCBI. Although shorter, interspersed, unannotated ORFs were also found to align with the ATG8 sequence, we only counted intact, full-length ATG8-family proteins to generate (Figure 3/C). However, all ORFs are listed in Supplementary Table 12.

The ATG8-ATG4 complexes (A4HYJ2 [ATG8] with A4I521 [ATG4.2] and A4HTT6 [ATG8C.x] with A4I8K7 [ATG4.1] respectively) were generated using AlphaFold2 multimer through the Colabfold platform. One representative structure of the 5 output models is shown on Figure 3/D for each complex.

### Performing proteome-wide scan for motifs presented in the paper

The developed LeiSig method to perform signal peptide prediction on Leishmania proteomes is available at https://leishmaniadb.ttk.hu/files/LeiSig.zip

The modified KDEL motif can be searched using the regular expression (also see Supplementary Material for a short script).

### Structure models from proteins analyzed in the manuscript

All structures used for the analysis are available in the Supplementary Structure file.

## Supporting information

Supplementary Structure

