## Supplementary Material for "Exploring Leishmania linear motifs directing protein secretion, sorting and autophagy"

**Content**

| | |
|---|---|
| 2 | Supplementary Figures |
| 5 | Supplementary Tables |
| 5 | Supplementary Structures |
| 6 | Scripts to screen LIR/KDEL motifs defined by regular expression in Leishmania proteomes |
| 6 | Web-servers to screen LIR/KDEL motifs defined by regular expression |
| 7 | KDEL receptors and identification of ER-resident proteins |

**Supplementary Figures**

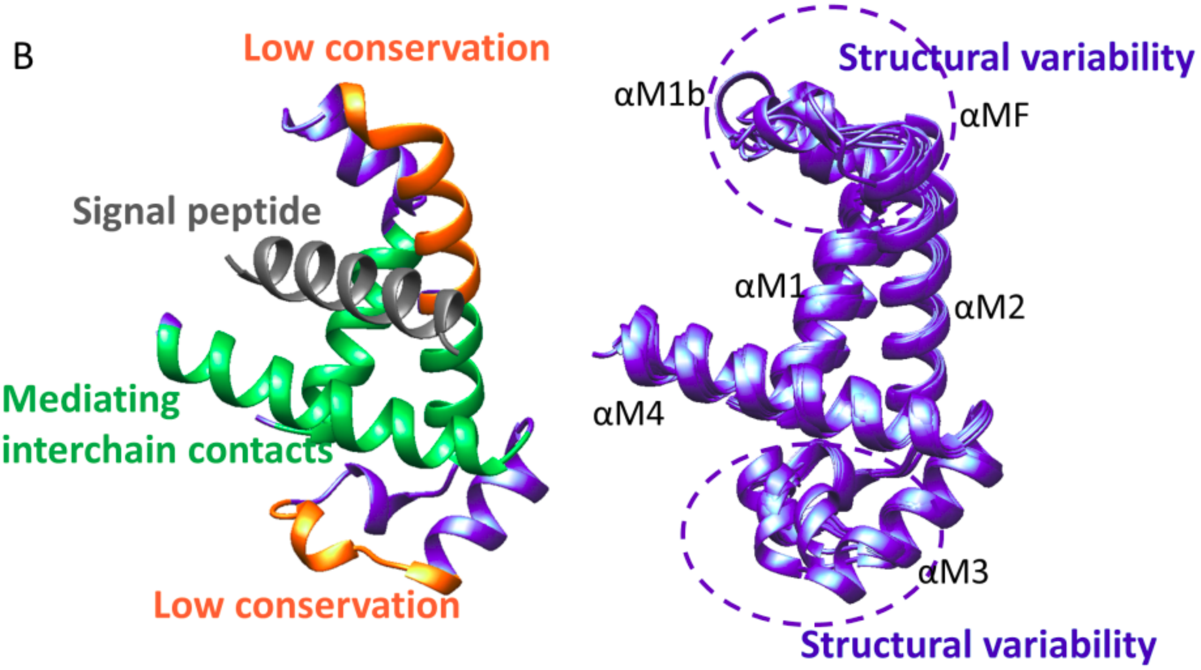


SFigure 1: Left: X-ray structure of *Saccharomyces cerevisiae* SRP54 together with the signal peptide (PDB: 3KL4) - green: SRP54 helices mediating the interaction with the signal peptide; orange: poorly conserved loops; grey: signal peptide. Right: structural superimposition of kinetoplastid SRP54 proteins. Protein segment names were taken from Janda et al, *Nature*, 2010.

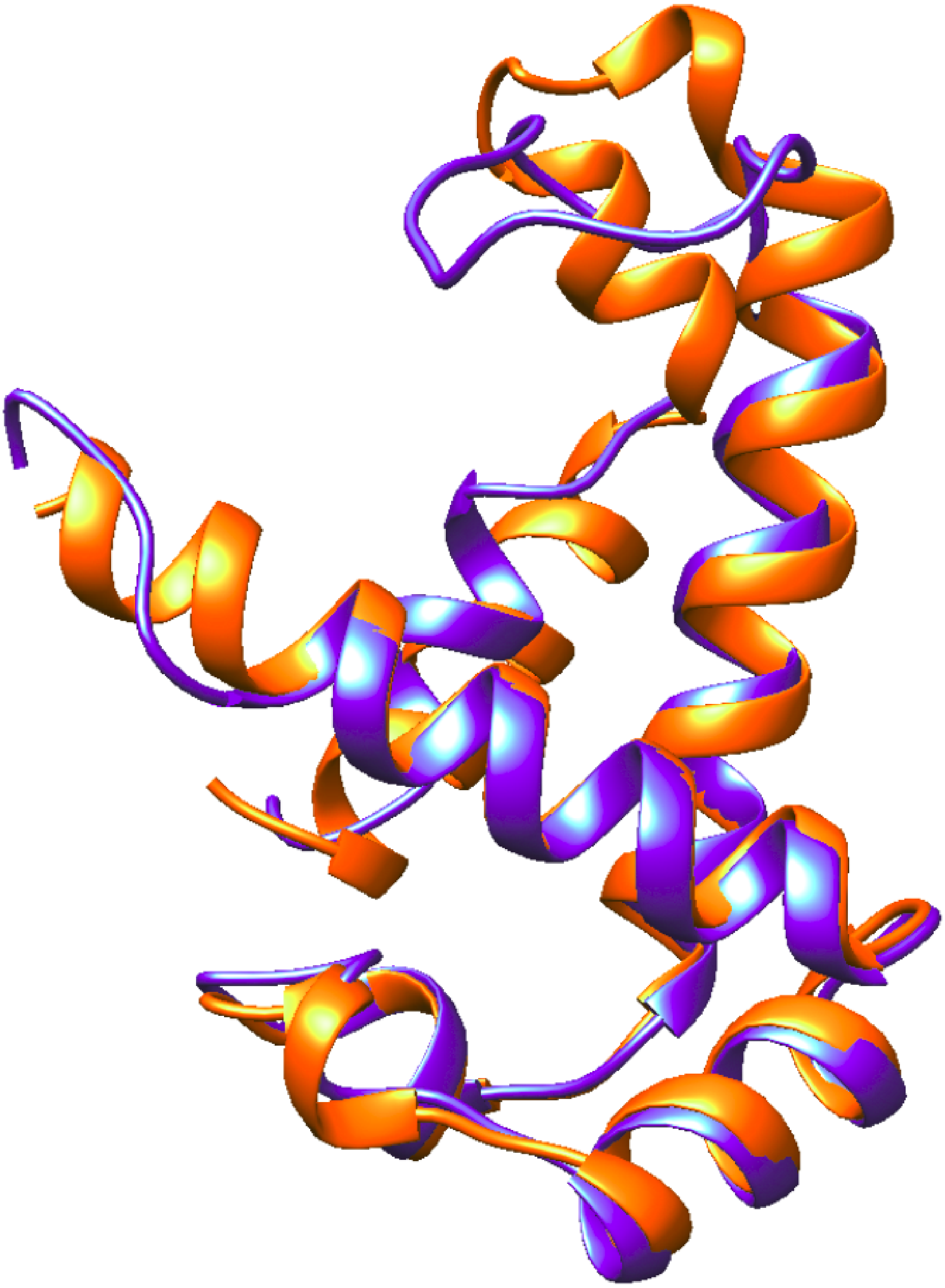

SFigure 2: Superimposed Phyre2 (blue) and AlphaFol2 (orange) model of SRP54 Leishmania infantum

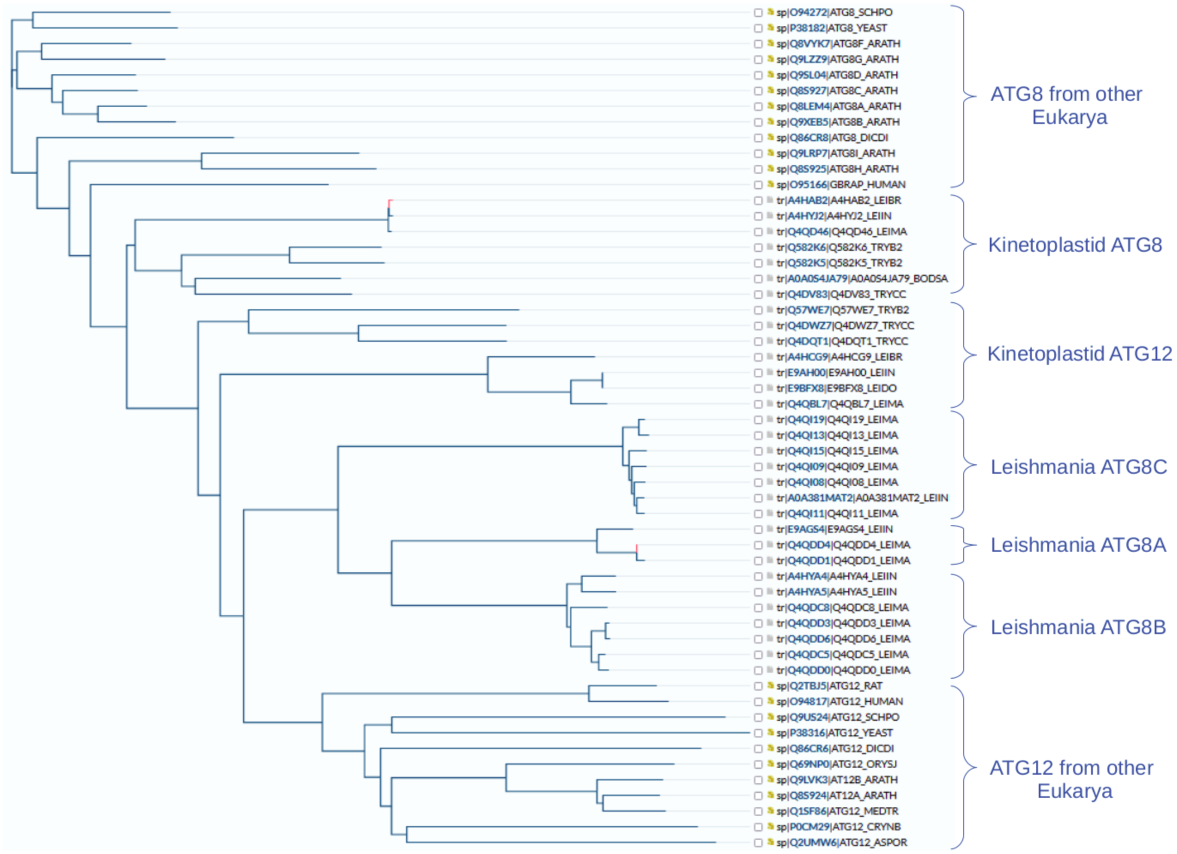


SFigure 3: tree of ATG8/ATG12 family proteins in kinetoplastids.

**Supplementary Structures**

SRP54-signal peptide models :
`signal_*_LEI.pdb`
`signal_*_SC.pdb`
Best energy HADDOCK structures for L infantum KDEL models
`KDEL-best-HADDOCK-run-01.pdb`
AlphaFold2 Multimer outputs for ATG8-ATG4 complexes:
`LeiATG41ATG8C*.pdb`
`LeiATG42ATG8*.pdb`
Cognate idealized LIR peptide modelled onto the L infantum ATG8 protein:
`Lei-ATG8-LIR-best-model.pdb`

**Supplementary Tables**

STable 1: Amino acid distribution in hydrophobic regions of signal peptides
STable 2: Energy calculations for SRP54 + signal peptides
STable 3: signal peptides detected in LeishMANIAdb by SignalP6
STable 4: signal peptides mapped by considering orthologs
STable 5: Training sets for Neural Networks
STable 6: Benchmark set 1
Stable 7: Benchmark set 2
Stable 8: Evaluation of the Neural Network
Stable 9: Positive Prediction result of 5 Leishmania species
Stable 10: Search results for DL$ regular expression (KDEL)
Stable 11: PSSM for modified KDEL motif
Stable 12: ATG8 chr. Locations
Stable 13: ATG8 99% identical sequences
Stable 14: Search results for DE.{0,2}W..[LIM] regular expression (LIR1)
Stable 15: Search results for W..[LIM].[DE] regular expression (LIR2)

### Scripts to screen LIR/KDEL motifs defined by regular expression in Leishmania proteomes

Download Leishmania reference proteome from any website. For example you can use the UniProt FTP site to download the most current reference proteome:

https://ftp.uniprot.org/pub/databases/uniprot/current_release/knowledgebase/reference_proteomes/

Read the supplied fasta file into any kind of data structure using python, then iterate through all sequences. Using the following script, you can retrieve hits:

```
import re

ID=""          #this is a string that contains the protein identifier
sequence=""    #this is a string that contains the amino acid sequence
regex=""       #this is a string that contains the regular expression (considering
               #this paper: KDEL: "DL$" or LIR1:"DE.{0,2}W..[LIM]" or
               #LIR2:"W..[LIM].[DE]"


p = re.compile(regex)
for m in p.finditer(sequence):
       print(ID,m.start(),sequence[m.start():m.end()])
```

### Web-servers to screen LIR/KDEL motifs defined by regular expression

The provided regular expressions can be also used on the http://slim.icr.ac.uk/slimsearch/ website to find potential motif hits in selected proteomes.

**KDEL receptors and identification of ER-resident proteins**

01. Canopy architecture related Saponin B1 (CARSAB)

These proteins are alignment-wise restricted to kinetoplastids (homology is too weak outside this group). No ortholog was identified past B saltans. However, domain architecture analyses (InterPro) betray the presence of a Saponin B domain (often, but not always, involved in lipid transfer). The group is distantly related to Canopy proteins (Conserved ER proteins with KDEL, of unclear function) found in animals and plants. Forced alignment of animal (Human, Drosophila) Canopy proteins show that disulphide bridges are matching perfectly, implicating a conserved architecture (other Saponin B domains often have different disulphide architecture). Thus these two distant families (animal Canopy and kinetoplastid CARSAB) must be evolutionarily related, despite the distance.
Note that the other Saponin B domain bearing ER protein family (SES1/SESSAB) identified in kinetoplastids have different architecture and do not align well. The ultimate proof for the distinct nature of these two families is the presence of both in the same organism (Arabidopsis thaliana has both SES1 and Canopy homologs).

Kinetoplastid

```
tr|A0A0S4JVF1|A0A0S4JVF1_BODSA      ------------------------------------------------------------    0
tr|Q381R4|Q381R4_TRYB2              ------------------------------------------------------------    0
tr|A0A1X0NUB5|A0A1X0NUB5_9TRYP      ------------------------------------------------------------    0
tr|Q4CZ44|Q4CZ44_TRYCC              MFACISVCLYFISLRGMNLCELCFKHRTSFSFFF-AIFFLYYPLFGWASAESSTKIKQKT    59
tr|Q4D5S5|Q4D5S5_TRYCC              MFACISPC--FISLRGMNLCKICFKHRTSFSFSFLPFFFFYYPLFGWASAESSTKTKQKK    58
tr|A0A3R7K0H2|A0A3R7K0H2_9TRYP      ------------------------------------------------------------    0
tr|A0A422NB75|A0A422NB75_TRYRA      ------------------------------------------------------------    0
tr|S9UAW3|S9UAW3_9TRYP              ------------------------------------------------------------    0
tr|A0A0N0DZ25|A0A0N0DZ25_LEPPY      ------------------------------------------------------------    0
tr|A4HKJ0|A4HKJ0_LEIBR              ------------------------------------------------------------    0
tr|Q4Q576|Q4Q576_LEIMA              ------------------------------------------------------------    0
tr|A4I822|A4I822_LEIIN              ------------------------------------------------------------    0


tr|A0A0S4JVF1|A0A0S4JVF1_BODSA      ----------------------------------------------MAVAFSV-------    7
tr|Q381R4|Q381R4_TRYB2              ------------------------MYA-----RLSNTSQSYYSII-----SKIFQLLAII    26
tr|A0A1X0NUB5|A0A1X0NUB5_9TRYP      ------------------------MRH--CFFSSRGINASSLLL---------ILCLLIT    25
tr|Q4CZ44|Q4CZ44_TRYCC              K-------TKENTIKGEILEKLGAMWF--FFLLFSFLNQKFMPF---------VIFLFTL    101
tr|Q4D5S5|Q4D5S5_TRYCC              KKKTKTKTKKENTIKGGILERLEAMWF--FLLLFSFLNRKFMPF---------VIFLFTL    107
tr|A0A3R7K0H2|A0A3R7K0H2_9TRYP      ------------------------MRL--CLPINSPVRQTSLFF---------ALCLFTL    25
tr|A0A422NB75|A0A422NB75_TRYRA      ------------------------------------------------------------    0
tr|S9UAW3|S9UAW3_9TRYP              --------------------------------------------------MKPFVTLLIA    10
tr|A0A0N0DZ25|A0A0N0DZ25_LEPPY      ------------------------MHTFHAKPLHPTLEQRVR-FAAHACALLFLLILLSL    35
tr|A4HKJ0|A4HKJ0_LEIBR              ------------------------MR-APSTPLLAPLQRNQRHFLLAVIAL--LVNLFLA    33
tr|Q4Q576|Q4Q576_LEIMA              ------------------------MR-IHSTFLLAPPQRHQRCCLLASVAL--LASLLLA    33
tr|A4I822|A4I822_LEIIN              ------------------------MQ-RHSTFLLAPPQRHQQCCLLALVAL--LAILLLA    33


tr|A0A0S4JVF1|A0A0S4JVF1_BODSA      -----VFLAFIASALEPL--SSQPQLQCSACLAIAELVGQKMNDTAKL----KTTFQASH    56
tr|Q381R4|Q381R4_TRYB2              GSISLVEAATVGDPLFRIHEPTYAFLDCSACIAFAGHLGRRMNESIRYGGRGGATFLSSH    86
tr|A0A1X0NUB5|A0A1X0NUB5_9TRYP      GDVLSVFAADPNDPRFRIHEPSYAYLDCSACLAFAEYLGRRMNASLEHGG-SGASFLTTH    84
tr|Q4CZ44|Q4CZ44_TRYCC              TDVMSASAAILKDPRFLIHGPSYAYLDCSACLAFAGYLGERMNASLEYDGRGGASFLATH    161
tr|Q4D5S5|Q4D5S5_TRYCC              ADFMSVSAAILKDPRFLIHGPSYAYLDCSACLAFAGYLGQRMNASLEYDGRGGASFLATH    167
tr|A0A3R7K0H2|A0A3R7K0H2_9TRYP      GGVLSGSAATLDDPRFRIHGPSYAYMDCSACIAFAGYLGQRMNASLEYDGRGGATFLSTH    85
tr|A0A422NB75|A0A422NB75_TRYRA      --MLSASAATLGDPRFRIHGPSYAYMDCSACIAFAGYLGQRMNASLEYDGRGGATFLSTH    58
tr|S9UAW3|S9UAW3_9TRYP              ALC--VVAGTAAIDLPQQHPSGYAHMDCSSCLAVAHVLSDKMNKSLLA---SSSSFLSSH    65
tr|A0A0N0DZ25|A0A0N0DZ25_LEPPY      AAPALVATAHTGHSVRPLHGPGYEHLDCSACLTVSRALFVRLNATLSE---SPSTYLASH    92
tr|A4HKJ0|A4HKJ0_LEIBR              SSLTLAATPAQGKSIRPLHGAGYEHLDCSACITVARTLFNRLNKTLAE---NPSTYLVSH    90
tr|Q4Q576|Q4Q576_LEIMA              LPAVLAVTPPQGKPIRPLHGAGYEHLDCSACITVARTLFERLNQTLAE---NPSTYLISH    90
tr|A4I822|A4I822_LEIIN              SPTVLAAPPPQGKPIRPLHGAGYEHLDCSACITVARTLFERLNQTLAE---NPSTYLISH    90
                                                               ::**:*::.:  :  ::* :      :::  :*

tr|A0A0S4JVF1|A0A0S4JVF1_BODSA      RLDARNKVKRIDYESSELRAYEILDGFCNELE-KDYIFRLGEN-NLRLFSSN--------    106
tr|Q381R4|Q381R4_TRYB2              RLGKDNKLKRRDYATSELRGAEVLEKICNDSLFDDYALHFRADMRIRVYDNG--------    138
tr|A0A1X0NUB5|A0A1X0NUB5_9TRYP      RLSETNKLKRQAYATSELRAVEVLEKICNDELEENFVLRLDTDKRIRVYESE--------    136
tr|Q4CZ44|Q4CZ44_TRYCC              RLNKENKLQRNLYATSELRAVEVLEKICSVALEENYVLHLDIDRRIRVYQSG--------    213
tr|Q4D5S5|Q4D5S5_TRYCC              RLNKENKLQRNLYATSELRAVEVLEKICSVTLEENYVLHLDIDRRIRVYQSG--------    219
tr|A0A3R7K0H2|A0A3R7K0H2_9TRYP      RLDAKNKLQRSVYATSELRSVEVLEKICNDALEENYVLRLDLDRRIRVYESE--------    137
tr|A0A422NB75|A0A422NB75_TRYRA      RLNEENKLQRSVYATSELRSVEVLEKICNDALRENYVLRLDLDRRIRVYESE--------    110
tr|S9UAW3|S9UAW3_9TRYP              RLDTANQLKRVQYADSELRAAEVLEDACNKEVYEWYHLRVNPLTKVRGYHFFYYSHIQAV    125
tr|A0A0N0DZ25|A0A0N0DZ25_LEPPY      RLNRVNQLKRRPYRNSELLVTEVMDNFCSSYKNDERTLRLHPKSKVRLYHQQIWKDVTLG    152
```

```
tr|A4HKJ0|A4HKJ0_LEIBR              RLSKQNQLRRRQYRSSELLVSEVMENMCATYQNDNRLLRLHPKSKVRLYHQQIFGDARLG    150
tr|Q4Q576|Q4Q576_LEIMA              RLGKKNQLRRRQYRNSELLVTEVMENVCTSYKNDARLLRLHPKSKVRLYHQQVFGDTRLG    150
tr|A4I822|A4I822_LEIIN              RLGKKNQLRRRQYRNSELLVTEVMENVCTSYKNDARLLRLHPKSKVRLYHQQVFGDTRLG    150
                                    **.  *:::*  *  ***   *:::  *        .   ::.  .:* :

tr|A0A0S4JVF1|A0A0S4JVF1_BODSA      --------------------------KSLTRANYYGKNDKKELRSV--SKRLRDICNEIT    138
tr|Q381R4|Q381R4_TRYB2              -------------------------ESEYPLAQHYSKEDSRILKAV-KTKAVHVFCTRVM    172
tr|A0A1X0NUB5|A0A1X0NUB5_9TRYP      -------------------------KSDLPFAQHYSTQDSKALKET-SKTAVRSFCTRVM    170
tr|Q4CZ44|Q4CZ44_TRYCC              -------------------------NTDFPYAQHYSVQDSDALKVI-SKTAVRDFCTRLM    247
tr|Q4D5S5|Q4D5S5_TRYCC              -------------------------NTDFPYAQHYSVQDSAALKVV-SKTAVRDFCTRLM    253
tr|A0A3R7K0H2|A0A3R7K0H2_9TRYP      -------------------------KSDSPFAQHYSLQDAEALKSV-SKTAVRSFCTRVM    171
tr|A0A422NB75|A0A422NB75_TRYRA      -------------------------KSDSPFAQHYSMQDADALKSP-SKTAVWTFCTRVM    144
tr|S9UAW3|S9UAW3_9TRYP              TRPILLEGVAFPANS---TYEDDRPSKMAAVAQLYSTSEKKKLGPL-HKNAPASLCVQIM    181
tr|A0A0N0DZ25|A0A0N0DZ25_LEPPY      VRHSLREDEVYPEGAHDPQWDDYVELRRYQVAKVYSPKDQGALHGMEMLAATPTMCATLV    212
tr|A4HKJ0|A4HKJ0_LEIBR              VRHALREDEVYPADADPAAWDNKLDGHLYPIARLYSVRDADALHGMQSLSATPTMCALLV    210
tr|Q4Q576|Q4Q576_LEIMA              VRHALREDEVYPADADPAAWAEKLDGQLYPIARLYSRRDADALQGLQSLSATPTMCALLV    210
tr|A4I822|A4I822_LEIIN              VRHALREDEVYPADADSAAWAEKLDGQLYPIARLYSRQDADALQGLQSLSATPTMCALLV    210
                                                                        *. *.  :   *              :*  :

tr|A0A0S4JVF1|A0A0S4JVF1_BODSA      EEDDTKIISLIKKERYLDEFQQKLCFSEGFK-----------------------------    169
tr|Q381R4|Q381R4_TRYB2              DEEEDAMTSLVREVELLSELEQRLCGGYLEM-RNIT---------------APPFPVEKA    216
tr|A0A1X0NUB5|A0A1X0NUB5_9TRYP      GEEEEAMIALVREEQNLTNVEWRLCDGSTDEN---S---------------TRPESLGES    212
tr|Q4CZ44|Q4CZ44_TRYCC              DEEEEAMMNVVREVRELAELELRLCGGNDVNRSSVK---------------TPLPKIEAL    292
tr|Q4D5S5|Q4D5S5_TRYCC              DEEEEAMMNVVREVRELAELELRLCGGNDVNRSSVK---------------TSLPKFETL    298
tr|A0A3R7K0H2|A0A3R7K0H2_9TRYP      DEEEDAMMALVREVRELAELERRLCGGNDVNSSAAT---------------TPPPPLEEL    216
tr|A0A422NB75|A0A422NB75_TRYRA      SEEEDAMMALVREVHDLTELERRLCGGNDVNSSAVT---------------TPPPPLENL    189
tr|S9UAW3|S9UAW3_9TRYP              AEAEEEIEDMVRHAQYQWEVERHLCGLDIGPWWPEE---GEQEEDEYNVRRPRWEGLEEP    238
tr|A0A0N0DZ25|A0A0N0DZ25_LEPPY      EDFEDEIEELVKLAHNLSDIEYGLCGLPLAD--PTDAAASGEAEAVR--------DAIRP    262
tr|A4HKJ0|A4HKJ0_LEIBR              EEAEEEIEELVKTARTQAEVEYRLCGMAFPLNKSFPTLDGETKENEV--------SSAPP    262
tr|Q4Q576|Q4Q576_LEIMA              EEFEDEIESLVKSVRSLEETEHSLCGMTLPVNKPRTVLDGETEGDEV--------SSVPL    262
tr|A4I822|A4I822_LEIIN              EEFEDEIESLVKSVRSLEETEHSLCGMALPVNKPRAALDGETEGDEV--------SSVPL    262
                                    : :  :  :::  .   : :  **

tr|A0A0S4JVF1|A0A0S4JVF1_BODSA      ----CCGTKKVEASKEKERERREKWLELKKVKDEKIKARAEREKRKKEREEEEKKRREES    225
tr|Q381R4|Q381R4_TRYB2              VTAVCVGVEPSIEAEVRRLEKYEKWQASISKGTSQD-KMKGA---EGAES-VVVK-----    266
tr|A0A1X0NUB5|A0A1X0NUB5_9TRYP      ITSVCVGTEASMKAELGRIQRYEKWQEGFTKRREEAIRNKSN---KTEEP-VELK-----    263
tr|Q4CZ44|Q4CZ44_TRYCC              VTAVCIGTEASIAAELGRIQRYEKWQESISRRREEAIKKIGG---KSEEP-IVAK-----    343
tr|Q4D5S5|Q4D5S5_TRYCC              VTAVCIGTEASIAAELGRIQRYEKWQESISRRREEAIKKLGG---KSEEP-IVAK-----    349
tr|A0A3R7K0H2|A0A3R7K0H2_9TRYP      ITAVCIGTEASLAAELGRIHRYEQWQDTFAKRREEAIEKSGG---RSSEP-VVAK-----    267
tr|A0A422NB75|A0A422NB75_TRYRA      VTAVCIGTEASLNAELGRIQRYELWQDSISKRREEALKNIRE---ESDEP-VFAK-----    240
tr|S9UAW3|S9UAW3_9TRYP              VLQVCAASHPIAEDARKDQVLWQRWTDIRERKKQEAAVRHSR---RFLDA-EDE------    288
tr|A0A0N0DZ25|A0A0N0DZ25_LEPPY      ITNICALTEQLREAAHQDQLRWSQYMRREERRKERLEEKEKV---LKRDA-EEPA-----    313
tr|A4HKJ0|A4HKJ0_LEIBR              ITNVCADVEVLRAAARRDQERWEQYQRREAKRKVEIAQRMET---AADVA-GAAT-----    313
tr|Q4Q576|Q4Q576_LEIMA              ITNVCADAEVLRAAARRDQQRWEQYERREAERKAKLAQRRKL---ET-AA-AAAT-----    312
tr|A4I822|A4I822_LEIIN              ITNVCADVEVLRAAARRDQQRWEQYERREAKRKAKLDQRRKL---ET-TA-AAAT-----    312
                                             *   .             . :      .

tr|A0A0S4JVF1|A0A0S4JVF1_BODSA      CSTTVDYLLFHFIPFNLRVEHTW-------------------------------------    248
tr|Q381R4|Q381R4_TRYB2              ---GVQYSKDDPLSFTVLRKPNAKSNGDTTEGGSDAAE----DVRK--------QPDKDS    311
tr|A0A1X0NUB5|A0A1X0NUB5_9TRYP      ---PVNHSAN-PEPFFKMSDLTGGISDD-DDDEE--------------------------    292
tr|Q4CZ44|Q4CZ44_TRYCC              ---EVVHGEKDPQPWFSVSKSTFSSSDS-EEGDDDD-E----GGLE--------------    380
tr|Q4D5S5|Q4D5S5_TRYCC              ---EVVHGEKDPQPWFSVSESTFSSGDR-EEGDDDD-E----GGLE--------------    386
tr|A0A3R7K0H2|A0A3R7K0H2_9TRYP      ---PVEHNANDTKPWVLFPELGADADQD-GDGDGDAGE----GDTG--------------    305
tr|A0A422NB75|A0A422NB75_TRYRA      ---PVEPDDERYKPWSLFPEMDAATGED-GDIDD---S----DDTA--------------    275
tr|S9UAW3|S9UAW3_9TRYP              -----EL-----------------------------------------------------    290
tr|A0A0N0DZ25|A0A0N0DZ25_LEPPY      ------------EKTEVETQEGSSSSGNTGEHNGDKGPAKSEDVVNEGTARSDAAPEAAA    361
tr|A4HKJ0|A4HKJ0_LEIBR              ----SELASEASTNTEVAAALDRSTPAAAGKS------------VDSSTEKATNNP----    353
tr|Q4Q576|Q4Q576_LEIMA              ----SESADEAPANAEAAAALASSAPGASGEP------------IDSPAEEAPYSP----    352
tr|A4I822|A4I822_LEIIN              ----SEPAEEAPTNAEAAAALASSAPGAPGEP------------IDSPTEEAPYSS----    352


tr|A0A0S4JVF1|A0A0S4JVF1_BODSA      ---------------------    248
tr|Q381R4|Q381R4_TRYB2              V--------------FMNFDL    318
tr|A0A1X0NUB5|A0A1X0NUB5_9TRYP      -----------------DMDL    296
tr|Q4CZ44|Q4CZ44_TRYCC              ---------------SIGIDL    386
tr|Q4D5S5|Q4D5S5_TRYCC              ---------------SIGIDL    392
tr|A0A3R7K0H2|A0A3R7K0H2_9TRYP      ---------------ERGMDL    311
tr|A0A422NB75|A0A422NB75_TRYRA      ---------------ERNMDL    281
tr|S9UAW3|S9UAW3_9TRYP              ---------------------    290
tr|A0A0N0DZ25|A0A0N0DZ25_LEPPY      EEAGSAAEDTTPHGDVEEGDL    382
tr|A4HKJ0|A4HKJ0_LEIBR              --VGGG--DKEVSDNYTDGEL    370
tr|Q4Q576|Q4Q576_LEIMA              --VGGV--LKEAAEDRTDGDL    369
tr|A4I822|A4I822_LEIIN              --ADGA--VKEAAEDRTDGDL    369
```

### Animal

```
tr|Q4Q576|Q4Q576_LEIMA     MRIHSTFLLAPPQRHQRCCLLASVALLASLLLALPAVLAVTPPQGKPIRPLHGAGYEHLD    60
tr|A4I822|A4I822_LEIIN     MQRHSTFLLAPPQRHQQCCLLALVALLAILLLASPTVLAAPPPQGKPIRPLHGAGYEHLD    60
sp|Q9BT09|CNPY3_HUMAN      -------MDSMPEPASRCLLLLP--LLLLLLLLLPAPELGPSQAGAE---ENDWVRLPSK    48
sp|Q9Y2B0|CNPY2_HUMAN      ------------------MKGWG--WLALL----LGALLGT---------AWARRSQDLH    27
sp|Q7JXF7|SEELE_DROME      -------------MLTKALILFG--LLA----------LAQ---------GYSFTSREVK    26
                                                      *                                    .

tr|Q4Q576|Q4Q576_LEIMA     CSACITVARTLFERLNQTLAENPSTYLISHRLGKK----NQLRRRQYRNSELLVTEVMEN    116
```

```
tr|A4I822|A4I822_LEIIN     CSACITVARTLFERLNQTLAENPSTYLISHRLGKK----NQLRRRQYRNSELLVTEVMEN    116
sp|Q9BT09|CNPY3_HUMAN      CEVCKYVAVELKSAFEETG---KTKEVIGTGYGILDQ---KASGVKYTKSDLRLIEVTET    102
sp|Q9Y2B0|CNPY2_HUMAN      CGACRALVDELEWEIAQVD---PKKTIQMGSFRINPDGSQSVVEVPYARSEAHLTELLEE    84
sp|Q7JXF7|SEELE_DROME      CHVCKAVVTELEEAIAKED---PHKMADVSGFRLDAQGNSISKKVRLVKSEMFLTELMEK    83
                           * .*  :.  *   : :         .                          .*:  : *: *

tr|Q4Q576|Q4Q576_LEIMA     VCTSYKNDARLLRLHPKSKVRLYHQQVFGDTRLGVRHALREDEVYPADADPAAWAEKLDG    176
tr|A4I822|A4I822_LEIIN     VCTSYKNDARLLRLHPKSKVRLYHQQVFGDTRLGVRHALREDEVYPADADSAAWAEKLDG    176
sp|Q9BT09|CNPY3_HUMAN      ICKRLLDYSLHKERT---GSNRFAKGM-SETFETLHNLVHKGVKVVMDIPYELWNET---    155
sp|Q9Y2B0|CNPY2_HUMAN      ICDRMKEYGEQIDPS---THRK----------NYVRVVGRNGESS--ELDLQGIRID---    126
sp|Q7JXF7|SEELE_DROME      ICEKMDDYLKATYKS---NGKF----------TLLKMIINGQMNP--DSSLVDFVQD---    125
                           :*       :                 .                ::   .           :

tr|Q4Q576|Q4Q576_LEIMA     QLYPIARLYSRRDADALQGLQSLSATPTMCALLVEEFEDEIESLVKSVRSLEETEHSLCG    236
tr|A4I822|A4I822_LEIIN     QLYPIARLYSRQDADALQGLQSLSATPTMCALLVEEFEDEIESLVKSVRSLEETEHSLCG    236
sp|Q9BT09|CNPY3_HUMAN      -------------S------AEVADLKKQCDVLVEEFEEVIEDWYRNH-QEEDLTEFLCA    195
sp|Q9Y2B0|CNPY2_HUMAN      -------------S------DISGTLKFACESIVEEYEDELIEFFSRE--ADNVKDKLCS    165
sp|Q7JXF7|SEELE_DROME      -------------G------DLNKSLGHFCNEVLEDNDEIFVKAFQAEELGNDLDIKICS    166
                                             .                 *  ::*: :: : .            ::       :*.

tr|Q4Q576|Q4Q576_LEIMA     MTLPVNKPRTVLDG-ETEGDEVSSVPLITNVCADAEVLRAAARRDQQRWEQYERREAERK    295
tr|A4I822|A4I822_LEIIN     MALPVNKPRAALDG-ETEGDEVSSVPLITNVCADVEVLRAAARRDQQRWEQYERREAKRK    295
sp|Q9BT09|CNPY3_HUMAN      NHVLKGKDTSCLA-EQWSGKKGDTAALGGKK----------SKKKSSR----AKAAGGRS    240
sp|Q9Y2B0|CNPY2_HUMAN      KRTDLCDHALHISHDEL-------------------------------------------    182
sp|Q7JXF7|SEELE_DROME      EQASYCDESPVQEEYDFDGKEEL-------------------------------------    189
                                    .        :

tr|Q4Q576|Q4Q576_LEIMA     AKLAQRRKLETAAAAATSESADEAPANAEAAAALASSAPGASGEPIDSPAEEAPYSPVGG    355
tr|A4I822|A4I822_LEIIN     AKLDQRRKLETTAAAATSEPAEEAPTNAEAAAALASSAPGAPGEPIDSPTEEAPYSSADG    355
sp|Q9BT09|CNPY3_HUMAN      SSSKQRKELGGLE-------GDPSPEEDE-------------GIQKASPLTHSPPDEL--    278
sp|Q9Y2B0|CNPY2_HUMAN      ------------------------------------------------------------    182
sp|Q7JXF7|SEELE_DROME      ------------------------------------------------------------    189


tr|Q4Q576|Q4Q576_LEIMA     VLKEAAEDRTDGDL    369
tr|A4I822|A4I822_LEIIN     AVKEAAEDRTDGDL    369
sp|Q9BT09|CNPY3_HUMAN      --------------    278
sp|Q9Y2B0|CNPY2_HUMAN      --------------    182
sp|Q7JXF7|SEELE_DROME      --------------    189
```

### 02. Sensitive to salt 1 / Saponin B (SES1 / SESSAB)

This obscure group does have homologs beyond Kinetoplastida: Many related sequences can be retrieved from green algae and higher plants (including A thaliana as well). All these homologs have a well-visible KDEL signal on them. Interestingly, a few animals also possess such proteins, but only among cnidarians, e.g. Hydra vulgaris (they might also have been acquired it from algae by horizontal gene transfer, as the gene is otherwise lacking from all other animals or even fungi) Seredipitiously, the Arabidopsis thaliana ortholog has already been studied experimentally (PMID:30287478). Because of the phenotype of the mutant, the gene was baptized SENSITIVE TO SALT 1 (SES1). Despite the presence of a Saponin B domain, the SES1/SESSAB proteins are rather different from Canopy/CARSAB family members. Both are suggested to be ER-localized chaperones serving different roles.

```
tr|S9W4D1|S9W4D1_9TRYP             -----------------------MKHLIHLLALCVGFLFLVCDVSGKKDLMLPT--PLPS    35
tr|S9UQQ8|S9UQQ8_9TRYP             -----------------------MFTPI----VLLLLLFSHGVASDGVEVIVPT--SLSP    31
tr|A0A0N0VFN2|A0A0N0VFN2_LEPPY     ---------------MHTMQLRVATPLLLVLCV-LVSSVGLERATARKELPLNT--AFPP    42
tr|A4HQ81|A4HQ81_LEIBR             -------MPCNPRSYVSSTQPLVTAALLLLVCV-LAVATG--CASARKELPLNI--PYPP    48
tr|Q4Q0J6|Q4Q0J6_LEIMA             ------MPCCSPPSHVAPTRPLVTTTLLLLVCV-LAAALG--CASARKQLPLNT--PYPP    49
tr|A4IDX9|A4IDX9_LEIIN             ------MPCCSPPSHVAPTRPLVTSALLLLVCV-LAAALG--CASARKQLPLNT--PYPP    49
tr|A0A0S4IVP5|A0A0S4IVP5_BODSA     ------------------M-----STCVWVIGV-VLWLFVYHAVSAKKEKLTFN--SYGP    34
tr|Q38A59|Q38A59_TRYB2             -------------MGKAPV-----KG--VFLVL-VALVPMFPAVDAKKEKPMNV--PFPQ    37
tr|A0A1X0P0S9|A0A1X0P0S9_9TRYP     ------------------M-----RSLMYFVLL-ISAATLLTLTNAKKEKPLNE--PFPK    34
tr|Q4CTW1|Q4CTW1_TRYCC             MDRIKVEMKSKQTREGRPM-----RFIKLCIFF-AVTLLFSAAGVAKKEKPPHA--PFPK    52
tr|Q4CWS6|Q4CWS6_TRYCC             MDRIKVEMKSKQTREGRPM-----RFLKLCIFF-AVTLLFSEAGVAKKEKPPHA--PLPK    52
tr|A0A422PSY9|A0A422PSY9_9TRYP     ------------------M-----RTFVVSLFI-LVTLLLSSAIDARKEKPLHD--AFPK    34
tr|A0A3R7NVI0|A0A3R7NVI0_TRYRA     ------------MRKDHQM-----RTFIITVCI-LVVLLFSSAVAAKKEKPLHI--PFPK    40
tr|A0A8J4C7T8|A0A8J4C7T8_9CHLO     --------------HFHIC----MMVRLLYV-LLFAFIAVASAQFGGPKRPKTK-APVNR    40
tr|A8J2L8|A8J2L8_CHLRE             ------------------M----ARLNLLQLCVLGLCLAVASAQFGGKRPPKPK-APVAK    37
tr|A0A150GFK4|A0A150GFK4_GONPE     ------------------M----KRNNLLRLLLLALV-VAASAQFGGPKRPKKK-SPVAK    36
```

```
tr|Q5VML4|Q5VML4_ORYSJ              ---------MARGGGVAVA----MAVAVAAVV-----LLLHPAASAAAAGPKKVATAARK    42
tr|A0A8B6XU83|A0A8B6XU83_HYDVU      ------------------M----IGIEICTAY-----LLIHFG-FGISQMPNKSPILPIV    32

tr|S9W4D1|S9W4D1_9TRYP              EVQAILNCDLCAYTLINAVLRVHEVQDALEAQELRLKEDDILS-VLETLCQPYTEYGQWM    94
tr|S9UQQ8|S9UQQ8_9TRYP              VVSEAFACEICGFTVMYALMAVSEKTAELEKADLVINEDDILS-LFEDICNPFMEKGEWI    90
tr|A0A0N0VFN2|A0A0N0VFN2_LEPPY      VVQDALRCDVCAFIVANSLNHVEAKRDELRAKRLALREDDVLE-ETENLCIPFKDQGQWI    101
tr|A4HQ81|A4HQ81_LEIBR              IVQNALRCNVCSFIVDNALIYVQGMRDEMSQHRLQLREDDVLD-ELEKLCVPFKDPGQWI    107
tr|Q4Q0J6|Q4Q0J6_LEIMA              VVQNALRCDVCSFIVVNSLYQIEAKREEQQRKRLQVREDDVLE-EVENMCVPFKDQGQWI    108
tr|A4IDX9|A4IDX9_LEIIN              VVQDALRCDVCSFIVVNALYQVEAKREEQQRKRLQVREDDVLE-EVENMCVPFKDQGQWI    108
tr|A0A0S4IVP5|A0A0S4IVP5_BODSA      AVERRHSCEICRTLVVEAYPIAEELHKNASRRGYALREEEVLSKVTEAICNPFVKAGQWI    94
tr|Q38A59|Q38A59_TRYB2              DAIDDVRCDVCRLMAKRAYGYVYELFVTSEHTRIPVNEENVLV-AIEDICNPLAASGQWI    96
tr|A0A1X0P0S9|A0A1X0P0S9_9TRYP      DAIDAISCDICSLLVRKTYLDVQALFAASVETRVRVNEDDILT-AIEDVCNPFAETGQWI    93
tr|Q4CTW1|Q4CTW1_TRYCC              DVVNAISCGVCNFMVGKAYGDVQLLVNASVQTRVRVNEDDVLT-AIEDLCNPFSDVGQWI    111
tr|Q4CWS6|Q4CWS6_TRYCC              DVVDAISCGVCNFMVGKAYGDVQLLFNASVQTRVRVNEDDVLT-AIEDVCNPFSDVGQWI    111
tr|A0A422PSY9|A0A422PSY9_9TRYP      EVIDGIRCGVCNFAVKQAYGNVLGLFNASIQTRVRMNEEDVLT-VLEDICNPFTEVGQWI    93
tr|A0A3R7NVI0|A0A3R7NVI0_TRYRA      DAVDGISCGVCTFVVKQVYRDVLVLFNASIRRRVRMSEDDVLT-ALEDVCNPFAETGQWI    99
tr|A0A8J4C7T8|A0A8J4C7T8_9CHLO      DDIKYIKCKVCEAMAKEARN-VVKELGEL-AGAKKINEADILE-RLEKMCNPDLNEGDWI    97
tr|A8J2L8|A8J2L8_CHLRE              GDIKYIKCQVCEAIAKQSIK-ATKDLIEQ-AGPKKVPEADILE-RVEKMCDPDTDDGEWI    94
tr|A0A150GFK4|A0A150GFK4_GONPE      DDIKYIKCQVCEAIAKQARN-IKKDLQAV-AGPKKVSEGDLLD-KFEKMCDPEQDDGDWI    93
tr|Q5VML4|Q5VML4_ORYSJ              EDIPYIRCQVCERIAREISAQVAKKQQAL-PATKKVPEIEIIE-IAENVCNLKKQEADWM    100
tr|A0A8B6XU83|A0A8B6XU83_HYDVU      EDLPFIKCDVCQKAAKVLFKTIEN----Q-RSEKKLDEDDVLS-IVEKSCDTDVLYGDWI    86
                                             * :*                       : * :::  *  *     .:*:

tr|S9W4D1|S9W4D1_9TRYP              RRVSIQLQADTHS---------EGKEKMLQTTVLPYVSTCKRTCSTTQAVCEQVLDSSEM    145
tr|S9UQQ8|S9UQQ8_9TRYP              RRAELTLDEKKR---------------TVSFNIGNNYTKCKRFCGTFSNVCEQVYDSDEA    135
tr|A0A0N0VFN2|A0A0N0VFN2_LEPPY      RQVSLQVAPKSS--------QDASEGHQMEVGVVNFYGRCGRTCDTVAALCEELMDRDVM    153
tr|A4HQ81|A4HQ81_LEIBR              RQVSLRVDETKVC-RNADTSVIEAPRQVMRVALVEYYSKCGRTCDTVALLCEEWMDSSYM    166
tr|Q4Q0J6|Q4Q0J6_LEIMA              RQVALQVEEVAPSERGTDRPAAATPRHHMSVGVVDYYSKCGRICETVAVLCEEWMDSGYM    168
tr|A4IDX9|A4IDX9_LEIIN              RQVALQMEEVAPSKRGADRPAAATPRYHMSVGVVDYYSKCGRICETVAVLCEEWMDSGYM    168
tr|A0A0S4IVP5|A0A0S4IVP5_BODSA      RLVHIKANE---------------KTLQTVLEPLEFYSQCKRDCTTVANSCSSVVDTDYG    139
tr|Q38A59|Q38A59_TRYB2              RQVAVDTSSP----------AAAKGPFHITLTELPHHSKCKRTCKLSDACEAVVDDDDM    146
tr|A0A1X0P0S9|A0A1X0P0S9_9TRYP      RRVAINYSA-------------MTAPFLLGIEELPVFTKCKRTCTTIVEACEAIMDHDSM    140
tr|Q4CTW1|Q4CTW1_TRYCC              RQITITHES-------------DTAPFLLGVSRLPAYAKCKRVCSTVVEACEAVMDHENM    158
tr|Q4CWS6|Q4CWS6_TRYCC              RQITITHES-------------DTAPFLLGVSRLPAYAKCKRVCSTVVEACEAVMDHKNM    158
tr|A0A422PSY9|A0A422PSY9_9TRYP      RRITITHRR-------------DTDHL-LGVEELPVYTKCKRTCSTVVEACEAVMDHESM    139
tr|A0A3R7NVI0|A0A3R7NVI0_TRYRA      RRITITHKR-------------ETAPF-LGVEELQVYTKCKRTCSTVVEACEGVLDHESM    145
tr|A0A8J4C7T8|A0A8J4C7T8_9CHLO      AMYDIVKEGD-----------------ILNLKDTGMVGRCKSKCRTIARTCELIAEDLDL    140
tr|A8J2L8|A8J2L8_CHLRE              TKYDIVEEDS-----------------ALVLKDTGAVGRCKSECRTIARACEMISDDVDL    137
tr|A0A150GFK4|A0A150GFK4_GONPE      TQFDIVEEGS-----------------ALKLKDTGMMGKCKSECRTIAYACEKILDDVDL    136
tr|Q5VML4|Q5VML4_ORYSJ              LKIDIVEKGD-----------------KLELVEQDEEGHCNAECKTIERACQEVMGYAD-    142
tr|A0A8B6XU83|A0A8B6XU83_HYDVU      SRLDIVEKEN-----------------DLKVVEHPQEGKCDRECKTISRSCEDIIGDID-    128
                                             :                                   *   * *  *.

tr|S9W4D1|S9W4D1_9TRYP              DDFSAAVLKLSQRHKTIAEPSVWEPLITQTCHQM-DMCLDATNKRNAFIRQYRKDPAVYQ    204
tr|S9UQQ8|S9UQQ8_9TRYP              TNLAADMLKLIKKRVGLVKTNEAEEIVNKVCKK--EFCEHKERNMAALHQMLAKDDNLWG    193
tr|A0A0N0VFN2|A0A0N0VFN2_LEPPY      DDFPGQLLKVSK-DANMADEAHRDAVFHQFCYAS-TYCKLHQKYVVALEKELRTNVNL-L    210
tr|A4HQ81|A4HQ81_LEIBR              DDFPSYLVREANSGYNISSEEHRSAVFSQFCAPT-PHCENIGSALRELSEELTKNSEL-R    224
tr|Q4Q0J6|Q4Q0J6_LEIMA              DGFSSRLVKEANAGRSISDAPHRDAVFDSFCAPS-PHCKKHAIFVRELDLALTTDTEL-R    226
tr|A4IDX9|A4IDX9_LEIIN              DGFSSRLVKEAKAGRNISDAAHRDVVFGSFCAPS-PHCKKHAIFVRKLDMALTKDTEL-R    226
tr|A0A0S4IVP5|A0A0S4IVP5_BODSA      DDLPGLLLKLKPV----------EVIDEAVCKRI-CIDTTAV---RTKPLSA----KM-I    180
tr|Q38A59|Q38A59_TRYB2              DQFSSKLLILKKY-------KDASALADATCGVS-AFCTSR------TGFSSKRYKEL-L    191
tr|A0A1X0P0S9|A0A1X0P0S9_9TRYP      DQLTPKLLHLPEY-------ADGETLARSLCQTS-PICTKR------KSLSAERYAEL-R    185
tr|Q4CTW1|Q4CTW1_TRYCC              DQLSPRLLRLNEY-------AGAEAFANALCLPS-SICTER------QGLSAARYEEV-R    203
tr|Q4CWS6|Q4CWS6_TRYCC              DQLSPRLLRLNEY-------AGAEAFANALCQSS-SICTER------QGLSAARYEEV-R    203
tr|A0A422PSY9|A0A422PSY9_9TRYP      DTLSPRLLHLTEY-------ADAGTFAQALCGPS-PICTKR------RRLLADRYDEL-V    184
tr|A0A3R7NVI0|A0A3R7NVI0_TRYRA      DMLSPRLLHLTEY-------ADADKFAEALCDRS-PICTKR------WGLTASRYDEL-T    190
tr|A0A8J4C7T8|A0A8J4C7T8_9CHLO      TDLSAMLFKGK-K---------RSAVTNWMCYDATDVCISKPPP-------------VPA    177
tr|A8J2L8|A8J2L8_CHLRE              TDLSAMLFKGK-K---------RAAISNWMCHDASDACSKKAPA-------------LTK    174
tr|A0A150GFK4|A0A150GFK4_GONPE      TELSSLVYAGK-K---------RAAVTNWLCHESSDACESKPPP-------------VPK    173
tr|Q5VML4|Q5VML4_ORYSJ              TDVAEFVYKKKPS---------ADQLVKFLCKDLSEACVVDPPP-------------VPK    180
tr|A0A8B6XU83|A0A8B6XU83_HYDVU      TDIGELLWKNEMK---------LATFINHVCYKLTNSCKAKKKY-------------VKG    166
                                    .   :                      .        *                         :

tr|S9W4D1|S9W4D1_9TRYP              KLIEDEAPAKLDEDEIEVEQMMDNFNRVD-----GKKTSVFSREEMVELQSALLRGDKKA    259
tr|S9UQQ8|S9UQQ8_9TRYP              TLVEDKS-EPADHDRIEVENLIAKME-RD-----GKRSNVYSRDHVKKLQAAVKQGDRRA    246
tr|A0A0N0VFN2|A0A0N0VFN2_LEPPY      EEMKADKPRRMKKEELEMETMMYRLMREQ-----RQSADVFSRDEIRKMQQAFIHGTKED    265
tr|A4HQ81|A4HQ81_LEIBR              EAIDADRPQEIKSEEREMEMMLHRLTREH-----GQSADVFSRDEVHRMKEAFLKGNREE    279
tr|Q4Q0J6|Q4Q0J6_LEIMA              ASIDADRPQEIETEEREMETMLHRLTREQ-----GQSADVFSRDEIRRMKEAFVTGNKED    281
tr|A4IDX9|A4IDX9_LEIIN              ESIDADRPQEIETEEREMETMLHRLTREQ-----GQSADVFSRDEIRRMKEAFVTGNKED    281
tr|A0A0S4IVP5|A0A0S4IVP5_BODSA      KEIAAEVPVEIDKKNLDIEEIMDDMERARGGYGGAPKMDVFSRDEMMDVQRAILERDGSK    240
tr|Q38A59|Q38A59_TRYB2              TLVASDVVEVIEQKEMDIERFMDEMERKG-----NRRQEIYSREEVIKMQKSLIEGDIET    246
tr|A0A1X0P0S9|A0A1X0P0S9_9TRYP      TMIKADVMEEIDPKEMEVERMMDQMERKE-----NRRHDIFSREEITAMQQGLLRGDREA    240
tr|Q4CTW1|Q4CTW1_TRYCC              NMIDEDTVEEIDPKEMEIERMMDHMERKE-----NRRQTIFSRDEITSMQKAFLRGDKEA    258
tr|Q4CWS6|Q4CWS6_TRYCC              NMIDEDTVEEIDPKEMEIERMMDHMERKE-----NRRQTIFSRDEITSMQKAFLRGDKEA    258
tr|A0A422PSY9|A0A422PSY9_9TRYP      TMMDADPMEEIDPKEMEVERMMDHMERKE-----NRRQTIFSREEITKMQEAFLRGDKEA    239
tr|A0A3R7NVI0|A0A3R7NVI0_TRYRA      TMIDEDTMEEIDPKEMEVERMMDHMERKE-----NRRHSIFYRDEIVKMQEAILRGDKEA    245
tr|A0A8J4C7T8|A0A8J4C7T8_9CHLO      DRPVGEPHEPLDEDEYRNIMMMRDMEAIG----V--SGSLYSRDTLTEELEEMQDMYGDD    231
tr|A8J2L8|A8J2L8_CHLRE              REGADEAHAPMDEDELRTERMMRSMKAAG----L--SGTMYNKETMQEELAEMADQYEDN    228
tr|A0A150GFK4|A0A150GFK4_GONPE      GRRADERHVPMSEDEVRTEKMLRNMKAAG----L--SGSMYSRETLEEELAEMAGEYSDD    227
tr|Q5VML4|Q5VML4_ORYSJ              DRVPGEPFAAKPSKDAEMDRILKSMEGIP----GAPSMKMYSRDDLMK--NNFGVDGDDD    234
tr|A0A8B6XU83|A0A8B6XU83_HYDVU      S-HKDYKFVEMSEKEKQARDLMRNMKSVP----GMPGMEMYSQDDLQNMREQMGVRNEDE    221
                                                      .        ::  :             :: :: :          .

tr|S9W4D1|S9W4D1_9TRYP              AARIDPSIVDLSDEEFESIQNLAAKQGYVRGGRG-DGEKEGLFHMTDDES----------    308
tr|S9UQQ8|S9UQQ8_9TRYP              ATKLDPTIRSLSDEEFEEVVRMAREEYNTDT-L-----EEDL------------------    282
tr|A0A0N0VFN2|A0A0N0VFN2_LEPPY      VAAVDPKAFDLSDEEFVALREHMRADERVTGGNR-RGQKQEGDED-GD------------    311
tr|A4HQ81|A4HQ81_LEIBR              LKAIDPAAFDLTDSEFSMLQNYIRGEAPEEKEWE-QQRGSSNPSDAED------------    326
```

```
tr|Q4Q0J6|Q4Q0J6_LEIMA                 LQAVDPTAFDLTDDEFSTLKDYMHGEERKQQQQQ-RQRRPSAPSDADD------------    328
tr|A4IDX9|A4IDX9_LEIIN                 LQAVDPTAFDLTDDEFSTLQDYMRGEERKQQ--Q-RQRRPSAPSDADD------------    326
tr|A0A0S4IVP5|A0A0S4IVP5_BODSA         LRDLDPSAEDLSDEELEYLQRMYAGEGDGGEEYP-TELGEDL------------------    281
tr|Q38A59|Q38A59_TRYB2                 AAGIDPSLNKLTEEEQLALQQMLKNKPMGKKRPP-EDAGQAASPEGDS-AEVNPDL----    300
tr|A0A1X0P0S9|A0A1X0P0S9_9TRYP         VAKIDPSIMDLNNEEFAALQAMIRGKMPASQEKV-TGQQQGQSSDNMNSIGMDQDIYNGA    299
tr|Q4CTW1|Q4CTW1_TRYCC                 VAQVDPSIMDLSDEEFSALQSMIRGRAMGQEQPE-ENGDDGEWKGRNK----NYPMYHSA    313
tr|Q4CWS6|Q4CWS6_TRYCC                 VAQVDPSIMDLSDEDFSALQSMIRGRAMGQEQPE-ENGDDGEWKGRNK----NYPMYHST    313
tr|A0A422PSY9|A0A422PSY9_9TRYP         VAKVDPSIMDLSEDEFAAVQAMMRGRKDGKPQAG-ETDADGEAKVRRT----QRRKRRES    294
tr|A0A3R7NVI0|A0A3R7NVI0_TRYRA         VAKVDPSIADLSEEEFAAVQAMVRGKNNEKLRSG-THRADDEGKARDT----QRRKHRES    300
tr|A0A8J4C7T8|A0A8J4C7T8_9CHLO         P-------------DFAQV---MKDTGMDSFASRKPDEEPTSGETAVDSST---------    266
tr|A8J2L8|A8J2L8_CHLRE                 E-------------DFARM---LEETGLDKHMPSRPGSDAQEPDSGVGGG--------AA    264
tr|A0A150GFK4|A0A150GFK4_GONPE         P-------------DFAKA---MADSGLDKHLQKGTEEEPTAADATAEQPSVASSVANAA    271
tr|Q5VML4|Q5VML4_ORYSJ                 D-----D-DEDEDDDFPKN---LGKVFKDK------------------------------    255
tr|A0A8B6XU83|A0A8B6XU83_HYDVU         N-----Q-QD---SEQSEN---YDEVYRTK------------------------------    239
                                                         :

tr|S9W4D1|S9W4D1_9TRYP                 ------------------------------------------------------DL    310
tr|S9UQQ8|S9UQQ8_9TRYP                 --------------------------------------------------------    282
tr|A0A0N0VFN2|A0A0N0VFN2_LEPPY         -----------------L--------------------------------------    312
tr|A4HQ81|A4HQ81_LEIBR                 -----------------L--------------------------------------    327
tr|Q4Q0J6|Q4Q0J6_LEIMA                 -----------------L--------------------------------------    329
tr|A4IDX9|A4IDX9_LEIIN                 -----------------L--------------------------------------    327
tr|A0A0S4IVP5|A0A0S4IVP5_BODSA         --------------------------------------------------------    281
tr|Q38A59|Q38A59_TRYB2                 --------------------------------------------------------    300
tr|A0A1X0P0S9|A0A1X0P0S9_9TRYP         DYGNDE----------DSM-------------------------------IMDDDL    314
tr|Q4CTW1|Q4CTW1_TRYCC                 DNGNDDGE--------NLN-------------------------------LEDEDF    330
tr|Q4CWS6|Q4CWS6_TRYCC                 DNGNDDGG--------NLN-------------------------------LEDEDF    330
tr|A0A422PSY9|A0A422PSY9_9TRYP         GGGEDDWE--------DVG-------------------------------TGESDL    311
tr|A0A3R7NVI0|A0A3R7NVI0_TRYRA         DGGEGDLE--------NFE-------------------------------LEQEDL    317
tr|A0A8J4C7T8|A0A8J4C7T8_9CHLO         ---MTKLQETAAKAAGDIKDGA---AKLVEGAKKFMGKLFGGSKQDG--KSGEL--    312
tr|A8J2L8|A8J2L8_CHLRE                 AGAVADAAARLQDTLTQVKEGA---SKLFDSAKGLVGKWFGKKEAAK--KPAGEL-    314
tr|A0A150GFK4|A0A150GFK4_GONPE         AGAASSLTEAASKAAESIKEGA---AKVVEGAKNVVGKLFGGGGKQE--EAGGNEL    322
tr|Q5VML4|Q5VML4_ORYSJ                 GSPKKDLKQQ---VVKQIKDTGKKLKGHVNKVSKVVKKWWQGKKKPS--KSSKTEL    306
tr|A0A8B6XU83|A0A8B6XU83_HYDVU         LGFFET-------------------------LKFFLNDLWKKIKYLFSYISYKKEL    270
```

### 03. Chitinase with ER retention signal

These proteins are found in kinetoplastids with ER retention (KDEL) signals, and have a lot of homologs across other eukaryotes. However, most multicellular organisms (animals, plants, fungi) have secreted chitinases without any retention signal. The fact that chitinases from some green algae and other plant-like unicellular eukaryotes do have KDEL signal, shows that this feature is not a kinetoplastid innovation. Nevertheless, the role of such a glycosidase in the ER is unclear (externally digested, e.g. insect chitin is unlikely to reach the ER), but it likely processes endogenous glycans. Technically, these glycosidases can also be called Glycohydrolase-18 (PFAM/Interpro label).

```
tr|A0A0S4J833|A0A0S4J833_BODSA         ------------------------------------------------------------    0
tr|A0A250X009|A0A250X009_9CHLO         ------------------------------------------------------------    0
tr|C5K6G1|C5K6G1_PERM5                 ------------------------------------------------------------    0
tr|C1N616|C1N616_MICPC                 ---------------------------------------------------------MAS    3
tr|A0A7G2CNI0|A0A7G2CNI0_9TRYP         ------------------------------------------------------------    0
tr|S9TUR0|S9TUR0_9TRYP                 --------------------MTFVSLSTTT----------QG-ILYRSTLFSFLFNFLCI    29
tr|A0A0M9FXG9|A0A0M9FXG9_LEPPY         ---------------------------------------------------------MCP    3
tr|A4H8K3|A4H8K3_LEIBR                 ---------------------------------------------------------MEP    3
tr|A4HWX6|A4HWX6_LEIIN                 ---------------------------------------------------------MVQ    3
tr|Q4QEU0|Q4QEU0_LEIMA                 ---------------------------------------------------------MVQ    3
sp|O81862-2|CHIC_ARATH                 ------------------------------------------------------------    0
tr|S8AUV1|S8AUV1_PENO1                 ------------------------------------------------------------    0
tr|A0A179UX46|A0A179UX46_BLAGS         MSTYTVASGDSMWAISVARGISLDALIAANPQVSVPSQIEVGQVLNIPGGDAPADPFPAP    60
tr|M9NG70|M9NG70_DROME                 -----------------------------------------------------MLPFR--    5
tr|Q22468|Q22468_CAEEL                 ---------------------------------------------------------M--    1


tr|A0A0S4J833|A0A0S4J833_BODSA         ----MATVETIALVTITVLLLLSASTTPVT-----------------------AETTPEA    33
tr|A0A250X009|A0A250X009_9CHLO         ------------------------------------------------------------    0
tr|C5K6G1|C5K6G1_PERM5                 -------MARLACLL-LLLLS--A--------------------------F---TVIADE    21
tr|C1N616|C1N616_MICPC                 RGGGAGATARLALLL-VVLLLLPGRGAPDTAP---------------SSSFASPSPGAPA    47
tr|A0A7G2CNI0|A0A7G2CNI0_9TRYP         -------MKPTYILLCFVLLFCSCSGT-----------------------------SVPP    24
tr|S9TUR0|S9TUR0_9TRYP                 MQHSWSSRRSAPAMVCTLLLLLLCSRLVLPTAAGSHS---NPEEEQRIRAAAEAYDVTDP    86
tr|A0A0M9FXG9|A0A0M9FXG9_LEPPY         SGKCRDSGNHAASFLLVLTAALLCIGSLPTEAQRSEPPSVRTAPATTDAARSSSSSNSSA    63
tr|A4H8K3|A4H8K3_LEIBR                 RR------ELVLALCLAVVLHLSCLGAPLSTTVAAAASAAVRDSAARSSSQNASITTASP    57
tr|A4HWX6|A4HWX6_LEIIN                 RS------ALLQLACLVAVLHSSCLSALLSSTIAAAAPAAARDTAISSR-HNTPITVASP    56
tr|Q4QEU0|Q4QEU0_LEIMA                 RS------ALVRLACLVAVLYSSCLSAPLSSTIAAAAPAAARDTAISSR-HNTSVTAASP    56
sp|O81862-2|CHIC_ARATH                 ------------------------------------------------------------    0
tr|S8AUV1|S8AUV1_PENO1                 -----------------------------------------------------------M    1
```

```
tr|A0A179UX46|A0A179UX46_BLAGS      ----------TANVPPPQEVFTLLSGGVPPPVPAAAA---------VQPSLPPPVGPSPG    101
tr|M9NG70|M9NG70_DROME              ------------RGAAWQTLFLLCA---------------------LA-Y-----CINEA    26
tr|Q22468|Q22468_CAEEL              ------------RFVMWMWLFVVWI---------------------TVLFGVSNGGILDR    28


tr|A0A0S4J833|A0A0S4J833_BODSA      AQPFTVFGYLPEYRLR------------GYNYTAAFQTGLTHLIYFSLEVDART--FLP-    78
tr|A0A250X009|A0A250X009_9CHLO      ---------MPEYRLGS-----------NYNYEEAFKNGLTHVIFFSLEVSTSD--FIP-    37
tr|C5K6G1|C5K6G1_PERM5              QERFRVFGYLPEYRLSD-----------DVDFDGIFSHGVTDLIFFSVEVSYLG--II--    66
tr|C1N616|C1N616_MICPC              SSPFAVVAYLPEWRFAS-----------T-DWDA-VCVNVTHLILFSLEVTDDG--SL--    90
tr|A0A7G2CNI0|A0A7G2CNI0_9TRYP      PQPFNVFGYLPEYRANK-----------LFPYETFFANGLTHLIFFSVEINNST--LGID    71
tr|S9TUR0|S9TUR0_9TRYP              GRPFAVFGYLPEYRHQSYGADTCGRHKFYFPYEEFFTRGLTHLIFFSLEADPST--LRLA    144
tr|A0A0M9FXG9|A0A0M9FXG9_LEPPY      APPFAVFGYLPEYRQL------------RFDYEAFFRSGLTHLIFFSAEVDPTS--LQLT    109
tr|A4H8K3|A4H8K3_LEIBR              L-PFAVFGYLPEYRQS------------WFNYEAFFKSGLTHLIFFSAEVDPAT--LRLA    102
tr|A4HWX6|A4HWX6_LEIIN              SPLFTVFGYLPEYRQG------------TFNYEAFFKAGLTHLIFFSAEVDPAT--LRLI    102
tr|Q4QEU0|Q4QEU0_LEIMA              SPLFTVFGYLPEYRQG------------SFNYEAFFKAGLTHLIFFSAEVDPAT--LRLI    102
sp|O81862-2|CHIC_ARATH              ----------------------------------MSSTKLISLIVSIT--------FFLN    18
tr|S8AUV1|S8AUV1_PENO1              SAGLKSIAYFVNWAIYGR----------NYNPQDLPAGKLTHVLYSFANIHPETGEVYLT    51
tr|A0A179UX46|A0A179UX46_BLAGS      SEGFRTVGYFTNWGIYGR----------NYQPMDIPGNYITHILYSFANVRPDSGEVYLT    151
tr|M9NG70|M9NG70_DROME              SSEGRVVCYYTNWSVYRPGT-------AKFNPQNINPYLCTHLVYAFGGFTKDN-QMKPF    78
tr|Q22468|Q22468_CAEEL              SCGRRRVGYITSWGKHPF------------RD--DQAEKLTHLVFAFFVVDSDG-SVKLE    73
                                                                                 ::

tr|A0A0S4J833|A0A0S4J833_BODSA      ----KAKDRLPTMIEAK-----------QAREAA-DAVGGKIILSFGGN-ARSNGFAEMV    121
tr|A0A250X009|A0A250X009_9CHLO      ----SALDRLPPPDILK-----------RARAAA-DAYGGKLLLCFGGN-SRTGGFPGMV    80
tr|C5K6G1|C5K6G1_PERM5              ----QQKQRLPGPAILE-----------KARAAA-DAYGGRLLVSIGGA-GRSSGFADAV    109
tr|C1N616|C1N616_MICPC              ----TSLDRFPSPEAMT-----------TLRAAA-AKRGVKLLLSLGGH-GRTRGFPIVA    133
tr|A0A7G2CNI0|A0A7G2CNI0_9TRYP      ----YIEERLPPMEEWN-----------KIRELA-DKYNVKLMVCVGGS-DRSEAYPTLV    114
tr|S9TUR0|S9TUR0_9TRYP              ----R-EDRLPRATEWA-----------MVRALA-DQHNVKLMVSIGGG-GRSSGFGPLV    186
tr|A0A0M9FXG9|A0A0M9FXG9_LEPPY      ----HVDDRLPSREKWK-----------TLRELA-DTYGVKLLLCIGGG-GRSAGFPRLV    152
tr|A4H8K3|A4H8K3_LEIBR              ----HVDDRLPSDGEWA-----------RIRRLA-DLHGAKLMLCIGGG-GRSSGFSDLV    145
tr|A4HWX6|A4HWX6_LEIIN              ----HVDDRLPSGDEWA-----------RIRRLA-DLHGVKLMLCIGGG-GRSAGFADLV    145
tr|Q4QEU0|Q4QEU0_LEIMA              ----HVDDRLPSGDEWA-----------RIRRLA-DLHGAKLMLCIGGG-GRSAGFADLV    145
sp|O81862-2|CHIC_ARATH              SQTNQV----T-VSSANQP---KFSTF--TQTVQRRNPSVKTLLSIGGGIADKTAYASMA    68
tr|S8AUV1|S8AUV1_PENO1              DSWSDVEKHYA-GDSWNDTGNNVYGCIKQLFLLKKKHRQLKVLLSIGGW-TYSANFAGPA    109
tr|A0A179UX46|A0A179UX46_BLAGS      DTWADVEKRYP-GDSWEEPGENVYGCIKQLYLLKKHYRHLKTLLSIGGW-TYSANFPVPA    209
tr|M9NG70|M9NG70_DROME              DKYQDI----------------EQGGYAKFTGLKTYNKQLKTMIAIGGWNEASSRFSPLV    122
tr|Q22468|Q22468_CAEEL              ---GDA----------------AKARLEHVKEVASRHPDLKLLYAVGGWEN-SQYFSVLT    113
                                                                                 : : ..** .  :   .

tr|A0A0S4J833|A0A0S4J833_BODSA      ATPSSRRVFLGALEALLTQYDFDGVDYNWEYPRDAE-------EWRRWALLLKESKDTLK    174
tr|A0A250X009|A0A250X009_9CHLO      VDKARRRRFLEALSNLMVSHSFDGVDYNWEYPTKYV-------EWEGLADLMKETKALLN    133
tr|C5K6G1|C5K6G1_PERM5              AHNGDVRRLIKQVDDLLNKYQLDGVDFNWEYPQSET-------EWYNFKQMLRWLKVKLR    162
tr|C1N616|C1N616_MICPC              VEKTKRRRLARTLASFCATHGLHGVDYNWEYPASPA-------EWNGMFALLRTTRKVFD    186
tr|A0A7G2CNI0|A0A7G2CNI0_9TRYP      RSNH-TENFFSQLKSLYVDRKLDGFDFNWEYPALEE-------DYRKMNEFLIKARAALN    166
tr|S9TUR0|S9TUR0_9TRYP              RDAAKRRAFVAQLNELYVERRLDGVDVNWEYPQTME-------EWRLLGVLLREMRAALG    239
tr|A0A0M9FXG9|A0A0M9FXG9_LEPPY      GDVLGRRRFVSEVGRVLHERDLDGVDFNWEYPSSMP-------EWLNFGQFLTELRSALN    205
tr|A4H8K3|A4H8K3_LEIBR              GDPVRRLSFIEEVNAVLLTRELDGIDFNWEYPQTMT-------EWLNFGRFLLEIRSALG    198
tr|A4HWX6|A4HWX6_LEIIN              GDTVQRQAFIEEVNAVLLARKLDGIDFNWEYPQTMT-------EWLNFGRFLMELRSSLG    198
tr|Q4QEU0|Q4QEU0_LEIMA              GDTVQRKAFIEEVNAVLLARKLDGIDFNWDYPQTMT-------EWLNFGRFLMELRSSLG    198
sp|O81862-2|CHIC_ARATH              SNPTSRKSFIDSSIRVARSYGFHGLDLDWEYPSSAT-------EMTNFGTLLREWRSAVV    121
tr|S8AUV1|S8AUV1_PENO1              STPAGRARFAESATRLVLDLGLDGIDIDWEYPQDET-------QALNLVELLRACRHNLD    162
tr|A0A179UX46|A0A179UX46_BLAGS      STETGRKTFARTAVQLLADLGFDGIDIDWEYPQDAA-------QAENFVRLLKETREALD    262
tr|M9NG70|M9NG70_DROME              ASNERRQQFIKNILKFLRQNHFDGIDLDWEYPAHRE--GGKSRDRDNYAQFVQELRAEFE    180
tr|Q22468|Q22468_CAEEL              ADHSRRSILISNFVKVIKEYGFDGVDIDWEYPVTGGAVEGTPADRRNYVNLMRELRNELR    173
                                             :        .        :.*.* :*:**                :       ::    :  .

tr|A0A0S4J833|A0A0S4J833_BODSA      GGD-------------------RSQNIVTFTMYLDPKHADV-IQRFNMLEHADYVHCMAY    214
tr|A0A250X009|A0A250X009_9CHLO      G-----------------------QVIVTSAFYPDPNQYII-IKALKLHEICHYLLSMTY    169
tr|C5K6G1|C5K6G1_PERM5              KRE-------------------K-PAIITLAYQPGGLQEDM-IAKLRFAKQCDYFLAMSY    201
tr|C1N616|C1N616_MICPC              GSP-------------------Q-RLTITMAYYPDGRQERE-LKRGGAEEHVELLHAMSY    225
tr|A0A7G2CNI0|A0A7G2CNI0_9TRYP      ST--------------------GRPPIVSMPAHPHPTTSSF-LRLSGINDAVDYIHWMAY    205
tr|S9TUR0|S9TUR0_9TRYP              FETVSGGPDGARP---RRRRMRARAPTLTIALHPHIQGGLL-LRRNDVLDAVDYVHWMAY    295
tr|A0A0M9FXG9|A0A0M9FXG9_LEPPY      RTTA-----------GDHHGRRVGPALITMALHPHPRIPHV-LRVSHVLPSLDYLHWMAY    253
tr|A4H8K3|A4H8K3_LEIBR              YAPAVGREHAAGLIGERHHRTRVRHAVLSMALHPHSSIAAV-LQSARVLRSLDYVHWMAY    257
tr|A4HWX6|A4HWX6_LEIIN              YAAAAGGEHAAGPIGERRHRTRVRGAALSMALHPHPSMAAV-LQSARVLHSLDYVHLMAY    257
tr|Q4QEU0|Q4QEU0_LEIMA              YAAAAGGEHAAEPIGERRYRTRVRGAVLSMALHPHPSMAAV-LQSARMLHSLDYVHLMAY    257
sp|O81862-2|CHIC_ARATH              AEASSSGK---------------PRLLLAAAVFYSNNYYSVLYPVSAVASSLDWVNLMAY    166
tr|S8AUV1|S8AUV1_PENO1              TAAG---P--------------HRKFYLSIACPAGPNNFNK-LKLHEMTPLLDFYNLMAY    204
tr|A0A179UX46|A0A179UX46_BLAGS      AYSAQHAQ--------------GRRLLLTVAVPCGETNYKK-LLMSDMDKYLDFWNLMCY    307
tr|M9NG70|M9NG70_DROME              REAEKTGR---------------TRLLLTMAVPAGIEYIDKGYDVPKLNKYLDWFNVLTY    225
tr|Q22468|Q22468_CAEEL              DLESETGK---------------S-YLISFAGAAGHWVLKPGYDLQQLMKYCDFVNVMSY    217
                                                               ::                         .        : *

tr|A0A0S4J833|A0A0S4J833_BODSA      DQ-HGE-----------------HSTYEFAVSGVRMA-----------------------    233
tr|A0A250X009|A0A250X009_9CHLO      DMVPGK-----------------HSTYEFAVQTIEAW-----------------------    189
tr|C5K6G1|C5K6G1_PERM5              EHPEGK-----------------GE--DLARVVVEAW-----------------------    219
tr|C1N616|C1N616_MICPC              DHPRGS-----------------HSSTALARAAIRNA-----------------------    245
tr|A0A7G2CNI0|A0A7G2CNI0_9TRYP      GTVERL-SV----------------------LMQPLVL--------------------    220
tr|S9TUR0|S9TUR0_9TRYP              DFYLKP-NEQPD--KEP--PL-GHTDYRYASGMLSDEVLEGFLDPAASRGS---------    340
tr|A0A0M9FXG9|A0A0M9FXG9_LEPPY      DHIIAN-DS--------------HSSVAYAASVLQDDVIGDLDDAVYNARLKKIRRQAHG    298
tr|A4H8K3|A4H8K3_LEIBR              DHILGT-EP--------------HSSVEYAASVLSEEMIGLFNEATYNKRSGQTHQE---    299
tr|A4HWX6|A4HWX6_LEIIN              DHVVGT-GP--------------HSSVEYAASVLSEETIGFFNEAAYNRRLGQMHRR---    299
tr|Q4QEU0|Q4QEU0_LEIMA              DHVAGT-GP--------------HSSVEYAASVLSEETIGLFNEAVYNRRLGRMHQR---    299
sp|O81862-2|CHIC_ARATH              DFYGPGWSRV----TGPPAALFDPSNAGPS----GDA-----------------------    195
tr|S8AUV1|S8AUV1_PENO1              DYAG-SWDRV----AGHQANLYPSRSNPASTPFSTIA-----------------------    236
tr|A0A179UX46|A0A179UX46_BLAGS      DFAG-SWDRK----AGHMANIFPSRDVPESTPFNADE-----------------------    339
tr|M9NG70|M9NG70_DROME              DFHS-SHEPSVNH-HAPLYSLEEDSEYNYDAELNIDY-----------------------    260
tr|Q22468|Q22468_CAEEL              DYFG-AWASKWGAYTGPPAPLQFAMPKKFSGRMNVHA-----------------------    253
```

```
tr|A0A0S4J833|A0A0S4J833_BODSA       -----------IEKKMTLSKFTLGVPFYARHVGNGE--PKTYGEIIKEIPK---------    271
tr|A0A250X009|A0A250X009_9CHLO       -----------KQQGLPLDKLALGVPFYGRHMQTGA--PDTYYDLFPKLEKR--------    228
tr|C5K6G1|C5K6G1_PERM5               -----------EQRGLDIRKLALGIPFYGRDLQTGE--ARTYSEISTIP-----------    255
tr|C1N616|C1N616_MICPC               -----------EEAGLDARKITIGLPFYGRHVETGE--WKTYAELDAAH-----------    281
tr|A0A7G2CNI0|A0A7G2CNI0_9TRYP       -----------EGFYDNIKKLTLGLDFYATNGPQNA---TTYSEVVVR--AL--------    256
tr|S9TUR0|S9TUR0_9TRYP               ------RRSKLDIYRDPRRKLTLGIPLFAKHIAENFRHPMPYKDVMTE--QV--------    384
tr|A0A0M9FXG9|A0A0M9FXG9_LEPPY       EAEEDAPQSRKPHEADHRRKLCLGIPFYGRHRADGRVPPETYEHLWQFLRQW--------    350
tr|A4H8K3|A4H8K3_LEIBR               ----------LRTEQDHRRKLTLGIPFYGRHRENRRRQPETYDRLWRSIQQW--------    341
tr|A4HWX6|A4HWX6_LEIIN               ----------PRTEQDHRRKLTLGIPFYGRHREDRRLQPEAYDRLWLFIQEW--------    341
tr|Q4QEU0|Q4QEU0_LEIMA               ----------PRTEQDHRRKLTLGIPFYGRHREDRRLQPEAYDRLWFFIQEW--------    341
sp|O81862-2|CHIC_ARATH               -----GTRSWI-QAGLPAKKAVLGFPYYGYAWRLTNANSHSYYAP--------TTGAAIS    241
tr|S8AUV1|S8AUV1_PENO1               -----ALDHYIGVGGVPPNKMILGMPLYGRAFEGTDGPGTPYSGV--------GPGSW--    281
tr|A0A179UX46|A0A179UX46_BLAGS       -----AITAYV-AGGVHPKKIVFGLPLYGRAFEQTDGPGHPFQGV--------GEGSW--    383
tr|M9NG70|M9NG70_DROME               -----SIKYYL-KAGADRDKLVLGIPTYGRSYTLINEESTELGAP---AEGPGEQGDATR    311
tr|Q22468|Q22468_CAEEL               -----TMKDYS-CQIKATDKINMGVPFYGRFWKNVGDAVDSTDDMWRTATATNSEG-TKF    306
                                                       *  :*.  :.

tr|A0A0S4J833|A0A0S4J833_BODSA       --------------------------------SKRWTQDRVGPYYLNSPSMIQKKTKLAI    299
tr|A0A250X009|A0A250X009_9CHLO       ------------YK----------------NLSKRHAVDELGAVFFNGRSTLKQKAELAI    260
tr|C5K6G1|C5K6G1_PERM5               ------------------------------D--------AQHKFVYDTPEEVQARTRYAL    277
tr|C1N616|C1N616_MICPC               ------------------------------GVSAEPSKDEAGGYAFNGADTIRLKTSEAK    311
tr|A0A7G2CNI0|A0A7G2CNI0_9TRYP       ------------MQSEKPATDE--------VVQRLKESSNEGGFDFTGYYGVQKKMDRAR    296
tr|S9TUR0|S9TUR0_9TRYP               ------------ARVAPPQSTASSAELERQMAAFIRPVNELDRYTFSGYNEIQKKVELAR    432
tr|A0A0M9FXG9|A0A0M9FXG9_LEPPY       ------------AGKRHPDWVE--------GGPELRSLSEYAGYDYNGYDDVRRKMRLAR    390
tr|A4H8K3|A4H8K3_LEIBR               ------------ARKKHPAWVE--------GGAELRALNNYGGYSFTGYNDVKRKMRLAR    381
tr|A4HWX6|A4HWX6_LEIIN               ------------ASKSHPTWVE--------GGAELRALSEYGGYSFTGYDDVKRKMQLTR    381
tr|Q4QEU0|Q4QEU0_LEIMA               ------------ASKNHPTWVE--------GGAELRALSEYGGYSFTCYDDVKRKMRLAR    381
sp|O81862-2|CHIC_ARATH               PDGSIGYGQIRKFIVDNG--ATTVYNS-TVVGDYCY--AGTNWIGYDDNQSIVTKVRYAK    296
tr|S8AUV1|S8AUV1_PENO1               ENGVWDYK----ALPRPG--ATEHLDT-EADASWSYDPASRTMVSYDTVVMSERKAAFVK    334
tr|A0A179UX46|A0A179UX46_BLAGS       ESGVWDYK----VLPQPG--SEEVNDD-DLKASWSYDRNARKMISYDTPAIVAQKADYIR    436
tr|M9NG70|M9NG70_DROME               EKGYLAYYEICQTLKDDPEWTVVQPNA-NVMGP--YAYRRNQWVGYDDEAIVRKKAEYVV    368
tr|Q22468|Q22468_CAEEL               EGGDVQWRDLHEKFDT----TKTKFHS-GSKTPFIWLSEQKTFVGYENAESLKHKVDYIV    361
                                                                                          :

tr|A0A0S4J833|A0A0S4J833_BODSA       DN----QLGGVMIWELGQDLQPE-S---------------HRNSLLRGILKATLTRG---    336
tr|A0A250X009|A0A250X009_9CHLO       RE----GLGGIMIWELGQDISPP-S---------------HESALMSGLGEAIVPMG---    297
tr|C5K6G1|C5K6G1_PERM5               EE----GLGGVMIWELGQDLPSS-----------------NERSLLAAIGKVVSSPS---    313
tr|C1N616|C1N616_MICPC               AA----GTGGVMIWEAGQDLHPS-----------------HARSLLAVVAAEAWGDE---    347
tr|A0A7G2CNI0|A0A7G2CNI0_9TRYP       QA----NFTGVMIWELGQDILSYPPEVLKKATDDATIEKLYNVSLMRSIHEKLT-EWKKA    351
tr|S9TUR0|S9TUR0_9TRYP               TGGPPGGIGGIMIWELGQDLLPPSE---------------HPLAAMNAISEVVQERWGGP    477
tr|A0A0M9FXG9|A0A0M9FXG9_LEPPY       SS----GIAGIMIWELGQDV-PPGT---------------SPMSLMTAVQEQLA-EWDKD    429
tr|A4H8K3|A4H8K3_LEIBR               AA----NLSGIMIWELGQDV-PPGT---------------SPMSLMAAVHEQLA-DWGLL    420
tr|A4HWX6|A4HWX6_LEIIN               AA----NLSGIMIWELGQDV-PPGT---------------SPMSLMTAVHEQLA-DWGLL    420
tr|Q4QEU0|Q4QEU0_LEIMA               AA----NLSGIMIWELGQDV-PPGT---------------SPMSLMTAVHEQLA-DWGLL    420
sp|O81862-2|CHIC_ARATH               QR----GLLGYFSWHVGADDNS-GLSR------------AASQAWDATTAT------TR-    332
tr|S8AUV1|S8AUV1_PENO1               YR----QLGGGMWWESSGDKGG-KDAN------------PAQGSLIGTF--------VD-    368
tr|A0A179UX46|A0A179UX46_BLAGS       KR----GMGGGMWWELSGDAPV-----------------GSERSLIATT--------VN-    466
tr|M9NG70|M9NG70_DROME               AQ----GLGGIMFWAIDNDDFR-GTCN------------GKPYPLIEAAKEAMVEALGL-    410
tr|Q22468|Q22468_CAEEL               EN----NIGGVMIWAIDFDDDQ-GTLL------------NSAA------AESICTTSTK-    397
                                              * : *  . *

tr|A0A0S4J833|A0A0S4J833_BODSA       LALPK----LQE-QLRAAK--EETPS---LKEPRDE---------L--------------    363
tr|A0A250X009|A0A250X009_9CHLO       SLQKT----VLGAGISSEVEIEEELN---VVLSTDDVATVNSKTEL--------------    336
tr|C5K6G1|C5K6G1_PERM5               --IEE----DGG---EKEEDDDEEL-----------------------------------    329
tr|C1N616|C1N616_MICPC               --GKN----AVG---EKKRAR-EEL-----------------------------------    362
tr|A0A7G2CNI0|A0A7G2CNI0_9TRYP       DPPKKD-I--------------------------------NEEGDL--------------    364
tr|S9TUR0|S9TUR0_9TRYP               ARPWTGEVRDGAGGVTREADAASTPTEKATAPPRE--ADYEELSDL--------------    521
tr|A0A0M9FXG9|A0A0M9FXG9_LEPPY       TASNA----DESDACLRTPARLEH---TRENATRLVGSDD-ADEDL--------------    467
tr|A4H8K3|A4H8K3_LEIBR               TDSGH----DSGGDVNGDNTG--H---LQQRQRRYLSSDVAEDGDL--------------    457
tr|A4HWX6|A4HWX6_LEIIN               TDGGR----GSGGNVNSDNAY--D---RPQPPPQHSSPDVAEDGDL--------------    457
tr|Q4QEU0|Q4QEU0_LEIMA               TDSGR----GIGGNVNSDNAH--D---RPQPSPQHSSPDVAEDGDL--------------    457
sp|O81862-2|CHIC_ARATH               ---------TIQ-KV---------------------------------------------    337
tr|S8AUV1|S8AUV1_PENO1               ---------GIG-GVTA-L---D----GIENALDYPESKYDNLRAGFPGQ----------    400
tr|A0A179UX46|A0A179UX46_BLAGS       ---------GLG-GVGN-L---D----HSENLLDYPASRYENLRKGFQ------------    496
tr|M9NG70|M9NG70_DROME               ---------GIN-EVAK-P---S----GPQKPSRSRSRDNASNRNRLNGKTEAPLSSRRP    452
tr|Q22468|Q22468_CAEEL               ---------SFN-YKCS-----------PVDDKRWWTYDDNEELAGMCGKSSPLIDGYYP    436


tr|A0A0S4J833|A0A0S4J833_BODSA       ------------------------------------------------------------    363
tr|A0A250X009|A0A250X009_9CHLO       ------------------------------------------------------------    336
tr|C5K6G1|C5K6G1_PERM5               ------------------------------------------------------------    329
tr|C1N616|C1N616_MICPC               ------------------------------------------------------------    362
tr|A0A7G2CNI0|A0A7G2CNI0_9TRYP       ------------------------------------------------------------    364
tr|S9TUR0|S9TUR0_9TRYP               ------------------------------------------------------------    521
tr|A0A0M9FXG9|A0A0M9FXG9_LEPPY       ------------------------------------------------------------    467
tr|A4H8K3|A4H8K3_LEIBR               ------------------------------------------------------------    457
tr|A4HWX6|A4HWX6_LEIIN               ------------------------------------------------------------    457
tr|Q4QEU0|Q4QEU0_LEIMA               ------------------------------------------------------------    457
sp|O81862-2|CHIC_ARATH               ------------------------------------------------------------    337
tr|S8AUV1|S8AUV1_PENO1               ------------------------------------------------------------    400
tr|A0A179UX46|A0A179UX46_BLAGS       ------------------------------------------------------------    496
tr|M9NG70|M9NG70_DROME               SA-------------------------------TRRPAVSSTQAPPPS--TTFKL-TEAE    478
tr|Q22468|Q22468_CAEEL               VCDPDDPGHACCGKYGYCGSGAEFCSCPECIDYGADPNLILKEPVKPSQKITWYTSDAGE    496


tr|A0A0S4J833|A0A0S4J833_BODSA       ------------------------------------------------------------    363
```

```
tr|A0A250X009|A0A250X009_9CHLO      ------------------------------------------------------------   336
tr|C5K6G1|C5K6G1_PERM5              ------------------------------------------------------------   329
tr|C1N616|C1N616_MICPC              ------------------------------------------------------------   362
tr|A0A7G2CNI0|A0A7G2CNI0_9TRYP      ------------------------------------------------------------   364
tr|S9TUR0|S9TUR0_9TRYP              ------------------------------------------------------------   521
tr|A0A0M9FXG9|A0A0M9FXG9_LEPPY      ------------------------------------------------------------   467
tr|A4H8K3|A4H8K3_LEIBR              ------------------------------------------------------------   457
tr|A4HWX6|A4HWX6_LEIIN              ------------------------------------------------------------   457
tr|Q4QEU0|Q4QEU0_LEIMA              ------------------------------------------------------------   457
sp|O81862-2|CHIC_ARATH              ------------------------------------------------------------   337
tr|S8AUV1|S8AUV1_PENO1              ------------------------------------------------------------   400
tr|A0A179UX46|A0A179UX46_BLAGS      ------------------------------------------------------------   496
tr|M9NG70|M9NG70_DROME              GSSLYIGGRASTTPP------PPTTPDPGSDFKCEEEGFFQHPRDC--------------   518
tr|Q22468|Q22468_CAEEL              GK----RGRCGRDVPPLEGEAPTCNPDDANAHCCSNGGYCGNSKEHCECNGCIDFAKQRD   552


tr|A0A0S4J833|A0A0S4J833_BODSA      ------------------------------------------------------------   363
tr|A0A250X009|A0A250X009_9CHLO      ------------------------------------------------------------   336
tr|C5K6G1|C5K6G1_PERM5              ------------------------------------------------------------   329
tr|C1N616|C1N616_MICPC              ------------------------------------------------------------   362
tr|A0A7G2CNI0|A0A7G2CNI0_9TRYP      ------------------------------------------------------------   364
tr|S9TUR0|S9TUR0_9TRYP              ------------------------------------------------------------   521
tr|A0A0M9FXG9|A0A0M9FXG9_LEPPY      ------------------------------------------------------------   467
tr|A4H8K3|A4H8K3_LEIBR              ------------------------------------------------------------   457
tr|A4HWX6|A4HWX6_LEIIN              ------------------------------------------------------------   457
tr|Q4QEU0|Q4QEU0_LEIMA              ------------------------------------------------------------   457
sp|O81862-2|CHIC_ARATH              ------------------------------------------------------------   337
tr|S8AUV1|S8AUV1_PENO1              ------------------------------------------------------------   400
tr|A0A179UX46|A0A179UX46_BLAGS      ------------------------------------------------------------   496
tr|M9NG70|M9NG70_DROME              ----KKYYWCLDSGPSGLGIVAH------------------MFTCPSGLYFNPAADSCDF   556
tr|Q22468|Q22468_CAEEL              FKYKPLEWWTFSENPANVGRCGYNAPRLSTGKIPKCDPDSESYCCSNSGYCGKGEQYCS-   611


tr|A0A0S4J833|A0A0S4J833_BODSA      ------------------------------------------------------------   363
tr|A0A250X009|A0A250X009_9CHLO      ------------------------------------------------------------   336
tr|C5K6G1|C5K6G1_PERM5              ------------------------------------------------------------   329
tr|C1N616|C1N616_MICPC              ------------------------------------------------------------   362
tr|A0A7G2CNI0|A0A7G2CNI0_9TRYP      ------------------------------------------------------------   364
tr|S9TUR0|S9TUR0_9TRYP              ------------------------------------------------------------   521
tr|A0A0M9FXG9|A0A0M9FXG9_LEPPY      ------------------------------------------------------------   467
tr|A4H8K3|A4H8K3_LEIBR              ------------------------------------------------------------   457
tr|A4HWX6|A4HWX6_LEIIN              ------------------------------------------------------------   457
tr|Q4QEU0|Q4QEU0_LEIMA              ------------------------------------------------------------   457
sp|O81862-2|CHIC_ARATH              ------------------------------------------------------------   337
tr|S8AUV1|S8AUV1_PENO1              ------------------------------------------------------------   400
tr|A0A179UX46|A0A179UX46_BLAGS      ------------------------------------------------------------   496
tr|M9NG70|M9NG70_DROME              ARNVPCKTKKSTTAAPVTSTTPATTTVRSNRVTAAPTSRPVYPRTTTTTSTTTTTTTTPS   616
tr|Q22468|Q22468_CAEEL              --CLGCADFKANP-----------------------------------------------   622


tr|A0A0S4J833|A0A0S4J833_BODSA      ------------------------------------------------------------   363
tr|A0A250X009|A0A250X009_9CHLO      ------------------------------------------------------------   336
tr|C5K6G1|C5K6G1_PERM5              ------------------------------------------------------------   329
tr|C1N616|C1N616_MICPC              ------------------------------------------------------------   362
tr|A0A7G2CNI0|A0A7G2CNI0_9TRYP      ------------------------------------------------------------   364
tr|S9TUR0|S9TUR0_9TRYP              ------------------------------------------------------------   521
tr|A0A0M9FXG9|A0A0M9FXG9_LEPPY      ------------------------------------------------------------   467
tr|A4H8K3|A4H8K3_LEIBR              ------------------------------------------------------------   457
tr|A4HWX6|A4HWX6_LEIIN              ------------------------------------------------------------   457
tr|Q4QEU0|Q4QEU0_LEIMA              ------------------------------------------------------------   457
sp|O81862-2|CHIC_ARATH              ------------------------------------------------------------   337
tr|S8AUV1|S8AUV1_PENO1              ------------------------------------------------------------   400
tr|A0A179UX46|A0A179UX46_BLAGS      ------------------------------------------------------------   496
tr|M9NG70|M9NG70_DROME              TVDEDLEYEEDTDELSPSKSTDAEEDPQVIKELIDLIRKVGGVEQLEKHLLRNKDGSITL   676
tr|Q22468|Q22468_CAEEL              ----AFEY----------------------------------------------------   626


tr|A0A0S4J833|A0A0S4J833_BODSA      ------------------------------------------------------------   363
tr|A0A250X009|A0A250X009_9CHLO      ------------------------------------------------------------   336
tr|C5K6G1|C5K6G1_PERM5              ------------------------------------------------------------   329
tr|C1N616|C1N616_MICPC              ------------------------------------------------------------   362
tr|A0A7G2CNI0|A0A7G2CNI0_9TRYP      ------------------------------------------------------------   364
tr|S9TUR0|S9TUR0_9TRYP              ------------------------------------------------------------   521
tr|A0A0M9FXG9|A0A0M9FXG9_LEPPY      ------------------------------------------------------------   467
tr|A4H8K3|A4H8K3_LEIBR              ------------------------------------------------------------   457
tr|A4HWX6|A4HWX6_LEIIN              ------------------------------------------------------------   457
tr|Q4QEU0|Q4QEU0_LEIMA              ------------------------------------------------------------   457
sp|O81862-2|CHIC_ARATH              ------------------------------------------------------------   337
tr|S8AUV1|S8AUV1_PENO1              ------------------------------------------------------------   400
tr|A0A179UX46|A0A179UX46_BLAGS      ------------------------------------------------------------   496
tr|M9NG70|M9NG70_DROME              KENSATGAATTPSTISKSLYDRVLSRPGTLNSFSRNRFKISEASETSTEPTTSSSSSRGS   736
tr|Q22468|Q22468_CAEEL              ------------------------------------------------------------   626


tr|A0A0S4J833|A0A0S4J833_BODSA      ------------------------------------------------------------   363
tr|A0A250X009|A0A250X009_9CHLO      ------------------------------------------------------------   336
tr|C5K6G1|C5K6G1_PERM5              ------------------------------------------------------------   329
tr|C1N616|C1N616_MICPC              ------------------------------------------------------------   362
```

```
tr|A0A7G2CNI0|A0A7G2CNI0_9TRYP      ------------------------------------------------------------   364
tr|S9TUR0|S9TUR0_9TRYP              ------------------------------------------------------------   521
tr|A0A0M9FXG9|A0A0M9FXG9_LEPPY      ------------------------------------------------------------   467
tr|A4H8K3|A4H8K3_LEIBR              ------------------------------------------------------------   457
tr|A4HWX6|A4HWX6_LEIIN              ------------------------------------------------------------   457
tr|Q4QEU0|Q4QEU0_LEIMA              ------------------------------------------------------------   457
sp|O81862-2|CHIC_ARATH              ------------------------------------------------------------   337
tr|S8AUV1|S8AUV1_PENO1              ------------------------------------------------------------   400
tr|A0A179UX46|A0A179UX46_BLAGS      ------------------------------------------------------------   496
tr|M9NG70|M9NG70_DROME              STLTSNTNSKYSSVLRGNSRQGPQNEGIEKLAEFDGFLKERKQYVTINRHRSASQGDEEE   796
tr|Q22468|Q22468_CAEEL              ------------------------------------------------------------   626

tr|A0A0S4J833|A0A0S4J833_BODSA      ------------------------------------------------------------   363
tr|A0A250X009|A0A250X009_9CHLO      ------------------------------------------------------------   336
tr|C5K6G1|C5K6G1_PERM5              ------------------------------------------------------------   329
tr|C1N616|C1N616_MICPC              ------------------------------------------------------------   362
tr|A0A7G2CNI0|A0A7G2CNI0_9TRYP      ------------------------------------------------------------   364
tr|S9TUR0|S9TUR0_9TRYP              ------------------------------------------------------------   521
tr|A0A0M9FXG9|A0A0M9FXG9_LEPPY      ------------------------------------------------------------   467
tr|A4H8K3|A4H8K3_LEIBR              ------------------------------------------------------------   457
tr|A4HWX6|A4HWX6_LEIIN              ------------------------------------------------------------   457
tr|Q4QEU0|Q4QEU0_LEIMA              ------------------------------------------------------------   457
sp|O81862-2|CHIC_ARATH              ------------------------------------------------------------   337
tr|S8AUV1|S8AUV1_PENO1              ------------------------------------------------------------   400
tr|A0A179UX46|A0A179UX46_BLAGS      ------------------------------------------------------------   496
tr|M9NG70|M9NG70_DROME              HADQQEEEENLAEVETTTRRPLSSITPSYTSLRRSRPTTVAPPAEESHEEAEQQTQTQVK   856
tr|Q22468|Q22468_CAEEL              ------------------------------------------------------------   626

tr|A0A0S4J833|A0A0S4J833_BODSA      ------------------------------------------------------------   363
tr|A0A250X009|A0A250X009_9CHLO      ------------------------------------------------------------   336
tr|C5K6G1|C5K6G1_PERM5              ------------------------------------------------------------   329
tr|C1N616|C1N616_MICPC              ------------------------------------------------------------   362
tr|A0A7G2CNI0|A0A7G2CNI0_9TRYP      ------------------------------------------------------------   364
tr|S9TUR0|S9TUR0_9TRYP              ------------------------------------------------------------   521
tr|A0A0M9FXG9|A0A0M9FXG9_LEPPY      ------------------------------------------------------------   467
tr|A4H8K3|A4H8K3_LEIBR              ------------------------------------------------------------   457
tr|A4HWX6|A4HWX6_LEIIN              ------------------------------------------------------------   457
tr|Q4QEU0|Q4QEU0_LEIMA              ------------------------------------------------------------   457
sp|O81862-2|CHIC_ARATH              ------------------------------------------------------------   337
tr|S8AUV1|S8AUV1_PENO1              ------------------------------------------------------------   400
tr|A0A179UX46|A0A179UX46_BLAGS      ------------------------------------------------------------   496
tr|M9NG70|M9NG70_DROME              SYATLSRTRGRTTSSPEVTEAAPSSTTNRYKYFERTRPTKSATAEDSEDPTEDEEEEYED   916
tr|Q22468|Q22468_CAEEL              ------------------------------------------------------------   626

tr|A0A0S4J833|A0A0S4J833_BODSA      ------------------------------------------------------------   363
tr|A0A250X009|A0A250X009_9CHLO      ------------------------------------------------------------   336
tr|C5K6G1|C5K6G1_PERM5              ------------------------------------------------------------   329
tr|C1N616|C1N616_MICPC              ------------------------------------------------------------   362
tr|A0A7G2CNI0|A0A7G2CNI0_9TRYP      ------------------------------------------------------------   364
tr|S9TUR0|S9TUR0_9TRYP              ------------------------------------------------------------   521
tr|A0A0M9FXG9|A0A0M9FXG9_LEPPY      ------------------------------------------------------------   467
tr|A4H8K3|A4H8K3_LEIBR              ------------------------------------------------------------   457
tr|A4HWX6|A4HWX6_LEIIN              ------------------------------------------------------------   457
tr|Q4QEU0|Q4QEU0_LEIMA              ------------------------------------------------------------   457
sp|O81862-2|CHIC_ARATH              ------------------------------------------------------------   337
tr|S8AUV1|S8AUV1_PENO1              ------------------------------------------------------------   400
tr|A0A179UX46|A0A179UX46_BLAGS      ------------------------------------------------------------   496
tr|M9NG70|M9NG70_DROME              EQKDIVTLRQPTTGSVSRRPVLSVRRRIINSPTSAAVETTTQQPAAEPPTTSKYSRLRSR   976
tr|Q22468|Q22468_CAEEL              ------------------------------------------------------------   626

tr|A0A0S4J833|A0A0S4J833_BODSA      ------------------------------------------------------------   363
tr|A0A250X009|A0A250X009_9CHLO      ------------------------------------------------------------   336
tr|C5K6G1|C5K6G1_PERM5              ------------------------------------------------------------   329
tr|C1N616|C1N616_MICPC              ------------------------------------------------------------   362
tr|A0A7G2CNI0|A0A7G2CNI0_9TRYP      ------------------------------------------------------------   364
tr|S9TUR0|S9TUR0_9TRYP              ------------------------------------------------------------   521
tr|A0A0M9FXG9|A0A0M9FXG9_LEPPY      ------------------------------------------------------------   467
tr|A4H8K3|A4H8K3_LEIBR              ------------------------------------------------------------   457
tr|A4HWX6|A4HWX6_LEIIN              ------------------------------------------------------------   457
tr|Q4QEU0|Q4QEU0_LEIMA              ------------------------------------------------------------   457
sp|O81862-2|CHIC_ARATH              ------------------------------------------------------------   337
tr|S8AUV1|S8AUV1_PENO1              ------------------------------------------------------------   400
tr|A0A179UX46|A0A179UX46_BLAGS      ------------------------------------------------------------   496
tr|M9NG70|M9NG70_DROME              PSATATAAAAAATTTTAAATTAFPAATSAPGGRTTSNIYLSKLKAKSGAAAAAAASGEAA   1036
tr|Q22468|Q22468_CAEEL              ------------------------------------------------------------   626

tr|A0A0S4J833|A0A0S4J833_BODSA      ------------------------------------------------------------   363
tr|A0A250X009|A0A250X009_9CHLO      ------------------------------------------------------------   336
tr|C5K6G1|C5K6G1_PERM5              ------------------------------------------------------------   329
tr|C1N616|C1N616_MICPC              ------------------------------------------------------------   362
tr|A0A7G2CNI0|A0A7G2CNI0_9TRYP      ------------------------------------------------------------   364
tr|S9TUR0|S9TUR0_9TRYP              ------------------------------------------------------------   521
tr|A0A0M9FXG9|A0A0M9FXG9_LEPPY      ------------------------------------------------------------   467
```

```
tr|A4H8K3|A4H8K3_LEIBR            ------------------------------------------------------------    457
tr|A4HWX6|A4HWX6_LEIIN            ------------------------------------------------------------    457
tr|Q4QEU0|Q4QEU0_LEIMA            ------------------------------------------------------------    457
sp|O81862-2|CHIC_ARATH            ------------------------------------------------------------    337
tr|S8AUV1|S8AUV1_PENO1            ------------------------------------------------------------    400
tr|A0A179UX46|A0A179UX46_BLAGS    ------------------------------------------------------------    496
tr|M9NG70|M9NG70_DROME            TLTPATSNISSSSNDITQKQHKFQPASFALRRQFQTRRLTTFAPAANGDESATEVPRTQN    1096
tr|Q22468|Q22468_CAEEL            ------------------------------------------------------------    626

tr|A0A0S4J833|A0A0S4J833_BODSA    ------------------------------------------------------------    363
tr|A0A250X009|A0A250X009_9CHLO    ------------------------------------------------------------    336
tr|C5K6G1|C5K6G1_PERM5            ------------------------------------------------------------    329
tr|C1N616|C1N616_MICPC            ------------------------------------------------------------    362
tr|A0A7G2CNI0|A0A7G2CNI0_9TRYP    ------------------------------------------------------------    364
tr|S9TUR0|S9TUR0_9TRYP            ------------------------------------------------------------    521
tr|A0A0M9FXG9|A0A0M9FXG9_LEPPY    ------------------------------------------------------------    467
tr|A4H8K3|A4H8K3_LEIBR            ------------------------------------------------------------    457
tr|A4HWX6|A4HWX6_LEIIN            ------------------------------------------------------------    457
tr|Q4QEU0|Q4QEU0_LEIMA            ------------------------------------------------------------    457
sp|O81862-2|CHIC_ARATH            ------------------------------------------------------------    337
tr|S8AUV1|S8AUV1_PENO1            ------------------------------------------------------------    400
tr|A0A179UX46|A0A179UX46_BLAGS    ------------------------------------------------------------    496
tr|M9NG70|M9NG70_DROME            PLFKRRLTLISTTPPSARTTNPPVSGLETTTTLYLNDDDEDQVAKSSIHTSRFNQIPEQV    1156
tr|Q22468|Q22468_CAEEL            ------------------------------------------------------------    626

tr|A0A0S4J833|A0A0S4J833_BODSA    ------------------------------------------------------------    363
tr|A0A250X009|A0A250X009_9CHLO    ------------------------------------------------------------    336
tr|C5K6G1|C5K6G1_PERM5            ------------------------------------------------------------    329
tr|C1N616|C1N616_MICPC            ------------------------------------------------------------    362
tr|A0A7G2CNI0|A0A7G2CNI0_9TRYP    ------------------------------------------------------------    364
tr|S9TUR0|S9TUR0_9TRYP            ------------------------------------------------------------    521
tr|A0A0M9FXG9|A0A0M9FXG9_LEPPY    ------------------------------------------------------------    467
tr|A4H8K3|A4H8K3_LEIBR            ------------------------------------------------------------    457
tr|A4HWX6|A4HWX6_LEIIN            ------------------------------------------------------------    457
tr|Q4QEU0|Q4QEU0_LEIMA            ------------------------------------------------------------    457
sp|O81862-2|CHIC_ARATH            ------------------------------------------------------------    337
tr|S8AUV1|S8AUV1_PENO1            ------------------------------------------------------------    400
tr|A0A179UX46|A0A179UX46_BLAGS    ------------------------------------------------------------    496
tr|M9NG70|M9NG70_DROME            RPREEYDLALPAQPLKSTSTTVSTTANAPLPVIGQGIRRQLIPRPRRPQSTTSTPPTTTT    1216
tr|Q22468|Q22468_CAEEL            ------------------------------------------------------------    626

tr|A0A0S4J833|A0A0S4J833_BODSA    ------------------------------------------------------------    363
tr|A0A250X009|A0A250X009_9CHLO    ------------------------------------------------------------    336
tr|C5K6G1|C5K6G1_PERM5            ------------------------------------------------------------    329
tr|C1N616|C1N616_MICPC            ------------------------------------------------------------    362
tr|A0A7G2CNI0|A0A7G2CNI0_9TRYP    ------------------------------------------------------------    364
tr|S9TUR0|S9TUR0_9TRYP            ------------------------------------------------------------    521
tr|A0A0M9FXG9|A0A0M9FXG9_LEPPY    ------------------------------------------------------------    467
tr|A4H8K3|A4H8K3_LEIBR            ------------------------------------------------------------    457
tr|A4HWX6|A4HWX6_LEIIN            ------------------------------------------------------------    457
tr|Q4QEU0|Q4QEU0_LEIMA            ------------------------------------------------------------    457
sp|O81862-2|CHIC_ARATH            ------------------------------------------------------------    337
tr|S8AUV1|S8AUV1_PENO1            ------------------------------------------------------------    400
tr|A0A179UX46|A0A179UX46_BLAGS    ------------------------------------------------------------    496
tr|M9NG70|M9NG70_DROME            SGSTLRSTTSAGHSAPPLSRRQQSRRRPHKYIEVYSRPPAKTAVISATSSSQFLDEGFPV    1276
tr|Q22468|Q22468_CAEEL            ------------------------------------------------------------    626

tr|A0A0S4J833|A0A0S4J833_BODSA    ------------------------------------------------------------    363
tr|A0A250X009|A0A250X009_9CHLO    ------------------------------------------------------------    336
tr|C5K6G1|C5K6G1_PERM5            ------------------------------------------------------------    329
tr|C1N616|C1N616_MICPC            ------------------------------------------------------------    362
tr|A0A7G2CNI0|A0A7G2CNI0_9TRYP    ------------------------------------------------------------    364
tr|S9TUR0|S9TUR0_9TRYP            ------------------------------------------------------------    521
tr|A0A0M9FXG9|A0A0M9FXG9_LEPPY    ------------------------------------------------------------    467
tr|A4H8K3|A4H8K3_LEIBR            ------------------------------------------------------------    457
tr|A4HWX6|A4HWX6_LEIIN            ------------------------------------------------------------    457
tr|Q4QEU0|Q4QEU0_LEIMA            ------------------------------------------------------------    457
sp|O81862-2|CHIC_ARATH            ------------------------------------------------------------    337
tr|S8AUV1|S8AUV1_PENO1            ------------------------------------------------------------    400
tr|A0A179UX46|A0A179UX46_BLAGS    ------------------------------------------------------------    496
tr|M9NG70|M9NG70_DROME            VHSQVAQRRRSGSILPTKNDPKVIVHGHGIIECRAQGNFPHPLNCRKFISCARFEETGGI    1336
tr|Q22468|Q22468_CAEEL            ------------------------------------------------------------    626

tr|A0A0S4J833|A0A0S4J833_BODSA    -------------------------------    363
tr|A0A250X009|A0A250X009_9CHLO    -------------------------------    336
tr|C5K6G1|C5K6G1_PERM5            -------------------------------    329
tr|C1N616|C1N616_MICPC            -------------------------------    362
tr|A0A7G2CNI0|A0A7G2CNI0_9TRYP    -------------------------------    364
tr|S9TUR0|S9TUR0_9TRYP            -------------------------------    521
tr|A0A0M9FXG9|A0A0M9FXG9_LEPPY    -------------------------------    467
tr|A4H8K3|A4H8K3_LEIBR            -------------------------------    457
tr|A4HWX6|A4HWX6_LEIIN            -------------------------------    457
tr|Q4QEU0|Q4QEU0_LEIMA            -------------------------------    457
```

```
sp|O81862-2|CHIC_ARATH              -------------------------------    337
tr|S8AUV1|S8AUV1_PENO1              -------------------------------    400
tr|A0A179UX46|A0A179UX46_BLAGS      -------------------------------    496
tr|M9NG70|M9NG70_DROME              VGWEYTCPKGLTYDGVGGMCTWSPSDQPCRD    1367
tr|Q22468|Q22468_CAEEL              -------------------------------    626
```

### 04. MANF (ARMET) family proteins

These proteins are called by the utter misnomer "Mesencephalic astrocyte-secreted neurotrophic factor" in animals, despite the fact that they are not secreted by default, but retained in the ER by a KDEL signal. The real physiological function of these proteins is unclear. However, they are a well conserved group in animals and some other eukaryotes, hinting at a critical (possibly protein chaperoning) function. Our alignments show that both the protein and its KDEL signal is an ancient heritage in kinetoplastids: already found in the distinctly related euglenids. While missing from fungi and higher plants, MANF (ARMET) proteins are still seen among SAR supergroup (Stramenopila-Alveolata-Rhizaria) organisms, showing that its loss is likely secondary.

```
tr|C1EB43|C1EB43_MICCC              ------------------------------------------------------------    0
tr|A0A0L1KUF0|A0A0L1KUF0_9EUGL      ------------------------------------------------------------    0
tr|A0A0S4IZ22|A0A0S4IZ22_BODSA      ------------------------------------------------------------    0
tr|Q57VE7|Q57VE7_TRYB2              ------------------------------------------------------------    0
tr|A0A1X0NHR2|A0A1X0NHR2_9TRYP      ------------------------------------------------------------    0
tr|Q4D5D6|Q4D5D6_TRYCC              ---------------------------------------------------MPPR-----    4
tr|Q4CP18|Q4CP18_TRYCC              ---------------------------------------------------MPPR-----    4
tr|A0A3R7N5F9|A0A3R7N5F9_9TRYP      ------------------------------------------------------M-----    1
tr|A0A422N9T4|A0A422N9T4_TRYRA      ------------------------------------------------------M-----    1
tr|S9WF29|S9WF29_9TRYP              ------------------------------------------------------------    0
tr|S9UQX2|S9UQX2_9TRYP              ------------------------------------------------------------    0
tr|S9UU37|S9UU37_9TRYP              --------------------------------MCAT--------F-----SSPPTFFFLL    15
tr|A0A0M9GAG4|A0A0M9GAG4_LEPPY      ------------------------------------------------------------    0
tr|A4HME3|A4HME3_LEIBR              ------------------------------------------------------------    0
tr|E9AEU4|E9AEU4_LEIMA              ------------------------------------------------------------    0
tr|A4IB15|A4IB15_LEIIN              ------------------------------------------------------------    0
tr|A0A833SEV5|A0A833SEV5_PHYIN      --MLTKHVMLLALVALVLGNAPYVQADNKECEVCVKVIDDLKATYAQLQEENPKGKTQAL    58
sp|Q9CXI5|MANF_MOUSE                --MWATRGLAVALALSVLP--DSRALRPGDCEVCISYLGRFYQDL---KD-RDVTFSPAT    52
sp|Q9XZ63|ARMET_DROME               MKTWYMVV-VIGFLATLAQ--TSLALKEEDCEVCVKTVRRFADSL---DD-ST-KKDYKQ    52

tr|C1EB43|C1EB43_MICCC              ---------------------------MR---------GWLKIAFIALL-----------    13
tr|A0A0L1KUF0|A0A0L1KUF0_9EUGL      ----------------------------M---------SFP-----ALL-----------    7
tr|A0A0S4IZ22|A0A0S4IZ22_BODSA      ----------------------------M---------QRSGLALLLCA-----------    12
tr|Q57VE7|Q57VE7_TRYB2              ----------M--------FSMSPCVL-------------TALLAAAVL-----------    18
tr|A0A1X0NHR2|A0A1X0NHR2_9TRYP      --------------------MRDIGLR-------------PLLLLL-VL-----------    15
tr|Q4D5D6|Q4D5D6_TRYCC              ---------RY--------LRNGAALL-------------PWLL--VLL-----------    21
tr|Q4CP18|Q4CP18_TRYCC              ---------RY--------LRNGAAPF-------------PWLL--VLL-----------    21
tr|A0A3R7N5F9|A0A3R7N5F9_9TRYP      ---------SH--------LRDRA----------------PLLLLLLLL-----------    17
tr|A0A422N9T4|A0A422N9T4_TRYRA      ---------PH--------LRDRVAVL-------------PLLFVLLLL-----------    20
tr|S9WF29|S9WF29_9TRYP              ---------------------------------------MRKLFIVAVL-----------    10
tr|S9UQX2|S9UQX2_9TRYP              ----------------------------M---------QRKLYLFLALT-----------    12
tr|S9UU37|S9UU37_9TRYP              VDFRFFFLCSY--------LCFRAALLKM---------QRKLYLFLALT-----------    47
tr|A0A0M9GAG4|A0A0M9GAG4_LEPPY      ------------------------------------------------------------    0
tr|A4HME3|A4HME3_LEIBR              ----------------------------------------MKAFFITVL-----------    9
tr|E9AEU4|E9AEU4_LEIMA              ----------------------------------------MKLFFAAVL-----------    9
tr|A4IB15|A4IB15_LEIIN              ----------------------------------------MKLFFAAVL-----------    9
tr|A0A833SEV5|A0A833SEV5_PHYIN      AEKAVTKLCGKKLSTKDNKLCYNLEPLKK-------DVARQVTFKKDTLKICKSLEKKNP    111
sp|Q9CXI5|MANF_MOUSE                IEEELIKFCRE-ARGKENRLCYYIGATDDAATKIINEVSKPLAHHIPVEKICEKLKKKDS    111
sp|Q9XZ63|ARMET_DROME               IETAFKKFCKA-QKNKEHRFCYYLGGLEESATGILNELSKPLSWSMPAEKICEKLKKKDA    111

tr|C1EB43|C1EB43_MICCC              ALLATPQCRAEEHEMPNFKTMKIKELKAILADRGRECKGCAEKADYVAMAADVWSLPIVE    73
tr|A0A0L1KUF0|A0A0L1KUF0_9EUGL      AVSILACHAAHCITDDEIRKMKMKELKEFLDDRDESCSGCVEKGDFVRIAIEVKDKKISQ    67
tr|A0A0S4IZ22|A0A0S4IZ22_BODSA      ALLFSAVTQAFAESEAEFKKMNVRQLKKFLDERDVDYRDITEKSEFVSRAVEWAGRN---    69
tr|Q57VE7|Q57VE7_TRYB2              AVLPLSSFASPELTEADFKRMKIKELRNFLEDRGLTCPGCQEKADFVRVAFTNRAKKPLS    78
tr|A0A1X0NHR2|A0A1X0NHR2_9TRYP      LFYLVCLTAMAEMTEQDFKRMKIKDLRHFLEERDLSCPGCQEKADFVRVAFQNRDKKPVS    75
tr|Q4D5D6|Q4D5D6_TRYCC              LMAFMVTATAAELTEADFKRMKIRELRDFLEDRGLTCPDCQEKADFARYAYQNRDKKPTS    81
tr|Q4CP18|Q4CP18_TRYCC              WMAFMVTAAAAELTEADFKRMKIRELRDFLEDRGLTCPDCQEKADFARYAYQNRDKKPTS    81
tr|A0A3R7N5F9|A0A3R7N5F9_9TRYP      LMVSVGPAAAAELTEADFKRMKIRELRRFLGERGLTCVGCQEKSDFSRYAYQNRDKKPLS    77
tr|A0A422N9T4|A0A422N9T4_TRYRA      QMAFVGPVTAAKLTEADFKRMKIRDLRSFLEERGLTCVGCQEKADFSRYAYQNRDKKPLS    80
tr|S9WF29|S9WF29_9TRYP              FALL--IAVNASLTEDDFKKMKMKDLRIFLSSRGLECVGCLEKSDFVRLAFQNKDKQPLA    68
tr|S9UQX2|S9UQX2_9TRYP              LCLLGAFVSASDMTMADFKKMKIKDLRTFLDERGLECNGCQEKSDFVRVAYEYRDKAPLG    72
tr|S9UU37|S9UU37_9TRYP              LCLLGAFVSASDMTMADFKKMKIKDLRTFLDERGLECNGCQEKSDFVRVAYEYRDKAPLG    107
tr|A0A0M9GAG4|A0A0M9GAG4_LEPPY      --------------------MKVKDLRIFLSDRGLECSGCQEKSDFVRMAHQYRSLNPAG    40
tr|A4HME3|A4HME3_LEIBR              VAILAATLVSAGMTESDFKKMKVKDIRAFLMDRGLECIGCQEKSDFVRMAYQHRDKSPIG    69
```

```
tr|E9AEU4|E9AEU4_LEIMA             IAVFAVILVSAGMTESDFKRMKVKDLRAFLQDRGLECAGCQEKSDFVRMAYQHRDTNPVG    69
tr|A4IB15|A4IB15_LEIIN             IAVFAVTLVSAGMTESDFKKMKVKDLRAFLQDRGLECAGCQEKSDFVRMAYQHRDTNPVG    69
tr|A0A833SEV5|A0A833SEV5_PHYIN     DFCSMRYPVKTDAN-TDYSKMRVKQLRKILAERGVECVGCVEKSDFIAKIKDTESLHTEL    170
sp|Q9CXI5|MANF_MOUSE               QICELKYDKQIDLSTVDLKKLRVKELKKILDDWGEMCKGCAEKSDYIRKINELMPKYAPK    171
sp|Q9XZ63|ARMET_DROME              QICDLRYEKQIDLNSVDLKKLKVRDLKKILNDWDESCDGCLEKGDFIKRIEELKPKYSRS    171
                                                       :.::::: :* . .    .  **.::

tr|C1EB43|C1EB43_MICCC             KPA-SGDETKEKPAEPDLND----------------------------ADQERIRRMMDE    104
tr|A0A0L1KUF0|A0A0L1KUF0_9EUGL     EKQKMKGYTGEYPKKSFWDFWTEESLQIAAS----------SELDTKGRKLIADAVETCF    117
tr|A0A0S4IZ22|A0A0S4IZ22_BODSA     KKV-----VRDLPKEPFWEVWAKISRDKCEAAVATKGLGES---GAKVCDAVASAVDSFF    121
tr|Q57VE7|Q57VE7_TRYB2             EEG-----KREIPKAPLWEVWRDNAKLVCEEAAKKRGLDVTAKPQSDICSAVALVVENFF    133
tr|A0A1X0NHR2|A0A1X0NHR2_9TRYP     EQG-----KREIPNASFWEVWKDNAKALCTEVVQKRGLDVSGKPQADICDAIAYVVENFF    130
tr|Q4D5D6|Q4D5D6_TRYCC             EQG-----KREVPNAPFWEVWRDIAKEVCEEAVKKRGLDVSAEPQSEVCSALAYVTESFF    136
tr|Q4CP18|Q4CP18_TRYCC             EQG-----KREVPNASFWEVWRDIAKEVCEEAVKKRGLDVSAEPQSEVCSALAYVTESFF    136
tr|A0A3R7N5F9|A0A3R7N5F9_9TRYP     EQG-----KRDVPDAPFWAVWRDLAKEVCEGAVRKRGLDVSAAPQSDVCAALAYVTESFF    132
tr|A0A422N9T4|A0A422N9T4_TRYRA     EQG-----KREVPDAPFWVVWRDIAKEVCEEAVKKRGLDVTSTPQSDVCNAIVYVTESFF    135
tr|S9WF29|S9WF29_9TRYP             SAT-----KREIPEGKFWEVWEANAKTKCEEAVKKRGGDPASKPFSDVCHTIEKAVEGYF    123
tr|S9UQX2|S9UQX2_9TRYP             IAA-----KREVPDKKFWEAWGANGKVQCEEAVTKRGSDPAAEPFVHVCDTIEKAIDSFF    127
tr|S9UU37|S9UU37_9TRYP             IAA-----KREVPDKKFWEAWGANGKVQCEEAVTKRGSDPAAEPFVHVCDTIEKAIDSFF    162
tr|A0A0M9GAG4|A0A0M9GAG4_LEPPY     SAE-----KRAVPAKKFWEAWADIAHAECEKAVRLRSNDPTTEPFKSVCSTLRSATDSYL    95
tr|A4HME3|A4HME3_LEIBR             SAK-----KREIPSKKFWEAWSDIAKQECQNAVKRRGNDADAEPFSIICDTIHSAADSYL    124
tr|E9AEU4|E9AEU4_LEIMA             SAV-----KREIPNKKFWEAWGDIAKNECQNAVRRRGNDEGTEPFSTICDTIHSAVDSYL    124
tr|A4IB15|A4IB15_LEIIN             SAV-----KREIPNKKFWEAWGDIAKNECQNAVRRRGNDEGTEPFSTICDTIHSAVDSYL    124
tr|A0A833SEV5|A0A833SEV5_PHYIN     ------------------------------------------------------------    170
sp|Q9CXI5|MANF_MOUSE               AASARTD----L------------------------------------------------    179
sp|Q9XZ63|ARMET_DROME              EL----------------------------------------------------------    173


tr|C1EB43|C1EB43_MICCC             MQNGPRPTGDPERD--AILKKLHSSGIKFSGGEGMPLDQLRNL----------EKAMGNI    152
tr|A0A0L1KUF0|A0A0L1KUF0_9EUGL     MQYGKSVATKLKKGPAELLK-TSLKSPYYQAGIRG-IQKLITLCASSPSLKNA-------    168
tr|A0A0S4IZ22|A0A0S4IZ22_BODSA     MMNGKRTASKLKKKPDALTK-TANGDIYYNAGSRI-IARLLGYCLNAK-NRASCSSSSQV    178
tr|Q57VE7|Q57VE7_TRYB2             MQHGKRVANKLRKNHEALLK-TSYKNVYYDAGHVL-LKRLTEYCLVSEENQNKCSSIGSL    191
tr|A0A1X0NHR2|A0A1X0NHR2_9TRYP     MQHGKRTANKLRKKADDLLK-TSYKNVYYDAGRVL-LERLANYCLASPANQEKCSSVGSL    188
tr|Q4D5D6|Q4D5D6_TRYCC             LQHGKRTAGKLRKKPEALLK-TSFKAVYYDAGRVL-LGRLADHCLASAGNQNTCSSMSKL    194
tr|Q4CP18|Q4CP18_TRYCC             LQHGKRTAGKLRKKPEALLK-TSFKAVYYDAGRVL-LGRLADHCLASAGNQNTCSSMSKL    194
tr|A0A3R7N5F9|A0A3R7N5F9_9TRYP     MQHGKRTASKLRKKPEALLK-TSFKGVYYDAGRLL-LERLANYCLASADHQSTCSSVGKL    190
tr|A0A422N9T4|A0A422N9T4_TRYRA     MQHGKRTANKLRKKPDALLK-TSHKGVYYDSGRLL-LERLVNYCLASADHQVKCSSVSKL    193
tr|S9WF29|S9WF29_9TRYP             MQNGRQVATKLKKQPLQLLK-TSYKDIYYDAGLRL-LEKLSNHCLSTPDQQKKCSSMSNV    181
tr|S9UQX2|S9UQX2_9TRYP             MQHGRQTANRLKKTPHHMLK-TSYKHVYYDVGVRL-LNKLINYCLASPSIQSKCESLSHV    185
tr|S9UU37|S9UU37_9TRYP             MQHGRQTANRLKKTPHHMLK-TSYKHVYYDVGVRL-LNKLINYCLASPSIQSKCESLSHV    220
tr|A0A0M9GAG4|A0A0M9GAG4_LEPPY     MQHGRKVANQLKKTPHHLLQ-TSFKDIYFEAGSHL-FQILADYCLASPAAQENCQSLGAV    153
tr|A4HME3|A4HME3_LEIBR             MQHGRRVANQLKKTPLHLLE-TSFKDVYYEAGLHL-FQTLADYCLASPSSQTACQSLGSV    182
tr|E9AEU4|E9AEU4_LEIMA             MQHGRRVANQLKKTPQDLLR-TSFKDVYFEAGSHL-FQMLADYCLGSPSLQKACQSLGSV    182
tr|A4IB15|A4IB15_LEIIN             MQHGRRVANQLKKTPQDLLK-TSFKDVYFEAGSHL-FQTLADYCLGSPSLQKACQSLGSV    182
tr|A0A833SEV5|A0A833SEV5_PHYIN     ------------------------------------------------------------    170
sp|Q9CXI5|MANF_MOUSE               ------------------------------------------------------------    179
sp|Q9XZ63|ARMET_DROME              ------------------------------------------------------------    173


tr|C1EB43|C1EB43_MICCC             KTKKQGDGDDE--L----------------------------    164
tr|A0A0L1KUF0|A0A0L1KUF0_9EUGL     ----ELQNACEKEFVPWITNVGIENTNPMYEILEQMKSEL--    204
tr|A0A0S4IZ22|A0A0S4IZ22_BODSA     VELMDKDTVKGTGFGAWITNVGIENTNPMYELSNSKSLHDEL    220
tr|Q57VE7|Q57VE7_TRYB2             TTMLES--GKMVDFAKWMTNVGIENTNPMYEVLDGRGDL---    228
tr|A0A1X0NHR2|A0A1X0NHR2_9TRYP     SSLIEG--SSVIDLVKWMTNVGIENTNPMYDFLELRDDL---    225
tr|Q4D5D6|Q4D5D6_TRYCC             MALTEE--SKIADLAKWMTNVGIENTNPMYEFIDQRDDL---    231
tr|Q4CP18|Q4CP18_TRYCC             MALAEE--SKIADLAKWMTNVGIENTNPMYEFIDQRDDL---    231
tr|A0A3R7N5F9|A0A3R7N5F9_9TRYP     MTLAEE--AKVVDLQGWMTNVGIENTNPMYELLDRRDDL---    227
tr|A0A422N9T4|A0A422N9T4_TRYRA     MTLAEE--AKVVDLQGWMTNVGIENTNPMYELLDRRDDL---    230
tr|S9WF29|S9WF29_9TRYP             MKLLE-----DEGFKVYITNVGIENTNPMYDIIDDAADL---    215
tr|S9UQX2|S9UQX2_9TRYP             LGVMEG--NKEGNFKSWITNVGIENTNPMYEILDDASDL---    222
tr|S9UU37|S9UU37_9TRYP             LGVMEG--NKEGNFKSWITNVGIENTNPMYEILDDASDL---    257
tr|A0A0M9GAG4|A0A0M9GAG4_LEPPY     MSAMDG--ACGADFKMWTTNVGIENTNPMYEIIDTRDDL---    190
tr|A4HME3|A4HME3_LEIBR             LSTMDG--SSGANFKVWTTNVGIENTNPMYDIIGEHADL---    219
tr|E9AEU4|E9AEU4_LEIMA             MSAMDG--SSGADFRIWTTNVGIENTNPMYEIIDGRDDL---    219
tr|A4IB15|A4IB15_LEIIN             MSAMDG--SSGADFRIWTTNVGIENTNPMYEIIDGRDDL---    219
tr|A0A833SEV5|A0A833SEV5_PHYIN     ------------------------------------------    170
sp|Q9CXI5|MANF_MOUSE               ------------------------------------------    179
sp|Q9XZ63|ARMET_DROME              ------------------------------------------    173
```

### 05. ER glutathionylspermidine amidase (ERGA)

These proteins, technically CHAP (C51) domain peptidases have a lot of close relatives in bacteria, and even within kinetoplastids. The latter paralogs are the well-known bifunctional trypanothione synthase enzymes (called glutathionylspermidine synthases/amidases). The distinction between the two subfamilies (Type II transmembrane versus soluble, ER-retained) is clear-cut in all kinetoplastid proteomes. Therefore this paralog has likely functionally distinct roles in trypanothione metabolism. The family can be more distantly traced back to prokaryotic D-alanyl-glycine endopeptidases, but it is sporadically found in multiple eukaryotic groups as well (typically without KDEL). It is unclear if its limited occurrance animals (only in rotifer worms) and other groups (e.g. pseudofungi) points to horizontal gene transfer events.

```
tr|A0A815F576|A0A815F576_9BILA      ------------------------------------------------------------    0
tr|A0A814QMX3|A0A814QMX3_9BILA      ------------------------------------------------------------    0
tr|K2MZ59|K2MZ59_TRYCR              ------------------------------------------------------------    0
tr|A0A0G4IUR7|A0A0G4IUR7_PLABS      ------------------------------------------------------------    0
tr|A0A0N0DV74|A0A0N0DV74_LEPPY      ------------------------------------------------------------    0
tr|A4HEJ1|A4HEJ1_LEIBR              ------------------------------------------------------------    0
tr|Q4Q9R0|Q4Q9R0_LEIMA              ------------------------------------------------------------    0
tr|A4I1M7|A4I1M7_LEIIN              ------------------------------------------------------------    0
tr|S9UNH8|S9UNH8_9TRYP              ------------------------------------------------------------    0
tr|S9UZF8|S9UZF8_9TRYP              ------------------------------------------------------------    0
tr|Q57ZC4|Q57ZC4_TRYB2              ------------------------------------------------------------    0
tr|A0A3R7LGJ5|A0A3R7LGJ5_9TRYP      ------------------------------------------------------------    0
tr|Q4CUZ5|Q4CUZ5_TRYCC              ------------------------------------------------------------    0
tr|Q4D7T1|Q4D7T1_TRYCC              ------------------------------------------------------------    0
tr|A0A3R7KMG1|A0A3R7KMG1_TRYRA      ------------------------------------------------------------    0
tr|Q4CYQ4|Q4CYQ4_TRYCC              ------------------------------------------------------------    0
tr|Q4CUZ4|Q4CUZ4_TRYCC              ------------------------------------------------------------    0
tr|Q4CUZ2|Q4CUZ2_TRYCC              ------------------------------------------------------------    0
tr|Q4CYQ2|Q4CYQ2_TRYCC              ------------------------------------------------------------    0
tr|Q4D7T2|Q4D7T2_TRYCC              ------------------------------------------------------------    0
tr|Q4D7T0|Q4D7T0_TRYCC              ------------------------------------------------------------    0
tr|A0A1X0NY88|A0A1X0NY88_9TRYP      ------------------------------------------------------------    0
tr|A0A1X0NYK9|A0A1X0NYK9_9TRYP      ------------------------------------------------------------    0
tr|A0A1X0NRB4|A0A1X0NRB4_9TRYP      ---------------------------------MHNNPPIT-------------------    8
tr|Q4D408|Q4D408_TRYCC              ---------------------------------------MN-------------------    2
tr|A0A422N172|A0A422N172_9TRYP      ---------------------------------------MK-------------------    2
tr|A0A422NKH5|A0A422NKH5_TRYRA      ----------------------------------------M-------------------    1
tr|A0A0S4ITA7|A0A0S4ITA7_BODSA      ------------------------------------------------------------    0
tr|A0A0N0DYP4|A0A0N0DYP4_LEPPY      ---------------------------------------MRNRRSSAPDVCRKNDA----    17
tr|A4HLZ6|A4HLZ6_LEIBR              ---------------------------------------MRSRRTASGAAEGPVGDAAAN    21
tr|Q4Q3Q0|Q4Q3Q0_LEIMA              MFPFAFEYPCEATFAHQMQNKQPSTLLASFFAYYHFLIPMRSRRSPAGDIEDRADVAAAT    60
tr|A4I9C6|A4I9C6_LEIIN              -----------------MKNKQPTTVFASLFAYHHFLIPMRSRQPPSGGVEDHADVAAST    43
tr|A0A0N0DYQ7|A0A0N0DYQ7_LEPPY      ------------------------------------MLPNNMGNSNPGSGGREAGDAAVP    24
tr|A4HLZ4|A4HLZ4_LEIBR              ------------------------------------------------------------    0
tr|Q4Q3Q2|Q4Q3Q2_LEIMA              ------------------------------------------------------------    0
tr|A4I9C4|A4I9C4_LEIIN              -------------------------------------------------------MAAAP    5
tr|Q384E7|Q384E7_TRYB2              ------------------------------------------------------------    0
tr|Q586Y2|Q586Y2_TRYB2              ------------------------------------------------------------    0
tr|Q586Y1|Q586Y1_TRYB2              ------------------------------------------------------------    0
tr|F9W970|F9W970_TRYCI              ------------------------------------------------------------    0
tr|F9W971|F9W971_TRYCI              ------------------------------------------------------------    0
tr|S9UGT3|S9UGT3_9TRYP              ------------------------------------------------------------    0
tr|A0A1X0NR96|A0A1X0NR96_9TRYP      ------------------------------------------------------------    0
tr|A0A1X0NR98|A0A1X0NR98_9TRYP      ------------------------------------------------------------    0
tr|Q4D409|Q4D409_TRYCC              ------------------------------------------------------------    0
tr|Q4D410|Q4D410_TRYCC              ------------------------------------------------------------    0
tr|Q4CKB1|Q4CKB1_TRYCC              ------------------------------------------------------------    0
tr|Q4DEC6|Q4DEC6_TRYCC              ------------------------------------------------------------    0
tr|A0A3R7N0H0|A0A3R7N0H0_9TRYP      ------------------------------------------------------------    0
tr|A0A3R7N2W0|A0A3R7N2W0_9TRYP      ------------------------------------------------------------    0
tr|A0A3S5IRC8|A0A3S5IRC8_TRYRA      ------------------------------------------------------------    0


tr|A0A815F576|A0A815F576_9BILA      ------------------------------------------------------------    0
tr|A0A814QMX3|A0A814QMX3_9BILA      ------------------------------------------------------------    0
tr|K2MZ59|K2MZ59_TRYCR              ------------------------------------------------------------    0
tr|A0A0G4IUR7|A0A0G4IUR7_PLABS      ------------------------------------------------------------    0
tr|A0A0N0DV74|A0A0N0DV74_LEPPY      ------------------------------------------------------------    0
tr|A4HEJ1|A4HEJ1_LEIBR              ------------------------------------------------------------    0
tr|Q4Q9R0|Q4Q9R0_LEIMA              ------------------------------------------------------------    0
tr|A4I1M7|A4I1M7_LEIIN              ------------------------------------------------------------    0
tr|S9UNH8|S9UNH8_9TRYP              ------------------------------------------------------------    0
tr|S9UZF8|S9UZF8_9TRYP              ------------------------------------------------------------    0
tr|Q57ZC4|Q57ZC4_TRYB2              ------------------------------------------------------------    0
tr|A0A3R7LGJ5|A0A3R7LGJ5_9TRYP      ------------------------------------------------------------    0
tr|Q4CUZ5|Q4CUZ5_TRYCC              ------------------------------------------------------------    0
tr|Q4D7T1|Q4D7T1_TRYCC              ------------------------------------------------------------    0
tr|A0A3R7KMG1|A0A3R7KMG1_TRYRA      ------------------------------------------------------------    0
tr|Q4CYQ4|Q4CYQ4_TRYCC              ------------------------------------------------------------    0
tr|Q4CUZ4|Q4CUZ4_TRYCC              ------------------------------------------------------------    0
tr|Q4CUZ2|Q4CUZ2_TRYCC              ------------------------------------------------------------    0
tr|Q4CYQ2|Q4CYQ2_TRYCC              ------------------------------------------------------------    0
tr|Q4D7T2|Q4D7T2_TRYCC              ------------------------------------------------------------    0
tr|Q4D7T0|Q4D7T0_TRYCC              ------------------------------------------------------------    0
```

```
tr|A0A1X0NY88|A0A1X0NY88_9TRYP      ------------------------------------------------------------   0
tr|A0A1X0NYK9|A0A1X0NYK9_9TRYP      ------------------------------------------------------------   0
tr|A0A1X0NRB4|A0A1X0NRB4_9TRYP      ---------------TETSTGPQNSRSSASSYSAT-----GREGANERTPMLGIRPMTV-   47
tr|Q4D408|Q4D408_TRYCC              ---------------SAVN------ASPSVTVSNL-----SAPPPNECTPMLRTRPVPM-   35
tr|A0A422N172|A0A422N172_9TRYP      ---------------RAVA------AP-------P-----PAAAPNERTPMLHPRPLPV-   28
tr|A0A422NKH5|A0A422NKH5_TRYRA      ---------------RAVE------VSPPATASKE-----AAAASTERTPLLKFHPRPI-   34
tr|A0A0S4ITA7|A0A0S4ITA7_BODSA      ---------------------------------------------------MAYNSTKVD   9
tr|A0A0N0DYP4|A0A0N0DYP4_LEPPY      --SDVRS------------------SSPVVGG----------------------------   29
tr|A4HLZ6|A4HLZ6_LEIBR              APHRLRS------------------PSPPSASSS--------------------------   37
tr|Q4Q3Q0|Q4Q3Q0_LEIMA              VLHSSSS------------------SSSAAAAAS--------------------------   76
tr|A4I9C6|A4I9C6_LEIIN              VLRSSAS------------------SSSAAAAAS--------------------------   59
tr|A0A0N0DYQ7|A0A0N0DYQ7_LEPPY      FLQRIRSESDTAEGNAAVSVQRNHSSSPHTATACYGGTETHALR--------KRNPKVSD   76
tr|A4HLZ4|A4HLZ4_LEIBR              ---------------------------------------------------MGGATRFV-   8
tr|Q4Q3Q2|Q4Q3Q2_LEIMA              ------------------------------------------------------------   0
tr|A4I9C4|A4I9C4_LEIIN              LSRTF---------------------SPKSSESSFSSHSQERMEQNYRESAAGFGPPPA-   43
tr|Q384E7|Q384E7_TRYB2              ---------------------------------------------MAKGS----------   5
tr|Q586Y2|Q586Y2_TRYB2              ---------------------------------------------MAARK----------   5
tr|Q586Y1|Q586Y1_TRYB2              ---------------------------------------------MEREP----------   5
tr|F9W970|F9W970_TRYCI              ------------------------------------------------------------   0
tr|F9W971|F9W971_TRYCI              ---------------------------------------------MQRET----------   5
tr|S9UGT3|S9UGT3_9TRYP              ------------------------------------------------------------   0
tr|A0A1X0NR96|A0A1X0NR96_9TRYP      ------------------------------------------------------------   0
tr|A0A1X0NR98|A0A1X0NR98_9TRYP      ---------------------------------------------MQRDT-ELWAVPRK-   13
tr|Q4D409|Q4D409_TRYCC              ------------------------------------------------------------   0
tr|Q4D410|Q4D410_TRYCC              ------------------------------------------------------------   0
tr|Q4CKB1|Q4CKB1_TRYCC              ------------------------------------------------------------   0
tr|Q4DEC6|Q4DEC6_TRYCC              ------------------------------------------MGTLRENN-AGFGDEKM-   16
tr|A0A3R7N0H0|A0A3R7N0H0_9TRYP      ------------------------------------------MDTTKGDH-AHFGEEKM-   16
tr|A0A3R7N2W0|A0A3R7N2W0_9TRYP      ------------------------------------------------------------   0
tr|A0A3S5IRC8|A0A3S5IRC8_TRYRA      ------------------------------------------------------------   0


tr|A0A815F576|A0A815F576_9BILA      ------------------------------------------------------------   0
tr|A0A814QMX3|A0A814QMX3_9BILA      ------------------------------------------------------------   0
tr|K2MZ59|K2MZ59_TRYCR              ------------------------------------------------------------   0
tr|A0A0G4IUR7|A0A0G4IUR7_PLABS      ------------------------------------------------------------   0
tr|A0A0N0DV74|A0A0N0DV74_LEPPY      ------------------------------------------------------------   0
tr|A4HEJ1|A4HEJ1_LEIBR              -----------------------------------------------------------M   1
tr|Q4Q9R0|Q4Q9R0_LEIMA              ------------------------------------------------------------   0
tr|A4I1M7|A4I1M7_LEIIN              ------------------------------------------------------------   0
tr|S9UNH8|S9UNH8_9TRYP              ------------------------------------------------------------   0
tr|S9UZF8|S9UZF8_9TRYP              ------------------------------------------------------------   0
tr|Q57ZC4|Q57ZC4_TRYB2              -----------------------------------------------------------M   1
tr|A0A3R7LGJ5|A0A3R7LGJ5_9TRYP      ------------------------------------------------------------   0
tr|Q4CUZ5|Q4CUZ5_TRYCC              ------------------------------MLPVCIFF-----------FFLQAGMWISF   19
tr|Q4D7T1|Q4D7T1_TRYCC              ------------------------------------------------------------   0
tr|A0A3R7KMG1|A0A3R7KMG1_TRYRA      -------------------------------MPVSSLYSADGQGNKRCWLSFSAGAWCST   29
tr|Q4CYQ4|Q4CYQ4_TRYCC              -----------------------------------------------------------M   1
tr|Q4CUZ4|Q4CUZ4_TRYCC              -----------------------------------------------------------M   1
tr|Q4CUZ2|Q4CUZ2_TRYCC              ------------------------------------------------------------   0
tr|Q4CYQ2|Q4CYQ2_TRYCC              ------------------------------------------------------------   0
tr|Q4D7T2|Q4D7T2_TRYCC              -----------------------------------------------------------M   1
tr|Q4D7T0|Q4D7T0_TRYCC              ------------------------------------------------------------   0
tr|A0A1X0NY88|A0A1X0NY88_9TRYP      ------------------------------------------------------------   0
tr|A0A1X0NYK9|A0A1X0NYK9_9TRYP      -----------------------------------------------------------M   1
tr|A0A1X0NRB4|A0A1X0NRB4_9TRYP      --------------------------------------------EIDSTTVR---EDYYE   60
tr|Q4D408|Q4D408_TRYCC              --------------------------------------------EAESMLRR---EGQDE   48
tr|A0A422N172|A0A422N172_9TRYP      --------------------------------------------EVGSMQLR---EEMDE   41
tr|A0A422NKH5|A0A422NKH5_TRYRA      --------------------------------------------EVGSMQQQ---EEMDE   47
tr|A0A0S4ITA7|A0A0S4ITA7_BODSA      SHDEEDSNDDFVVGV--KM-----KKAPY--------------PWVDDNHHE-----RTS   43
tr|A0A0N0DYP4|A0A0N0DYP4_LEPPY      ---------------AARIPPRRHKELDVEDAALCDAYAINLRRESEDME-------LRR   67
tr|A4HLZ6|A4HLZ6_LEIBR              -------------LLHSASLFCARKESDSSNVDVCDAYAIRLRNEAVDAV-------RYR   77
tr|Q4Q3Q0|Q4Q3Q0_LEIMA              -------------MMQDQVLRCVRKEPGPSNAAICDAYAMRLLQEATEEE-------RCR   116
tr|A4I9C6|A4I9C6_LEIIN              -------------LLQDEVLPRLRKESGPSNAAICDAYATRLRQEATEEE-------RRR   99
tr|A0A0N0DYQ7|A0A0N0DYQ7_LEPPY      SGERSSKDDGDVVVVPSRAGGGGGGSSGNNSRPDAGRDVEGECIVVNDNGTDTTADTCMG   136
tr|A4HLZ4|A4HLZ4_LEIBR              -SSGASTEDDDVVVRHSRRYGGATIQRNTS----CEIDNERTLFVQQENKRTDSGVACFG   63
tr|Q4Q3Q2|Q4Q3Q2_LEIMA              ------------------------------------------------------------   0
tr|A4I9C4|A4I9C4_LEIIN              -ESASNAANGDVVVCQSRSHRETTAQPNTS----GNNYGESTRVGARDNKSADSGIASAG   98
tr|Q384E7|Q384E7_TRYB2              ------DYDAVAYEPMPREGFDAGDEGPL----------------VGNHDQGF--LFDDQ   41
tr|Q586Y2|Q586Y2_TRYB2              ----------A------------GTARPLLQREGEAR--------------NF----DS-   24
tr|Q586Y1|Q586Y1_TRYB2              ----------AVTSKTNRLSSNEKDPHDVPETNDGAI--------------NG--SFNNR   39
tr|F9W970|F9W970_TRYCI              -------------------------MTPRANTEGCDT--------------LL--G-DDR   18
tr|F9W971|F9W971_TRYCI              ----------EGRPKRSRASSERQDINDILQEDNCNN--------------DY--TTDLK   39
tr|S9UGT3|S9UGT3_9TRYP              ------------------------------------------------------------   0
tr|A0A1X0NR96|A0A1X0NR96_9TRYP      ------------------------------MAAQRKRKESGATRPVNSSK-KT--IMSSF   27
tr|A0A1X0NR98|A0A1X0NR98_9TRYP      ----------DVREETPF--PEEEEVNVT-IPNGEDLKTEANEEPVDSLE-KR--V----   53
tr|Q4D409|Q4D409_TRYCC              ------------------MLPHG--------------------ARMRTSKESC--FFTGL   20
tr|Q4D410|Q4D410_TRYCC              ------------------------------------------------------------   0
tr|Q4CKB1|Q4CKB1_TRYCC              ---------------------------------------------MQSNDSQD--TF---   10
tr|Q4DEC6|Q4DEC6_TRYCC              -----DHSNNSIRVSTPRMQPGEAELVNSLDLNGPTNEFAGEQVEMQSNDSQD--TF---   66
tr|A0A3R7N0H0|A0A3R7N0H0_9TRYP      -----DHAHNHGFEPTTEMHPREMYVSDEHNTHKEANAFVSEQVEINSNEAD-----EAM   66
tr|A0A3R7N2W0|A0A3R7N2W0_9TRYP      ------------------MSPREATKRTRQA------------PRVNKGAAT-------S   23
tr|A0A3S5IRC8|A0A3S5IRC8_TRYRA      ------------------MSLRQVKKRTRQV------------SCVNKGSFT-------R   23


tr|A0A815F576|A0A815F576_9BILA      ----------------------------------------MKIVL--LIVLFTIIYRY--   16
tr|A0A814QMX3|A0A814QMX3_9BILA      ------------------------------------------------------------   0
tr|K2MZ59|K2MZ59_TRYCR              ------------------------------------------------------------   0
```

```
tr|A0A0G4IUR7|A0A0G4IUR7_PLABS      ------------------------------------------------------------   0
tr|A0A0N0DV74|A0A0N0DV74_LEPPY      ---------------------------------MNIVRLA-FCCITAALLSLGALFYF--   24
tr|A4HEJ1|A4HEJ1_LEIBR              HRHKERTCTHIDSRHIAIKYPPPIRTH---TLIMKLGYVV-VFCACAALLSLGSLFYY--   55
tr|Q4Q9R0|Q4Q9R0_LEIMA              ---------------------------------MNLVRVS-LLCACTTLLCLSALYYY--   24
tr|A4I1M7|A4I1M7_LEIIN              ---------------------------------MNLVRVA-LLCACTTLLCLGALYYY--   24
tr|S9UNH8|S9UNH8_9TRYP              -----------------------MRIS---QVFP-------FLAGFLLIVCVSSVVYF--   25
tr|S9UZF8|S9UZF8_9TRYP              ----------------------MKVT---HLFA--------LTATAACILLIYTTRF--   24
tr|Q57ZC4|Q57ZC4_TRYB2              VSL-----NRK--QKFQLAVPAT------------------LLLFLFVYVLLGGGS----   32
tr|A0A3R7LGJ5|A0A3R7LGJ5_9TRYP      ------------------------MKS---KSIC-------KVVPLVCALLCGGYLLW--   24
tr|Q4CUZ5|Q4CUZ5_TRYCC              FFFFLGYCLSR--SQKQHTMVAGFVEN---KSLG-------RVACLAFVLICGVLLLW--   65
tr|Q4D7T1|Q4D7T1_TRYCC              -------------------MAAGFMEN---KSLG-------RVACLACVLICGVLLLW--   29
tr|A0A3R7KMG1|A0A3R7KMG1_TRYRA      MYRIR-----K--QALGLSTP-AFLVF---PLLM-------LLLLLLGYVMLGGR-----   66
tr|Q4CYQ4|Q4CYQ4_TRYCC              IHTAR-----K--KQFGLSAL-A--------LFV-------LLLFLLVCVTLGLH-----   33
tr|Q4CUZ4|Q4CUZ4_TRYCC              IHTAR-----K--KQFGLSAL-A--------LFV-------LLLFLLVCVTLGLH-----   33
tr|Q4CUZ2|Q4CUZ2_TRYCC              ------------------------------------------------------------   0
tr|Q4CYQ2|Q4CYQ2_TRYCC              ------------------------------------------------------------   0
tr|Q4D7T2|Q4D7T2_TRYCC              IHTAR-----K--KQFGLSAL-A--------LFV-------LLLFLLVCITLGLH-----   33
tr|Q4D7T0|Q4D7T0_TRYCC              ------------------------------------------------------------   0
tr|A0A1X0NY88|A0A1X0NY88_9TRYP      --------------------MP-ALV-----------------ILTLLVYIIIGIGK----   19
tr|A0A1X0NYK9|A0A1X0NYK9_9TRYP      MM--------K--STGEIAVL-ALL-----------------LLLAIYYLLISDANIL--   31
tr|A0A1X0NRB4|A0A1X0NRB4_9TRYP      EPLWHRLS-----------SP-LHPVFRKVHDHCHYEWIAALLSVITLLLLFVFVFYY--   106
tr|Q4D408|Q4D408_TRYCC              MPFWERVS-----------SP-LHPVFRKVQEHCHYEWVAALLSAIMLLTLFVFIFYL--   94
tr|A0A422N172|A0A422N172_9TRYP      VPYWERIY-----------SP-LHPVLSKVQEHCHYEWVAALLSAMMLLALFVFVFYL--   87
tr|A0A422NKH5|A0A422NKH5_TRYRA      MSFWERVS-----------SP-LHPVLNKVQEHCHYEWVAALLSAIMLVALFVIVFYL--   93
tr|A0A0S4ITA7|A0A0S4ITA7_BODSA      SSWLPRSL-----------RD-GILVC-------IPACALLLLVVL-FLPH---------   74
tr|A0A0N0DYP4|A0A0N0DYP4_LEPPY      PSFWRIRC-----------SS-SIFVG-------LFIWALIGLVVLFKVS----------   98
tr|A4HLZ6|A4HLZ6_LEIBR              RSYWHVHC-----------SP-SILVA-------LLIWVTIGLVVYTKLD----------   108
tr|Q4Q3Q0|Q4Q3Q0_LEIMA              GPHWRARC-----------SP-SILVA-------LLIWATIGLVVYMKVD----------   147
tr|A4I9C6|A4I9C6_LEIIN              GPHWRARC-----------SP-SILVA-------LLIWATIALVVYMKVD----------   130
tr|A0A0N0DYQ7|A0A0N0DYQ7_LEPPY      KVRHFLLN-----------SD-SALYA-------LLKWGLAVLLLF-FLSFVVFGMIY--   174
tr|A4HLZ4|A4HLZ4_LEIBR              AVRRFVLT-----------RD-PLLYG-------LLKWGLSVFLLL-FLTFILLGLFF--   101
tr|Q4Q3Q2|Q4Q3Q2_LEIMA              -------------------------------------------------------MLY--   3
tr|A4I9C4|A4I9C4_LEIIN              VARCFVLT-----------HD-PLLYG-------ILKWGLATLLLL-FLAFVVFGMLY--   136
tr|Q384E7|Q384E7_TRYB2              ADGAHSTQ-----------LT-YIAPSVK-----KPVYALGITLLF-LLVAVVLTVIL-Q   82
tr|Q586Y2|Q586Y2_TRYB2              ----YKL------------------YA-------GASFAVGVVIIL-CLMKVPRA----E   50
tr|Q586Y1|Q586Y1_TRYB2              KRGIKRLC--------------SIAQA-------IFAWTFGVLVLL-FISFIAFGVLYSG   77
tr|F9W970|F9W970_TRYCI              EVVRRKSC--------------PA-YF-------LL--SFFLVILL-AMFYHAASPLWKE   53
tr|F9W971|F9W971_TRYCI              DKWRSRVC--------------STASF-------AFAWAFGVLIVL-FLVFIAFGVLYSG   77
tr|S9UGT3|S9UGT3_9TRYP              ------------------------------------MKWSFHFLLL--FFLIIIVGMLY--   21
tr|A0A1X0NR96|A0A1X0NR96_9TRYP      AKPFKSLP--------------YYIIS-------LFIWGTVASLVCMLLLAKSSREEYSH   66
tr|A0A1X0NR98|A0A1X0NR98_9TRYP      ----DRAT--------------SVLAV-------IIGWTLCVLVV-LFLGFIVFGILY--   85
tr|Q4D409|Q4D409_TRYCC              PTWFYSLS--------------HFFVA-------LGIWGSITCMVL-FLLTTSLRSKK-S   57
tr|Q4D410|Q4D410_TRYCC              ------------------------------------------------------------   0
tr|Q4CKB1|Q4CKB1_TRYCC              KKRMSRIN--------------AILTS-------VVIWSFLLLTLL-FLASIVFGIVY-S   47
tr|Q4DEC6|Q4DEC6_TRYCC              KKRMSRIN--------------AILTS-------VVIWSFLLLTLL-FLASIVFGIVY-S   103
tr|A0A3R7N0H0|A0A3R7N0H0_9TRYP      RKKFSRVN-----------AT-LTMVA-------GWSFGVLTLLFL-AFIV--FG-IL-Y   102
tr|A0A3R7N2W0|A0A3R7N2W0_9TRYP      MLWCPSLP--------------YCVIA-------VGIWGLLACLMA-LLLTRTSHLSR-L   60
tr|A0A3S5IRC8|A0A3S5IRC8_TRYRA      RLRWPSLS--------------CCIIA-------PGILGLLACLVA-LLHT-MCPIRQ-L   59


tr|A0A815F576|A0A815F576_9BILA      SKCDIVTQLKNF---------DSNLNSINARAFGNSTLAAYGAMVGIA-STNVPAYSNGN   66
tr|A0A814QMX3|A0A814QMX3_9BILA      -------------------------------MSNNVNTCVPFNEIEGIA-STNVPAYSNKK   29
tr|K2MZ59|K2MZ59_TRYCR              ---------------------MDNTDTQRGVHASETEHVSFGGVIGVT-DSGVPVFSNGE   38
tr|A0A0G4IUR7|A0A0G4IUR7_PLABS      ----------MW---------WPAVIVVAAVVVGADARAPFGDVLGVS-HGGVQVRSCHT   40
tr|A0A0N0DV74|A0A0N0DV74_LEPPY      STYDPEKHM------------LMRARVYSDYDPLEHCVTPFGVVLGYS--NGVPAFSNCI   70
tr|A4HEJ1|A4HEJ1_LEIBR              SVYDYQKHM------------SMVQRTYSTYDPLTDCAAPFGQLLGVA--DDVPAYSNCN   101
tr|Q4Q9R0|Q4Q9R0_LEIMA              SMYDYEKHM------------NMVQRKYSVYDPLTDCATPFGQLLGVA--DDVPAYSNCN   70
tr|A4I1M7|A4I1M7_LEIIN              SVYDYEKHM------------SMVQRKYSGYDPLTDCATPFGQLLGVA--DDVPAYSNCN   70
tr|S9UNH8|S9UNH8_9TRYP              ARDDYERHL------------LNQHKKYDTFDPLTNCTTPFDTLLGIA--DHVPAYSNCH   71
tr|S9UZF8|S9UZF8_9TRYP              AWSDYRPAV------------GTREREYDAYDPLTDCVTPFGTVIGYV--GPVPVLSNCN   70
tr|Q57ZC4|Q57ZC4_TRYB2              -----TQKS-------------KRHGMHMDFNPLEHCNAPMDSVVGYS--SDVPSMSNCH   72
tr|A0A3R7LGJ5|A0A3R7LGJ5_9TRYP      GWRGSTGGA-------------PSRETDPHFDPLEHCSTDLGTLLGYA--FSVPAFSNCH   69
tr|Q4CUZ5|Q4CUZ5_TRYCC              GWRGNKRNP-------------LSAEGNPLFDPLKHCRTRYGTILGYA--LSVPAFSNCH   110
tr|Q4D7T1|Q4D7T1_TRYCC              GWRGNKRNP-------------LSAEGDPLFDPLKHCRTRYGTILGYA--LSVPAFSNCH   74
tr|A0A3R7KMG1|A0A3R7KMG1_TRYRA      -WWRHTAGS-------------EKNSAQPSFNPLDHCGTPYGTLLGYA--HGIPAFSNCH   110
tr|Q4CYQ4|Q4CYQ4_TRYCC              -WWRDTSGP-------------ALHPPRSRFDPLAHCATPHGTLLGYA--LDIPAFSNCH   77
tr|Q4CUZ4|Q4CUZ4_TRYCC              -WWRDTSGP-------------ALHPPRSRFDPLAHCATPHGTLLGYA--LDIPAFSNCH   77
tr|Q4CUZ2|Q4CUZ2_TRYCC              ------------------------------------------------------------   0
tr|Q4CYQ2|Q4CYQ2_TRYCC              ------------------------------------------------------------   0
tr|Q4D7T2|Q4D7T2_TRYCC              -WWRDTSGP-------------ALHPPRSRFDPLAHCATPHGTLLGYA--LGIPAFSNCH   77
tr|Q4D7T0|Q4D7T0_TRYCC              ------------------------------------------------------------   0
tr|A0A1X0NY88|A0A1X0NY88_9TRYP      -RSNFFKAE-------------VQMPRRPVNHGLLDCVTPHSTLLGYA--SGIPAYSNCH   63
tr|A0A1X0NYK9|A0A1X0NYK9_9TRYP      GLNRYYNT--------------HLPSKDPRFDPLKHCTTPFGTFLGVA--HGIPAYSNCH   75
tr|A0A1X0NRB4|A0A1X0NRB4_9TRYP      ------N----------------------TPPVPVNCVTPFGQLLGEN--SGVMVFSNCR   136
tr|Q4D408|Q4D408_TRYCC              ------D----------------------TSPRPEHCVTPFGHLLGES--RGVKAFSNCN   124
tr|A0A422N172|A0A422N172_9TRYP      ------D----------------------TPPRPEHCVTPFGELLGES--KGVFAFSNCN   117
tr|A0A422NKH5|A0A422NKH5_TRYRA      ------D----------------------TPPRPVHCVTPFGELLGES--NGVLAFSNCN   123
tr|A0A0S4ITA7|A0A0S4ITA7_BODSA      ---NCRGGSM----------------SLREQRSSSGCDTAFGSVIGVSAVGSVTAYSNCN   115
tr|A0A0N0DYP4|A0A0N0DYP4_LEPPY      -GSAQVEAAST---ALIPAEAVLPDGCRGDYCFEEGGTEPFGAVLGAH--DGVYAYSNCY   152
tr|A4HLZ6|A4HLZ6_LEIBR              -SFHLRNGAAPTTSSAETVHSQLPDGCLDHHCLKDGGSELFGAVLGAH--NGVFAYSNCN   165
tr|Q4Q3Q0|Q4Q3Q0_LEIMA              -SSHLRPGAALATTSSVETPLSQLPESCRGHYCLQEGGKEPFGAVLGAH--NGVFAYSNCN   204
tr|A4I9C6|A4I9C6_LEIIN              -NSRLRPGAAPDTTSSVETPLSQLPESCRGHYCLREGGKEPFGAVLGAH--DGVFAYSNCN   187
tr|A0A0N0DYQ7|A0A0N0DYQ7_LEPPY      SEVRYNGNC--T---------VDAGVTDTPA-LPEGCESPFGAILGAF--QNTFAHSNCN   220
tr|A4HLZ4|A4HLZ4_LEIBR              NVFFYRGNC--E---------NTNETTKGKTDPKKECVTPFGAILGVY--DGVFGYSNCN   148
tr|Q4Q3Q2|Q4Q3Q2_LEIMA              SSIRYGRNC--T---------GTNE----TVKPLGPCVTPLGTILGVF--NGVFGYSNCN   46
tr|A4I9C4|A4I9C4_LEIIN              SSIRYGGNC--T---------GTNGTSGGTVKPREPCVTPFSTILGVF--NGVFGYSNCN   183
tr|Q384E7|Q384E7_TRYB2              P----------------------------TFLYGETCVTPFGTIIGVH--NGVFAYSNCR   112
tr|Q586Y2|Q586Y2_TRYB2              NSKDTVST----------------NTGSLSVGCQQNCSAPFGNVLGIY--NGVPAMSNCN   92
```

```
tr|Q586Y1|Q586Y1_TRYB2             IR--YPE-G--------------------SGKVEKHCLNPVGAILGAH--EGVFSYSNCG    112
tr|F9W970|F9W970_TRYCI             KGDDYLRTGA--------------GIVDVPPGCRRGCATPFGSVLGEY--NGVKGMSNCD    97
tr|F9W971|F9W971_TRYCI             --I-------------------------RYRGGSNTCSNSFGAITGTH--SGVFSYSNCG    108
tr|S9UGT3|S9UGT3_9TRYP             SGVHYQGSTATV---------DTNDSSSGTTLDQSSCHTQFGSIMGVH--QGVFAYSNCN    70
tr|A0A1X0NR96|A0A1X0NR96_9TRYP     ETINYNSS-HSL-----------------NNSCTVDCATTFGTILGIH--NGIFAYSNCN    106
tr|A0A1X0NR98|A0A1X0NR98_9TRYP     SGIHYHGDKWHE-----------------DSDGQEKCSTPFGAIIGVH--NDVFAYSNCD    126
tr|Q4D409|Q4D409_TRYCC             ETSVAKTGGTFL-----------------PEGCRKGCDTTFGAILGIV--NGVVSYSNCN    98
tr|Q4D410|Q4D410_TRYCC             ------------------------------FFGESACVSPYGAILGVA--GDVFAYSNCI    28
tr|Q4CKB1|Q4CKB1_TRYCC             GIHY-HGPGWHD-----------------ASSAKSACVSPYGAILGVA--GDVFAYSNCI    87
tr|Q4DEC6|Q4DEC6_TRYCC             GIHY-HGPGWHD-----------------ASSAKSACVSPYGAILGAA--GDVFAYSNCI    143
tr|A0A3R7N0H0|A0A3R7N0H0_9TRYP     SGVHYHGESWHN-----------------VNSNTPECVTPFGSIIGVT--NGVFAYSNCN    143
tr|A0A3R7N2W0|A0A3R7N2W0_9TRYP     DTVHEGNDTYRL-----------------SEGCSIGCDTTFGTILGVA--NGVFGFSNCN    101
tr|A0A3S5IRC8|A0A3S5IRC8_TRYRA     DTLVEGNGTSPF-----------------HEGCSIGCDTTFGSILGIS--NGVFGYSNCN    100


tr|A0A815F576|A0A815F576_9BILA     DSYISNEPNYL---------------------YNIYTGMRWQCVEYARRWLFIRKGCV--    103
tr|A0A814QMX3|A0A814QMX3_9BILA     SQYFRCGRHYV---------------------YGVFTGYQWQCVEFARRWLLMRKSCT--    66
tr|K2MZ59|K2MZ59_TRYCR             KNFFSERISLAPDCRHIEAAT-----STTVIGTYDTLGYQWLCLEFTRRYLLMTKGVW--    91
tr|A0A0G4IUR7|A0A0G4IUR7_PLABS     ESPEAYREALRAG-----DPK-----YHAFL-GDVFTGFQWQCVELARRYLVVNAGVA--    87
tr|A0A0N0DV74|A0A0N0DV74_LEPPY     STFKTTYVNYVNLMNPLDVGR---RGDPSET-TVIMLGVRYTAMDFYIRWMAWNRGLVP-    125
tr|A4HEJ1|A4HEJ1_LEIBR             TKFTSTYINYVNLMDPMDNGR---RGDPSEM-RVIMTAYRYSAFDYYMRWLVWNRGIMP-    156
tr|Q4Q9R0|Q4Q9R0_LEIMA             TKFSSTYINYVNLMDPMDNGR---RGDPSET-RIVMTAYRYTAFDYCMRWLVWNRGVMP-    125
tr|A4I1M7|A4I1M7_LEIIN             TKFSSTYINYVSLMDPMDNGR---RGDPSET-RIVMTAYRYTAFDYCMRWLVWNRGVMP-    125
tr|S9UNH8|S9UNH8_9TRYP             TRFNMERVSYCNLWNPLDVGR---TADPADGVPRVLVGEHFTANDYVNRWLIFNRGVLI-    127
tr|S9UZF8|S9UZF8_9TRYP             RRFKSDITNFVNHFNPYDMEQ---PYNPGEPVQRSMTSTAYDCLDFVSRYMVYYRGLIV-    126
tr|Q57ZC4|Q57ZC4_TRYB2             RHWLSETYAYTTIGYPHEVMK---KMPYGSEWKRAPTGLCWTADEFVARYLLHTRGIFV-    128
tr|A0A3R7LGJ5|A0A3R7LGJ5_9TRYP     RSYKAMRLVVTRFGRPDFVLD---YN--DGAAGNYFSGTPWLSTEYVARFLFHHRGVTH-    123
tr|Q4CUZ5|Q4CUZ5_TRYCC             SSYSAERLVITRFGYPELVAD---HN--DGSKGNYCSGSPWSSMEYTARILFHYKGIKY-    164
tr|Q4D7T1|Q4D7T1_TRYCC             RSYSAERLVITRFGHPELVAD---HN--DGSKGNYCSGSPWSSMEYTARILFHYKGIKY-    128
tr|A0A3R7KMG1|A0A3R7KMG1_TRYRA     RLWKTHVYAFVALSHPHDVRH---APFSLGTLHSRGTGTCWDSAEYIVRFFYLIRGMSF-    166
tr|Q4CYQ4|Q4CYQ4_TRYCC             RRWNSKSQAFVTLEHPHDVRQ---EPFLLGTLRSRSTGVRWVAAEFIARTYYLHRGMSF-    133
tr|Q4CUZ4|Q4CUZ4_TRYCC             RRWNSKSQAFVTLEHPHDVRQ---EPFLLGTLRSRSTGVRWVAAEFIARTYYLHRGMSF-    133
tr|Q4CUZ2|Q4CUZ2_TRYCC             --------------------------FLLGTLRSRSTGVRWVAAEFIARTYYLHRGMSF-    33
tr|Q4CYQ2|Q4CYQ2_TRYCC             ------------------------------------------------------MSF-    3
tr|Q4D7T2|Q4D7T2_TRYCC             RRWNSKSQAFVTLEHPHDVRQ---EPFLLGTLRSRSTGIRWVAAEFIARTYYLHRGMSF-    133
tr|Q4D7T0|Q4D7T0_TRYCC             ------------------------------------------------------MSF-    3
tr|A0A1X0NY88|A0A1X0NY88_9TRYP     RLWSSDVMAFASLAFPDDVLK---EPFPLGDRRQLSTGKCWVAAEYVSRFYYLMRGLVF-    119
tr|A0A1X0NYK9|A0A1X0NYK9_9TRYP     RSFRSNRLILTRFGSPDVFLE---HS--SDQEGSFISGSPWAATEYVARFLFHTKGITI-    129
tr|A0A1X0NRB4|A0A1X0NRB4_9TRYP     RDYQNMEEHYL-----------------FIKSLRVYSGVKWKSVEYARRFWLIAKHPG-V    178
tr|Q4D408|Q4D408_TRYCC             RDYRNVEEHFV-----------------LVRTSLIYSGLKWRSVEYARRFWLLAKRPA-V    166
tr|A0A422N172|A0A422N172_9TRYP     RDYRNVEEHFV-----------------LVKTSPVHSGLKWHSVEYARRFWLLAKWPA-V    159
tr|A0A422NKH5|A0A422NKH5_TRYRA     RDYRDVEEHFV-----------------LVKTSQVHSGLKWRSVEYARRFWLLAKQPA-V    165
tr|A0A0S4ITA7|A0A0S4ITA7_BODSA     SDYISNLSNFVNVTIPGGAGG-------SVVVPDVYTGMPWQCVEYARRYWATT--SPYT    166
tr|A0A0N0DYP4|A0A0N0DYP4_LEPPY     AHSCVSFVD-FEYPIPLPPGAHTPLDDPQATTRLMRTGMKWQCVEYARRYWMLRGTPAPA    211
tr|A4HLZ6|A4HLZ6_LEIBR             SNTCISHLE-HQMEIPLPPGSRTTLDAPHATTRPMKTGMKWQCVEYARRYWMLHGKPTPA    224
tr|Q4Q3Q0|Q4Q3Q0_LEIMA             SDTCISSLQ-YQMAIPLPPGARTALDAPHATTRLMTTGMKWQCVEYARRYWMLRGTPTPA    263
tr|A4I9C6|A4I9C6_LEIIN             SDTCTSLLK-YQMAIPLPPGARTALDAPHATTRLMTTGMKWQCVEFARRYWMLRGKPTPA    246
tr|A0A0N0DYQ7|A0A0N0DYQ7_LEPPY     GYYVSTVDSYMNFTVPVMDAA---TGMLTMSSKEFYTGLAWQCVEYARRYWMINGRPKPA    277
tr|A4HLZ4|A4HLZ4_LEIBR             DDYESTKYKYVNLMVPVLNNK---TGKVEYTSKSFFTGLEWQCVEFARRFWMLSGKPEGA    205
tr|Q4Q3Q2|Q4Q3Q2_LEIMA             DSYVSTELRYINLTVPELNNE---TGQLTYISKQFYTGLAWQCVEYARRYWMQRGTPQPA    103
tr|A4I9C4|A4I9C4_LEIIN             DSYVSTELGYINLTVLELNNE---TGQVTYISKQFYTGLAWQCVEYARRYWMQSGTPQPA    240
tr|Q384E7|Q384E7_TRYB2             RDHISTKKN-S-----------------IKGEGSMETGMEWQCFEYVKRYWIMRGVPQSV    154
tr|Q586Y2|Q586Y2_TRYB2             SDSCTAELW-NTVKVEDIRIP---AGRVDPHAVPPIYGMQWQCVEYARRYWMLRGTPQPA    148
tr|Q586Y1|Q586Y1_TRYB2             ATENTTTYN-N----------------VTVAGTSYQSGLKWQCVEYARRYWMLRGTPQPA    155
tr|F9W970|F9W970_TRYCI             SDTCIADHW-HTIDVSELRDA------IRMETFPSSYGVKWQCVEYARRYWVLRGTPKPA    150
tr|F9W971|F9W971_TRYCI             MDVNAIEHH-N----------------VTVEGRTQTTGVKWQCVEYARRYWVLRGTPKPA    151
tr|S9UGT3|S9UGT3_9TRYP             SDYISEDDS-VI---------------KVSGAKNLYAGLRWQCVEYARRYWMLVGTPVPA    114
tr|A0A1X0NR96|A0A1X0NR96_9TRYP     STTCISNEW-HELEDPQISIA---FKGNT--TIGNSTGMKWQCVEYARRYWMLRGSPVPA    160
tr|A0A1X0NR98|A0A1X0NR98_9TRYP     EDY-ISKDY-NTLAY----------------TALSNTGLKWQCVEYARRYWMLRGSPVPA    168
tr|Q4D409|Q4D409_TRYCC             SSTCTSNLW-HPMSSLLAHAD---YNNENRWNNTNTSGLEWQCVEFARRYWMLRGTPVRA    154
tr|Q4D410|Q4D410_TRYCC             RNYDSK-------------------SRNNVSSQNKESGLEWDCMEFARRYWMLRGTPVRA    69
tr|Q4CKB1|Q4CKB1_TRYCC             RNYDSK-------------------SRNNVSSQNKESGLEWDCMEFARRYWMLRGTPVRA    128
tr|Q4DEC6|Q4DEC6_TRYCC             RNYDSK-------------------SRNNVSSQNKESGLKWDCMEFARRYWMLRGTPVRA    184
tr|A0A3R7N0H0|A0A3R7N0H0_9TRYP     EDYSSKENS-----TLL-------------GKDKLFTGLGWQCVEFARRYWMLRGTPVPA    185
tr|A0A3R7N2W0|A0A3R7N2W0_9TRYP     ATTCISNIW-HPMHALMGPAG---ISSSSGIGPGNMSGIEWQCVEFARRYWMLRGTPVPA    157
tr|A0A3S5IRC8|A0A3S5IRC8_TRYRA     ATTCISNKW-HPIHTFMTSGG---FTGRNILAPENTSGIKWQCVEFARRYWMLRGSPVPA    156


tr|A0A815F576|A0A815F576_9BILA     -FDSVDAANDMWSQIFKVQRVVDG----KCFSLKQYQNG------------------ST    139
tr|A0A814QMX3|A0A814QMX3_9BILA     -FKDIPCASNIWTDLSYIERITDG----QHFSLHSIPNG------------------SS    102
tr|K2MZ59|K2MZ59_TRYCR             -LASIPTAEDIWEMDEVFISVPTG----TPVPMEREKHG-----------------ETT    128
tr|A0A0G4IUR7|A0A0G4IUR7_PLABS     -FDDVVNAYQIWDLDGVRR-VADG----ERVAWRRFANG------------------SR    122
tr|A0A0N0DV74|A0A0N0DV74_LEPPY     -R-LVEDTSQFWKTNSFYN-PA--------KTAQDWEAVYITNYAEATD-----VEERRF    169
tr|A4HEJ1|A4HEJ1_LEIBR             -R-LVENTNQLWKRVDYFN-PA--------KPEQDWSAEYITNYEEVTS-----LEERKF    200
tr|Q4Q9R0|Q4Q9R0_LEIMA             -R-LVENTNQLWKTVDYFN-PA--------RPEQGWSAEYITNYEEVTD-----VEERKF    169
tr|A4I1M7|A4I1M7_LEIIN             -R-LVENTNQLWKTVDYFN-PA--------KPEQGWSAEYITNYEEVTG-----MEERKF    169
tr|S9UNH8|S9UNH8_9TRYP             -S-LFPYYSRYVSEASFAY-PT--------PTPSHRMTVRLHNKQRALD-----IEDRKL    171
tr|S9UZF8|S9UZF8_9TRYP             -S-MHHIFYEQWEYGSFIN-PAR--------PEVRRETEKYANYKVAST-----LDERKR    170
tr|Q57ZC4|Q57ZC4_TRYB2             -YFGGSRHDLYWENLKFFG-SS--------GMHRLYERINFENKVEVTT-----TKRRKR    173
tr|A0A3R7LGJ5|A0A3R7LGJ5_9TRYP     -L-PADTPQDVWTSKYFYN-PLEG-E--VGGKRRRYEPVRIANYDAATT-----AKLRKR    172
tr|Q4CUZ5|Q4CUZ5_TRYCC             -F-EMPTPQEVWSTPYFFN-PLEA-A--ERDKERRYEPVRVANYEEATT-----AKERKR    213
tr|Q4D7T1|Q4D7T1_TRYCC             -F-ETPTPQEVWSTPYFFN-PLEA-A--ERDKERRYEPVRVANYEEAAT-----AKERKR    177
tr|A0A3R7KMG1|A0A3R7KMG1_TRYRA     -IYSTRDVVSHWSTLEFHG-GI--------ATDRSYEAIRLLNAAEATT-----VKLRKR    211
tr|Q4CYQ4|Q4CYQ4_TRYCC             -IYGAPDAPEYWLALEFHG-PM--------VREREYETLELLNSAEATT-----AKERKR    178
tr|Q4CUZ4|Q4CUZ4_TRYCC             -IYGAPDAPEYWLALEFHG-PM--------VREREYETLELLNSAEATT-----AKERKR    178
tr|Q4CUZ2|Q4CUZ2_TRYCC             -IYGAPDAPEYWLALEFHG-PM--------VREREYETLELLNSAEATT-----AKERKR    78
tr|Q4CYQ2|Q4CYQ2_TRYCC             -IYGAPDAPEYWLALEFHG-PM--------VREREYETLELLNSAEATT-----AKERKR    48
tr|Q4D7T2|Q4D7T2_TRYCC             -IYGAPDAPEYWSALEFHG-PM--------VREREYETLELLNSAEATT-----AKERKH    178
```

```
tr|Q4D7T0|Q4D7T0_TRYCC              -IYGAPDAPEYWSALEFHG-PM--------VREREYETLELLNSAEATT-----AKERKH    48
tr|A0A1X0NY88|A0A1X0NY88_9TRYP      -YYDSAEVRQFWSTIEFYS-TE--------GDKRKYEVIELPNYKEVKT-----AKERKR    164
tr|A0A1X0NYK9|A0A1X0NYK9_9TRYP      -H-DPGTTQEIWSIRYFYN-PLST-K--HSDNLRKYEPIQLENFMEVKT-----AKERKR    178
tr|A0A1X0NRB4|A0A1X0NRB4_9TRYP      AFEYVNSPVDIWTYVETAK-DVNG----KDLNLRKYPNSLTGKSISNISKDM--KKQWLA    231
tr|Q4D408|Q4D408_TRYCC              QFMSVQNAEDIWTRVVYAS-YITG----ERVKLFKYENMLACDQTPTKNTGM-KKSLWKH    220
tr|A0A422N172|A0A422N172_9TRYP      QFMSVRNAEDIWTRVNYAS-YTNG----KRVKLFKYENMLSCDRAPALNATM-KKHAWNS    213
tr|A0A422NKH5|A0A422NKH5_TRYRA      QFMSVTCAEDIWTRVDYAY-YING----KRVKLLKYENMMACDQAPLSKTTM-KKGQWKH    219
tr|A0A0S4ITA7|A0A0S4ITA7_BODSA      LFGSVDGASDIWTSLHHGT-FIESSDVVVQFDLQKFENGGV----------------NAT    209
tr|A0A0N0DYP4|A0A0N0DYP4_LEPPY      VFGAVEGAADMWTDLSFVT-LLDN-V--TTAPLWKYTNG--------------APVGSGG    253
tr|A4HLZ6|A4HLZ6_LEIBR              LFGTVVGAADIWHSIHSVT-FLDN-K--TTAPLLKFQNG--------------AKLGYGG    266
tr|Q4Q3Q0|Q4Q3Q0_LEIMA              FFGAVKGAADIWDSLTHVT-FLDN-A--TTAPLLKFKNG--------------ARLGYGG    305
tr|A4I9C6|A4I9C6_LEIIN              FFGPVVGAADIWDTLTHVT-FLDN-A--TTAPLLKFKNG--------------ARLGYGG    288
tr|A0A0N0DYQ7|A0A0N0DYQ7_LEPPY      YFGSVIGAADIWN-LTEVH-LLENTS--RTLPLRKYKSGDR-------------VVQDGL    320
tr|A4HLZ4|A4HLZ4_LEIBR              YFDSVVGAADIWN-LSFVR-LVSNTS--ATLPLQKYSNGGR-------------L-SDGL    247
tr|Q4Q3Q2|Q4Q3Q2_LEIMA              YFDSVLGAADVWN-LTFVR-LLSNAS--ITLPLRRYWNGDR-------------V-TDNH    145
tr|A4I9C4|A4I9C4_LEIIN              YFESVPGAADIWN-LTFVR-LLSNTS--MKLPLHRFRNGDP-------------V-TSNL    282
tr|Q384E7|Q384E7_TRYB2              ILPTARKSSELWS-FTQAT-FKNG-S--KV-QLERHD--------------------NGG    188
tr|Q586Y2|Q586Y2_TRYB2              TFGSVDGAADIWD-LKDVQ-LLNG-Q--KRKPLLKYHNG---------------NATSAN    188
tr|Q586Y1|Q586Y1_TRYB2              TFGSVDGAADIWD-LKDVQ-LLNG-Q--KRKPLLKYHNG---------------NATSAN    195
tr|F9W970|F9W970_TRYCI              AFGSVEGAADIWD-LDNVA-LLDG-T--T-RPLLKFQNE---------------NATAAG    189
tr|F9W971|F9W971_TRYCI              AFGSVEGAADIWD-LDNVA-LLDG-T--T-RPLLKFQNE---------------NATAAG    190
tr|S9UGT3|S9UGT3_9TRYP              YFGSINGAADIWA-LTIVY-TVSD-A--TTLPLHRFESMTKKDYESAQQTPSSAMGNITN    169
tr|A0A1X0NR96|A0A1X0NR96_9TRYP      TFSAVEGAADIWE-LQSVQ-LVNG-A--KT-PLLKYSNG--------------LSISAGG    200
tr|A0A1X0NR98|A0A1X0NR98_9TRYP      TFSSVEGAADIWE-LQSVQ-LVNG-A--KT-PLLKYSNG--------------LSISAGG    208
tr|Q4D409|Q4D409_TRYCC              TFDSVVGAADIWA-LNSVR-LLDG-S--KT-PLLKYPNG--------------LPRRDGG    194
tr|Q4D410|Q4D410_TRYCC              TFDSVVGAADIWA-LNSVR-LLDG-S--KT-PLLKYPNG--------------LPRRDGG    109
tr|Q4CKB1|Q4CKB1_TRYCC              TFDSVVGAADIWA-LNSVR-LLDG-S--KT-PLLKYPNG--------------LPRRDGG    168
tr|Q4DEC6|Q4DEC6_TRYCC              TFDSVVGAADIWA-LNFVR-LLDG-S--KT-PLLKYPNG--------------LPRRDGG    224
tr|A0A3R7N0H0|A0A3R7N0H0_9TRYP      SFEKVAGAADIWS-LTSVQ-LLDG-S--TK-PLLKYANN--------------VSARDGG    225
tr|A0A3R7N2W0|A0A3R7N2W0_9TRYP      SFEKVAGAADIWS-LTSVQ-LLDG-S--TT-PLLKYANN--------------VSARAGG    197
tr|A0A3S5IRC8|A0A3S5IRC8_TRYRA      SFATVEGAADIWA-LTSVQ-LLNG-S--TA-PLLKYANG--------------VSVRAGG    196


tr|A0A815F576|A0A815F576_9BILA      SPPKNESLLIYSLGQD-----M-PFGHVAVIVDVLK------------------------    169
tr|A0A814QMX3|A0A814QMX3_9BILA      KSPKKDSLLIYRRSKK-----M-TYGHVAIITDVTS------------------------    132
tr|K2MZ59|K2MZ59_TRYCR              NMPSVGDLLVWSRTED-----F-PYGHLAVVTHVSD------------------------    158
tr|A0A0G4IUR7|A0A0G4IUR7_PLABS      VRPEIGSLLIWRPAGF-----YERTGHVAVVVAVGD------------------------    153
tr|A0A0N0DV74|A0A0N0DV74_LEPPY      NAPRKADAILYNQDSK-----ILPDGHIAVIVKVEGDVEA--AGGTEKF---NELKKLRL    219
tr|A4HEJ1|A4HEJ1_LEIBR              NAPRRADAVIYRMDEK-----TLPAGHIAVVVKVEDDIEA--AGGLEKL---KDLKKMRL    250
tr|Q4Q9R0|Q4Q9R0_LEIMA              NAPRRGDAIVYRMDKN-----TIPAGHMAVVVKVEDDVEA--AGGPEKL---NELKKMRL    219
tr|A4I1M7|A4I1M7_LEIIN              NAPRRADAIVYRMDKN-----TIPAGHIAVVVKVEDDVEA--AGGPEKL---KELKKMRL    219
tr|S9UNH8|S9UNH8_9TRYP              NAPRRGDVVVFDAMYTEGQPHYLPSGHIAVVVKVENNLEHPDMQDKEKK---NKLIRERL    228
tr|S9UZF8|S9UZF8_9TRYP              FAPRVGDVIVYNADPA----MDLSAGHLAVVVRVEADMDDPVAKDKERL---RQLREQRL    223
tr|Q57ZC4|Q57ZC4_TRYB2              HTPLVGDVVVWDSDYK----AYFPRGHVAVVVKVEDDVSA--AGGEAAL---RELKKERR    224
tr|A0A3R7LGJ5|A0A3R7LGJ5_9TRYP      LAPRFADIVVWPAQSE----EGLPGGHVAVVVQVEDDVAA--AGGADRL---RALRRERL    223
tr|Q4CUZ5|Q4CUZ5_TRYCC              LAPRVFDIVVWPAQME----HELPEGHIAVVVQVEDDVEA--AGGEDRL---RELRKLRL    264
tr|Q4D7T1|Q4D7T1_TRYCC              LAPRVFDIVVWPAQME----HELPEGHIAVVVQVEDDVEA--AGGEDRL---RELRKLRL    228
tr|A0A3R7KMG1|A0A3R7KMG1_TRYRA      LAPKVADIVVWAAQRE----NDLPEGHLAVVVQVEHDVEA--AGGEDSL---RELQKERL    262
tr|Q4CYQ4|Q4CYQ4_TRYCC              LAPRVFDVVVWSPMEE----RDLPEGHIAVVVQVEDDVEA--AGGEDRL---RELRKLRL    229
tr|Q4CUZ4|Q4CUZ4_TRYCC              LAPRVFDVVVWSPMEE----RDLPEGHIAVVVQVEDDVEA--AGGEDRL---RELRKLRL    229
tr|Q4CUZ2|Q4CUZ2_TRYCC              LAPRVFDVVVWSPMEE----RDLPEGHIAVVVQVEDDVEA--AGGEDRL---RELRKLRL    129
tr|Q4CYQ2|Q4CYQ2_TRYCC              LAPRVFDVVVWSPMEE----RDLPEGHIAVVVQVEDDVEA--AGGEDRL---RELRKLRL    99
tr|Q4D7T2|Q4D7T2_TRYCC              LAPRVLDVVVWSPMEE----RDLPEGHIAVVVRVEDDVEA--AGGEDRL---RELRKLRL    229
tr|Q4D7T0|Q4D7T0_TRYCC              LAPRVLDVVVWSPMEE----RDLPEGHIAVVVQVEDDVEA--AGGEDRL---RELRKLRL    99
tr|A0A1X0NY88|A0A1X0NY88_9TRYP      LAPKFADVVVWASHPE----RELDEGHAAVVVHVEDDVEA--AGGEAQL---REMQKEHT    215
tr|A0A1X0NYK9|A0A1X0NYK9_9TRYP      LAPKFADVVVWASHPE----RELDDGHAAVVVHVEDDVEA--AGGEAQL---REMQKEHT    229
tr|A0A1X0NRB4|A0A1X0NRB4_9TRYP      SKPQVGDLLIYDHSK------KLPEGHVAVIVNVVPVSNI--NNG----------NDIFV    273
tr|Q4D408|Q4D408_TRYCC              YKPQVGDLLVYANDS------SIPGGHVAVVVGIVSMPSN--SGA----------NNSSM    262
tr|A0A422N172|A0A422N172_9TRYP      YRPQIGDLLVYANES------RLPGGHVAVVVGIASLPLQ--AAA----------NASRI    255
tr|A0A422NKH5|A0A422NKH5_TRYRA      YGPQIGDLLVYANDS------RLPGGHVAVVVGRVSQPPH--AVS----------NTSWI    261
tr|A0A0S4ITA7|A0A0S4ITA7_BODSA      SPPQVGDLFIYPIQP-----GGFPFGHVTVVLNDQLPPTL--NNG------------GND    250
tr|A0A0N0DYP4|A0A0N0DYP4_LEPPY      SAPRVGDLLIYPRDTD----GKFPFGHVAVIVGVELPGER--PQATTNSDE-----AHAA    302
tr|A4HLZ6|A4HLZ6_LEIBR              SAPRVGDLLIYPRDTL----GNFPFGHVAVVVGVEMPANA--EANDSYTDAEMAAAQLRQ    320
tr|Q4Q3Q0|Q4Q3Q0_LEIMA              SAPRVGDLLIYPRDTE----NVFPYGHVAVVVKVEMPTKA--EADDSYMDAGAASPKPRQ    359
tr|A4I9C6|A4I9C6_LEIIN              SAPRVGDLLIYPRDAE----GFFSYGHVAVVVRVEMTTKA--EADDSYMDAAVTSSKPRQ    342
tr|A0A0N0DYQ7|A0A0N0DYQ7_LEPPY      QPPQAGDIIIYPVQS-----GGFPVGHVAVVTKVEMGE----------------------    353
tr|A4HLZ4|A4HLZ4_LEIBR              QAPAPGDIIIYPVQG-----GGFPYGHVAVITKVDMAV----------------------    280
tr|Q4Q3Q2|Q4Q3Q2_LEIMA              QIPAIGDIIIYPVQD-----GGFPYGHVAVIANVELST----------------------    178
tr|A4I9C4|A4I9C4_LEIIN              QVPAVGDIIIYPVQD-----GGFPYGHVAVIASVEIST----------------------    315
tr|Q384E7|Q384E7_TRYB2              PQPLVGDLLVYREQP-----ALLPVGHVAVIVRVGK------------------------    219
tr|Q586Y2|Q586Y2_TRYB2              SKPRVGDLLIYPRQP-----NGFPYGHVAVVAGVTG------------------------    219
tr|Q586Y1|Q586Y1_TRYB2              SKPRVGDLLIYPRQP-----NGFPCGHVAVVAGVTG------------------------    226
tr|F9W970|F9W970_TRYCI              SRPRAGDLLIYPRQP-----NGFPYGHVAVVAGVEE------------------------    220
tr|F9W971|F9W971_TRYCI              SRPRAGDLLIYPRQP-----NGFPYGHVAVVAGVEE------------------------    221
tr|S9UGT3|S9UGT3_9TRYP              TVPRVGDILIYKREL-----PSFPYGHVAVVVEFVAGATT--AG------------AAAA    210
tr|A0A1X0NR96|A0A1X0NR96_9TRYP      SAPRVGDLLIYPRQK-----GGFVYGHVAVIVDVLS------------------------    231
tr|A0A1X0NR98|A0A1X0NR98_9TRYP      SAPRVGDLLIYPRQK-----GGFVYGHVAVIVDVLS------------------------    239
tr|Q4D409|Q4D409_TRYCC              SAPRAGDLLIYPRQR-----NDFPFGHVAVVVGVTK------------------------    225
tr|Q4D410|Q4D410_TRYCC              SAPRAGDLLIYPRQR-----NDFPFGHVAVVVGVTK------------------------    140
tr|Q4CKB1|Q4CKB1_TRYCC              SAPRAGDLLIYPRQR-----NDFPFGHVAVVVGVTK------------------------    199
tr|Q4DEC6|Q4DEC6_TRYCC              SAPRAGDLLIYPRQR-----EDFPFGHVAVVVGVTK------------------------    255
tr|A0A3R7N0H0|A0A3R7N0H0_9TRYP      SAPRVGDLLIYPRQE-----GGFPFGHVAVVVAVKS------------------------    256
tr|A0A3R7N2W0|A0A3R7N2W0_9TRYP      SAPRVGDLLIYPRQE-----GGFPFGHVAVVVAVKS------------------------    228
tr|A0A3S5IRC8|A0A3S5IRC8_TRYRA      SAPRIGDLLVYPKQG-----DGLPFGHVAVVVSVKS------------------------    227
                                    *   . .::                 ** :::

tr|A0A815F576|A0A815F576_9BILA      --DSIRVAEQNYHAYYWS--G---------------NYSRQIPYVVKNGN-------YY    202
tr|A0A814QMX3|A0A814QMX3_9BILA      --DYVYIAEQNNLYHYWP--G---------------DYARRERLRFDNGN-------YY    165
```

```
tr|K2MZ59|K2MZ59_TRYCR               --KRVWVAEQNYEFKRWQPGK----------------NYSRRFDCEKREDG-------VT    193
tr|A0A0G4IUR7|A0A0G4IUR7_PLABS       --AFVDIVEQNVDDTVWEPGQ----------------TYSRRLKARSDAGS-------FT    188
tr|A0A0N0DV74|A0A0N0DV74_LEPPY       HPRRVYVAEQNFKNEDWG-GK----------------NYSRILTFQWRQVKT-GTTYEGF    261
tr|A4HEJ1|A4HEJ1_LEIBR               HPRRVYVAEQNCQNQPWG-EH----------------NYSRVLQFKWRAVSE-M-AHEGY    291
tr|Q4Q9R0|Q4Q9R0_LEIMA               HPRRVYVAEQNWKNQPWG-GH----------------NYSRVLQFKWRAVSE-K-AHEGG    260
tr|A4I1M7|A4I1M7_LEIIN               HPRRVYVAEQNWKNQPWG-GH----------------NYSRVLQFKWRAVAE-K-AYEGC    260
tr|S9UNH8|S9UNH8_9TRYP               QPRRVYIAEQNLYNRHWE-GK----------------NYSRVMDFYWEPTAN-EKEFRAI    270
tr|S9UZF8|S9UZF8_9TRYP               HPRKVYIAEQNFGNADWG-GK----------------NYTRICRFVWEPVRGATGLYESI    266
tr|Q57ZC4|Q57ZC4_TRYB2               QPSLVYIAEQNFDNKNWE-GR----------------NFSRVLKFTWMRGD------RAS    261
tr|A0A3R7LGJ5|A0A3R7LGJ5_9TRYP       QPQLVYIAEQNFDNAPWA-GR----------------NYSRVLRFYWENGK------EAV    260
tr|Q4CUZ5|Q4CUZ5_TRYCC               QPRLLYIAEQNFDNTHWG-GK----------------NYSRVLRFQWRNGR------EAV    301
tr|Q4D7T1|Q4D7T1_TRYCC               QPRLLYIAEQNFDNTHWG-CK----------------NYSRVLRFQWRNGR------EAV    265
tr|A0A3R7KMG1|A0A3R7KMG1_TRYRA       QPQLVYIAEQNFDNVLWD-GK----------------NYSRVLRFYWKNGK------EAM    299
tr|Q4CYQ4|Q4CYQ4_TRYCC               QPRLLYIAEQNFDNTHWG-GK----------------NYSRVLRFQWRNGR------EAV    266
tr|Q4CUZ4|Q4CUZ4_TRYCC               QPRLLYIAEQNFDNTHWG-GK----------------NYSRVLRFQWRNGR------EAV    266
tr|Q4CUZ2|Q4CUZ2_TRYCC               QPRLLYIAEQNFDNTHWG-GK----------------NYSRVLRFQWRNGR------EAV    166
tr|Q4CYQ2|Q4CYQ2_TRYCC               QPRLLYIAEQNFDNTHWG-GK----------------NYSRVLRFQWRNGR------EAV    136
tr|Q4D7T2|Q4D7T2_TRYCC               QPRLLYIAEQNFDNTHWG-CK----------------NYSRVLRFQWRNGR------EAV    266
tr|Q4D7T0|Q4D7T0_TRYCC               QPRLLYIAEQNFDNTHWG-CK----------------NYSRVLRFQWRNGR------EAV    136
tr|A0A1X0NY88|A0A1X0NY88_9TRYP       QPQLVYIAEQNFDNKDWE-GK----------------NYSRVLRFSWKNGK------EML    252
tr|A0A1X0NYK9|A0A1X0NYK9_9TRYP       QPQLVYIAEQNFDNKDWE-GK----------------NYSRVLRFSWKNGK------EML    266
tr|A0A1X0NRB4|A0A1X0NRB4_9TRYP       KYYNVYLAEQNWDNRPWV-EEEDLVTSSD-NNEPRALYYSRMVLLKEDMTT-----HRMT    326
tr|Q4D408|Q4D408_TRYCC               TQYSMLLAEQNWNNEPWP-HRSNLEGKDTRNDTRLLGYYSREVSLQEVESA-----QMVC    316
tr|A0A422N172|A0A422N172_9TRYP       RQYSLLLAEQNWDNARWL-CNDTLEGGT-STVTALPCQYSRKVLLQQVDSA-----RRMC    308
tr|A0A422NKH5|A0A422NKH5_TRYRA       RQCSLLLAEQNWDNARWL-YNETLRGRDTPTSTAVPLHYSRKVLLQEVDST-----RKMC    315
tr|A0A0S4ITA7|A0A0S4ITA7_BODSA       VTSEVHVGEQNWDSFQWQ-HA--------------AEGYSRKLQLTYHQLS-----RRWS    290
tr|A0A0N0DYP4|A0A0N0DYP4_LEPPY       REGRAYIAEQNWHSSPWP-EP--------------YHNYSRWLPLEVTATPQ-GTSVQYT    346
tr|A4HLZ6|A4HLZ6_LEIBR               RRGLVYIAEQNWDSVPWP-KP--------------YHNYSRSLPLVVLESAE-GQPLQYT    364
tr|Q4Q3Q0|Q4Q3Q0_LEIMA               RHSHVYVAEQNWDSVTWP-NP--------------YHNYSRSLPLVVLESAE-GRPLQYT    403
tr|A4I9C6|A4I9C6_LEIIN               RHGLVYLAEQNWDSATWP-NP--------------YHNYSRSLPLVVLESAE-GLPLQYT    386
tr|A0A0N0DYQ7|A0A0N0DYQ7_LEPPY       -QGFIYVAEQNWESTQWS-GP--------------FYNYSRKIPLRYDPLT-----TAII    392
tr|A4HLZ4|A4HLZ4_LEIBR               -NGAIYVAEQNWGSAMWV-DP--------------HHNYSRRIPLMYDMTT-----SSVT    319
tr|Q4Q3Q2|Q4Q3Q2_LEIMA               -HGAIYVAEQNWANAVWS-SP--------------HHNYTRRIPMFYDMLT-----STIT    217
tr|A4I9C4|A4I9C4_LEIIN               -HGAIYVAEQNWANAVWP-SA--------------YHNYSRRIPLYYEMLT-----STIT    354
tr|Q384E7|Q384E7_TRYB2               --THVWVAEQNWFNKQWH-PP--------------FHNFSRTIKMHHNAES-----QTYE    257
tr|Q586Y2|Q586Y2_TRYB2               --DRMFVAEQNWENTAWP-GP--------------YHNYSRVLNLSCNPNG-----TACT    257
tr|Q586Y1|Q586Y1_TRYB2               --DRMFVAEQNWENAAWP-GP--------------YHNYSRVLNLSCNPNG-----TACT    264
tr|F9W970|F9W970_TRYCI               --RRLFVAEQNWDNQQWP-GP--------------YHNYSRELKLTCDASS-----TRCS    258
tr|F9W971|F9W971_TRYCI               --RRLFVAEQNWDNQQWP-GP--------------YHNYSRELKLTCDASS-----TRCS    259
tr|S9UGT3|S9UGT3_9TRYP               SAPYVLVGEQNWDNEMWP-SP--------------YHNYSRRVNMVYDASK-----EVYT    250
tr|A0A1X0NR96|A0A1X0NR96_9TRYP       --AAVLVAEQNWDNTVWP-GP--------------FHNYSRSIPMQYDANS-----KAYT    269
tr|A0A1X0NR98|A0A1X0NR98_9TRYP       --AAVLVAEQNWDNTVWP-GP--------------FHNYSRSIPMQYDANS-----KAYT    277
tr|Q4D409|Q4D409_TRYCC               --NSVLVAEQNWDNKMWP-GP--------------YHNHSREIRMLYSPIH-----DAYN    263
tr|Q4D410|Q4D410_TRYCC               --NSVLVAEQNWDNKMWP-GP--------------YHNHSREIRMLYSPIH-----DAYN    178
tr|Q4CKB1|Q4CKB1_TRYCC               --NSVLVAEQNWDNKMWP-GP--------------YHNHSREIRMLYSPIH-----DAYN    237
tr|Q4DEC6|Q4DEC6_TRYCC               --NSVLVAEQNWDNKMWP-GP--------------YHNHSREIRMLYSPIH-----DAYN    293
tr|A0A3R7N0H0|A0A3R7N0H0_9TRYP       --ESLLVAEQNWDNKVWP-AP--------------HHNYSREIPMHHNTKE-----DIYE    294
tr|A0A3R7N2W0|A0A3R7N2W0_9TRYP       --ESLLVAEQNWDNKVWP-AP--------------HHNYSREIPMHHNTKE-----DIYE    266
tr|A0A3S5IRC8|A0A3S5IRC8_TRYRA       --ESLLVAEQNWDNNVWP-AP--------------HHDYSREIPMHHNVTE-----DTYK    265
                                           : ***     *                     .:*

tr|A0A815F576|A0A815F576_9BILA       IMD--SYKIYGWMSVEDNNQN---YPLNQSTINKIMEKNIS--FPNFICSKSVTFNQLKP    255
tr|A0A814QMX3|A0A814QMX3_9BILA       IDD--EDPIYGWMEIENNDEL---KPFDESNINNILQKYLE--FKSMDGV----------    208
tr|K2MZ59|K2MZ59_TRYCR               VFV-GELHLLGWVSIK--------APSYDFSLGDLPDKYRHILGPGHIVRRHLEREFLLP    244
tr|A0A0G4IUR7|A0A0G4IUR7_PLABS       VVDDYDGDVIGWIIMSIVTTE--RAPAYDDESDMDRDEL---------------------    225
tr|A0A0N0DV74|A0A0N0DV74_LEPPY       LTDPDSLSIIGFVRVGKPLPLRELSDPYDDALRDD-GGDL--------------------    300
tr|A4HEJ1|A4HEJ1_LEIBR               YVDPDGLDIIGCMRVGKSMPLRAVPDPYQETLDMENDGDL--------------------    331
tr|Q4Q9R0|Q4Q9R0_LEIMA               YVDPDELDIIGVVRVGKAMPLRAAPDPYEEALNMDNDGDL--------------------    300
tr|A4I1M7|A4I1M7_LEIIN               YVDPDGLGIIGVVRVGKAMPLRAAPDPYQEALDMDNDGDL--------------------    300
tr|S9UNH8|S9UNH8_9TRYP               LQDNDGLTVLGRIRAGRRVPTSENKPSYQLGLQEAAKEVKEKYGEEVEVVTPTDQE----    326
tr|S9UZF8|S9UZF8_9TRYP               LKDPDSLSIIGRLRIGRELSLTEEEPSAEEKGLQAMDEDKMKHKQGLK------------    314
tr|Q57ZC4|Q57ZC4_TRYB2               LEDPDGPPLMGHVRVGKLLED--ASFFGDL------------------------------    289
tr|A0A3R7LGJ5|A0A3R7LGJ5_9TRYP       LQDPSGARVLGLVRKGKLALNLEDDDAGDL------------------------------    290
tr|Q4CUZ5|Q4CUZ5_TRYCC               LQDPLGLKVLGLVRIGKRTVIFEDEDGGDL------------------------------    331
tr|Q4D7T1|Q4D7T1_TRYCC               LQDPLGLKVLGLVRIGKRTVIFEDEDGGDL------------------------------    295
tr|A0A3R7KMG1|A0A3R7KMG1_TRYRA       LHDSSGPQELGFVRKGQACRK--PQGRGCS------------------------------    327
tr|Q4CYQ4|Q4CYQ4_TRYCC               LQDPLGLPIIGRVRPGRPKQI---------------------------------------    287
tr|Q4CUZ4|Q4CUZ4_TRYCC               LQDPLGLPIIGRVRPGRPKQN--TDVEDDL------------------------------    294
tr|Q4CUZ2|Q4CUZ2_TRYCC               LQDPLGLPIIGRVRPGRPKQN--TDVEDDL------------------------------    194
tr|Q4CYQ2|Q4CYQ2_TRYCC               LQDPLGLPIIGRVRPGRPKQN--TDVEDDL------------------------------    164
tr|Q4D7T2|Q4D7T2_TRYCC               LQDPLELPIIGRVRPGKPKQN--TDVEDDL------------------------------    294
tr|Q4D7T0|Q4D7T0_TRYCC               LQDPLELPIIGRVRPGKPKQN--TDVEDDL------------------------------    164
tr|A0A1X0NY88|A0A1X0NY88_9TRYP       LEDSDDYNIIGRVRPGKVKRD--DMDEGDL------------------------------    280
tr|A0A1X0NYK9|A0A1X0NYK9_9TRYP       LEDSDDYNIIGRVRPGNLKRD--DMDEGDL------------------------------    294
tr|A0A1X0NRB4|A0A1X0NRB4_9TRYP       IEDAGG-MVLGWVRV---------------------------------------------    340
tr|Q4D408|Q4D408_TRYCC               VKDPWG-TVLGWVRA---------------------------------------------    330
tr|A0A422N172|A0A422N172_9TRYP       VTDPGG-TVLGWVRV---------------------------------------------    322
tr|A0A422NKH5|A0A422NKH5_TRYRA       VKDPRG-TVLGWVRV---------------------------------------------    329
tr|A0A0S4ITA7|A0A0S4ITA7_BODSA       VYDPQG-KIAGWVRRPSQRS----------------------------------------    309
tr|A0A0N0DYP4|A0A0N0DYP4_LEPPY       LHDKYH-AILGWMRYGEP------------------------------------------    363
tr|A4HLZ6|A4HLZ6_LEIBR               IEDSFH-GVQGWARYDDVP-----------------------------------------    382
tr|Q4Q3Q0|Q4Q3Q0_LEIMA               IEDSLH-GIQGWVRYDDEP-----------------------------------------    421
tr|A4I9C6|A4I9C6_LEIIN               IEDSLH-GIQGWVRYDDDP-----------------------------------------    404
tr|A0A0N0DYQ7|A0A0N0DYQ7_LEPPY       LNDPDG-KIIGWMRYG--------------------------------------------    407
tr|A4HLZ4|A4HLZ4_LEIBR               LNDPNG-IIIGWMRYG--------------------------------------------    334
tr|Q4Q3Q2|Q4Q3Q2_LEIMA               LDDSEH-QIIGWMRYG--------------------------------------------    232
tr|A4I9C4|A4I9C4_LEIIN               LDDPKG-KIIGWMRYG--------------------------------------------    369
tr|Q384E7|Q384E7_TRYB2               LEDMAGTTIKGWMRYKT-------------------------------------------    274
```

```
tr|Q586Y2|Q586Y2_TRYB2              VREKDNVTVQGWVRYE--------------------------------------------   273
tr|Q586Y1|Q586Y1_TRYB2              VREKDNVTVQGWVRYE--------------------------------------------   280
tr|F9W970|F9W970_TRYCI              VHEEGNIVVQGWVRYE--------------------------------------------   274
tr|F9W971|F9W971_TRYCI              VHEEGNIVVQGWVRYE--------------------------------------------   275
tr|S9UGT3|S9UGT3_9TRYP              LVDVEG-TLYGWMRYG--------------------------------------------   265
tr|A0A1X0NR96|A0A1X0NR96_9TRYP      LKESDDIIIQGWMRYSK-------------------------------------------   286
tr|A0A1X0NR98|A0A1X0NR98_9TRYP      LKESDDIIIQGWMRYSK-------------------------------------------   294
tr|Q4D409|Q4D409_TRYCC              ITEEENIIIDGWMRYAT-------------------------------------------   280
tr|Q4D410|Q4D410_TRYCC              ITEEENIIIDGWMRYAT-------------------------------------------   195
tr|Q4CKB1|Q4CKB1_TRYCC              ITEEEN------------------------------------------------------   243
tr|Q4DEC6|Q4DEC6_TRYCC              ITEEENIIIDGWMRYTT-------------------------------------------   310
tr|A0A3R7N0H0|A0A3R7N0H0_9TRYP      VAEADKRTITGWVRYAE-------------------------------------------   311
tr|A0A3R7N2W0|A0A3R7N2W0_9TRYP      VAEADKRTITGWVRYAE-------------------------------------------   283
tr|A0A3S5IRC8|A0A3S5IRC8_TRYRA      VIEADNVIVTGWVRYAM-------------------------------------------   282


tr|A0A815F576|A0A815F576_9BILA      IVSHHNDIRFWGKEV--AHEYISQL-----------------------------------   278
tr|A0A814QMX3|A0A814QMX3_9BILA      ------------------------------------------------------------   208
tr|K2MZ59|K2MZ59_TRYCR              WLNPSQRCDFFLKRSLTVGGYMGEDAVAEAHSVPDGFYIMDYDMWCRFRFATKKLHEVAM   304
tr|A0A0G4IUR7|A0A0G4IUR7_PLABS      ------------------------------------------------------------   225
tr|A0A0N0DV74|A0A0N0DV74_LEPPY      ------------------------------------------------------------   300
tr|A4HEJ1|A4HEJ1_LEIBR              ------------------------------------------------------------   331
tr|Q4Q9R0|Q4Q9R0_LEIMA              ------------------------------------------------------------   300
tr|A4I1M7|A4I1M7_LEIIN              ------------------------------------------------------------   300
tr|S9UNH8|S9UNH8_9TRYP              ------------------------------------------------------------   326
tr|S9UZF8|S9UZF8_9TRYP              ------------------------------------------------------------   314
tr|Q57ZC4|Q57ZC4_TRYB2              ------------------------------------------------------------   289
tr|A0A3R7LGJ5|A0A3R7LGJ5_9TRYP      ------------------------------------------------------------   290
tr|Q4CUZ5|Q4CUZ5_TRYCC              ------------------------------------------------------------   331
tr|Q4D7T1|Q4D7T1_TRYCC              ------------------------------------------------------------   295
tr|A0A3R7KMG1|A0A3R7KMG1_TRYRA      ------------------------------------------------------------   327
tr|Q4CYQ4|Q4CYQ4_TRYCC              ------------------------------------------------------------   287
tr|Q4CUZ4|Q4CUZ4_TRYCC              ------------------------------------------------------------   294
tr|Q4CUZ2|Q4CUZ2_TRYCC              ------------------------------------------------------------   194
tr|Q4CYQ2|Q4CYQ2_TRYCC              ------------------------------------------------------------   164
tr|Q4D7T2|Q4D7T2_TRYCC              ------------------------------------------------------------   294
tr|Q4D7T0|Q4D7T0_TRYCC              ------------------------------------------------------------   164
tr|A0A1X0NY88|A0A1X0NY88_9TRYP      ------------------------------------------------------------   280
tr|A0A1X0NYK9|A0A1X0NYK9_9TRYP      ------------------------------------------------------------   294
tr|A0A1X0NRB4|A0A1X0NRB4_9TRYP      ------------------------------------------------------------   340
tr|Q4D408|Q4D408_TRYCC              ------------------------------------------------------------   330
tr|A0A422N172|A0A422N172_9TRYP      ------------------------------------------------------------   322
tr|A0A422NKH5|A0A422NKH5_TRYRA      ------------------------------------------------------------   329
tr|A0A0S4ITA7|A0A0S4ITA7_BODSA      ------------------------------------------------------------   309
tr|A0A0N0DYP4|A0A0N0DYP4_LEPPY      ------------------------------------------------------------   363
tr|A4HLZ6|A4HLZ6_LEIBR              ------------------------------------------------------------   382
tr|Q4Q3Q0|Q4Q3Q0_LEIMA              ------------------------------------------------------------   421
tr|A4I9C6|A4I9C6_LEIIN              ------------------------------------------------------------   404
tr|A0A0N0DYQ7|A0A0N0DYQ7_LEPPY      ------------------------------------------------------------   407
tr|A4HLZ4|A4HLZ4_LEIBR              ------------------------------------------------------------   334
tr|Q4Q3Q2|Q4Q3Q2_LEIMA              ------------------------------------------------------------   232
tr|A4I9C4|A4I9C4_LEIIN              ------------------------------------------------------------   369
tr|Q384E7|Q384E7_TRYB2              ------------------------------------------------------------   274
tr|Q586Y2|Q586Y2_TRYB2              ------------------------------------------------------------   273
tr|Q586Y1|Q586Y1_TRYB2              ------------------------------------------------------------   280
tr|F9W970|F9W970_TRYCI              ------------------------------------------------------------   274
tr|F9W971|F9W971_TRYCI              ------------------------------------------------------------   275
tr|S9UGT3|S9UGT3_9TRYP              ------------------------------------------------------------   265
tr|A0A1X0NR96|A0A1X0NR96_9TRYP      ------------------------------------------------------------   286
tr|A0A1X0NR98|A0A1X0NR98_9TRYP      ------------------------------------------------------------   294
tr|Q4D409|Q4D409_TRYCC              ------------------------------------------------------------   280
tr|Q4D410|Q4D410_TRYCC              ------------------------------------------------------------   195
tr|Q4CKB1|Q4CKB1_TRYCC              ------------------------------------------------------------   243
tr|Q4DEC6|Q4DEC6_TRYCC              ------------------------------------------------------------   310
tr|A0A3R7N0H0|A0A3R7N0H0_9TRYP      ------------------------------------------------------------   311
tr|A0A3R7N2W0|A0A3R7N2W0_9TRYP      ------------------------------------------------------------   283
tr|A0A3S5IRC8|A0A3S5IRC8_TRYRA      ------------------------------------------------------------   282


tr|A0A815F576|A0A815F576_9BILA      ------------------------------------------------------------   278
tr|A0A814QMX3|A0A814QMX3_9BILA      ------------------------------------------------------------   208
tr|K2MZ59|K2MZ59_TRYCR              ETTRLILNSRDSEALLVQYFGLPTELHLQLRRSFETIPSMCGRFDFGYDGKEIMMLAYKC   364
tr|A0A0G4IUR7|A0A0G4IUR7_PLABS      ------------------------------------------------------------   225
tr|A0A0N0DV74|A0A0N0DV74_LEPPY      ------------------------------------------------------------   300
tr|A4HEJ1|A4HEJ1_LEIBR              ------------------------------------------------------------   331
tr|Q4Q9R0|Q4Q9R0_LEIMA              ------------------------------------------------------------   300
tr|A4I1M7|A4I1M7_LEIIN              ------------------------------------------------------------   300
tr|S9UNH8|S9UNH8_9TRYP              ------------------------------------------------------------   326
tr|S9UZF8|S9UZF8_9TRYP              ------------------------------------------------------------   314
tr|Q57ZC4|Q57ZC4_TRYB2              ------------------------------------------------------------   289
tr|A0A3R7LGJ5|A0A3R7LGJ5_9TRYP      ------------------------------------------------------------   290
tr|Q4CUZ5|Q4CUZ5_TRYCC              ------------------------------------------------------------   331
tr|Q4D7T1|Q4D7T1_TRYCC              ------------------------------------------------------------   295
tr|A0A3R7KMG1|A0A3R7KMG1_TRYRA      ------------------------------------------------------------   327
tr|Q4CYQ4|Q4CYQ4_TRYCC              ------------------------------------------------------------   287
tr|Q4CUZ4|Q4CUZ4_TRYCC              ------------------------------------------------------------   294
tr|Q4CUZ2|Q4CUZ2_TRYCC              ------------------------------------------------------------   194
tr|Q4CYQ2|Q4CYQ2_TRYCC              ------------------------------------------------------------   164
```

```
tr|Q4D7T2|Q4D7T2_TRYCC                 ------------------------------------------------------------   294
tr|Q4D7T0|Q4D7T0_TRYCC                 ------------------------------------------------------------   164
tr|A0A1X0NY88|A0A1X0NY88_9TRYP         ------------------------------------------------------------   280
tr|A0A1X0NYK9|A0A1X0NYK9_9TRYP         ------------------------------------------------------------   294
tr|A0A1X0NRB4|A0A1X0NRB4_9TRYP         ------------------------------------------------------------   340
tr|Q4D408|Q4D408_TRYCC                 ------------------------------------------------------------   330
tr|A0A422N172|A0A422N172_9TRYP         ------------------------------------------------------------   322
tr|A0A422NKH5|A0A422NKH5_TRYRA         ------------------------------------------------------------   329
tr|A0A0S4ITA7|A0A0S4ITA7_BODSA         ------------------------------------------------------------   309
tr|A0A0N0DYP4|A0A0N0DYP4_LEPPY         ------------------------------------------------------------   363
tr|A4HLZ6|A4HLZ6_LEIBR                 ------------------------------------------------------------   382
tr|Q4Q3Q0|Q4Q3Q0_LEIMA                 ------------------------------------------------------------   421
tr|A4I9C6|A4I9C6_LEIIN                 ------------------------------------------------------------   404
tr|A0A0N0DYQ7|A0A0N0DYQ7_LEPPY         ------------------------------------------------------------   407
tr|A4HLZ4|A4HLZ4_LEIBR                 ------------------------------------------------------------   334
tr|Q4Q3Q2|Q4Q3Q2_LEIMA                 ------------------------------------------------------------   232
tr|A4I9C4|A4I9C4_LEIIN                 ------------------------------------------------------------   369
tr|Q384E7|Q384E7_TRYB2                 ------------------------------------------------------------   274
tr|Q586Y2|Q586Y2_TRYB2                 ------------------------------------------------------------   273
tr|Q586Y1|Q586Y1_TRYB2                 ------------------------------------------------------------   280
tr|F9W970|F9W970_TRYCI                 ------------------------------------------------------------   274
tr|F9W971|F9W971_TRYCI                 ------------------------------------------------------------   275
tr|S9UGT3|S9UGT3_9TRYP                 ------------------------------------------------------------   265
tr|A0A1X0NR96|A0A1X0NR96_9TRYP         ------------------------------------------------------------   286
tr|A0A1X0NR98|A0A1X0NR98_9TRYP         ------------------------------------------------------------   294
tr|Q4D409|Q4D409_TRYCC                 ------------------------------------------------------------   280
tr|Q4D410|Q4D410_TRYCC                 ------------------------------------------------------------   195
tr|Q4CKB1|Q4CKB1_TRYCC                 ------------------------------------------------------------   243
tr|Q4DEC6|Q4DEC6_TRYCC                 ------------------------------------------------------------   310
tr|A0A3R7N0H0|A0A3R7N0H0_9TRYP         ------------------------------------------------------------   311
tr|A0A3R7N2W0|A0A3R7N2W0_9TRYP         ------------------------------------------------------------   283
tr|A0A3S5IRC8|A0A3S5IRC8_TRYRA         ------------------------------------------------------------   282


tr|A0A815F576|A0A815F576_9BILA         ------------------------------------------------------------   278
tr|A0A814QMX3|A0A814QMX3_9BILA         ------------------------------------------------------------   208
tr|K2MZ59|K2MZ59_TRYCR                 DSSAAMLECGDTQEKFTRHYGVKLGNSTGSFLYSRIVNYFSCLMQNECLCPHHKIVHFMI   424
tr|A0A0G4IUR7|A0A0G4IUR7_PLABS         ------------------------------------------------------------   225
tr|A0A0N0DV74|A0A0N0DV74_LEPPY         ------------------------------------------------------------   300
tr|A4HEJ1|A4HEJ1_LEIBR                 ------------------------------------------------------------   331
tr|Q4Q9R0|Q4Q9R0_LEIMA                 ------------------------------------------------------------   300
tr|A4I1M7|A4I1M7_LEIIN                 ------------------------------------------------------------   300
tr|S9UNH8|S9UNH8_9TRYP                 ------------------------------------------------------------   326
tr|S9UZF8|S9UZF8_9TRYP                 ------------------------------------------------------------   314
tr|Q57ZC4|Q57ZC4_TRYB2                 ------------------------------------------------------------   289
tr|A0A3R7LGJ5|A0A3R7LGJ5_9TRYP         ------------------------------------------------------------   290
tr|Q4CUZ5|Q4CUZ5_TRYCC                 ------------------------------------------------------------   331
tr|Q4D7T1|Q4D7T1_TRYCC                 ------------------------------------------------------------   295
tr|A0A3R7KMG1|A0A3R7KMG1_TRYRA         ------------------------------------------------------------   327
tr|Q4CYQ4|Q4CYQ4_TRYCC                 ------------------------------------------------------------   287
tr|Q4CUZ4|Q4CUZ4_TRYCC                 ------------------------------------------------------------   294
tr|Q4CUZ2|Q4CUZ2_TRYCC                 ------------------------------------------------------------   194
tr|Q4CYQ2|Q4CYQ2_TRYCC                 ------------------------------------------------------------   164
tr|Q4D7T2|Q4D7T2_TRYCC                 ------------------------------------------------------------   294
tr|Q4D7T0|Q4D7T0_TRYCC                 ------------------------------------------------------------   164
tr|A0A1X0NY88|A0A1X0NY88_9TRYP         ------------------------------------------------------------   280
tr|A0A1X0NYK9|A0A1X0NYK9_9TRYP         ------------------------------------------------------------   294
tr|A0A1X0NRB4|A0A1X0NRB4_9TRYP         ------------------------------------------------------------   340
tr|Q4D408|Q4D408_TRYCC                 ------------------------------------------------------------   330
tr|A0A422N172|A0A422N172_9TRYP         ------------------------------------------------------------   322
tr|A0A422NKH5|A0A422NKH5_TRYRA         ------------------------------------------------------------   329
tr|A0A0S4ITA7|A0A0S4ITA7_BODSA         ------------------------------------------------------------   309
tr|A0A0N0DYP4|A0A0N0DYP4_LEPPY         ------------------------------------------------------------   363
tr|A4HLZ6|A4HLZ6_LEIBR                 ------------------------------------------------------------   382
tr|Q4Q3Q0|Q4Q3Q0_LEIMA                 ------------------------------------------------------------   421
tr|A4I9C6|A4I9C6_LEIIN                 ------------------------------------------------------------   404
tr|A0A0N0DYQ7|A0A0N0DYQ7_LEPPY         ------------------------------------------------------------   407
tr|A4HLZ4|A4HLZ4_LEIBR                 ------------------------------------------------------------   334
tr|Q4Q3Q2|Q4Q3Q2_LEIMA                 ------------------------------------------------------------   232
tr|A4I9C4|A4I9C4_LEIIN                 ------------------------------------------------------------   369
tr|Q384E7|Q384E7_TRYB2                 ------------------------------------------------------------   274
tr|Q586Y2|Q586Y2_TRYB2                 ------------------------------------------------------------   273
tr|Q586Y1|Q586Y1_TRYB2                 ------------------------------------------------------------   280
tr|F9W970|F9W970_TRYCI                 ------------------------------------------------------------   274
tr|F9W971|F9W971_TRYCI                 ------------------------------------------------------------   275
tr|S9UGT3|S9UGT3_9TRYP                 ------------------------------------------------------------   265
tr|A0A1X0NR96|A0A1X0NR96_9TRYP         ------------------------------------------------------------   286
tr|A0A1X0NR98|A0A1X0NR98_9TRYP         ------------------------------------------------------------   294
tr|Q4D409|Q4D409_TRYCC                 ------------------------------------------------------------   280
tr|Q4D410|Q4D410_TRYCC                 ------------------------------------------------------------   195
tr|Q4CKB1|Q4CKB1_TRYCC                 ------------------------------------------------------------   243
tr|Q4DEC6|Q4DEC6_TRYCC                 ------------------------------------------------------------   310
tr|A0A3R7N0H0|A0A3R7N0H0_9TRYP         ------------------------------------------------------------   311
tr|A0A3R7N2W0|A0A3R7N2W0_9TRYP         ------------------------------------------------------------   283
tr|A0A3S5IRC8|A0A3S5IRC8_TRYRA         ------------------------------------------------------------   282


tr|A0A815F576|A0A815F576_9BILA         ------------------------------------------------------------   278
```

```
tr|A0A814QMX3|A0A814QMX3_9BILA      ------------------------------------------------------------    208
tr|K2MZ59|K2MZ59_TRYCR              DDNDEERYTALYVMNAAESVGFRTKLCVKLSDFRFGSSTGEVDLDASPTERRSVVDLENE    484
tr|A0A0G4IUR7|A0A0G4IUR7_PLABS      ------------------------------------------------------------    225
tr|A0A0N0DV74|A0A0N0DV74_LEPPY      ------------------------------------------------------------    300
tr|A4HEJ1|A4HEJ1_LEIBR              ------------------------------------------------------------    331
tr|Q4Q9R0|Q4Q9R0_LEIMA              ------------------------------------------------------------    300
tr|A4I1M7|A4I1M7_LEIIN              ------------------------------------------------------------    300
tr|S9UNH8|S9UNH8_9TRYP              ------------------------------------------------------------    326
tr|S9UZF8|S9UZF8_9TRYP              ------------------------------------------------------------    314
tr|Q57ZC4|Q57ZC4_TRYB2              ------------------------------------------------------------    289
tr|A0A3R7LGJ5|A0A3R7LGJ5_9TRYP      ------------------------------------------------------------    290
tr|Q4CUZ5|Q4CUZ5_TRYCC              ------------------------------------------------------------    331
tr|Q4D7T1|Q4D7T1_TRYCC              ------------------------------------------------------------    295
tr|A0A3R7KMG1|A0A3R7KMG1_TRYRA      ------------------------------------------------------------    327
tr|Q4CYQ4|Q4CYQ4_TRYCC              ------------------------------------------------------------    287
tr|Q4CUZ4|Q4CUZ4_TRYCC              ------------------------------------------------------------    294
tr|Q4CUZ2|Q4CUZ2_TRYCC              ------------------------------------------------------------    194
tr|Q4CYQ2|Q4CYQ2_TRYCC              ------------------------------------------------------------    164
tr|Q4D7T2|Q4D7T2_TRYCC              ------------------------------------------------------------    294
tr|Q4D7T0|Q4D7T0_TRYCC              ------------------------------------------------------------    164
tr|A0A1X0NY88|A0A1X0NY88_9TRYP      ------------------------------------------------------------    280
tr|A0A1X0NYK9|A0A1X0NYK9_9TRYP      ------------------------------------------------------------    294
tr|A0A1X0NRB4|A0A1X0NRB4_9TRYP      ------------------------------------------------------------    340
tr|Q4D408|Q4D408_TRYCC              ------------------------------------------------------------    330
tr|A0A422N172|A0A422N172_9TRYP      ------------------------------------------------------------    322
tr|A0A422NKH5|A0A422NKH5_TRYRA      ------------------------------------------------------------    329
tr|A0A0S4ITA7|A0A0S4ITA7_BODSA      ------------------------------------------------------------    309
tr|A0A0N0DYP4|A0A0N0DYP4_LEPPY      ------------------------------------------------------------    363
tr|A4HLZ6|A4HLZ6_LEIBR              ------------------------------------------------------------    382
tr|Q4Q3Q0|Q4Q3Q0_LEIMA              ------------------------------------------------------------    421
tr|A4I9C6|A4I9C6_LEIIN              ------------------------------------------------------------    404
tr|A0A0N0DYQ7|A0A0N0DYQ7_LEPPY      ------------------------------------------------------------    407
tr|A4HLZ4|A4HLZ4_LEIBR              ------------------------------------------------------------    334
tr|Q4Q3Q2|Q4Q3Q2_LEIMA              ------------------------------------------------------------    232
tr|A4I9C4|A4I9C4_LEIIN              ------------------------------------------------------------    369
tr|Q384E7|Q384E7_TRYB2              ------------------------------------------------------------    274
tr|Q586Y2|Q586Y2_TRYB2              ------------------------------------------------------------    273
tr|Q586Y1|Q586Y1_TRYB2              ------------------------------------------------------------    280
tr|F9W970|F9W970_TRYCI              ------------------------------------------------------------    274
tr|F9W971|F9W971_TRYCI              ------------------------------------------------------------    275
tr|S9UGT3|S9UGT3_9TRYP              ------------------------------------------------------------    265
tr|A0A1X0NR96|A0A1X0NR96_9TRYP      ------------------------------------------------------------    286
tr|A0A1X0NR98|A0A1X0NR98_9TRYP      ------------------------------------------------------------    294
tr|Q4D409|Q4D409_TRYCC              ------------------------------------------------------------    280
tr|Q4D410|Q4D410_TRYCC              ------------------------------------------------------------    195
tr|Q4CKB1|Q4CKB1_TRYCC              ------------------------------------------------------------    243
tr|Q4DEC6|Q4DEC6_TRYCC              ------------------------------------------------------------    310
tr|A0A3R7N0H0|A0A3R7N0H0_9TRYP      ------------------------------------------------------------    311
tr|A0A3R7N2W0|A0A3R7N2W0_9TRYP      ------------------------------------------------------------    283
tr|A0A3S5IRC8|A0A3S5IRC8_TRYRA      ------------------------------------------------------------    282

tr|A0A815F576|A0A815F576_9BILA      ------------------------------------------------------------    278
tr|A0A814QMX3|A0A814QMX3_9BILA      ------------------------------------------------------------    208
tr|K2MZ59|K2MZ59_TRYCR              KVLLVWKTWAWDTVVQQYMEQRTQEGHGVSTKPTLSDILLNEHIRVLEPLWKAVTGSKAI    544
tr|A0A0G4IUR7|A0A0G4IUR7_PLABS      ------------------------------------------------------------    225
tr|A0A0N0DV74|A0A0N0DV74_LEPPY      ------------------------------------------------------------    300
tr|A4HEJ1|A4HEJ1_LEIBR              ------------------------------------------------------------    331
tr|Q4Q9R0|Q4Q9R0_LEIMA              ------------------------------------------------------------    300
tr|A4I1M7|A4I1M7_LEIIN              ------------------------------------------------------------    300
tr|S9UNH8|S9UNH8_9TRYP              ------------------------------------------------------------    326
tr|S9UZF8|S9UZF8_9TRYP              ------------------------------------------------------------    314
tr|Q57ZC4|Q57ZC4_TRYB2              ------------------------------------------------------------    289
tr|A0A3R7LGJ5|A0A3R7LGJ5_9TRYP      ------------------------------------------------------------    290
tr|Q4CUZ5|Q4CUZ5_TRYCC              ------------------------------------------------------------    331
tr|Q4D7T1|Q4D7T1_TRYCC              ------------------------------------------------------------    295
tr|A0A3R7KMG1|A0A3R7KMG1_TRYRA      ------------------------------------------------------------    327
tr|Q4CYQ4|Q4CYQ4_TRYCC              ------------------------------------------------------------    287
tr|Q4CUZ4|Q4CUZ4_TRYCC              ------------------------------------------------------------    294
tr|Q4CUZ2|Q4CUZ2_TRYCC              ------------------------------------------------------------    194
tr|Q4CYQ2|Q4CYQ2_TRYCC              ------------------------------------------------------------    164
tr|Q4D7T2|Q4D7T2_TRYCC              ------------------------------------------------------------    294
tr|Q4D7T0|Q4D7T0_TRYCC              ------------------------------------------------------------    164
tr|A0A1X0NY88|A0A1X0NY88_9TRYP      ------------------------------------------------------------    280
tr|A0A1X0NYK9|A0A1X0NYK9_9TRYP      ------------------------------------------------------------    294
tr|A0A1X0NRB4|A0A1X0NRB4_9TRYP      ------------------------------------------------------------    340
tr|Q4D408|Q4D408_TRYCC              ------------------------------------------------------------    330
tr|A0A422N172|A0A422N172_9TRYP      ------------------------------------------------------------    322
tr|A0A422NKH5|A0A422NKH5_TRYRA      ------------------------------------------------------------    329
tr|A0A0S4ITA7|A0A0S4ITA7_BODSA      ------------------------------------------------------------    309
tr|A0A0N0DYP4|A0A0N0DYP4_LEPPY      ------------------------------------------------------------    363
tr|A4HLZ6|A4HLZ6_LEIBR              ------------------------------------------------------------    382
tr|Q4Q3Q0|Q4Q3Q0_LEIMA              ------------------------------------------------------------    421
tr|A4I9C6|A4I9C6_LEIIN              ------------------------------------------------------------    404
tr|A0A0N0DYQ7|A0A0N0DYQ7_LEPPY      ------------------------------------------------------------    407
tr|A4HLZ4|A4HLZ4_LEIBR              ------------------------------------------------------------    334
tr|Q4Q3Q2|Q4Q3Q2_LEIMA              ------------------------------------------------------------    232
tr|A4I9C4|A4I9C4_LEIIN              ------------------------------------------------------------    369
```

```
tr|Q384E7|Q384E7_TRYB2                  ------------------------------------------------------------   274
tr|Q586Y2|Q586Y2_TRYB2                  ------------------------------------------------------------   273
tr|Q586Y1|Q586Y1_TRYB2                  ------------------------------------------------------------   280
tr|F9W970|F9W970_TRYCI                  ------------------------------------------------------------   274
tr|F9W971|F9W971_TRYCI                  ------------------------------------------------------------   275
tr|S9UGT3|S9UGT3_9TRYP                  ------------------------------------------------------------   265
tr|A0A1X0NR96|A0A1X0NR96_9TRYP          ------------------------------------------------------------   286
tr|A0A1X0NR98|A0A1X0NR98_9TRYP          ------------------------------------------------------------   294
tr|Q4D409|Q4D409_TRYCC                  ------------------------------------------------------------   280
tr|Q4D410|Q4D410_TRYCC                  ------------------------------------------------------------   195
tr|Q4CKB1|Q4CKB1_TRYCC                  ------------------------------------------------------------   243
tr|Q4DEC6|Q4DEC6_TRYCC                  ------------------------------------------------------------   310
tr|A0A3R7N0H0|A0A3R7N0H0_9TRYP          ------------------------------------------------------------   311
tr|A0A3R7N2W0|A0A3R7N2W0_9TRYP          ------------------------------------------------------------   283
tr|A0A3S5IRC8|A0A3S5IRC8_TRYRA          ------------------------------------------------------------   282


tr|A0A815F576|A0A815F576_9BILA          ------------------------------------------------------------   278
tr|A0A814QMX3|A0A814QMX3_9BILA          ------------------------------------------------------------   208
tr|K2MZ59|K2MZ59_TRYCR                  LPFMYAVAPHHPNMVPASFYRTKEIISSPYLTKPVNGRVGKNMTIYDPEENPEAIAAAPY   604
tr|A0A0G4IUR7|A0A0G4IUR7_PLABS          ------------------------------------------------------------   225
tr|A0A0N0DV74|A0A0N0DV74_LEPPY          ------------------------------------------------------------   300
tr|A4HEJ1|A4HEJ1_LEIBR                  ------------------------------------------------------------   331
tr|Q4Q9R0|Q4Q9R0_LEIMA                  ------------------------------------------------------------   300
tr|A4I1M7|A4I1M7_LEIIN                  ------------------------------------------------------------   300
tr|S9UNH8|S9UNH8_9TRYP                  ------------------------------------------------------------   326
tr|S9UZF8|S9UZF8_9TRYP                  ------------------------------------------------------------   314
tr|Q57ZC4|Q57ZC4_TRYB2                  ------------------------------------------------------------   289
tr|A0A3R7LGJ5|A0A3R7LGJ5_9TRYP          ------------------------------------------------------------   290
tr|Q4CUZ5|Q4CUZ5_TRYCC                  ------------------------------------------------------------   331
tr|Q4D7T1|Q4D7T1_TRYCC                  ------------------------------------------------------------   295
tr|A0A3R7KMG1|A0A3R7KMG1_TRYRA          ------------------------------------------------------------   327
tr|Q4CYQ4|Q4CYQ4_TRYCC                  ------------------------------------------------------------   287
tr|Q4CUZ4|Q4CUZ4_TRYCC                  ------------------------------------------------------------   294
tr|Q4CUZ2|Q4CUZ2_TRYCC                  ------------------------------------------------------------   194
tr|Q4CYQ2|Q4CYQ2_TRYCC                  ------------------------------------------------------------   164
tr|Q4D7T2|Q4D7T2_TRYCC                  ------------------------------------------------------------   294
tr|Q4D7T0|Q4D7T0_TRYCC                  ------------------------------------------------------------   164
tr|A0A1X0NY88|A0A1X0NY88_9TRYP          ------------------------------------------------------------   280
tr|A0A1X0NYK9|A0A1X0NYK9_9TRYP          ------------------------------------------------------------   294
tr|A0A1X0NRB4|A0A1X0NRB4_9TRYP          ------------------------------------------------------------   340
tr|Q4D408|Q4D408_TRYCC                  ------------------------------------------------------------   330
tr|A0A422N172|A0A422N172_9TRYP          ------------------------------------------------------------   322
tr|A0A422NKH5|A0A422NKH5_TRYRA          ------------------------------------------------------------   329
tr|A0A0S4ITA7|A0A0S4ITA7_BODSA          ------------------------------------------------------------   309
tr|A0A0N0DYP4|A0A0N0DYP4_LEPPY          ------------------------------------------------------------   363
tr|A4HLZ6|A4HLZ6_LEIBR                  ------------------------------------------------------------   382
tr|Q4Q3Q0|Q4Q3Q0_LEIMA                  ------------------------------------------------------------   421
tr|A4I9C6|A4I9C6_LEIIN                  ------------------------------------------------------------   404
tr|A0A0N0DYQ7|A0A0N0DYQ7_LEPPY          ------------------------------------------------------------   407
tr|A4HLZ4|A4HLZ4_LEIBR                  ------------------------------------------------------------   334
tr|Q4Q3Q2|Q4Q3Q2_LEIMA                  ------------------------------------------------------------   232
tr|A4I9C4|A4I9C4_LEIIN                  ------------------------------------------------------------   369
tr|Q384E7|Q384E7_TRYB2                  ------------------------------------------------------------   274
tr|Q586Y2|Q586Y2_TRYB2                  ------------------------------------------------------------   273
tr|Q586Y1|Q586Y1_TRYB2                  ------------------------------------------------------------   280
tr|F9W970|F9W970_TRYCI                  ------------------------------------------------------------   274
tr|F9W971|F9W971_TRYCI                  ------------------------------------------------------------   275
tr|S9UGT3|S9UGT3_9TRYP                  ------------------------------------------------------------   265
tr|A0A1X0NR96|A0A1X0NR96_9TRYP          ------------------------------------------------------------   286
tr|A0A1X0NR98|A0A1X0NR98_9TRYP          ------------------------------------------------------------   294
tr|Q4D409|Q4D409_TRYCC                  ------------------------------------------------------------   280
tr|Q4D410|Q4D410_TRYCC                  ------------------------------------------------------------   195
tr|Q4CKB1|Q4CKB1_TRYCC                  ------------------------------------------------------------   243
tr|Q4DEC6|Q4DEC6_TRYCC                  ------------------------------------------------------------   310
tr|A0A3R7N0H0|A0A3R7N0H0_9TRYP          ------------------------------------------------------------   311
tr|A0A3R7N2W0|A0A3R7N2W0_9TRYP          ------------------------------------------------------------   283
tr|A0A3S5IRC8|A0A3S5IRC8_TRYRA          ------------------------------------------------------------   282


tr|A0A815F576|A0A815F576_9BILA          ------------------------------------------------------------   278
tr|A0A814QMX3|A0A814QMX3_9BILA          ------------------------------------------------------------   208
tr|K2MZ59|K2MZ59_TRYCR                  TTAAPVLGRSISAVLFDDAPVQPCRMPFDNVHENLTGRLFDSAVVYQSRVFLTRYEKKYS   664
tr|A0A0G4IUR7|A0A0G4IUR7_PLABS          ------------------------------------------------------------   225
tr|A0A0N0DV74|A0A0N0DV74_LEPPY          ------------------------------------------------------------   300
tr|A4HEJ1|A4HEJ1_LEIBR                  ------------------------------------------------------------   331
tr|Q4Q9R0|Q4Q9R0_LEIMA                  ------------------------------------------------------------   300
tr|A4I1M7|A4I1M7_LEIIN                  ------------------------------------------------------------   300
tr|S9UNH8|S9UNH8_9TRYP                  ------------------------------------------------------------   326
tr|S9UZF8|S9UZF8_9TRYP                  ------------------------------------------------------------   314
tr|Q57ZC4|Q57ZC4_TRYB2                  ------------------------------------------------------------   289
tr|A0A3R7LGJ5|A0A3R7LGJ5_9TRYP          ------------------------------------------------------------   290
tr|Q4CUZ5|Q4CUZ5_TRYCC                  ------------------------------------------------------------   331
tr|Q4D7T1|Q4D7T1_TRYCC                  ------------------------------------------------------------   295
tr|A0A3R7KMG1|A0A3R7KMG1_TRYRA          ------------------------------------------------------------   327
tr|Q4CYQ4|Q4CYQ4_TRYCC                  ------------------------------------------------------------   287
tr|Q4CUZ4|Q4CUZ4_TRYCC                  ------------------------------------------------------------   294
tr|Q4CUZ2|Q4CUZ2_TRYCC                  ------------------------------------------------------------   194
```

```
tr|Q4CYQ2|Q4CYQ2_TRYCC                ------------------------------------------------------------    164
tr|Q4D7T2|Q4D7T2_TRYCC                ------------------------------------------------------------    294
tr|Q4D7T0|Q4D7T0_TRYCC                ------------------------------------------------------------    164
tr|A0A1X0NY88|A0A1X0NY88_9TRYP        ------------------------------------------------------------    280
tr|A0A1X0NYK9|A0A1X0NYK9_9TRYP        ------------------------------------------------------------    294
tr|A0A1X0NRB4|A0A1X0NRB4_9TRYP        ------------------------------------------------------------    340
tr|Q4D408|Q4D408_TRYCC                ------------------------------------------------------------    330
tr|A0A422N172|A0A422N172_9TRYP        ------------------------------------------------------------    322
tr|A0A422NKH5|A0A422NKH5_TRYRA        ------------------------------------------------------------    329
tr|A0A0S4ITA7|A0A0S4ITA7_BODSA        ------------------------------------------------------------    309
tr|A0A0N0DYP4|A0A0N0DYP4_LEPPY        ------------------------------------------------------------    363
tr|A4HLZ6|A4HLZ6_LEIBR                ------------------------------------------------------------    382
tr|Q4Q3Q0|Q4Q3Q0_LEIMA                ------------------------------------------------------------    421
tr|A4I9C6|A4I9C6_LEIIN                ------------------------------------------------------------    404
tr|A0A0N0DYQ7|A0A0N0DYQ7_LEPPY        ------------------------------------------------------------    407
tr|A4HLZ4|A4HLZ4_LEIBR                ------------------------------------------------------------    334
tr|Q4Q3Q2|Q4Q3Q2_LEIMA                ------------------------------------------------------------    232
tr|A4I9C4|A4I9C4_LEIIN                ------------------------------------------------------------    369
tr|Q384E7|Q384E7_TRYB2                ------------------------------------------------------------    274
tr|Q586Y2|Q586Y2_TRYB2                ------------------------------------------------------------    273
tr|Q586Y1|Q586Y1_TRYB2                ------------------------------------------------------------    280
tr|F9W970|F9W970_TRYCI                ------------------------------------------------------------    274
tr|F9W971|F9W971_TRYCI                ------------------------------------------------------------    275
tr|S9UGT3|S9UGT3_9TRYP                ------------------------------------------------------------    265
tr|A0A1X0NR96|A0A1X0NR96_9TRYP        ------------------------------------------------------------    286
tr|A0A1X0NR98|A0A1X0NR98_9TRYP        ------------------------------------------------------------    294
tr|Q4D409|Q4D409_TRYCC                ------------------------------------------------------------    280
tr|Q4D410|Q4D410_TRYCC                ------------------------------------------------------------    195
tr|Q4CKB1|Q4CKB1_TRYCC                ------------------------------------------------------------    243
tr|Q4DEC6|Q4DEC6_TRYCC                ------------------------------------------------------------    310
tr|A0A3R7N0H0|A0A3R7N0H0_9TRYP        ------------------------------------------------------------    311
tr|A0A3R7N2W0|A0A3R7N2W0_9TRYP        ------------------------------------------------------------    283
tr|A0A3S5IRC8|A0A3S5IRC8_TRYRA        ------------------------------------------------------------    282

tr|A0A815F576|A0A815F576_9BILA        -------------------------------------------------------    278
tr|A0A814QMX3|A0A814QMX3_9BILA        -------------------------------------------------------    208
tr|K2MZ59|K2MZ59_TRYCR                PIFCGWNVGGEFGGVIVREETLNTTGLSSLVIPSRVVRQHIPLHAVQESTTFGEV    719
tr|A0A0G4IUR7|A0A0G4IUR7_PLABS        -------------------------------------------------------    225
tr|A0A0N0DV74|A0A0N0DV74_LEPPY        -------------------------------------------------------    300
tr|A4HEJ1|A4HEJ1_LEIBR                -------------------------------------------------------    331
tr|Q4Q9R0|Q4Q9R0_LEIMA                -------------------------------------------------------    300
tr|A4I1M7|A4I1M7_LEIIN                -------------------------------------------------------    300
tr|S9UNH8|S9UNH8_9TRYP                -------------------------------------------------------    326
tr|S9UZF8|S9UZF8_9TRYP                -------------------------------------------------------    314
tr|Q57ZC4|Q57ZC4_TRYB2                -------------------------------------------------------    289
tr|A0A3R7LGJ5|A0A3R7LGJ5_9TRYP        -------------------------------------------------------    290
tr|Q4CUZ5|Q4CUZ5_TRYCC                -------------------------------------------------------    331
tr|Q4D7T1|Q4D7T1_TRYCC                -------------------------------------------------------    295
tr|A0A3R7KMG1|A0A3R7KMG1_TRYRA        -------------------------------------------------------    327
tr|Q4CYQ4|Q4CYQ4_TRYCC                -------------------------------------------------------    287
tr|Q4CUZ4|Q4CUZ4_TRYCC                -------------------------------------------------------    294
tr|Q4CUZ2|Q4CUZ2_TRYCC                -------------------------------------------------------    194
tr|Q4CYQ2|Q4CYQ2_TRYCC                -------------------------------------------------------    164
tr|Q4D7T2|Q4D7T2_TRYCC                -------------------------------------------------------    294
tr|Q4D7T0|Q4D7T0_TRYCC                -------------------------------------------------------    164
tr|A0A1X0NY88|A0A1X0NY88_9TRYP        -------------------------------------------------------    280
tr|A0A1X0NYK9|A0A1X0NYK9_9TRYP        -------------------------------------------------------    294
tr|A0A1X0NRB4|A0A1X0NRB4_9TRYP        -------------------------------------------------------    340
tr|Q4D408|Q4D408_TRYCC                -------------------------------------------------------    330
tr|A0A422N172|A0A422N172_9TRYP        -------------------------------------------------------    322
tr|A0A422NKH5|A0A422NKH5_TRYRA        -------------------------------------------------------    329
tr|A0A0S4ITA7|A0A0S4ITA7_BODSA        -------------------------------------------------------    309
tr|A0A0N0DYP4|A0A0N0DYP4_LEPPY        -------------------------------------------------------    363
tr|A4HLZ6|A4HLZ6_LEIBR                -------------------------------------------------------    382
tr|Q4Q3Q0|Q4Q3Q0_LEIMA                -------------------------------------------------------    421
tr|A4I9C6|A4I9C6_LEIIN                -------------------------------------------------------    404
tr|A0A0N0DYQ7|A0A0N0DYQ7_LEPPY        -------------------------------------------------------    407
tr|A4HLZ4|A4HLZ4_LEIBR                -------------------------------------------------------    334
tr|Q4Q3Q2|Q4Q3Q2_LEIMA                -------------------------------------------------------    232
tr|A4I9C4|A4I9C4_LEIIN                -------------------------------------------------------    369
tr|Q384E7|Q384E7_TRYB2                -------------------------------------------------------    274
tr|Q586Y2|Q586Y2_TRYB2                -------------------------------------------------------    273
tr|Q586Y1|Q586Y1_TRYB2                -------------------------------------------------------    280
tr|F9W970|F9W970_TRYCI                -------------------------------------------------------    274
tr|F9W971|F9W971_TRYCI                -------------------------------------------------------    275
tr|S9UGT3|S9UGT3_9TRYP                -------------------------------------------------------    265
tr|A0A1X0NR96|A0A1X0NR96_9TRYP        -------------------------------------------------------    286
tr|A0A1X0NR98|A0A1X0NR98_9TRYP        -------------------------------------------------------    294
tr|Q4D409|Q4D409_TRYCC                -------------------------------------------------------    280
tr|Q4D410|Q4D410_TRYCC                -------------------------------------------------------    195
tr|Q4CKB1|Q4CKB1_TRYCC                -------------------------------------------------------    243
tr|Q4DEC6|Q4DEC6_TRYCC                -------------------------------------------------------    310
tr|A0A3R7N0H0|A0A3R7N0H0_9TRYP        -------------------------------------------------------    311
tr|A0A3R7N2W0|A0A3R7N2W0_9TRYP        -------------------------------------------------------    283
tr|A0A3S5IRC8|A0A3S5IRC8_TRYRA        -------------------------------------------------------    282
```

### 06. Leishmaniid small secreted / ER proteins (LESSER)

This is a very unique family identified only among kinetoplastids. These proteins are very small, and consist only of a single domain. Even though kinetoplastid organisms typically only have one ortholog, there is a significant variability: it is likely secreted in most groups (without KDEL), apparently retained in the ER in leishmaniids (Leishmania + Leptomonas group) and very rarely can even be anchored to the membrane in others. The biological function of these proteins is utterly unclear.

```
tr|A0A0S4JVT8|A0A0S4JVT8_BODSA      ------------------------------------------------------------    0
tr|A0A0M9G0R4|A0A0M9G0R4_LEPPY      ------------------------------------------------------------    0
tr|A4HNW9|A4HNW9_LEIBR              ------------------------------------------------------------    0
tr|Q4Q1W7|Q4Q1W7_LEIMA              ------------------------------------------------------------    0
tr|A4ICM5|A4ICM5_LEIIN              ------------------------------------------------------------    0
tr|A0A1X0P821|A0A1X0P821_9TRYP      ------------------------------------------------------------    0
tr|S9TGZ6|S9TGZ6_9TRYP              ------------------------------------------------------------    0
tr|A0A7G2C2Y3|A0A7G2C2Y3_9TRYP      --------------------------MFIRCRFAPFELKRISFF----------------    18
tr|A0A1X0P888|A0A1X0P888_9TRYP      ------------------------------------------------------------    0
tr|A0A3R7KQP7|A0A3R7KQP7_9TRYP      ------------------------------------------------------------    0
tr|A0A3R7NU99|A0A3R7NU99_TRYRA      ------------------------------------------------------------    0
tr|Q4CWZ8|Q4CWZ8_TRYCC              MDGKYIYIRFCPMHGPFFFFFLVNRMIFACPPFAPMEFSLYLFLFFYSSPITHPGQNNNK    60
tr|Q4CZ14|Q4CZ14_TRYCC              -----------------------------------MEFPLYLFLFFYSSPITHPGQNNK-    24

tr|A0A0S4JVT8|A0A0S4JVT8_BODSA      -------------------MLSLKFVL-SVVVVAFLVAAVTGEDGAPAVSVKRIG--TNS    38
tr|A0A0M9G0R4|A0A0M9G0R4_LEPPY      ----------------MRPFLR-AVSMVSLCFAYFLLVGVSAQA--P-----------PR    30
tr|A4HNW9|A4HNW9_LEIBR              -----------------MPFLR-TFTVFCI-IAGVLLQSCAALT--P-----------SR    28
tr|Q4Q1W7|Q4Q1W7_LEIMA              ----------------MVSFLR-TFAVLCI-IAGVLLQGGTARA--P-----------PR    29
tr|A4ICM5|A4ICM5_LEIIN              ----------------MASFLR-TFAILCI-IAGVLLQGSTARA--P-----------PR    29
tr|A0A1X0P821|A0A1X0P821_9TRYP      ----------------------------MIIETA------------A-----------AA    9
tr|S9TGZ6|S9TGZ6_9TRYP              --------------MTRRCFLVLFLAAVCLCFSA------------D-----------AA    23
tr|A0A7G2C2Y3|A0A7G2C2Y3_9TRYP      -----------------------TILITCF-IPTLLLLL--PPITAVYANPDRYA--TVS    50
tr|A0A1X0P888|A0A1X0P888_9TRYP      ----------------MSQ----NVLQTLA-LVLLSLLLVSAEHTLMQTRTNRIVSRG--    37
tr|A0A3R7KQP7|A0A3R7KQP7_9TRYP      ----------------MKG-VRTVWLVVCV-TLLLQVLSSRAEPSVVPARSQRQSVQHPS    42
tr|A0A3R7NU99|A0A3R7NU99_TRYRA      ----------------MRG-VSIVCFFLCV-TLAFQVLPLHAEPSPIPARAQRSSTQKIP    42
tr|Q4CWZ8|Q4CWZ8_TRYCC              KKKKENNYLYLALMLGMRG-VCTVWIFVCV-VLMFGLSLSHAELSTTEIQFQELSPPQTS    118
tr|Q4CZ14|Q4CZ14_TRYCC              KKKKENNYLYLALMLGMRG-VCTVWIFVCV-VLTFGLSLSHAELSTIETQFQELSPPQTS    82

tr|A0A0S4JVT8|A0A0S4JVT8_BODSA      RVRPI----RTTLQACRADFRKLCLKRGENTGAVAPVSRNSPNECLQNQIENIA------    88
tr|A0A0M9G0R4|A0A0M9G0R4_LEPPY      GWRPYESKFNSLSQACYGVLEDACP-----------PNSADRLYCLMLNQDKND------    73
tr|A4HNW9|A4HNW9_LEIBR              AVRPFETKFSSLSEACRGVLETECP-----------PDRLGRADCLMARTEGNE------    71
tr|Q4Q1W7|Q4Q1W7_LEIMA              AVRAFETKFNSVSQACRGVLETECP-----------SDGLARTDCLMEHIVDNA------    72
tr|A4ICM5|A4ICM5_LEIIN              AVRAFETKFNSVSQACRGVLETECP-----------SDGLARTDCLMEHIVDNA------    72
tr|A0A1X0P821|A0A1X0P821_9TRYP      ATVPLYTAPQTITQACWGELQDECP-----------GP---DVDCLANNIIRIK------    49
tr|S9TGZ6|S9TGZ6_9TRYP              TLGELRAAPRNIAQACRDELTATCD-----------HRYNDFPDCLLTHFKKVK------    66
tr|A0A7G2C2Y3|A0A7G2C2Y3_9TRYP      HAAGLPHDPPSVYHACAGEMALLCP-----------DTPHR-VVCFLSAITREGR-----    93
tr|A0A1X0P888|A0A1X0P888_9TRYP      -TEPSVNAPYFLFSVCEAEIQQFCP-----------SQKEAPLKCLLHHFSAARQNPRGG    85
tr|A0A3R7KQP7|A0A3R7KQP7_9TRYP      SRASPTKFLVFPFAACDADAKRLCP-----------TQKESPLKCLLQHFESQRFS----    87
tr|A0A3R7NU99|A0A3R7NU99_TRYRA      FRASPTDLLVFPFSACDADVKQLCP-----------TQKESPLKCLLQHFESQRSS----    87
tr|Q4CWZ8|Q4CWZ8_TRYCC              SEVSETNSLTFPFFACEAELRQLCP-----------KQMEAPLKCLLQHFESQRNT----    163
tr|Q4CZ14|Q4CZ14_TRYCC              SEVSETNSLTFPFFACEAEVRQLCL-----------KQMEAPLKCLLQHFESQRNT----    127
                                                      .*       *                 *:

tr|A0A0S4JVT8|A0A0S4JVT8_BODSA      ------------------------------------------SEVCKTWVSAEQSCTSAL    106
tr|A0A0M9G0R4|A0A0M9G0R4_LEPPY      ------------------------------------------NHECNMWLGWRTVCLSFV    91
tr|A4HNW9|A4HNW9_LEIBR              ------------------------------------------NHECKLWLSWRATCFAYA    89
tr|Q4Q1W7|Q4Q1W7_LEIMA              ------------------------------------------NHECKMWLSWRASCFAYA    90
tr|A4ICM5|A4ICM5_LEIIN              ------------------------------------------NHECKMWLSWRANCFAYA    90
tr|A0A1X0P821|A0A1X0P821_9TRYP      ------------------------------------------SDICRSWITAREVCITHV    67
tr|S9TGZ6|S9TGZ6_9TRYP              ------------------------------------------SPTCKTWLLARSICNSFV    84
tr|A0A7G2C2Y3|A0A7G2C2Y3_9TRYP      ------------------------------------TPDHTFSPTCQSYLSGRESCVRAV    117
tr|A0A1X0P888|A0A1X0P888_9TRYP      GAAQGRPHLLHAYTQQPQQTLQPTQRPQEEPAETQPRRPIHFGKECSSWLRAREVCVTYV    145
tr|A0A3R7KQP7|A0A3R7KQP7_9TRYP      --------------------------------EAPWRRRPAFSEACRRWLWARQECVSYV    115
tr|A0A3R7NU99|A0A3R7NU99_TRYRA      --------------------------------EYPWRRRPAFSKACSRWLWARQECVSYV    115
tr|Q4CWZ8|Q4CWZ8_TRYCC              --------------------------------KLPWRRRITFSGACSRWLWARQECVSYV    191
tr|Q4CZ14|Q4CZ14_TRYCC              --------------------------------KLLWRRRITFSSACSRWLWARQECVSFV    155
                                                                                 .  *  ::  .  *

tr|A0A0S4JVT8|A0A0S4JVT8_BODSA      K------SNTKCPKNS------DIRRCFGKVSATELPDTCTSTEFYKSFAAVRIRGGL--    152
tr|A0A0M9G0R4|A0A0M9G0R4_LEPPY      EVK--LVPNGFCGFTKEESTPELVRQCLGIVDKKELPELCHKSPYYKSLKLHSPRLMD--    147
tr|A4HNW9|A4HNW9_LEIBR              MVK--LIPQGKCNFTSDQMTSELVLQCLRVTDAKELPTACSASPYFKSIMLHAPRRMD--    145
tr|Q4Q1W7|Q4Q1W7_LEIMA              TVF--LIPQGKCNFKPDQATSQLIRNCLRNTDKSELPAVCSGSPYYKSLTLRSPRRID--    146
tr|A4ICM5|A4ICM5_LEIIN              TVY--LIPQGKCNFKPHQATSQLIRNCLRNTEKSELPAVCSGSPYYKSLTLRVPRRMD--    146
tr|A0A1X0P821|A0A1X0P821_9TRYP      K------EVMKCNDD-------KYRQCIREAHRDELPFLCVNSPYYKSLYRNLPRQFL--    112
tr|S9TGZ6|S9TGZ6_9TRYP              K------NNNKCGYDQ---TIFNYRECIRNVPLDELPVQCRESDYYKSLKRFRFE-----    130
tr|A0A7G2C2Y3|A0A7G2C2Y3_9TRYP      RLGQNKFVSKLCGDTI------NVRECLRRVPSHSLPSKCARSLYYRSVLLAPMLSHGQN    171
tr|A0A1X0P888|A0A1X0P888_9TRYP      RRS------GRCRASE------TARECLRRVPASLLPPGCRNTDYHRSVMLYGEMKRQKR    193
tr|A0A3R7KQP7|A0A3R7KQP7_9TRYP      RLP------GRCRASE------SARDCLRRVPPRDLPPGCRDTEYYRSVLLYGEMKSKQG    163
```

```
tr|A0A3R7NU99|A0A3R7NU99_TRYRA      RLA------GKCRVSE------SARDCLRRVPPRDLPPGCRDTEYYRSVLLYGEMKRTQR    163
tr|Q4CWZ8|Q4CWZ8_TRYCC              RQT------GICRASE------SARNCLRRIPPRGIPSGCRETEYYRSVLLYGEMKRKQG    239
tr|Q4CZ14|Q4CZ14_TRYCC              RQT------GICRASE------SARNCLRRIPPRGIPSGCRDTEYYRSVLLYGEMKRKQG    203
                                             *                *:       :*   *  : :.:*.

tr|A0A0S4JVT8|A0A0S4JVT8_BODSA      --------RNRTKTE--    159
tr|A0A0M9G0R4|A0A0M9G0R4_LEPPY      --------DEGDL----    152
tr|A4HNW9|A4HNW9_LEIBR              --------SDGTADL--    152
tr|Q4Q1W7|Q4Q1W7_LEIMA              --------SDAAADL--    153
tr|A4ICM5|A4ICM5_LEIIN              --------SDAAADL--    153
tr|A0A1X0P821|A0A1X0P821_9TRYP      --------R--------    113
tr|S9TGZ6|S9TGZ6_9TRYP              -----------------    130
tr|A0A7G2C2Y3|A0A7G2C2Y3_9TRYP      -----------------    171
tr|A0A1X0P888|A0A1X0P888_9TRYP      T----------------    194
tr|A0A3R7KQP7|A0A3R7KQP7_9TRYP      MRPRNKSNDNETQKLAE    180
tr|A0A3R7NU99|A0A3R7NU99_TRYRA      MRPPEKSKENRTEQHAE    180
tr|Q4CWZ8|Q4CWZ8_TRYCC              LWKPSNTPKKQQKD---    253
tr|Q4CZ14|Q4CZ14_TRYCC              LWKPSNTPKKQQKN---    217
```

### 07. Essential for viability in ER (EVER)

This another very unique protein, consisting of a single domain with a beta-sheet-only architecture (predicted).
While its predicted structure loosely resembles certain plant lectins, the exact biological function is unclear.
There has been an important publication (Plos Pathogens 2022, PMID: 35202447), where the knockout of this gene (LdBPK_211610) was shown to be lethal in Leishmania donovani. The authors assumed a secreted protein (judged by the signal), but now we know that all kinetoplastid members of this family carry a well-discernible KDEL-like signal, hinting at an ER-localized protein. In accordance with its purported essential function, it is found in all better-studied kinetoplastid genomes, except those where it has likely not been mapped yet.
Some organisms have even duplicated this gene.

```
tr|A0A0M9FY26|A0A0M9FY26_LEPPY      ----------------------------------------MGSAQRAFHAVVVATL--LV    18
tr|A4HBZ4|A4HBZ4_LEIBR              ---MVA------PLSAALALGSPGLRGG-----SFNHTTISSRATTAACRAVLVAV--LI    44
tr|Q4QC59|Q4QC59_LEIMA              -----------------------------------------------MLCTLIAAV--LI    11
tr|A4HZD4|A4HZD4_LEIIN              -----------------------------------------------MLCTLIAAV--LI    11
tr|A0A0S4IMV9|A0A0S4IMV9_BODSA      --------------------------MR--------VLII--AA----LCTIVVMHLSMS    20
tr|A0A0S4IJV6|A0A0S4IJV6_BODSA      ------------------------------------------------------------    0
tr|A0A1X0NHQ9|A0A1X0NHQ9_9TRYP      --------------------------MKC----TFSHVLVT-F-----ISLLFIFGCSLS    24
tr|Q38CG9|Q38CG9_TRYB2              ------------MRSAYCRKPSSGFMAHS----NVVVALLS----------LVALHFVPY    34
tr|K2NQ48|K2NQ48_TRYCR              -----------MQEHPFCVKRAPGMALTHRVHGGLAPLLLFSKA----KRVLWLVL-MVS    44
tr|V5BHB3|V5BHB3_TRYCR              ------------------------MAVTHWVHGGVVPPPLL-MA----KRVLWLVL-MMS    30
tr|Q4E1T8|Q4E1T8_TRYCC              ------------------------MAVTHWVHGGVVPPPLP-MA----KRVLWLVL-MMS    30
tr|Q4DB70|Q4DB70_TRYCC              ------------------------MAVTHWVHGGVVPPPLP-MA----KRVLWLVL-MMS    30
tr|A0A3R7N2P8|A0A3R7N2P8_9TRYP      --------------------------MRREARGGAASLELSSM-----KRPLWLLLLVMS    29
tr|A0A3R7KKV6|A0A3R7KKV6_TRYRA      --------------------------MRRRVCGDAVSLQLPIM-----KRVLWLLL-TLS    28
tr|A0A061IV15|A0A061IV15_TRYRA      MFFFVFFFFCCVEEHRFGLKGRPSRVMRRRVCGDAVSLQLPIM-----KGVLWLLL-ALS    54


tr|A0A0M9FY26|A0A0M9FY26_LEPPY      LLCTA------AGLASAAPSSRPFRCLRPFSAMDSFVAVAKADATDNCMFHYFTVTAAAA    72
tr|A4HBZ4|A4HBZ4_LEIBR              LMVAP------S-GAQGAVMPASFRCLRPFSATDTFSNVAKAQLSQNCDYFYFTVEDAKA    97
tr|Q4QC59|Q4QC59_LEIMA              LVVAP------P-GAQAAASPSSFRCLRPFSASATFPNVAKAHPAQNCDYFYFSVEDAKA    64
tr|A4HZD4|A4HZD4_LEIIN              LVVAP------P-GAQAAAGPSSFRCLRPFSATGTFTNVAKAHPTQNCDYFYFSVEDAKA    64
tr|A0A0S4IMV9|A0A0S4IMV9_BODSA      GVVAASSSSSPAPVNTPSPERHPFRCFRPFSALDKFAP-SSMHRRDNCEFQFVQVQDARV    79
tr|A0A0S4IJV6|A0A0S4IJV6_BODSA      ------------------------------------------------------------    0
tr|A0A1X0NHQ9|A0A1X0NHQ9_9TRYP      LVADAT--NNNNNTTTETAALTPFRCMRPFGSLSRFSP-SAMNAAENCKFGSFRILNASA    81
tr|Q38CG9|Q38CG9_TRYB2              YIQGT------VSADSGSNTTVGFRCMRPFASLPRFSS-TSMDAAENCKFKRYNVLNASA    87
tr|K2NQ48|K2NQ48_TRYCR              FYVDV------ASASDTLKSVAPFRCMRPFESLPRFSS-ANMDAGENCRLGPFRVINAKA    97
tr|V5BHB3|V5BHB3_TRYCR              LYVDA------ASASDTLKGVAPFRCMRPFESLPRFSP-ANMDSGENCRFGPFRVINAKA    83
tr|Q4E1T8|Q4E1T8_TRYCC              LYVDA------ASASDTLKGVAPFRCMRPFESLPRFSP-ANMDSGENCRLGPFRVINAKA    83
tr|Q4DB70|Q4DB70_TRYCC              LCVDA------ASASDILKGVAPFRCMRPFESLPRFSP-ANMDSGENCRLGPFRVINAKA    83
tr|A0A3R7N2P8|A0A3R7N2P8_9TRYP      FSLGA------ASTSAAVKDAAPFRCMRPFASLPRFSP-ANMDAAENCKFGSLRVINATA    82
tr|A0A3R7KKV6|A0A3R7KKV6_TRYRA      FFLGA------ASSSTAVKDTAPFRCMRPFGSLPRFSP-ANMDAAENCKFGLLRVINATA    81
tr|A0A061IV15|A0A061IV15_TRYRA      FFLGA------ASSSAAVKDTAPFRCMRPFGSLPRFSP-ANMDAAENCKFGLLRVINATA    107


tr|A0A0M9FY26|A0A0M9FY26_LEPPY      ELRCATTADRGTPPLSSQ-----AGEKAAATSCNVTVRWSMRRADDVAAVLRRQIGTDAD    127
tr|A4HBZ4|A4HBZ4_LEIBR              ELKCTSSSAAAT-------------VVSDDISCSVTVRWSMRRASDVTARLQRQVGTDKD    144
tr|Q4QC59|Q4QC59_LEIMA              ELKCASPSAVTT-------------AVSDEMSCNVTVRVSMRRASDAVSHLQRQVGTDED    111
tr|A4HZD4|A4HZD4_LEIIN              ELKCASPSAATT-------------AVSDEMSCNVTVRMSMRRASDVVPRLQRQVGTDKD    111
tr|A0A0S4IMV9|A0A0S4IMV9_BODSA      QVECEKGTGDTT-----------NENEGGVGGCYATLALDLSVTGQKLIEHE---L----    121
tr|A0A0S4IJV6|A0A0S4IJV6_BODSA      ---------------------------------------------------E---L----    2
```

```
tr|A0A1X0NHQ9|A0A1X0NHQ9_9TRYP      VLNCTHAKSTTMSNIDGQSTQAEENDKMLVPGCNAVIRIVFKMEEENKFGT---------    132
tr|Q38CG9|Q38CG9_TRYB2              VVKCESTGNAT-----GK-----AEPTAVTSTCSVTIHAAFRTMGGGKDDN---------    128
tr|K2NQ48|K2NQ48_TRYCR              MTQCVREYDDE-----DD------TARATSPTCSVTLRVTFQTEGDKKRHHA---G----    139
tr|V5BHB3|V5BHB3_TRYCR              MTQCVRD---D-----DD------TARAASPTCSVTLRVSFQTEGDKRRHRV---G----    122
tr|Q4E1T8|Q4E1T8_TRYCC              MTQCVRD---D-----DD------TTRAASPTCSVTLRVSFQTEGDKRRHRV---G----    122
tr|Q4DB70|Q4DB70_TRYCC              MTQCVRD---D-----DD------TARAASPTCSVTLRVSFQTEGDKRRHRV---G----    122
tr|A0A3R7N2P8|A0A3R7N2P8_9TRYP      RTQCLREDAAR-----GAS--TASEGEKPPPTCSVTLHLTFQTVGDEVDRPV---G----    128
tr|A0A3R7KKV6|A0A3R7KKV6_TRYRA      RTRCVRDDAAR-----GAS--TASNNDKMPPTCSVTLNVTFQAEGNEAYHAV---G----    127
tr|A0A061IV15|A0A061IV15_TRYRA      RTRCVRDDAAR-----GAS--TASNDDKQPPTCSVTLNVTFQAEGNEVYHAV---G----    153

tr|A0A0M9FY26|A0A0M9FY26_LEPPY      YFAANTAELSASSARTAQHSPQQQQQKKRSRAPTELFLTEDEDVWRYAVSLCAKGDSTVQ    187
tr|A4HBZ4|A4HBZ4_LEIBR              YFVHDASTASTATLPT---------RTRNTHQSARVFLTEDDDVWRYTVSLKAKGDSSIH    195
tr|Q4QC59|Q4QC59_LEIMA              YFTHEASTVSTTTSPT---------RSGNTCPSAAPFVTEDEDVWRYAVSLKAKGDSSIH    162
tr|A4HZD4|A4HZD4_LEIIN              YFTHEASTDSTTASPT---------RVGNTRPSAVPFLTEDEDVWRYAVSLKAKGDSSIH    162
tr|A0A0S4IMV9|A0A0S4IMV9_BODSA      --------------PQDSH------------EDVVSGRVRAHYHPYYTLELHALNHSKIT    155
tr|A0A0S4IJV6|A0A0S4IJV6_BODSA      --------------PQDSH------------EDVVSGRVRAHYHPYYTLELHALNHSKIT    36
tr|A0A1X0NHQ9|A0A1X0NHQ9_9TRYP      ---------------TESN------------NGLVE-----NDIYRFSLQLRAFNDSVVH    160
tr|Q38CG9|Q38CG9_TRYB2              ---------------------------------ADA-----ADVYRYSLQLRAINNSVVH    150
tr|K2NQ48|K2NQ48_TRYCR              ------------GSTVAGS------------EETFG-----SEAYRFSLRLRAFNDSVVE    170
tr|V5BHB3|V5BHB3_TRYCR              ------------GAALAGS------------DEAFG-----SEAYRFSLRLRAFNDSIVE    153
tr|Q4E1T8|Q4E1T8_TRYCC              ------------GAALAGS------------DEAFG-----SGAYRFSLRLRAFNDSIVE    153
tr|Q4DB70|Q4DB70_TRYCC              ------------GAALAGS------------DEAFG-----SEAYRFSLRLRAFNDSIVE    153
tr|A0A3R7N2P8|A0A3R7N2P8_9TRYP      ------------VSSVAGH------------RPAVG-----GEAYRFSLRLRAFNDSVVE    159
tr|A0A3R7KKV6|A0A3R7KKV6_TRYRA      ------------VSSVAGH------------EEAFG-----GEAYRFSLRLRAFNDSVVE    158
tr|A0A061IV15|A0A061IV15_TRYRA      ------------VSSVAGH------------KEAFG-----GEAYRFSLRLRAFNDSVVE    184
                                                                             ::: * * ..* :

tr|A0A0M9FY26|A0A0M9FY26_LEPPY      YKGYNGGDGYSMCCDGIVE--GECAWMAAEQG-----DNDLA--------ADEEVNPDFA    232
tr|A4HBZ4|A4HBZ4_LEIBR              YKGFSKGNGYSLCCNALEE--ADCAWMTLEKG-----DNDLD--------CDEAIGSAFA    240
tr|Q4QC59|Q4QC59_LEIMA              YKGFNGGNGYSLCCAALEE--ADCAWITPQKG-----DKDLD--------CDEDVDVGFA    207
tr|A4HZD4|A4HZD4_LEIIN              YKGFNGGNGYSLCCAALEE--ADCAWMTPQKG-----DKDLD--------CDEDVDVRFA    207
tr|A0A0S4IMV9|A0A0S4IMV9_BODSA      YKGFD-GLGYDLCCDFLNESSAHCEWEDTKNT-VAGGDE-SSSSSLPPSEQEDAATQQQR    212
tr|A0A0S4IJV6|A0A0S4IJV6_BODSA      YKGFD-GLGYDLCCDFLNASSAHCEWEDTKKDTVAGGDESSSSSSLPPSEQEDAATQQQR    95
tr|A0A1X0NHQ9|A0A1X0NHQ9_9TRYP      YKGFD-QNGYSMCCDLIQE--GECVWEKNGNLSQTEGKENISKN--------TETSEITT    209
tr|Q38CG9|Q38CG9_TRYB2              YKGFD-QNGYGLCCSFIEG--AKCTWEERETSGVQEA-----------KQEVNATTSQAP    196
tr|K2NQ48|K2NQ48_TRYCR              YKGFD-QNGYSMCCQFIEG--SECSWEMKEKEKEEEGANSTQQESATSQEGDDGRMSHAP    227
tr|V5BHB3|V5BHB3_TRYCR              YKGFD-QNGYSMCCHFIEG--SECSWDVKEEE-----GNSTQQEAATPQKDDDGRASHAP    205
tr|Q4E1T8|Q4E1T8_TRYCC              YKGFD-QNGYSMCCHFIEG--SECSWDVKEEE-----GNSTQQEAATPQKDDDGRASHAP    205
tr|Q4DB70|Q4DB70_TRYCC              YKGFD-QNGYSMCCHFIEG--SECSWEVKEEE-----GNSTQQEAATPQKEDDGRASHAP    205
tr|A0A3R7N2P8|A0A3R7N2P8_9TRYP      YKGFD-QNGYSMCCDLIQG--AECAWAGEGG------SSAAAHSPAVTQEGDGGSTPDAP    210
tr|A0A3R7KKV6|A0A3R7KKV6_TRYRA      YKGFD-QNGYSMCCNFIQS--AECSWDEERK------SSGTPHYSASPQEDNGGNAPDAS    209
tr|A0A061IV15|A0A061IV15_TRYRA      YKGFD-QNGYSMCCNFIQG--AECSWDEERK------SSDTPHYSASPQEDDGGNAPDAA    235
                                    ***:.   **.:**  : ..* *

tr|A0A0M9FY26|A0A0M9FY26_LEPPY      LQHRRMLRGCPLPFPATMPPTAAAEEADDNEADGAFGAAPGARVALTDLDEEDAAEGREE    292
tr|A4HBZ4|A4HBZ4_LEIBR              SQSTRTLRSCPLPLPARRCA--------AGDVDDDCGAWAGAGVVLTDLDE-----DEDS    287
tr|Q4QC59|Q4QC59_LEIMA              SQRSRVLRGCPLPFPAPCSA--------GGGV---FDAPAGARVLLTDLDE-----GESS    251
tr|A4HZD4|A4HZD4_LEIIN              PQRGRVLRGCPLPFPAPRSA--------GDGV---FGASAGARVLLTDLDE-----GEGS    251
tr|A0A0S4IMV9|A0A0S4IMV9_BODSA      IPRRQRLHHCAVTNPLTP----------------------------------------SG    232
tr|A0A0S4IJV6|A0A0S4IJV6_BODSA      IPRGQRLHHCAVTNPLTP----------------------------------------SG    115
tr|A0A1X0NHQ9|A0A1X0NHQ9_9TRYP      PQREPVVVSCPLYNKKIA----------------------------------------KG    229
tr|Q38CG9|Q38CG9_TRYB2              TVKEPAVVSCPLHHG-EE----------------------------------------EH    215
tr|K2NQ48|K2NQ48_TRYCR              QRREAMVVSCPIRNP-VV----------------------------------------PG    246
tr|V5BHB3|V5BHB3_TRYCR              RRREAMVVSCPIRNP-VV----------------------------------------PG    224
tr|Q4E1T8|Q4E1T8_TRYCC              RRREAMVVSCPIRNP-VV----------------------------------------AG    224
tr|Q4DB70|Q4DB70_TRYCC              RRREAMVVSCPIRNP-VV----------------------------------------PG    224
tr|A0A3R7N2P8|A0A3R7N2P8_9TRYP      RRREAVIVSCPLRSP-VV----------------------------------------PG    229
tr|A0A3R7KKV6|A0A3R7KKV6_TRYRA      RRREAMVVSCPIHSQ-VV----------------------------------------PG    228
tr|A0A061IV15|A0A061IV15_TRYRA      RRREAMVVTCPIGGQ-VV----------------------------------------PG    254
                                              :  * :

tr|A0A0M9FY26|A0A0M9FY26_LEPPY      GVFHGQITKPLHRLVEGPWEVTLQMWRRRQRRPTHAAATPSSSSVPTDSSAEAEVLGRVV    352
tr|A4HBZ4|A4HBZ4_LEIBR              GVFHGVITKALHRIVEGPWEMVVQMWRRRQWIPSNGA--EASYTIPTDEGVEAEVLGRIV    345
tr|Q4QC59|Q4QC59_LEIMA              GVFHAVITKPLHRIVEGPWEVMVQMWRRRQQMPRGGR--EDSFSVPADHSIEAEVLGRIM    309
tr|A4HZD4|A4HZD4_LEIIN              GVFHAVITKPLHRFVEGPWEVMVQMWRRRQHMPRVGR--EDSLSVPADHSIEAEVLGRIM    309
tr|A0A0S4IMV9|A0A0S4IMV9_BODSA      RIVSGSIRKPLHKIVAGDWEVRWTLRRGG------------------------ETVGRLT    268
tr|A0A0S4IJV6|A0A0S4IJV6_BODSA      RIVSGSIRKPLHKIVAGDWEVRWTLRRGS------------------------ETVGRLI    151
tr|A0A1X0NHQ9|A0A1X0NHQ9_9TRYP      NVFHGSLIKPLHRLAVTEWEARLEFWRGVK--------------------NEREVLGRLL    269
tr|Q38CG9|Q38CG9_TRYB2              IAFRGSTVKPLHRLVVGNWEARLEFWRGGP--------------------SNREVLGRLL    255
tr|K2NQ48|K2NQ48_TRYCR              IVFHGAITKPLHRLVITEWEARLELWRGRA--------------------HEKVFLGRVL    286
tr|V5BHB3|V5BHB3_TRYCR              IVFHGAITKPLYRLVITDWEARLELWRGRP--------------------HEKVFLGRVL    264
tr|Q4E1T8|Q4E1T8_TRYCC              IVFHGAITKPLHRLVITEWEARLELWRGRP--------------------HEKVFLGRVL    264
tr|Q4DB70|Q4DB70_TRYCC              IVFHGAITKPLHRLVITDWEARLELWRGRP--------------------HEKVFLGRVL    264
tr|A0A3R7N2P8|A0A3R7N2P8_9TRYP      VVFRGVVRKPLHRLLLTEWEARLEFWRGRR--------------------EERETLGRVL    269
tr|A0A3R7KKV6|A0A3R7KKV6_TRYRA      GVFHGVITKPLHRLVVTEWEARLEFWRGQR--------------------QERVMLGRVL    268
tr|A0A061IV15|A0A061IV15_TRYRA      DVFHGVITKPLHRLVVTEWEARLEFWRSQR--------------------EEREMLGRVL    294
                                    . .   * *:::      **       : *                         :**:

tr|A0A0M9FY26|A0A0M9FY26_LEPPY      VPFTVHLAELQREGRVXXXXXXXXXXX-------XXXXXXXXXXXXXXXXXXXXXXXXXX    405
tr|A4HBZ4|A4HBZ4_LEIBR              VPFTLNVADLQGQGRITYVPSMALAVE-------DIAEVVMVTDVKPAVEA---------    389
tr|Q4QC59|Q4QC59_LEIMA              VPFTLNLTELQKQALITQIPSMALTAE-------DVAGVAQAAGENAADET---------    353
tr|A4HZD4|A4HZD4_LEIIN              VPFTLNLTELQKQGRITQMPSMALTVE-------DVADVAQAAGEKAADEA---------    353
tr|A0A0S4IMV9|A0A0S4IMV9_BODSA      VPFHITDEILSGGGTPQQQGDTSAGAA--ATAAASLADSPTTVGTAVV-TVKE-------    318
tr|A0A0S4IJV6|A0A0S4IJV6_BODSA      VPFHITDEMLSGGTSQQVDTSASSAAA--PAASAPLVDSPT-TGTAVV-TVEE-------    200
tr|A0A1X0NHQ9|A0A1X0NHQ9_9TRYP      LPFKLTESDLAFLNESNRDRGTTGVSTNSWNNVGALV-EGKKEG----------------    312
tr|Q38CG9|Q38CG9_TRYB2              VPFRLTDENIAQGNNNTITERSDHTTA-VVVVAEQ----------------GQ-------    291
tr|K2NQ48|K2NQ48_TRYCR              VPFRLTEDDIASAQSNAAAKTTTTTST-AMTATESSK-SGKMSGGVLV-VTED-------    336
```

```
tr|V5BHB3|V5BHB3_TRYCR              VPFRLTEDDIASAQSNAAATTT---TT-AMTVTDSSA-SSKMAGGVLL-ATED-------    311
tr|Q4E1T8|Q4E1T8_TRYCC              VPFRLTEDDIASAQSNAAATTT---TT-AMTATDSSA-SGKMAGGVLL-VTED-------    311
tr|Q4DB70|Q4DB70_TRYCC              VPFRLTEDDIASAQFNAAATTT---TT-AMTATDSSA-SGKMAGGVLL-VTED-------    311
tr|A0A3R7N2P8|A0A3R7N2P8_9TRYP      VPFRLTEEDVASAYAADAAATT---SR-AVTAIDDSP-SGEQDVVAPL-ETGE-------    316
tr|A0A3R7KKV6|A0A3R7KKV6_TRYRA      LPFRLTEDDIASAYTSDAAATT-LTAT-AVAAIDDST-SGGRAGAAPI-ETGD-------    317
tr|A0A061IV15|A0A061IV15_TRYRA      LPFRLTEDDIASAYTSDAAATT-LTAT-AVAAIDDST-SGGRAGAAPT-ETGD-------    343
                                    :** :    :

tr|A0A0M9FY26|A0A0M9FY26_LEPPY      XXXXXXXXXXXXXXXXXXXXXXXXXXXXXXXXXXXXXXXLEEDVVEVAPAPAPPSSVDAG    465
tr|A4HBZ4|A4HBZ4_LEIBR              ------------------------------------------------------------    389
tr|Q4QC59|Q4QC59_LEIMA              ------------------------------------------------------------    353
tr|A4HZD4|A4HZD4_LEIIN              ------------------------------------------------------------    353
tr|A0A0S4IMV9|A0A0S4IMV9_BODSA      ---------------------------------------------------------QQR    321
tr|A0A0S4IJV6|A0A0S4IJV6_BODSA      ---------------------------------------------------------QR-    202
tr|A0A1X0NHQ9|A0A1X0NHQ9_9TRYP      ----------------------------------------------------------VK    314
tr|Q38CG9|Q38CG9_TRYB2              ---------------------------------------------------------LTG    294
tr|K2NQ48|K2NQ48_TRYCR              ---------------------------------------------------------VIK    339
tr|V5BHB3|V5BHB3_TRYCR              ---------------------------------------------------------VVK    314
tr|Q4E1T8|Q4E1T8_TRYCC              ---------------------------------------------------------VVK    314
tr|Q4DB70|Q4DB70_TRYCC              ---------------------------------------------------------VVK    314
tr|A0A3R7N2P8|A0A3R7N2P8_9TRYP      ---------------------------------------------------------NVG    319
tr|A0A3R7KKV6|A0A3R7KKV6_TRYRA      ---------------------------------------------------------VIG    320
tr|A0A061IV15|A0A061IV15_TRYRA      ---------------------------------------------------------VIG    346


tr|A0A0M9FY26|A0A0M9FY26_LEPPY      EAEDTGDL----------    473
tr|A4HBZ4|A4HBZ4_LEIBR              -----GDL----------    392
tr|Q4QC59|Q4QC59_LEIMA              -----GDL----------    356
tr|A4HZD4|A4HZD4_LEIIN              -----GDL----------    356
tr|A0A0S4IMV9|A0A0S4IMV9_BODSA      EHEQGDKVTHAPAKVEEL    339
tr|A0A0S4IJV6|A0A0S4IJV6_BODSA      -EHGSDKVTHAPAKVEEL    219
tr|A0A1X0NHQ9|A0A1X0NHQ9_9TRYP      EFNREDDL----------    322
tr|Q38CG9|Q38CG9_TRYB2              SRG---DL----------    299
tr|K2NQ48|K2NQ48_TRYCR              ERNGSEDL----------    347
tr|V5BHB3|V5BHB3_TRYCR              ERNGYEDL----------    322
tr|Q4E1T8|Q4E1T8_TRYCC              ERNGYEDL----------    322
tr|Q4DB70|Q4DB70_TRYCC              ERNGYEEL----------    322
tr|A0A3R7N2P8|A0A3R7N2P8_9TRYP      ERKGSADL----------    327
tr|A0A3R7KKV6|A0A3R7KKV6_TRYRA      KRKGSGDL----------    328
tr|A0A061IV15|A0A061IV15_TRYRA      KRKGSGDL----------    354
                                             .:
```

### 08. ER-implicated kinetoplastid ATPase (ERIKA)

These proteins belong to the greater Torsin (ClpA/ClpB) family of chaperone proteins. They are typically found at the nuclear membrane lumen as well as the ER, but usually have no KDEL signals. All parazitic kinetoplastids seem to have multiple torsin paralogues, and only one subtype (that we now cover under the acronym ERIKA) has the typical KDEL-like signal. While the KDEL-like signal is seen in Bodo saltans, it is not yet found in the early-branching kinetoplastid Perkinsela sp. The only other eukaryotic torsin protein with a clear KDEL-like retention signal was detected in the unicellular amoeba Acanthamoeba castellanii (the motif probably evolved independently).

```
tr|D7G3V5|D7G3V5_ECTSI              -------------------------------------------MAMIVSQAQMA------    11
tr|Q55FE2|Q55FE2_DICDI              -----------------------------MGRYFYLQLLFISILFLK--SYSAQN-N---    25
tr|L8GYA8|L8GYA8_ACACA              ------------------------MMQARL----RSLVLA---LVLLPALVMAA------    23
tr|A0A0L1L0A3|A0A0L1L0A3_9EUGL      ----MP------------------LPAFSMKAFILNLSLL---MTLSITVRSSNEDATRI    35
tr|A0A0S4KNW8|A0A0S4KNW8_BODSA      ------------------------MLAHK---GMLVATVA---VCMCI--LA--------    20
tr|A0A422N6P1|A0A422N6P1_TRYRA      ------------------------MGSCATA-TWVLAFLVVV-VAALTPWVA--------    26
tr|A0A3R7NQX6|A0A3R7NQX6_9TRYP      ----MP-----------------QMGPCATS-TCLLALLAAV-VAALVPWVA--------    29
tr|S9U3X1|S9U3X1_9TRYP              ------------------------------------------------------------    0
tr|S9U3J6|S9U3J6_9TRYP              ------------------------MNVGR-----TGAAFVVFLLALCVYLYG--------    23
tr|A0A0N0VF20|A0A0N0VF20_LEPPY      ------------------------MRSFL---ITALSIVCVCLACI---VYV--------    22
tr|A4I131|A4I131_LEIIN              ------------------------MHVFT----RILAVFVIVLAALLIPQFL--------    24
tr|Q4QAA8|Q4QAA8_LEIMA              ------------------------MHLFT----RILAVFVVVLVALLIPRFL--------    24
tr|A0A7G2CLT7|A0A7G2CLT7_9TRYP      ----MT------HT-RPFRV----------WRFSILLLLVSS---VLVSVRA--------    28
tr|S9VXS8|S9VXS8_9TRYP              ----MP-----------------RRPRVTHCGFALVLVLCMGVLAVAISLTAAA--AA--    35
tr|A0A0M9FWU4|A0A0M9FWU4_LEPPY      ----MP-----CHARREKRRAVPQLSYSSTV---RIAVLSLSLLVLLLSCVALP------    42
tr|A4H5R3|A4H5R3_LEIBR              MASFLPPQRQLSHISRRQPGWIPPSATAGKT---ALLILFAAAFFLALGSATAA--EGQQ    55
tr|Q4QHU2|Q4QHU2_LEIMA              -----------------------------MT---VILVVCTVVFVLVLGSATVA--DGRP    26
tr|A4HU03|A4HU03_LEIIN              -----------------------------MT---VALIVCSVVFVLVLGSATVA--GGRP    26
tr|Q382P2|Q382P2_TRYB2              ----MP-HSGYPFG-SSSVVGVEGDDRHSSVSLFVILFLCVLLTAVEPSVA---------    45
tr|A0A1X0NMJ2|A0A1X0NMJ2_9TRYP      --MIFF-SR------RSSS-----------III-AV--IFVVVLLALPTCLA--------    29
```

```
tr|Q4D6G4|Q4D6G4_TRYCC              ----MS-AR------RLFTRGIP------VATMAIA--VCIVIFLIFPA-----------    30
tr|Q4D903|Q4D903_TRYCC              ----MS-AR------RLFTRGIP------VATMAIA--VCIVIFLIFPA-----------    30
tr|A0A422ND04|A0A422ND04_9TRYP      ----MP-AR------RHSPRGVS------AVATVVA--ALILLLLIFPA-----------    30
tr|A0A3R7RCD3|A0A3R7RCD3_TRYRA      ----MF-AR------CRSLRGAP------AVTTLVA--ALTVLLLVFPA-----------    30


tr|D7G3V5|D7G3V5_ECTSI              ------ASVEE---------EPAAT-ATSAATI--SVLDA-------VRVMGSLTWWGVD    46
tr|Q55FE2|Q55FE2_DICDI              --DKIKNDD-------DSIYVNKIFGDCKEIYYNNKLNNDENYKNYIGSLFGTYFNNNNN    76
tr|L8GYA8|L8GYA8_ACACA              ------FDDCECNESWTSFACTRCFGPCPRTLE--PV-KRFEELTTK-------------    61
tr|A0A0L1L0A3|A0A0L1L0A3_9EUGL      NDDLTLLSYSNQTIYANKRICPCRP-LCSRGTG--FI-N--PICWGIPSLRY-RYQWSND    88
tr|A0A0S4KNW8|A0A0S4KNW8_BODSA      ----------EMVLGWSPSECPPPP-LCEGRNA--SY-QQQAWCQLGRDPLS--------    58
tr|A0A422N6P1|A0A422N6P1_TRYRA      ---R------VELHPGEAGDCTLRP-LCSSHP---TL-VKRLMCFLFQYPVD--------    64
tr|A0A3R7NQX6|A0A3R7NQX6_9TRYP      ---H------VELHPGEADDCMLRP-LCSSHP---TL-VKGLMCLLFRYPVD--------    67
tr|S9U3X1|S9U3X1_9TRYP              ---------------------MDVP-MCSSPSL--TA-LGHLYCTLAPTSLL--------    27
tr|S9U3J6|S9U3J6_9TRYP              --ASNTFSEEALLYRSPVQDCMDVP-MCSSPSL--TA-LGHLYCTLAPTSLL--------    69
tr|A0A0N0VF20|A0A0N0VF20_LEPPY      ----ETV---GSRLDCTALYCPAPP-LCDQATR--TF-SQRVVCGLGKSPLK--------    63
tr|A4I131|A4I131_LEIIN              --EPIYV---IPAADAVAQFCSEPP-LCARKDR--TM-KQQLLCTIRSAPLK--------    67
tr|Q4QAA8|Q4QAA8_LEIMA              --EPIYV---IPAADAVAPFCSEPP-LCARKDR--TM-KQQLLCTIRSAPLK--------    67
tr|A0A7G2CLT7|A0A7G2CLT7_9TRYP      ------APASHRERILASQACPAPP-LCEGSRL---SAAHIKYCNPLSATAVRFFRFTTN    78
tr|S9VXS8|S9VXS8_9TRYP              --EDTSEHVPHYGGGARRSLCPAGP-ICSRAHT--PFALKFKYCNKVTSFFTSFFGGGGG    90
tr|A0A0M9FWU4|A0A0M9FWU4_LEPPY      --QLARASPTVSTITSTSSLCPRCP-LCARRQT--SLRDRLKYCHSTSSYALSLFSSGGS    97
tr|A4H5R3|A4H5R3_LEIBR              QQRPLKATAAGASSHVSNSLCPRCP-LCSRPGT--PMKDWLKYCHTTTGYIFSLFSWGKS   112
tr|Q4QHU2|Q4QHU2_LEIMA              KQPISKGNVAEASSHASSSLCPRCP-LCSRPGT--PIKERLKYCHATSGYIFSLFSWGTS    83
tr|A4HU03|A4HU03_LEIIN              KQPISKGTTAGASSHASSSLCPRCP-LCSRSGT--PIKERLKYCHTTSGYIISLFSWGTS    83
tr|Q382P2|Q382P2_TRYB2              -GADVRSTVDEVVADTSEPTCPPAP-VCSRYKW---LSSSQLYCWMSKLP----------    90
tr|A0A1X0NMJ2|A0A1X0NMJ2_9TRYP      LQGKEQDKHFSSETQLREWKCPSSP-FCSKGNA--PLWTKLKYCLFSSIP----------    76
tr|Q4D6G4|Q4D6G4_TRYCC              --SSLRPLPVAAEGEEVERRCPRCP-ICSARGT--PFFLRLRYCVLTSLP----------    75
tr|Q4D903|Q4D903_TRYCC              --SSLRPLPVAAEGEEVERHCPRCP-ICSARGT--PFFLRLRYCVLTSLP----------    75
tr|A0A422ND04|A0A422ND04_9TRYP      --SPLRPSSVAAEVELVEWRCPRCP-ICAARGT--PLLLRLRHCVLTALP----------    75
tr|A0A3R7RCD3|A0A3R7RCD3_TRYRA      --SPLRPSFVSAEGEVVEWHCPRCP-ICAVRGT--PLLLQLRYCVLTSLP----------    75


tr|D7G3V5|D7G3V5_ECTSI              LAMET------------AGYGVCPFADDPVEA--ARRALQDGILAQPLALESLDGALSSW    92
tr|Q55FE2|Q55FE2_DICDI              KDNNNNN-------------NNCKFKKD---KNELLKSFKQYFEGFVSFQDDAIKEIESI   120
tr|L8GYA8|L8GYA8_ACACA              ----------------------------------------PGPAKKLMGQERAAVLIADA    81
tr|A0A0L1L0A3|A0A0L1L0A3_9EUGL      NIG-------------------------------DIHSCLYSLFMQIKGQQRAIETLVGH   117
tr|A0A0S4KNW8|A0A0S4KNW8_BODSA      -------------------------HNSVEYIRMVQNRLSYFLSRSLLGQDHLAPRILAD    93
tr|A0A422N6P1|A0A422N6P1_TRYRA      -------------------------VDRSDYKQQVLVRLRRLTQRNLKGQTHVVEGVIRG    99
tr|A0A3R7NQX6|A0A3R7NQX6_9TRYP      -------------------------VDRGDYKQQVLVRLRRLTQRSLKGQRHVVEGVTRS   102
tr|S9U3X1|S9U3X1_9TRYP              --------------------------SRPGYADYVYQQLRARAHHSVKGQYI-IDPVLQL    60
tr|S9U3J6|S9U3J6_9TRYP              --------------------------SRPGYADYVYQQLRARAHHSVKGQYI-IDPVLQL   102
tr|A0A0N0VF20|A0A0N0VF20_LEPPY      -------------------------PGSKEFRAYVVTRLHGFVKESLKGQLM-APAVVEM    97
tr|A4I131|A4I131_LEIIN              -------------------------PASPVYRQYVVSRLKRYVDGSIRGQSV-APSVIER   101
tr|Q4QAA8|Q4QAA8_LEIMA              -------------------------PASPAYRQYVVSRLKRYVDGSIRGQSV-AASVIER   101
tr|A0A7G2CLT7|A0A7G2CLT7_9TRYP      KLSSAVSHAPH--TKTEYRPLQLPFPTDAVIRKNAKRNVEEYLKNYLAGQEHLLSPITRL   136
tr|S9VXS8|S9VXS8_9TRYP              ----------GGKIDFRHFPLQLPFATDTAMRQLAKAGLEHLLLMRLIGQDHLQQELLDV   140
tr|A0A0M9FWU4|A0A0M9FWU4_LEPPY      ELATATAAGASYREQRHRYPLELPFATDDAVRAQCVRDATDLLASRVRGQPQVVKPLLEV   157
tr|A4H5R3|A4H5R3_LEIBR              TLAGTQRTGGSIREQRQLYPLELPFATDDAVRRQCMRDLQDLLLARVRGQPQVVIPLLDV   172
tr|Q4QHU2|Q4QHU2_LEIMA              ALAGKQRTGVSNREQRQLYPLELPFATDDAVRRQCMRDMQDLLLARVRGQPQVVMPLLDV   143
tr|A4HU03|A4HU03_LEIIN              ALAGKQRTGVSNREQRQLYPLELPFATDDAVRRQCMRDMQDLLLARVRGQPQVVMPLLDV   143
tr|Q382P2|Q382P2_TRYB2              ---------------SRLVSRELPFATDDKARRLAISNVEELLRTRLVGQGHLTEAIASL   135
tr|A0A1X0NMJ2|A0A1X0NMJ2_9TRYP      ---------------LAKIRQELPFATSDAARRIAMENVEELLFTHVVGQDHVTHAIAEA   121
tr|Q4D6G4|Q4D6G4_TRYCC              ---------------AVLVKREFSVEKDEKLRHRVMEDVKELLLTRLVGQEHVKGAILEA   120
tr|Q4D903|Q4D903_TRYCC              ---------------AVLVKREFSVEKDEKLRHRVMEDVKELLLTRLVGQEHLKGVILEA   120
tr|A0A422ND04|A0A422ND04_9TRYP      ---------------AALVRREFALAKDEALRQRVMKDVEELLLTRVAGQEHVKGAILEA   120
tr|A0A3R7RCD3|A0A3R7RCD3_TRYRA      ---------------AALVRQEFAFSKQEALRHRVMNDVEELLFTRVAGQEHVKKAILEA   120
                                                                                          :

tr|D7G3V5|D7G3V5_ECTSI              HYSRQSDRYEPLVVALTGSTGTGKTETAWVLADALLTKRCRITGG---TRDIPRGLLVLN   149
tr|Q55FE2|Q55FE2_DICDI              IYRKYLHPDTKVVMHLFGDHGIGKTLSSKLVSRVLF-ENGS--------GLEGDGLLLIN   171
tr|L8GYA8|L8GYA8_ACACA              LR--AASGKKPLTFHFCGENGVGKSHTALLLAEAYF-AYKDK------KTDMYKGLLWIS   132
tr|A0A0L1L0A3|A0A0L1L0A3_9EUGL      INRKLYDPRASFLIHLAGDNGVGKTMSAALLSIAIS-LYPHKAH-----RNAGETLLIIS   171
tr|A0A0S4KNW8|A0A0S4KNW8_BODSA      VIHKLRRPNEPLILHFAGDNGVGKTSTAQLISASMS-FRCHRSN-TGYYCGLGDASLSLS   151
tr|A0A422N6P1|A0A422N6P1_TRYRA      IAAKLENPDKPLVLHFAGDNGVGKTTLAQLISLALG-LRCHD-----AACTTGDSALVLS   153
tr|A0A3R7NQX6|A0A3R7NQX6_9TRYP      LAAKLENPDKPLVLHFAGDNGVGKTTLAQLISLALG-LRCRD-----AACTTGDSTLVLS   156
tr|S9U3X1|S9U3X1_9TRYP              VKHKLRHRDHPLILHFAGDNGVGKTSLAEVISLAMA-LRCKSA----EDCHIGDSTLELS   115
tr|S9U3J6|S9U3J6_9TRYP              VKHKLRHRDHPLILHFAGDNGVGKTSLAEVISLAMA-LRCKSA----EDCHIGDSTLELS   157
tr|A0A0N0VF20|A0A0N0VF20_LEPPY      VEYKLHHPYEPMVLHFAGDNGVGKTRLAELISLAYG-QRCGD-----ELCTVGDSTLVLS   151
tr|A4I131|A4I131_LEIIN              VRYKLAHMHEPMILHFAGDNGVGKTRLAELISLAIG-QKCGD-----ATCSIGDTTLVLS   155
tr|Q4QAA8|Q4QAA8_LEIMA              VRYKLAHVHEPMVLHFAGDNGVGKTRLAELISLAMG-QKCGD-----ATCSIGDTTLVLS   155
tr|A0A7G2CLT7|A0A7G2CLT7_9TRYP      VQYKVDHPTEPLVLHLAGDNGVGKTFTAKLISLALS-LYCGAEGRDTLPCAHGDALLIVA   195
tr|S9VXS8|S9VXS8_9TRYP              LFKKLYFPATPLVLHLAGDNGVGKSYTAKLISLALS-LRCGADGDG--GCEAGDALLTIS   197
tr|A0A0M9FWU4|A0A0M9FWU4_LEPPY      LRRKLAFPREPVVVHLAGDNGVGKTHTARLVSQALS-LRCAADR-D--VCDAGDNLLVIA   213
tr|A4H5R3|A4H5R3_LEIBR              LRHKLAYPREPVVVHLAGDNGVGKSHTARLVSEALS-LRCALDR-D--VCDAGDNLLIIS   228
tr|Q4QHU2|Q4QHU2_LEIMA              LRRKLAYPREPVVVHLAGDNGVGKTHTARLVSQALS-LRCAPDR-D--VCDAGDNLLTIA   199
tr|A4HU03|A4HU03_LEIIN              LRRKLAYPREPVVVHLAGDNGVGKTHTARLVSQALS-LRCAPDR-D--VCDAGDNLLTIA   199
tr|Q382P2|Q382P2_TRYB2              MRKKLSYPHEPLVLHFAGDNGVGKTHTARLLSLATS-LRCANSRGR--QCDSGDNMLVIS   192
tr|A0A1X0NMJ2|A0A1X0NMJ2_9TRYP      IRQKMASPGDPLVLHFAGDNGVGKTHTARLLSLALS-LRCAQAR-P--QCDVGDNMLVIS   177
tr|Q4D6G4|Q4D6G4_TRYCC              VRRKLIYPRDPLVLHFAGDNGVGKTHTARLLSLATS-PHCAPSR-P--ACDMGENMLVIS   176
tr|Q4D903|Q4D903_TRYCC              VRRKLIYPRDPLVLHFAGDNGVGKTHTARLLSLATS-PHCAPSR-P--ACDMGENMLVIS   176
tr|A0A422ND04|A0A422ND04_9TRYP      VRRKLSYPQDPLVLHFAGDNGVGKTHTARLLSLATS-LRCAPSR-P--ACDMGENMLVIS   176
tr|A0A3R7RCD3|A0A3R7RCD3_TRYRA      VRRKLSYPEDPLVLHFAGDNGVGKTHTARLLSLATS-LRCAPSR-P--MCDAGDNMLIIS   176
                                             . . : *. * **:  : :::                       * :

tr|D7G3V5|D7G3V5_ECTSI              GADYMVAAKVEEY---------------------Q---SLIRRKLGQRLEYCGGNVVVLF   185
tr|Q55FE2|Q55FE2_DICDI              GEEFRIIEQQPPTNNNDNNENTKIEDQEYLNIKIQHLRDKLYNTIINKLIECPY-SVIVF   230
tr|L8GYA8|L8GYA8_ACACA              GKQYQMAKSEEEI--------------------KAAREYIHEQIIDHLATCPQ-AIIVI   170
tr|A0A0L1L0A3|A0A0L1L0A3_9EUGL      GSEFASVSGDSAEE-------------------RQLVTRQIQSRLVTHVQKFSK-CVVLF   211
```

```
tr|A0A0S4KNW8|A0A0S4KNW8_BODSA         GINYHGVSPEEFR---------------------K----SIVPQVLAFAERHPH-GLVIF    185
tr|A0A422N6P1|A0A422N6P1_TRYRA         GVSYDGYSVQEFR---------------------R----VVVQRIVQHVLRFPKNGVVIV    188
tr|A0A3R7NQX6|A0A3R7NQX6_9TRYP         GVSYDGYSAQEFR---------------------R----VVVPRIVQHALRFPKNGVVIV    191
tr|S9U3X1|S9U3X1_9TRYP                 GAAYDALTVAEFR---------------------R----FVVGAVTSHLQRHPRNAVIII    150
tr|S9U3J6|S9U3J6_9TRYP                 GAAYDALTVAEFR---------------------R----FVVGAVTSHLQRHPRNAVIII    192
tr|A0A0N0VF20|A0A0N0VF20_LEPPY         GTGYDGLSTAEFR---------------------K----AVVELVTRHARRHPRDGVVVI    186
tr|A4I131|A4I131_LEIIN                 GTSYDGMTVAEFR---------------------N----AVVPVVVRHAQRYPDNGVVIF    190
tr|Q4QAA8|Q4QAA8_LEIMA                 GTSYDGMTVAEFR---------------------N----AVVPVVVRHAQRYPDNGVVIF    190
tr|A0A7G2CLT7|A0A7G2CLT7_9TRYP         CSSLRTLPVAQAR---------------------E----IVVTQVLEFAVVHPH-GVVLL    229
tr|S9VXS8|S9VXS8_9TRYP                 GTAYDNEPVALAR---------------------A----RIVEQVTDFSTRHPH-GVVLL    231
tr|A0A0M9FWU4|A0A0M9FWU4_LEPPY         GTGFDGMPVSEAR---------------------G----RIVRQVTAHMEHYPH-GVVLI    247
tr|A4H5R3|A4H5R3_LEIBR                 GTGFDGLSVAEAR---------------------Q----RIVGQIVAHVARYPH-GVVLI    262
tr|Q4QHU2|Q4QHU2_LEIMA                 GTGFDGLSVAEAR---------------------Q----RIIGQIIAHTDCYPH-GVVLI    233
tr|A4HU03|A4HU03_LEIIN                 GTGFDGLSVAEAR---------------------Q----RIIGQIITHTDRYPH-GVVLI    233
tr|Q382P2|Q382P2_TRYB2                 GTGFGGLEGRDGL---------------------N----LLVRKITEHQRKYPH-GIVLL    226
tr|A0A1X0NMJ2|A0A1X0NMJ2_9TRYP         GTSFDELDITEAR---------------------Q----SIVRRITAHQRYYPH-GIILI    211
tr|Q4D6G4|Q4D6G4_TRYCC                 GTGFDGMPIEDAR---------------------T----RIIQRVTAHQKVYPH-GIVLL    210
tr|Q4D903|Q4D903_TRYCC                 GTGFDGMPIEDAR---------------------T----RIIQRVTAHQKVYPH-GIVLL    210
tr|A0A422ND04|A0A422ND04_9TRYP         GTGFDGMPVADAR---------------------L----RIMQRVTAHQKAYPH-GIVLL    210
tr|A0A3R7RCD3|A0A3R7RCD3_TRYRA         GTGFDGMPIADAR---------------------L----RIMQRITAHQKAYPH-GIVLL    210
                                                                                    :   :            :::.

tr|D7G3V5|D7G3V5_ECTSI                 DELQKAAPGTLDALAEAMSEHPRVTFE---------------------------------    212
tr|Q55FE2|Q55FE2_DICDI                 DEIQKIDPYIISVIEPFLDGATITISDEKKTTT---------------------------    263
tr|L8GYA8|L8GYA8_ACACA                 DEAEMMRADILRVVGAFMDDSQTTVSSLKDPSK---------------------------    203
tr|A0A0L1L0A3|A0A0L1L0A3_9EUGL         DEVTQMHPVLLQELNPLFSAIEHGESIN--------------------------------    239
tr|A0A0S4KNW8|A0A0S4KNW8_BODSA         NDMTELSPAQANVLMPLLGRSKHFPEDTHQ------------------------------    215
tr|A0A422N6P1|A0A422N6P1_TRYRA         NDLGALHPDLVRVLLPLLGRAPSFPEAS--------------------------------    216
tr|A0A3R7NQX6|A0A3R7NQX6_9TRYP         NDLGALHPDLVRVLLPLLGRAPSFPEAP--------------------------------    219
tr|S9U3X1|S9U3X1_9TRYP                 NELTSLHPEKVMVLLPLLGRGSYFPEHP--------------------------------    178
tr|S9U3J6|S9U3J6_9TRYP                 NELTSLHPEKVMVLLPLLGRGSYFPEHP--------------------------------    220
tr|A0A0N0VF20|A0A0N0VF20_LEPPY         NELSSLEPGKVRVLLPLLGRASEFPEHF--------------------------------    214
tr|A4I131|A4I131_LEIIN                 NELTSLEPSKVRVLSPLLGRGTSFPENP--------------------------------    218
tr|Q4QAA8|Q4QAA8_LEIMA                 NELTSLEPNKVRVLSPLLGRGTSFPESL--------------------------------    218
tr|A0A7G2CLT7|A0A7G2CLT7_9TRYP         DDLSALHPELIQGLSPLFGRAPYFPEQLLKEDK---------------------------    262
tr|S9VXS8|S9VXS8_9TRYP                 DDVTAMDPELVRSLAPLLGRAAYFSDQLFENATAPDSAAEP-------------------    272
tr|A0A0M9FWU4|A0A0M9FWU4_LEPPY         DDLTAMEPALVAALAPLFGRASHFAEQLTDPQADSTHSSNVNN---------------DN    292
tr|A4H5R3|A4H5R3_LEIBR                 DDLTAMDPVLVSVLAPLFGRASHFPEQLARSSSATAARVRHTTAKPLQTAHQWMKVFGRD    322
tr|Q4QHU2|Q4QHU2_LEIMA                 DDLTAMDPSLVSALAPLFGRAAHFPEQLTRSSSATAAGARHAAAKSLQTIHASTKGLGPD    293
tr|A4HU03|A4HU03_LEIIN                 DDLTAMDPSLVSALAPLFGRAAHFPEQLTHPSTATAAGARHAAAKSLQTMHESTKGLGPD    293
tr|Q382P2|Q382P2_TRYB2                 DDLNAMHPSLVALLAPLFGRADRFEGQAAD------------------------------    256
tr|A0A1X0NMJ2|A0A1X0NMJ2_9TRYP         DDLTAMHPKLVAALAPLFGRAERFNEQPEN------------------------------    241
tr|Q4D6G4|Q4D6G4_TRYCC                 DDLAAMHPKLVAALAPLLGRAERFEEQPED------------------------------    240
tr|Q4D903|Q4D903_TRYCC                 DDLAAMHPKLVAALAPLLGRAERFEEQPEG------------------------------    240
tr|A0A422ND04|A0A422ND04_9TRYP         DDLTAMHPKLVAALAPLLGRAERFEEQPAG------------------------------    240
tr|A0A3R7RCD3|A0A3R7RCD3_TRYRA         DDLTAMHPKLVAALAPLLGRAERFEEQPEG------------------------------    240
                                       ::                :   :.

tr|D7G3V5|D7G3V5_ECTSI                 ------------------------------------RGGQNVSVDSSRVVFLLVSDVGAE    236
tr|Q55FE2|Q55FE2_DICDI                 -----------------------------------------TKINTSLGTYILTSDFDKE    282
tr|L8GYA8|L8GYA8_ACACA                 ------------------------------------------RVNTKEAIIILISDFGRD    221
tr|A0A0L1L0A3|A0A0L1L0A3_9EUGL         ------------------------------------------GVPMTGVFCFLTSDFGSN    257
tr|A0A0S4KNW8|A0A0S4KNW8_BODSA         ------------------------------------------RVDLHKLMVVVTTDFGKQ    233
tr|A0A422N6P1|A0A422N6P1_TRYRA         ------------------------------------------NVPLGRLVVIVTTDFGRQ    234
tr|A0A3R7NQX6|A0A3R7NQX6_9TRYP         ------------------------------------------GVPLGRLFVIVTTDFGRQ    237
tr|S9U3X1|S9U3X1_9TRYP                 ------------------------------------------QVSLRGAIVIITTDLGRE    196
tr|S9U3J6|S9U3J6_9TRYP                 ------------------------------------------QVSLRGAIVIITTDLGRE    238
tr|A0A0N0VF20|A0A0N0VF20_LEPPY         ------------------------------------------DVKISTQLVVLTTDFGRE    232
tr|A4I131|A4I131_LEIIN                 ------------------------------------------GVSIAPLLVILTTDFGRE    236
tr|Q4QAA8|Q4QAA8_LEIMA                 ------------------------------------------GVSIAPLLVILTTDFGRE    236
tr|A0A7G2CLT7|A0A7G2CLT7_9TRYP         ---------------------------------------KEKLVSLSNLLVIITTDFGKQ    283
tr|S9VXS8|S9VXS8_9TRYP                 ------------------------------------RLLRRLGPSLRHLLVIVTTDFGKQ    296
tr|A0A0M9FWU4|A0A0M9FWU4_LEPPY         HLRGSSSQKGL-----------------LSWAWRPFSSPLKPPPPLSQLLVFITTDFGRQ    335
tr|A4H5R3|A4H5R3_LEIBR                 GGRGSAK-EGNDRKGEAAAGIDGAPPAASEMTSSLSRSGQPPPPPLSQLLVFITTDFGRQ    381
tr|Q4QHU2|Q4QHU2_LEIMA                 GGTGSAAKKGESEKWEAAADTGGAQPAASEMASSSLSSGQP-PPPLSQLIIFITTDFGRQ    352
tr|A4HU03|A4HU03_LEIIN                 GGRGSAAKKGEDEKREAAAGTGGAQPSTSEMASSSLSSGQP-PPPLSQLIIFVTTDFGRQ    352
tr|Q382P2|Q382P2_TRYB2                 ------------------------------------------LPSLKELTVIVTTDFGKQ    274
tr|A0A1X0NMJ2|A0A1X0NMJ2_9TRYP         ------------------------------------------GPSLAQLTVVVTTDFGQQ    259
tr|Q4D6G4|Q4D6G4_TRYCC                 ------------------------------------------TPSLAQLIVVVTTDFGKQ    258
tr|Q4D903|Q4D903_TRYCC                 ------------------------------------------TPSLAQLIVIVTTDFGKQ    258
tr|A0A422ND04|A0A422ND04_9TRYP         ------------------------------------------TPSLAQLLVVVTTDFGQQ    258
tr|A0A3R7RCD3|A0A3R7RCD3_TRYRA         ------------------------------------------TPSLAQLLVVVTTDFGQQ    258
                                                                                     .: :*.. :

tr|D7G3V5|D7G3V5_ECTSI                 GVNAAVLRYRKRSDVVPGALQSAVKRSLDEQWERLRFGKMVDKVVPYLPMDPASNLLVVE    296
tr|Q55FE2|Q55FE2_DICDI                 GMTYNQSIL---------ELKRKATAMFKSIYGDSKFSKLVTESLPFLPLSKSNSKLQII    333
tr|L8GYA8|L8GYA8_ACACA                 EIRTGDSWD---------EIAERVHRETKAILQEDLMVQRIQYHIPFSP-----------    261
tr|A0A0L1L0A3|A0A0L1L0A3_9EUGL         GKTLGMTST---------EVRTLVWEVIQETYKYTPA-LKKANVIPFLALSPADYQDAIR    307
tr|A0A0S4KNW8|A0A0S4KNW8_BODSA         GRTRGKSIE---------ELQQMVEQEVRGTFGALAG--SYLRTYAFIPATLPAVRDIVR    282
tr|A0A422N6P1|A0A422N6P1_TRYRA         GRTRGKSLM---------EMRRIVEDDFKSLYSQLSS--SMIETFPFLPAALDTAKEIVR    283
tr|A0A3R7NQX6|A0A3R7NQX6_9TRYP         GRTQGKSLA---------EMRRLVEEDFKSLYSQLSS--SMIETFPFLPAALDTAKEIVR    286
tr|S9U3X1|S9U3X1_9TRYP                 GRTRGKNLA---------QMRALIESDFQDLYSKLST--SYLHTFPFLPITLDAATDIVR    245
tr|S9U3J6|S9U3J6_9TRYP                 GRTRGKNLA---------QMRALIESDFQDLYSKLST--SYLHTFPFLPITLDAATDIVR    287
tr|A0A0N0VF20|A0A0N0VF20_LEPPY         GRTRGKGLS---------EMRSFINSEFTDLYSAQFA--SHVRTLPFLPISLDTAGEIVR    281
tr|A4I131|A4I131_LEIIN                 GRTRGKSLF---------EMRAFITDEFADLYSKEAA--SHVRTFPFLPISLSTAGDIVR    285
tr|Q4QAA8|Q4QAA8_LEIMA                 GRTRGKSLF---------EMRAFINDEFADLYSKEAA--SHVRTFPFLPISLSTAGDIVR    285
tr|A0A7G2CLT7|A0A7G2CLT7_9TRYP         GVTIGKSLD---------EIKALVDKDFASLYGSLLT--AHIYTFPYLAFNEQMGMEMIR    332
tr|S9VXS8|S9VXS8_9TRYP                 GRTVGLTTS---------EIRAMVLEEFSALYGSLLP--AYTRTFIYVPFTEQTAVEVVL    345
```

```
tr|A0A0M9FWU4|A0A0M9FWU4_LEPPY      GRTVGKSRA---------DIEAMVQHDFASLYGTLLP--AYTRTFVFFPFTTQVAEEVVR    384
tr|A4H5R3|A4H5R3_LEIBR              GRTVGKSRA---------EIEAMIQHDFADLYGSLLP--AYTRTFVYFPFTQAMAEDVVR    430
tr|Q4QHU2|Q4QHU2_LEIMA              GRTVGKSRT---------EIEAMIQHDFADLYGTLLP--AYTRTFVYYPFTSAMAEDVVR    401
tr|A4HU03|A4HU03_LEIIN              GRTVGKSRT---------EIEAMIQHDFADLYGTLLP--AYTRTFVYYPFTSAMAEDVVR    401
tr|Q382P2|Q382P2_TRYB2              GRTFGKSVV---------EVEKMVRMEFNSLYGSFVP--AFVRTLAFAAFSKRSAEEMVR    323
tr|A0A1X0NMJ2|A0A1X0NMJ2_9TRYP      GRTWGKSVV---------EIEQMVRVDFAGLYGTLVP--AFARTMVFVSLSQQSAEEMVR    308
tr|Q4D6G4|Q4D6G4_TRYCC              GRTWGKSMA---------EIEQLVRDEFAGLYGTLLS--AFTRTMLFLPFSRHDAEEMIR    307
tr|Q4D903|Q4D903_TRYCC              GRTWGKSMA---------EIEQLVRDEFAGLYGTLLS--AFTRTMLFLPFSRHDAEEMIR    307
tr|A0A422ND04|A0A422ND04_9TRYP      GRTWGKSTA---------EVEQLVRDEFAGLYGTLTS--AFTRTMLFAAFSRRDAEKLVQ    307
tr|A0A3R7RCD3|A0A3R7RCD3_TRYRA      GRTWGKSMA---------EVEQLVRDEFSGLYGTLVS--AFTRTMLFTAFSRLDAEQMVR    307
                                                      :                          :

tr|D7G3V5|D7G3V5_ECTSI              LKLKKLAETLDGG--------------------LYTTS------GLRWHLVQ--------    322
tr|Q55FE2|Q55FE2_DICDI              KEIENSFCRENHL----------------------SILSIELKSQVSEIIFQKMNTLYPN    371
tr|L8GYA8|L8GYA8_ACACA              --VPDLNCTARVG-----------------------KVSAPVEELVSLLVRQLKEHPAAK    296
tr|A0A0L1L0A3|A0A0L1L0A3_9EUGL      YRLNTLKCQHNKKNS-LFGNTNI----------NILEFTFDEEVLVDFLYCKILSGI-PQ    355
tr|A0A0S4KNW8|A0A0S4KNW8_BODSA      LIFNDWACSEK-----------------------MNSLSVT-PEAIDVVVDGCVGRV-AF    317
tr|A0A422N6P1|A0A422N6P1_TRYRA      LTIQDYKCRHG---------------------ETIRELRVS-EDAVLWFVELVHDDL-PM    320
tr|A0A3R7NQX6|A0A3R7NQX6_9TRYP      LTILDYKCRHG---------------------EAIRELRVS-EDAVLWFVDLVRDDL-PV    323
tr|S9U3X1|S9U3X1_9TRYP              MGVADSRCAWS-----------------------M-NVTIS-DDAVAWMLEGAKPYL-AS    279
tr|S9U3J6|S9U3J6_9TRYP              MGVADSRCAWS-----------------------M-NVTIS-DDAVAWMLEGAKPYL-AS    321
tr|A0A0N0VF20|A0A0N0VF20_LEPPY      VVAKEIGCSAT---------------------PPV-RLAIT-DTAVLWLVEKTKGSL-AV    317
tr|A4I131|A4I131_LEIIN              VVVREIGCSAP---------------------QPL-CLAIS-DSAVVWLVERTKILL-PA    321
tr|Q4QAA8|Q4QAA8_LEIMA              VVVREIGCSAQ---------------------QPL-CLAIS-DSAVVWLVERTKMLL-SA    321
tr|A0A7G2CLT7|A0A7G2CLT7_9TRYP      HKISQLSCVPGHFLSSWFHLEEDEALRSILARSRVTASWME-EDAAKFLIDLHRPLWEAS    391
tr|S9VXS8|S9VXS8_9TRYP              ATIDYLPCQLLLA-----S----------GGAAALRASEID-ADAAAFLVHKYREVWQGR    389
tr|A0A0M9FWU4|A0A0M9FWU4_LEPPY      SVVTDLPCALG--------------------EHLIAESWIS-DDAVAFLVEQHRAAWAGK    423
tr|A4H5R3|A4H5R3_LEIBR              SAVTDLPCALG--------------------ERLIASSTIS-DEAVAFLVQQHRQLWSGK    469
tr|Q4QHU2|Q4QHU2_LEIMA              SAVTDLPCALG--------------------ERLIASSTIS-DEAVSFLVQQHRQLWSGK    440
tr|A4HU03|A4HU03_LEIIN              SAVTDLPCALG--------------------ERLIASSTIS-DEAVSFLVQQHRQLWSGK    440
tr|Q382P2|Q382P2_TRYB2              TTVITLPCTAYRY-----GFAGPN-----AYGGGVVASSID-DVAVSFLVERYREVWEGR    372
tr|A0A1X0NMJ2|A0A1X0NMJ2_9TRYP      KAAAALACLNNWG-----------------G-GTVMASTIE-ELAVAYLVERYRDTWEGR    349
tr|Q4D6G4|Q4D6G4_TRYCC              IAVAALPCGTYWN-----------------K-GAVVTSSIE-DLAVTFLVERHRETWEGK    348
tr|Q4D903|Q4D903_TRYCC              IAVAALPCGTYWN-----------------K-GAVVTSSIE-DLAVTFLVERHRETWEGR    348
tr|A0A422ND04|A0A422ND04_9TRYP      MAVAALPCTRYWS-----------------DGGAVVASSID-ELAVAFLVERHRETWEGK    349
tr|A0A3R7RCD3|A0A3R7RCD3_TRYRA      IAVAALPCAKYWS-----------------DSGSVVASSID-DLAVAFLVERHRETWEGK    349


tr|D7G3V5|D7G3V5_ECTSI              ------PQYIQYSSYHVTLPNDGREEIIRHQL------AAYGARDVEKVSMRRLTGAIRN    370
tr|Q55FE2|Q55FE2_DICDI              ----------------------------------------ENFRAIEKILNYLIFNKVSN    391
tr|L8GYA8|L8GYA8_ACACA              RDNIVIKKYLPFFATTVGADIHAIAEKLIHFMVAHEHYCERNYRGIEGLFATKVVGPI--    354
tr|A0A0L1L0A3|A0A0L1L0A3_9EUGL      ----------------------------------------RNGREIDRIFDDYIEGPL--    373
tr|A0A0S4KNW8|A0A0S4KNW8_BODSA      ----------------------------------------ENGRAVVLHMDVELVRL---    334
tr|A0A422N6P1|A0A422N6P1_TRYRA      ----------------------------------------ENGRCVAHAVSALVGPGI--    338
tr|A0A3R7NQX6|A0A3R7NQX6_9TRYP      ----------------------------------------ENGRCVAQAVSALVGPGI--    341
tr|S9U3X1|S9U3X1_9TRYP              ----------------------------------------ENGRAVIKEVLAAVEPLL--    297
tr|S9U3J6|S9U3J6_9TRYP              ----------------------------------------ENGRAVIKEVLAAVEPLL--    339
tr|A0A0N0VF20|A0A0N0VF20_LEPPY      ----------------------------------------ENGRAVAQETKLHVGSLM--    335
tr|A4I131|A4I131_LEIIN              ----------------------------------------ENGRAVAFETKLQLEALL--    339
tr|Q4QAA8|Q4QAA8_LEIMA              ----------------------------------------ENGRAVAFETKLQLEALL--    339
tr|A0A7G2CLT7|A0A7G2CLT7_9TRYP      ----------------------------------------ENGRSIEKNIIQ--------    403
tr|S9VXS8|S9VXS8_9TRYP              ----------------------------------------ENGHALRRALEDEVLTPL--    407
tr|A0A0M9FWU4|A0A0M9FWU4_LEPPY      ----------------------------------------ENGHALRRLVEDELVSQL--    441
tr|A4H5R3|A4H5R3_LEIBR              ----------------------------------------ENGHALRRLIEDEIVSQL--    487
tr|Q4QHU2|Q4QHU2_LEIMA              ----------------------------------------ENGHALRRLVEDEIVSQL--    458
tr|A4HU03|A4HU03_LEIIN              ----------------------------------------ENGHALRRLVEDEIVSQL--    458
tr|Q382P2|Q382P2_TRYB2              ----------------------------------------ENGHALRRAVEDSLLSLL--    390
tr|A0A1X0NMJ2|A0A1X0NMJ2_9TRYP      ----------------------------------------ENGHALRRVVQDTLVPLL--    367
tr|Q4D6G4|Q4D6G4_TRYCC              ----------------------------------------ENGHSLRRAVEDSLVSLL--    366
tr|Q4D903|Q4D903_TRYCC              ----------------------------------------ENGHSLRRAVEDSLVSLL--    366
tr|A0A422ND04|A0A422ND04_9TRYP      ----------------------------------------ENGHALRRAVEDSLISLL--    367
tr|A0A3R7RCD3|A0A3R7RCD3_TRYRA      ----------------------------------------ENGHALRRAVEDSLISLL--    367
                                                                                 . : :

tr|D7G3V5|D7G3V5_ECTSI              HVLNPTCAATDEGAEETGSPEGQGRGGGGGRPAQGGYGGSSSSSRGGGGGRCWS------    424
tr|Q55FE2|Q55FE2_DICDI              FII----QDYDKLI-----EET-KINNNNND-------------N--ENEKIQQQNENYK    426
tr|L8GYA8|L8GYA8_ACACA              --L----KQMPEPR--PSTQ-----------------------------------------    366
tr|A0A0L1L0A3|A0A0L1L0A3_9EUGL      --V----MKLAEFE--MCASEK-STSTGGSR-----------------ESNLWE------    401
tr|A0A0S4KNW8|A0A0S4KNW8_BODSA      -------QHPAGRS-----LQG-L--N-----------------T---TVTARN------    353
tr|A0A422N6P1|A0A422N6P1_TRYRA      --L----RHLLDSP-----LAP-V-------------------------SLHVDI-----    356
tr|A0A3R7NQX6|A0A3R7NQX6_9TRYP      --L----RHLPDNP-----SAL-V-------------------------SLYVDI-----    359
tr|S9U3X1|S9U3X1_9TRYP              --E----QGDKVR--------P-YIID---------------------------------    309
tr|S9U3J6|S9U3J6_9TRYP              --E----QGDKVR--------P-YIID---------------------------------    351
tr|A0A0N0VF20|A0A0N0VF20_LEPPY      --E----RLQADEA-----SHP-S--S-----------------K---TLRPRDGAAN--    359
tr|A4I131|A4I131_LEIIN              --E----QVMDNCT-----HES-GIIA-----------------T---DELYLDVDAT--    365
tr|Q4QAA8|Q4QAA8_LEIMA              --E----QVMGNYT-----YES-GIIA-----------------T---DELHLDVDGT--    365
tr|A0A7G2CLT7|A0A7G2CLT7_9TRYP      --L----ELWNTLR--TG-LKT-LYHHLGLS-------------Q------RFEVFRQ--    432
tr|S9VXS8|S9VXS8_9TRYP              --L----LAMEELQ-----QQP-ARHGRGQR-------------RAAAASTLWDPAAA--    440
tr|A0A0M9FWU4|A0A0M9FWU4_LEPPY      --I----VHWELHA-----VR----------------------------ERLL-------    455
tr|A4H5R3|A4H5R3_LEIBR              --V----AYWETHE-----EQK-AF------------------------ERLR-------    504
tr|Q4QHU2|Q4QHU2_LEIMA              --I----VYWETHA-----QQR-HF------------------------ERLR-------    475
tr|A4HU03|A4HU03_LEIIN              --V----VYWETHA-----QQK-HF------------------------ERLR-------    475
tr|Q382P2|Q382P2_TRYB2              --L----KYFDEHG-----HD----------------------------RRVW-------    404
tr|A0A1X0NMJ2|A0A1X0NMJ2_9TRYP      --L----QYFDREG-----HD----------------------------QAVW-------    381
tr|Q4D6G4|Q4D6G4_TRYCC              --L----QYFDQNG-----HD----------------------------KFVW-------    380
tr|Q4D903|Q4D903_TRYCC              --L----QYFDQHG-----HD----------------------------KFVW-------    380
tr|A0A422ND04|A0A422ND04_9TRYP      --L----QYFDQHG-----HD----------------------------TVVW-------    381
tr|A0A3R7RCD3|A0A3R7RCD3_TRYRA      --L----KYFDKHG-----QD----------------------------TVVW-------    381
```

```
tr|D7G3V5|D7G3V5_ECTSI            EPFM--VDIRYDAS----------TEQVSFHRCEPDYADGL-------------------    453
tr|Q55FE2|Q55FE2_DICDI            IPFNYVYDSIFKIIKNNKILENKNNHNDEVLITLSIKPNNL-------------------    467
tr|L8GYA8|L8GYA8_ACACA            --------------------------EVTVLVKVPFDQS--------VVQPARAPGGGR-    391
tr|A0A0L1L0A3|A0A0L1L0A3_9EUGL    ---------YLFLG----------SSSIDVSVSFLVV----------------------N    420
tr|A0A0S4KNW8|A0A0S4KNW8_BODSA    ---G---AVEISID----------RDADDAMSDL--------------------------    371
tr|A0A422N6P1|A0A422N6P1_TRYRA    -------------D----------DNGSVTVLPLPAG-----------------------    370
tr|A0A3R7NQX6|A0A3R7NQX6_9TRYP    -------------D----------DNGSVAVLPLPAR-----------------------    373
tr|S9U3X1|S9U3X1_9TRYP            ------------VD----------AHGHLALFSEEYTV----------------------    325
tr|S9U3J6|S9U3J6_9TRYP            ------------VD----------AHGHLALFSEEYTV----------------------    367
tr|A0A0N0VF20|A0A0N0VF20_LEPPY    CSVP---ACNIYVD----------DGGQIGLAC---------------------------    379
tr|A4I131|A4I131_LEIIN            CPYR---RCTVFLE----------GDGTLAMTCQGTGTHSR------------VPPG---    397
tr|Q4QAA8|Q4QAA8_LEIMA            CPYR---RCTVFLE----------GDGTLAMTCQGTGTHTR------------VSPD---    397
tr|A0A7G2CLT7|A0A7G2CLT7_9TRYP    ---L---EIVFYLR----------EETATVAVRING--------------IA------VD    456
tr|S9VXS8|S9VXS8_9TRYP            ---T---DVRFFLD----------AAQWRIHFSLLQY--------GEVVYTAGEPSGAWG    476
tr|A0A0M9FWU4|A0A0M9FWU4_LEPPY    --------VRFELD----------EAELCVVLRLPNQHATAV---F-------DVDAVMP    487
tr|A4H5R3|A4H5R3_LEIBR            --------VRFELD----------EASMRVLLRTPHASTTTA---L-------PPSSALS    536
tr|Q4QHU2|Q4QHU2_LEIMA            --------VHFDLD----------EASMHVLLRTPHTSSTIA---S-------QPPAALS    507
tr|A4HU03|A4HU03_LEIIN            --------VHFELD----------EASMRVLLRTPHTSRTTA---S-------QPPAALS    507
tr|Q382P2|Q382P2_TRYB2            --------ARFHLD----------EKVGEIVLDAGADPHSMNDL----------------    430
tr|A0A1X0NMJ2|A0A1X0NMJ2_9TRYP    --------ARFRLD----------TTAGKIILDTGRDENGLDPSYVDSVEAGTGTSGALG    423
tr|Q4D6G4|Q4D6G4_TRYCC            --------AHFRLD----------ERAAKIVLSTGWKYDHMETSSEEFFGVATEAERTVL    422
tr|Q4D903|Q4D903_TRYCC            --------AHFRLD----------ERAAKIVLSTGWNYDHMETSSEEFFGVAIEAEGTVL    422
tr|A0A422ND04|A0A422ND04_9TRYP    --------AYFRLD----------ELTAKIVLSTGAERYQTDTRSIGYSGVAAGPGGAAA    423
tr|A0A3R7RCD3|A0A3R7RCD3_TRYRA    --------AYFRLD----------ELTAKIVLSTGSEYNQADVRSEESSGITTGGGGAAT    423


tr|D7G3V5|D7G3V5_ECTSI            -TEL-------------------QREHPDVVEGPKCDLAWRGVLHEHGALA---------    484
tr|Q55FE2|Q55FE2_DICDI            -----------------------------------------------------NNLLIID    474
tr|L8GYA8|L8GYA8_ACACA            ------------------EDL---------------------------------------    394
tr|A0A0L1L0A3|A0A0L1L0A3_9EUGL    STE-----TQGLDKKLVIEVLLDQRKH---------------------------------    442
tr|A0A0S4KNW8|A0A0S4KNW8_BODSA    ------------------------------------------------------------    371
tr|A0A422N6P1|A0A422N6P1_TRYRA    ------------------------------------------------------------    370
tr|A0A3R7NQX6|A0A3R7NQX6_9TRYP    ------------------------------------------------------------    373
tr|S9U3X1|S9U3X1_9TRYP            ------------------------------------------------------------    325
tr|S9U3J6|S9U3J6_9TRYP            ------------------------------------------------------------    367
tr|A0A0N0VF20|A0A0N0VF20_LEPPY    ------------------------------------------------------------    379
tr|A4I131|A4I131_LEIIN            ------------------------------------------------------------    397
tr|Q4QAA8|Q4QAA8_LEIMA            ------------------------------------------------------------    397
tr|A0A7G2CLT7|A0A7G2CLT7_9TRYP    EIQAQAEAKQA--EEG--D-FVGK-KKVFLIIK-KISRKWDCYIVDFETKNSIFFLLLFF    509
tr|S9VXS8|S9VXS8_9TRYP            EVEG------------------------------DL------------------------    482
tr|A0A0M9FWU4|A0A0M9FWU4_LEPPY    APEDDAASDEG---------------T----AG-DL------------------------    503
tr|A4H5R3|A4H5R3_LEIBR            STEARAHVRDGVHNEG--------EEE----PS-DL------------------------    559
tr|Q4QHU2|Q4QHU2_LEIMA            SAEARAYDHGGADDRD--------GEE----RG-DL------------------------    530
tr|A4HU03|A4HU03_LEIIN            SVEARAYGHGGADDGD--------GEE----RS-DL------------------------    530
tr|Q382P2|Q382P2_TRYB2            ------------------------------------------------------------    430
tr|A0A1X0NMJ2|A0A1X0NMJ2_9TRYP    DSVG-------K---------KRRDEERHVIND-DF------------------------    442
tr|Q4D6G4|Q4D6G4_TRYCC            ---------SGEEKDGGGENVAERGKHASTLNE-DL------------------------    448
tr|Q4D903|Q4D903_TRYCC            ---------SGEDKEGGGEDVAEKGKHASTFNG-DL------------------------    448
tr|A0A422ND04|A0A422ND04_9TRYP    AAAAAAATTADRPGEGKAEEFAERGIGPYTLNG-DL------------------------    458
tr|A0A3R7RCD3|A0A3R7RCD3_TRYRA    -----------RPGE-EVDEVAARGKGPYTLNG-DL------------------------    446


tr|D7G3V5|D7G3V5_ECTSI            ------------------------    484
tr|Q55FE2|Q55FE2_DICDI            IF------------PSKNVYKNTN    486
tr|L8GYA8|L8GYA8_ACACA            ------------------------    394
tr|A0A0L1L0A3|A0A0L1L0A3_9EUGL    ------------------------    442
tr|A0A0S4KNW8|A0A0S4KNW8_BODSA    ------------------------    371
tr|A0A422N6P1|A0A422N6P1_TRYRA    ------------------------    370
tr|A0A3R7NQX6|A0A3R7NQX6_9TRYP    ------------------------    373
tr|S9U3X1|S9U3X1_9TRYP            ------------------------    325
tr|S9U3J6|S9U3J6_9TRYP            ------------------------    367
tr|A0A0N0VF20|A0A0N0VF20_LEPPY    ------------------------    379
tr|A4I131|A4I131_LEIIN            ------------------------    397
tr|Q4QAA8|Q4QAA8_LEIMA            ------------------------    397
tr|A0A7G2CLT7|A0A7G2CLT7_9TRYP    LFHKKIIKLKKKKWPPRKVCSS--    531
tr|S9VXS8|S9VXS8_9TRYP            ------------------------    482
tr|A0A0M9FWU4|A0A0M9FWU4_LEPPY    ------------------------    503
tr|A4H5R3|A4H5R3_LEIBR            ------------------------    559
tr|Q4QHU2|Q4QHU2_LEIMA            ------------------------    530
tr|A4HU03|A4HU03_LEIIN            ------------------------    530
tr|Q382P2|Q382P2_TRYB2            ------------------------    430
tr|A0A1X0NMJ2|A0A1X0NMJ2_9TRYP    ------------------------    442
tr|Q4D6G4|Q4D6G4_TRYCC            ------------------------    448
tr|Q4D903|Q4D903_TRYCC            ------------------------    448
tr|A0A422ND04|A0A422ND04_9TRYP    ------------------------    458
tr|A0A3R7RCD3|A0A3R7RCD3_TRYRA    ------------------------    446
```

### 09. BIP-like HSP70 chaperones

BIP-like chaperones (Binding immunoglobulin protein, BIP, also known as 78 kDa glucose-regulated proteins, GRP78) are core components of the eukaryotic folding machinery associated with translocons. Hence the kinetoplastid proteins align well with BIPs from all other major groups, including fungal, plant and animal orthologues. The KDEL-like retention signal is also extremely conserved. Although sharing a HSP70-type core, BIP proteins differ greatly from the other key ER-localized HSP70 group (HYOU1), and should be handled as an ancestrally separate family.

```
tr|A0A0L1KCY6|A0A0L1KCY6_9EUGL      ------------------------------------------------------------    0
tr|A0A0S4JK39|A0A0S4JK39_BODSA      ------------------------------------------------------MF----    2
tr|A0A7G2CL24|A0A7G2CL24_9TRYP      ------------------------------------------------------------    0
tr|A0A0S4IPK6|A0A0S4IPK6_BODSA      ------------------------------------------------------------    0
tr|A0A0N0DZK2|A0A0N0DZK2_LEPPY      ------------------------------------------------------------    0
tr|A4I3J9|A4I3J9_LEIIN              ------------------------------------------------------------    0
tr|Q4Q8E6|Q4Q8E6_LEIMA              ------------------------------------------------------------    0
tr|A4HGG7|A4HGG7_LEIBR              ------------------------------------------------------------    0
tr|S9U467|S9U467_9TRYP              MVLCFLGLLLFEYTVVLLHSLFLVCTFCTVYPLVLIANCDCEYCILSLQYRKKNFFSSTK    60
tr|S9W3A0|S9W3A0_9TRYP              ------------------------------------------------------------    0
tr|A0A1X0NYK3|A0A1X0NYK3_9TRYP      ------------------------------------------------------------    0
tr|Q4D620|Q4D620_TRYCC              ------------------------------------------------------------    0
tr|K2NLB6|K2NLB6_TRYCR              ------------------------------------------------------------    0
tr|A0A422P6Q7|A0A422P6Q7_9TRYP      ------------------------------------------------------------    0
tr|A0A422N1T5|A0A422N1T5_TRYRA      ------------------------------------------------------------    0
tr|F9WRV9|F9WRV9_TRYVY              ------------------------------------------------------------    0
tr|F9W3P7|F9W3P7_TRYCI              ------------------------------------------------------------    0
tr|Q384Q5|Q384Q5_TRYB2              ------------------------------------------------------------    0
tr|A0A1Y2AI48|A0A1Y2AI48_9FUNG      ------------------------------------------------------------    0
tr|A0A1Y1X8R1|A0A1Y1X8R1_9FUNG      ------------------------------------------------------------    0
sp|P11021|BIP_HUMAN                 ------------------------------------------------------------    0
tr|A0A0G4EPG2|A0A0G4EPG2_VITBC      ------------------------------------------------------------    0
tr|M1BLB0|M1BLB0_SOLTU              ------------------------------------------------------------    0
sp|Q9LKR3|BIP1_ARATH                ------------------------------------------------------------    0
tr|I1KPN3|I1KPN3_SOYBN              ------------------------------------------------------------    0


tr|A0A0L1KCY6|A0A0L1KCY6_9EUGL      -------------------MLQ--T--TAILFAACFLCSAKEEKKQLEFPVVGIDLGTTY    37
tr|A0A0S4JK39|A0A0S4JK39_BODSA      -----------RNVRPLVALLIVTAVLLLLAGAAHAGAANTAASNKVEPPCVGIDLGTTY    51
tr|A0A7G2CL24|A0A7G2CL24_9TRYP      ----------MNLFRKRS-LI---ATTLLLLSASVLVLAAPSGTGKVAGPCVGVDLGTTY    46
tr|A0A0S4IPK6|A0A0S4IPK6_BODSA      ----------MQVTRGL---V---AAVLVAAVVVGTVLAGEASTNKVEAPCVGIDLGTTY    44
tr|A0A0N0DZK2|A0A0N0DZK2_LEPPY      ----------MAVRDRLL-LL---AVCLVAALLIVSAAAAPDGSGKVEPPCIGVDLGTTY    46
tr|A4I3J9|A4I3J9_LEIIN              ----------MTRKDNLT-LM---AVCLVSAMLVMSAAAAADGSGKVESPCIGVDLGTTY    46
tr|Q4Q8E6|Q4Q8E6_LEIMA              ----------MTRKDNLT-LM---AVCLVSAILVVSAAAVPDGSGKVESPCIGVDLGTTY    46
tr|A4HGG7|A4HGG7_LEIBR              ----------MMRKDSLT-LV---GVCLVSVMLVLSAAAAPDGSGKVEPPCIGVDLGTTY    46
tr|S9U467|S9U467_9TRYP              RIDTYTSSLKMPSLKELV-LL---LT-VLLVSVALLVSAAPEGAGKVEAPCVGVDLGTTY    115
tr|S9W3A0|S9W3A0_9TRYP              ----------MPSLKELV-LL---LT-VLLVSVALLVSAAPEGAGKVEAPCVGVDLGTTY    45
tr|A0A1X0NYK3|A0A1X0NYK3_9TRYP      -----------MTRFPL--VV---VL-GIVVVLAATVYAAPDGTGKVEAPCVGIDLGTTY    43
tr|Q4D620|Q4D620_TRYCC              --------------MLL--QA---LL---VLSAVVVAVAAPDGTGKVEAPCVGIDLGTTY    38
tr|K2NLB6|K2NLB6_TRYCR              --------------MFL--QV---LL---VLSTVVAAVAAPDGTGKVEAPCVGIDLGTTY    38
tr|A0A422P6Q7|A0A422P6Q7_9TRYP      -----------MPRLLL--AA---LL---VLSA-VAAAAAPDGTGKVEAPCVGIDLGTTY    40
tr|A0A422N1T5|A0A422N1T5_TRYRA      -----------MARLLL--AA---LL---VLFT-VVAAAAPDGTGKVEAPCVGIDLGTTY    40
tr|F9WRV9|F9WRV9_TRYVY              ----------MAKAMRL--AA---AA-LLLVAAATGAWAAPEASGKVEAPCVGIDLGTTY    44
tr|F9W3P7|F9W3P7_TRYCI              -----------MSGTSL--RT---AA-VLLVVAAAVATAAPESGGKVEAPCVGIDLGTTY    43
tr|Q384Q5|Q384Q5_TRYB2              -----------MSRMWL--TT---AA-VFLTVTVAAVSAAPESGGKVEAPCVGIDLGTTY    43
tr|A0A1Y2AI48|A0A1Y2AI48_9FUNG      ----------MK-------LSS--IFSTVVVALAAIVRAADSDDNGIKGPVIGIDLGTTY    41
tr|A0A1Y1X8R1|A0A1Y1X8R1_9FUNG      ----------MRFSIK--KLLI--AAASVVAFSAPKVFAASDDDGKVKGPIIGIDLGTTY    46
sp|P11021|BIP_HUMAN                 -----------------MKLSL--VAAMLL--LLSAARAEEEDKKEDVGTVVGIDLGTTY    39
tr|A0A0G4EPG2|A0A0G4EPG2_VITBC      ------------------MRSL--IVGAVLVAAIATSALAAEEEKKIEGPVVGIDLGTTY    40
tr|M1BLB0|M1BLB0_SOLTU              ----------MATALKRRASSI--VFAIVLLGSLFA-FSIAKEEATKLGTVIGIDLGTTY    47
sp|Q9LKR3|BIP1_ARATH                ----------MARS-FGANSTV--VLAIIFFGCLFA-LSSAIEEATKLGSVIGIDLGTTY    46
tr|I1KPN3|I1KPN3_SOYBN              ----------MAGS-WARRSLI--VLAIISFGCLFA-ISIAKEEATKLGTVIGIDLGTTY    46
                                                                                     :*:******

tr|A0A0L1KCY6|A0A0L1KCY6_9EUGL      SVVGVMKNGEVQIIPNEMGNRITPSVVAFT-ESERLIGDGAKNQLPQNPMNTIYTIKRLI    96
tr|A0A0S4JK39|A0A0S4JK39_BODSA      SVVGVWQNGEVQIIPNELGNRITPSVVAFT-ESERLVGDGAKNQLPQNPENTIYAIKRLI    110
tr|A0A7G2CL24|A0A7G2CL24_9TRYP      SVVGVWQKGEVHIIPNEMGNRITPSVVAFT-DDERLIGDGAKNQLPQNPENTVYAIKRLI    105
tr|A0A0S4IPK6|A0A0S4IPK6_BODSA      SVVGVWQKGEGQIIPNELGNRITPSVVAFT-ETERLIGDGAKNQLPQNPYNTIYTIKRLI    103
tr|A0A0N0DZK2|A0A0N0DZK2_LEPPY      SVAGVWQKGEVHIITNEMGNRITPSVVAFT-ETERLVGDGAKNQLPQNPENTIYAIKRLI    105
tr|A4I3J9|A4I3J9_LEIIN              SVAGVWQKGEVHIVTNEMGNRITPSVVAFT-DAERLVGDGAKNQLPQNPENTIYAIKRLI    105
tr|Q4Q8E6|Q4Q8E6_LEIMA              SVAGVWQKGEVHIVTNEMGNRITPSVVAFT-DAERLVGDGAKNQLPQNPENTIYAIKRLI    105
tr|A4HGG7|A4HGG7_LEIBR              SVAGVWQKGEVHVVTNEMGNRITPSVVAFT-DTERLVGDGAKNQLPQNPENTIYAIKRLI    105
tr|S9U467|S9U467_9TRYP              SVVGVWQKGDVHIVPNEMGNRITPSVVAFT-ETERLIGDGAKNQLPQNPENTIYAIKRLI    174
tr|S9W3A0|S9W3A0_9TRYP              SVVGVWQKGDVHIVPNEMGNRITPSVVAFT-ETERLIGDGAKNQLPQNPENTIYAIKRLI    104
tr|A0A1X0NYK3|A0A1X0NYK3_9TRYP      SVVGVWQKGDVHIIPNEMGNRITPSVVAFT-ETERLIGDGAKNQLPQNPHNTIYAIKRLI    102
tr|Q4D620|Q4D620_TRYCC              SVVGVWQKGDVHIIPNDMGNRITPSVVAFT-ETERLIGDGAKNQLPQNPHNTIYAIKRLI    97
tr|K2NLB6|K2NLB6_TRYCR              SVVGVWQKGDVHIIPNDMGNRITPSVVAFT-ETERLIGDGAKNQLPQNPHNTIYAIKRLI    97
tr|A0A422P6Q7|A0A422P6Q7_9TRYP      SVVGVWQKGDVHIIPNEMGNRITPSVVAFT-DTERLIGDGAKNQLPQNPHNTIYAIKRLI    99
tr|A0A422N1T5|A0A422N1T5_TRYRA      SVVGVWQKGDVHIIPNEMGNRITPSVVAFT-DTERLIGDGAKNQLPQNPHNTIYAIKRLI    99
tr|F9WRV9|F9WRV9_TRYVY              SVVGVWQKGDVHIIPNEMGNRITPSVVAFT-ETERLIGDGAKNQLPQNPHNTIYTIKRLI    103
tr|F9W3P7|F9W3P7_TRYCI              SVVGVWQKGDVHIIPNEMGNRITPSVVAFT-DTERLIGDGAKNQLPQNPHNTIYTIKRLI    102
tr|Q384Q5|Q384Q5_TRYB2              SVVGVWQKGDVHIIPNEMGNRITPSVVAFT-DTERLIGDGAKNQLPQNPHNTIYTIKRLI    102
tr|A0A1Y2AI48|A0A1Y2AI48_9FUNG      SCVGVYKNGRVEIIANDQGHRITPSYVAFT-DEERLVGDAAKNQAPMNPTNTIFDAKRLV    100
tr|A0A1Y1X8R1|A0A1Y1X8R1_9FUNG      SCVGIYQNGRVEILANDQGHRITPSYVAFT-EEERLVGDAAKNQAPSNPKNTIFDIKRLI    105
sp|P11021|BIP_HUMAN                 SCVGVFKNGRVEIIANDQGNRITPSYVAFTPEGERLIGDAAKNQLTSNPENTVFDAKRLI    99
tr|A0A0G4EPG2|A0A0G4EPG2_VITBC      SCVGIYKNGRVEIIPNDQGNRITPSYVAFT-EDERLIGEAAKNQATVNPSQTLFDVKRLI    99
tr|M1BLB0|M1BLB0_SOLTU              SCVGVYKNGHVEIIANDQGNRITPSWVAFT-DGERLIGEAAKNQAAVNPERTVFDVKRLI    106
```

```
sp|Q9LKR3|BIP1_ARATH            SCVGVYKNGHVEIIANDQGNRITPSWVGFT-DSERLIGEAAKNQAAVNPERTVFDVKRLI    105
tr|I1KPN3|I1KPN3_SOYBN          SCVGVYKNGHVEIIANDQGNRITPSWVAFT-DSERLIGEAAKNQAAVNPERTIFDVKRLI    105
                                * .*: ::*  .:: *: *:***** *.** : ***:*:.****   ** .*::  ***:

tr|A0A0L1KCY6|A0A0L1KCY6_9EUGL  GRKYTDKTVQYDKKLLSFDIVSH-NNEPRVQVNYK-GEKKVFTPEEISAMILTKMKEIAE    154
tr|A0A0S4JK39|A0A0S4JK39_BODSA  GRRYSDATVQKDKKLLSYELVEDKDGKPRVRVAVD-GKARLFTPEEVSAMVLQKMKEIAE    169
tr|A0A7G2CL24|A0A7G2CL24_9TRYP  GRKYTDATVQADKKLLSYEVIADKGGKPIVQVKAG-GEKKTYSPEEISAMVLQKMKQIAE    164
tr|A0A0S4IPK6|A0A0S4IPK6_BODSA  GRKYRDETVQRDKKLLSYELVEDKDGKPRVQVEVD-GKKKQFTPEEVSAMVLQKMKEIAE    162
tr|A0A0N0DZK2|A0A0N0DZK2_LEPPY  GRKFSDATVQNDKKLLSYKIVSDKAGKPLVQVTVS-GAKKEFTPEEVSAMVLQKMKDISE    164
tr|A4I3J9|A4I3J9_LEIIN          GRKYADPTVQNDKKLLSYHIVADKTGKPLVQVTVK-GQQKRFTPEEVSAMVLQKMKEISE    164
tr|Q4Q8E6|Q4Q8E6_LEIMA          GRKYADPTVQNDKKLLSYHIVADKTGKPLVQVTVK-GQQKRFTPEEVSAMVLQKMKEISE    164
tr|A4HGG7|A4HGG7_LEIBR          GRKYSDPTVQNDKKLLSYQIVADKVGKPLVQVTVK-GQQKRFTPEEVSAMVLQKMKEISE    164
tr|S9U467|S9U467_9TRYP          GRKYSDVTVQADKKLLSYEVVADKTGKPLVRVTVK-GAKKDFTAEEISSMVLQKMKEIAE    233
tr|S9W3A0|S9W3A0_9TRYP          GRKYSDVTVQADKKLLSYEVVADKTGKPLVRVTVK-GAKKDFTAEEISSMVLQKMKEIAE    163
tr|A0A1X0NYK3|A0A1X0NYK3_9TRYP  GRKYSDATVQKDKKLLSYEVVADKDGKPKVQVEVS-GKKKQFTPEEVSAMVLQKMKEIAE    161
tr|Q4D620|Q4D620_TRYCC          GRKYSDATVQTDKKLLSYEVVADKDGKPKVQVEVG-GKKKQFTPEEVSAMVLQKMKEIAE    156
tr|K2NLB6|K2NLB6_TRYCR          GRKYSDATVQTDKKLLSYEVVADKDGKPKVQVEVG-GKKKQFTPEEVSAMVLQKMKEIAE    156
tr|A0A422P6Q7|A0A422P6Q7_9TRYP  GRKYGDTTVQKDRKLLSYEVVADKDGKPKVQVEVG-GKKKQFTPEEVSAMVLQKMKEIAE    158
tr|A0A422N1T5|A0A422N1T5_TRYRA  GRKYSDSTVQKDKKLLSYEVVADKDGKPKVQVEVS-GKKKQFTPEEVSAMVLQKMKEIAE    158
tr|F9WRV9|F9WRV9_TRYVY          GRKYSDPAVQADKKLLSYEVVADRDGKPKVQVTLG-GKKKEFTPEEVSAMVLQKMKEIAE    162
tr|F9W3P7|F9W3P7_TRYCI          GRKYTDAAVQADKKLLSYEVVADRDGKPKVQVMVG-GKPKQFTPEEISAMVLQKMKEIAE    161
tr|Q384Q5|Q384Q5_TRYB2          GRKYTDAAVQADKKLLSYEVIADRDGKPKVQVMVG-GKKKQFTPEEISAMVLQKMKEIAE    161
tr|A0A1Y2AI48|A0A1Y2AI48_9FUNG  GRRFDEQDVQNDMKHFPFKVVA-KDGKPIIQVKVK-GEDKKFTPEEISAMVLQKMKEIAE    158
tr|A0A1Y1X8R1|A0A1Y1X8R1_9FUNG  GRKYDEPAVQKDLKHFPFNVIK-KDEKPYVKVNVK-GTDKTFSPEEISAMILQKMKNIAE    163
sp|P11021|BIP_HUMAN             GRTWNDPSVQQDIKFLPFKVVE-KKTKPYIQVDIGGGQTKTFAPEEISAMVLTKMKETAE    158
tr|A0A0G4EPG2|A0A0G4EPG2_VITBC  GRKYKDTEVQRDKKLLPFILAE-KEGKPFIEVQVK-GETKSFAPEEISAMVLVKMKEVAE    157
tr|M1BLB0|M1BLB0_SOLTU          GRKFEDKEVQRDMKLVPYKIVN-RDGKPYIQVKIKDGETKVFSPEEVSAMILIKMKETAE    165
sp|Q9LKR3|BIP1_ARATH            GRKFEDKEVQKDRKLVPYQIVN-KDGKPYIQVKIKDGETKVFSPEEISAMILTKMKETAE    164
tr|I1KPN3|I1KPN3_SOYBN          GRKFEDKEVQKDMKLVPYKIVN-KDGKPYIQVKIKDGETKVFSPEEISAMVLIKMKETAE    164
                                ** : :  ** * * . : :       :* :.*   *  : :: **:*:*:* ***: :*

tr|A0A0L1KCY6|A0A0L1KCY6_9EUGL  TYLGTTVKNAVVTVPAYFNDAQRQSTKDAGAIAGLNVARIINEPTAAAIAYGLDKK-GER    213
tr|A0A0S4JK39|A0A0S4JK39_BODSA  TYLGEKVKNAVVTVPAYFNDAQRQSTKDAGTIAGLNVVRIINEPTAAALAYGLNEA-GER    228
tr|A0A7G2CL24|A0A7G2CL24_9TRYP  TYIGEKVKNAVVTVPAYFNDAQRQSTKDAGTIAGLNVVRIINEPTAAALAYGLNKV-GEK    223
tr|A0A0S4IPK6|A0A0S4IPK6_BODSA  TYLGEKVKNAVVTVPAYFNDAQRQSTKDAGTIAGLNVVRIINEPTAAAIAYGLNKA-GEK    221
tr|A0A0N0DZK2|A0A0N0DZK2_LEPPY  TFLGEKVKNAVVTVPAYFNDAQRQATKDAGKIAGLNVVRIINEPTAAAIAYGLNKA-GEK    223
tr|A4I3J9|A4I3J9_LEIIN          TFLGEKVKNAVVTVPAYFNDAQRQATKDSGKIAGLNVVRIINEPTAAAIAYGLNKA-GEK    223
tr|Q4Q8E6|Q4Q8E6_LEIMA          TFLGEKVKNAVVTVPAYFNDAQRQATKDSGKIAGLNVVRIINEPTAAAIAYGLNKA-GEK    223
tr|A4HGG7|A4HGG7_LEIBR          TFLGEKVKNAVVTVPAYFNDAQRQATKDSGKIAGLNVVRIINEPTAAAIAYGLNKA-GEK    223
tr|S9U467|S9U467_9TRYP          TYLGEKVKNAVVTVPAYFNDAQRQSTKDAGTIAGLNVVRIINEPTAAAIAYGLNKA-GEK    292
tr|S9W3A0|S9W3A0_9TRYP          TYLGEKVKNAVVTVPAYFNDAQRQSTKDAGTIAGLNVVRIINEPTAAAIAYGLNKA-GEK    222
tr|A0A1X0NYK3|A0A1X0NYK3_9TRYP  TYLGEKVKNAVVTVPAYFNDAQRQSTKDAGTIAGLNVVRIINEPTAAAIAYGLNKA-GEK    220
tr|Q4D620|Q4D620_TRYCC          TYLGEKVKNAVVTVPAYFNDAQRQSTKDAGTIAGLNVVRIINEPTAAAIAYGLNKA-GEK    215
tr|K2NLB6|K2NLB6_TRYCR          TYLGEKVKNAVVTVPAYFNDAQRQSTKDAGTIAGLNVVRIINEPTAAAIAYGLNKA-GEK    215
tr|A0A422P6Q7|A0A422P6Q7_9TRYP  TYLGEKVKNAVVTVPAYFNDAQRQSTKDAGTIAGLNVVRIINEPTAAAIAYGLNKA-GEK    217
tr|A0A422N1T5|A0A422N1T5_TRYRA  TYLGEKVKNAVVTVPAYFNDAQRQSTKDAGTIAGLNVVRIINEPTAAAIAYGLNKA-GEK    217
tr|F9WRV9|F9WRV9_TRYVY          TYLGEKVKNAVVTVPAYFNDAQRQSTKDAGTIAGLNVVRIINEPTAAAIAYGLNKA-GEK    221
tr|F9W3P7|F9W3P7_TRYCI          TYLGEKVKNAVVTVPAYFNDAQRQSTKDAGTIAGLNVVRIINEPTAAAIAYGLNKA-GEK    220
tr|Q384Q5|Q384Q5_TRYB2          TYLGEKVKNAVVTVPAYFNDAQRQSTKDAGTIAGLNVVRIINEPTAAAIAYGLNKA-GEK    220
tr|A0A1Y2AI48|A0A1Y2AI48_9FUNG  SYLGEKVTHAVVTVPAYFNDAQRQATKDAGTIAGLTVARIINEPTAAAIAYGLDKTGSEK    218
tr|A0A1Y1X8R1|A0A1Y1X8R1_9FUNG  SYLGKSVGHAVVTVPAYFNDAQRQATKDAGTIAGLNILRVINEPTAAAIAYGLDKTGGEK    223
sp|P11021|BIP_HUMAN             AYLGKKVTHAVVTVPAYFNDAQRQATKDAGTIAGLNVMRIINEPTAAAIAYGLDKREGEK    218
tr|A0A0G4EPG2|A0A0G4EPG2_VITBC  AYLGKDVKHAVITVPAYFNDAQRQATKDAGTISGLNVIRIINEPTAAAIAYGMDKK-QEK    216
tr|M1BLB0|M1BLB0_SOLTU          AFLGKTIKDAVVTVPAYFNDAQRQATKDAGVIAGLNVARIINEPTAAAIAYGLDKKGGEK    225
sp|Q9LKR3|BIP1_ARATH            AYLGKKIKDAVVTVPAYFNDAQRQATKDAGVIAGLNVARIINEPTAAAIAYGLDKKGGEK    224
tr|I1KPN3|I1KPN3_SOYBN          AFLGKKINDAVVTVPAYFNDAQRQATKDAGVIAGLNVARIINEPTAAAIAYGLDKKGGEK    224
                                :::*  : .**:************:***:* *:**.: *:********:***:::   *:

tr|A0A0L1KCY6|A0A0L1KCY6_9EUGL  NILVFDLGGGTFDVSLLTIDDGFFEVIATNGDTHLGGEDFDNRMMKHFVDLIKKKSKIDI    273
tr|A0A0S4JK39|A0A0S4JK39_BODSA  NVLVFDLGGGTFDVSLLTIDDGFFEVLAVNGDTHLGGEDFDNNMMRHFVEGLRKKQGIDI    288
tr|A0A7G2CL24|A0A7G2CL24_9TRYP  NILVFDLGGGTFDVSLLTIDGDFFEVVATNGDTHLGGEDFDNNIMRYYVDNLKRKNSVDI    283
tr|A0A0S4IPK6|A0A0S4IPK6_BODSA  NILVFDLGGGTFDVSLLTIDDGFFEVVATNGDTHLGGEDFDNNMMRFYVDNLKKKKNIDI    281
tr|A0A0N0DZK2|A0A0N0DZK2_LEPPY  NILVFDLGGGTFDVSLLTIDEGFFEVVATNGDTHLGGEDFDNNMMKFFVDGLKRKQSIDI    283
tr|A4I3J9|A4I3J9_LEIIN          NILVFDLGGGTFDVSLLTIDEGFFEVVATNGDTHLGGEDFDNNMMKFFVDGLKRKQNVDI    283
tr|Q4Q8E6|Q4Q8E6_LEIMA          NILVFDLGGGTFDVSLLTIDEGFFEVVATNGDTHLGGEDFDNNMMKFFVDGLKRKQNVDI    283
tr|A4HGG7|A4HGG7_LEIBR          NILVFDLGGGTFDVSLLTIDEGFFEVVATNGDTHLGGEDFDNNMMKFFVDGLKRKQNVDI    283
tr|S9U467|S9U467_9TRYP          NILVFDLGGGTFDVSLLTIDEGFFEVVATNGDTHLGGEDFDNLMRYFVDGLKRKKSIDI     352
tr|S9W3A0|S9W3A0_9TRYP          NILVFDLGGGTFDVSLLTIDEGFFEVVATNGDTHLGGEDFDNLMRYFVDGLKRKKSIDI     282
tr|A0A1X0NYK3|A0A1X0NYK3_9TRYP  NILVFDLGGGTFDVSLLTIDEGFFEVVATNGDTHLGGEDFDNNMMRYFVDMLKKKKNVDV    280
tr|Q4D620|Q4D620_TRYCC          NILVFDLGGGTFDVSLLTIDEGFFEVVATNGDTHLGGEDFDNNMMRYFVDMLKKKKNVDV    275
tr|K2NLB6|K2NLB6_TRYCR          NILVFDLGGGTFDVSLLTIDEGFFEVVATNGDTHLGGEDFDNNMMRYFVDMLKKKKNVDV    275
tr|A0A422P6Q7|A0A422P6Q7_9TRYP  NILVFDLGGGTFDVSLLTIDEGFFEVVATNGDTHLGGEDFDNNMMRYFVDALKKKKGVDI    277
tr|A0A422N1T5|A0A422N1T5_TRYRA  NILVFDLGGGTFDVSLLTIDEGFFEVVATNGDTHLGGEDFDNNMMRYFVDMLKKKKGVDI    277
tr|F9WRV9|F9WRV9_TRYVY          NILVFDLGGGTFDVSLLTIDEGFFEVVATNGDTHLGGEDFDNNMMRYFVDMLKKKRNVDV    281
tr|F9W3P7|F9W3P7_TRYCI          NILVFDLGGGTFDVSLLTIDEGFFEVVATNGDTHLGGEDFDNNMMRYFVDMLKKKKNVDI    280
tr|Q384Q5|Q384Q5_TRYB2          NILVFDLGGGTFDVSLLTIDEGFFEVVATNGDTHLGGEDFDNNMMRHFVDMLKKKKNVDI    280
tr|A0A1Y2AI48|A0A1Y2AI48_9FUNG  NILVYDLGGGTFDVSVLTIDDGVFEVLATNGDTHLGGEDFDNRLIEHFTKSYKKKFKKDC    278
tr|A0A1Y1X8R1|A0A1Y1X8R1_9FUNG  NILVYDLGGGTFDVSLLSIDDGVFEVLATSGDTHLGGEDFDNRVIEYLTKQYQRKAKVDV    283
sp|P11021|BIP_HUMAN             NILVFDLGGGTFDVSLLTIDNGVFEVVATNGDTHLGGEDFDQRVMEHFIKLYKKKTGKDV    278
tr|A0A0G4EPG2|A0A0G4EPG2_VITBC  NILVYDLGGGTFDVSLLTIDNGVFEVVATSGDTHLGGEDFDQRVMDHFIKVIKKKYDKDI    276
tr|M1BLB0|M1BLB0_SOLTU          NILVFDLGGGTFDVSILTIDNGVFEVLATNGDTHLGGEDFDQRIMEYFIKLIKKKHGKDI    285
sp|Q9LKR3|BIP1_ARATH            NILVFDLGGGTFDVSVLTIDNGVFEVLSTNGDTHLGGEDFDHRVMEYFIKLIKKKHQKDI    284
tr|I1KPN3|I1KPN3_SOYBN          NILVFDLGGGTFDVSILTIDNGVFEVLATNGDTHLGGEDFDQRIMEYFIKLIKKKHGKDI    284
                                *:**:**********:*:** ..***::..***********:.:: .  .  ::*   *

tr|A0A0L1KCY6|A0A0L1KCY6_9EUGL  SKDQKALARLRKASEAAKRQLSSQPEAHVEVDGLVEGYDFSEKISRAKFEELSMDLFKST    333
tr|A0A0S4JK39|A0A0S4JK39_BODSA  SKDQRALARLRKACEAAKRQLSSQPEARVEVDSLVEGHDFSERVTRSKFEELNMALFKGT    348
tr|A0A7G2CL24|A0A7G2CL24_9TRYP  SKDQKALARLRKACEAAKRQLSSHPEARVEVDSLVEGFDFSEKLTRAKFEELNMDLFKNT    343
```

```
tr|A0A0S4IPK6|A0A0S4IPK6_BODSA      STDQKALARLRKACEAAKRQLSSQPEARVEVDSLVEGYDFSEKMTRSKFEELNMALFKGT    341
tr|A0A0N0DZK2|A0A0N0DZK2_LEPPY      SNDQKALARLRKACEAAKRQLSSHPEARVEVDSLVEGYDFSEKITRAKFEELNMAMFKNT    343
tr|A4I3J9|A4I3J9_LEIIN              SNDQKALARLRKACEAAKRQLSSHPEARVEVDSLVEGYDFSEKITRAKFEELNMALFKNT    343
tr|Q4Q8E6|Q4Q8E6_LEIMA              SNDQKALARLRKACEAAKRQLSSHPEARVEVDSLVEGYDFSEKITRAKFEELNMALFKNT    343
tr|A4HGG7|A4HGG7_LEIBR              SGDQKALARLRKACEAAKRQLSSHPEARVEVDSLVEGHDFSEKITRAKFEELNMALFKNT    343
tr|S9U467|S9U467_9TRYP              SKDLKALARLRKACEAAKRQLSSHPEARVEVDSLTEGFDFSEKITRAKFEELNMEMFKGT    412
tr|S9W3A0|S9W3A0_9TRYP              SKDLKALARLRKACEAAKRQLSSHPEARVEVDSLTEGFDFSEKITRAKFEELNMEMFKGT    342
tr|A0A1X0NYK3|A0A1X0NYK3_9TRYP      SKDQKALARLRKACEAAKRQLSSHPEARVEVDSLTEGFDFSEKITRAKFEELNMELFKGT    340
tr|Q4D620|Q4D620_TRYCC              SKDQKALARLRKACEAAKRQLSSHPEARVEVDSLTEGFDFSEKITRAKFEELNMELFKGT    335
tr|K2NLB6|K2NLB6_TRYCR              SKDQKALARLRKACEAAKRQLSSHPEARVEVDSLTEGFDFSEKITRAKFEELNMELFKGT    335
tr|A0A422P6Q7|A0A422P6Q7_9TRYP      GKDQKALARLRKACEAAKRQLSSHPEARVEVDSLTEGFDFSEKITRAKFEELNMELFKGT    337
tr|A0A422N1T5|A0A422N1T5_TRYRA      GKDQKALARLRKACEAAKRQLSSHPEARVEVDSLTEGFDFSEKITRAKFEELNMELFKGT    337
tr|F9WRV9|F9WRV9_TRYVY              SKDQKALARLRKACEAAKRQLSSHPEARVEVDSLTEGFDFSEKITRAKFEELNMDLFKGT    341
tr|F9W3P7|F9W3P7_TRYCI              SKDQKALARLRKACEAAKRQLSSHPEARVEVDSLTEGFDFSEKITRAKFEELNMDLFKGT    340
tr|Q384Q5|Q384Q5_TRYB2              SKDQKALARLRKACEAAKRQLSSHPEARVEVDSLTEGFDFSEKITRAKFEELNMDLFKGT    340
tr|A0A1Y2AI48|A0A1Y2AI48_9FUNG      TKDLKSMGKLKREVEKAKRALSSQMSVKVEIEAFYDGDDFSETLTRAKFEELNIDLFKKT    338
tr|A0A1Y1X8R1|A0A1Y1X8R1_9FUNG      SKDQRAIGKLKREVEKAKRTLSSQMSVRIEIESFYQGEDFSESLTRAKFEELNMDLFKKT    343
sp|P11021|BIP_HUMAN                 RKDNRAVQKLRREVEKAKRALSSQHQARIEIESFYEGEDFSETLTRAKFEELNMDLFRST    338
tr|A0A0G4EPG2|A0A0G4EPG2_VITBC      SKDKRALQKLRREVEKGKRALSSTHQARIEIESLVEGVDFSETLTRARFEELNNDLFQKT    336
tr|M1BLB0|M1BLB0_SOLTU              SKDNKALGKLRREAERAKRALSSQHQVRVEIESLFDGVDFSEPLTRARFEELNNDLFRKT    345
sp|Q9LKR3|BIP1_ARATH                SKDNKALGKLRRECERAKRALSSQHQVRVEIESLFDGVDFSEPLTRARFEELNNDLFRKT    344
tr|I1KPN3|I1KPN3_SOYBN              SKDNRALGKLRREAERAKRALSSQHQVRVEIESLFDGVDFSEPLTRARFEELNNDLFRKT    344
                                    * ::: :*::  * .** ***  ..::*::.: :* **** ::*::****.  :*: *

tr|A0A0L1KCY6|A0A0L1KCY6_9EUGL      LKPVEAVLADAKLKKSDIEDIVLVGGSTRIPKVQQLLKDFFNGKELNKGINPDEAVAYGA    393
tr|A0A0S4JK39|A0A0S4JK39_BODSA      LEPVRKVLEDAKLKKGDIDEIVLVGGSTRIPKVQQLIRDFFN-KEPNRGVNPDEAVAFGA    407
tr|A0A7G2CL24|A0A7G2CL24_9TRYP      LIPVQKVLEDARMKKTDIDEIVLVGGSTRIPKVQQLIKDFFGGKEPNKGINPDEAVAYGA    403
tr|A0A0S4IPK6|A0A0S4IPK6_BODSA      LEPVRKVLEDSKLKKSEIDEIVLVGGSTRIPKVQQLIRDFFN-KEPNKGINPDEAVAYGA    400
tr|A0A0N0DZK2|A0A0N0DZK2_LEPPY      LIPVQKVLEDAKLKKSDIDEIVLVGGSTRIPKVQQLIKDFFGGKEPNKGINPDEAVAYGA    403
tr|A4I3J9|A4I3J9_LEIIN              LVPVQKVLEDAKLKKSDIDEIVLVGGSTRIPKVQQLIKDFFSGKEPNKGINPDEAVAYGA    403
tr|Q4Q8E6|Q4Q8E6_LEIMA              LVPVQKVLEDAKLKKSDIDEIVLVGGSTRIPKVQQLIKDFFSGKEPNKGINPDEAVAYGA    403
tr|A4HGG7|A4HGG7_LEIBR              LLPVQKVLEDAKLKKSDIDEIVLVGGSTRIPKVQQLIKDFFGGKEPNKGINPDEAVAYGA    403
tr|S9U467|S9U467_9TRYP              LVPVQKVLEDAKLKKSDIDEIVLVGGSTRIPKVQQLIKDFFGGKEPNKGINPDEAVAYGA    472
tr|S9W3A0|S9W3A0_9TRYP              LVPVQKVLEDAKLKKSDIDEIVLVGGSTRIPKVQQLIKDFFGGKEPNKGINPDEAVAYGA    402
tr|A0A1X0NYK3|A0A1X0NYK3_9TRYP      LVPVQRVLEDAKLKKSDIDEIVLVGGSTRVPKVQQLIRDFFGGKEPNRGINPDEAVAYGA    400
tr|Q4D620|Q4D620_TRYCC              LVPVQRVLEDAKLKKSDIHEIVLVGGSTRVPKVQQLIRDFFGGKEPNRGINPDEAVAYGA    395
tr|K2NLB6|K2NLB6_TRYCR              LVPVQRVLEDAKLKKSDIHEIVLVGGSTRVPKVQQLIRDFFGGKEPNRGINPDEAVAYGA    395
tr|A0A422P6Q7|A0A422P6Q7_9TRYP      LVPVQRVLEDAKLKKSDIDEIVLVGGSTRVPKVQQLIRDFFGGKEPNRGINPDEAVAYGA    397
tr|A0A422N1T5|A0A422N1T5_TRYRA      LVPVQRVLEDAKLKKSEIDEIVLVGGSTRVPKVQQLIRDFFGGKEPNRGINPDEAVAYGA    397
tr|F9WRV9|F9WRV9_TRYVY              LVPVQRVLEDAKLKKSDIHEIVLVGGSTRVPKVQQLISDFFGGKELNRGINPDEAVAYGA    401
tr|F9W3P7|F9W3P7_TRYCI              LIPVQRVLEDAKLKKSDIHEIVLVGGSTRVPKVQQLISDFFGGKELNRGINPDEAVAYGA    400
tr|Q384Q5|Q384Q5_TRYB2              LVPVQRVLEDAKLKKSDIHEIVLVGGSTRVPKVQQLISDFFGGKELNRGINPDEAVAYGA    400
tr|A0A1Y2AI48|A0A1Y2AI48_9FUNG      LKPVEKVMKDAGLDKHEIHDIVLVGGSTRIPKVVQLLEDYFNGKKASKGINPDEAVAYGA    398
tr|A0A1Y1X8R1|A0A1Y1X8R1_9FUNG      IRPVEKVLKDAGLGKHEIHDIVLVGGSTRIPKIQQLIEDFFNGKKASKGINPDEAVAYGA    403
sp|P11021|BIP_HUMAN                 MKPVQKVLEDSDLKKSDIDEIVLVGGSTRIPKIQQLVKEFFNGKEPSRGINPDEAVAYGA    398
tr|A0A0G4EPG2|A0A0G4EPG2_VITBC      LKPVKNVLDDAGFKKTQIDEIVLVGGSTRIPKIQQLIKDFFNGKEPNRGINPDEAVAYGA    396
tr|M1BLB0|M1BLB0_SOLTU              MTPVKKAMEDAGLAKNQIDEIVLVGGSTRIPKVQQLLKDYFDGKEPSKGVNPDEAVAFGA    405
sp|Q9LKR3|BIP1_ARATH                MGPVKKAMDDAGLQKSQIDEIVLVGGSTRIPKVQQLLKDFFEGKEPNKGVNPDEAVAYGA    404
tr|I1KPN3|I1KPN3_SOYBN              MGPVKKAMEDAGLQKSQIDEIVLVGGSTRIPKVQQLLKDYFDGKEPNKGVNPDEAVAYGA    404
                                    : **. .: *: : * :*.:*********:**: **: ::*  *: .:*:*******:**

tr|A0A0L1KCY6|A0A0L1KCY6_9EUGL      AVQGAVLCGDS-AVKDKVLLVDVIPLSLGIETVGGVMTKLVERNTQIPAKKSQIFSTASD    452
tr|A0A0S4JK39|A0A0S4JK39_BODSA      AVQAAVLTGAR-DFKNTVVVADVVPLSLGIETVGGVMTRVIERNTQIPTKKSQTFTTYED    466
tr|A0A7G2CL24|A0A7G2CL24_9TRYP      AVQAAVMTGES-EVGGKVLLVDVISLSLGIETVGGVMTKLVERNTQIPAKKSQVFSTYQD    462
tr|A0A0S4IPK6|A0A0S4IPK6_BODSA      AVQAAVLTGDS-DLGNKVVLVDVIPLSLGIETVGGVMTKLIERNTQIPTKKSQVFSTYQD    459
tr|A0A0N0DZK2|A0A0N0DZK2_LEPPY      AVQAAVLMGES-EVGGKVVLVDVIPLSLGIETVGGVMTKLIERNTQIPTKKSQVFSTYQD    462
tr|A4I3J9|A4I3J9_LEIIN              AVQAAVLTGES-EVGGKVVLVDVIPLSLGIETVGGVMTKLIERNTQIPTKKSQIFSTYQD    462
tr|Q4Q8E6|Q4Q8E6_LEIMA              AVQAAVLTGES-EVGGKVVLVDVIPLSLGIETVGGVMTKLIERNTQIPTKKSQIFSTYQD    462
tr|A4HGG7|A4HGG7_LEIBR              AVQAAVLTGES-EVGGKVVLVDVIPLSLGIETVGGVMTKLIERNTQIPTKKSQVFSTYQD    462
tr|S9U467|S9U467_9TRYP              AVQAAVLTGES-EVGGKVVLVDVIPLSLGIETVGGVMTKLIERNTQIPTKKSQVFSTYQD    531
tr|S9W3A0|S9W3A0_9TRYP              AVQAAVLTGES-EVGGKVVLVDVIPLSLGIETVGGVMTKLIERNTQIPTKKSQVFSTYQD    461
tr|A0A1X0NYK3|A0A1X0NYK3_9TRYP      AVQAAVLTGES-EVGGRVVLVDVIPLSLGIETVGGVMTKLIERNTQIPTKKSQVFSTYQD    459
tr|Q4D620|Q4D620_TRYCC              AVQAAVLTGES-EVGGRVVLVDVIPLSLGIETVGGVMTKLIERNTQIPTKKSQVFSTYQD    454
tr|K2NLB6|K2NLB6_TRYCR              AVQAAVLTGES-EVGGRVVLVDVIPLSLGIETVGGVMTKLIERNTQIPTKKSQVFSTYQD    454
tr|A0A422P6Q7|A0A422P6Q7_9TRYP      AVQAAVLTGES-EVGGRVVLVDVIPLSLGIETVGGVMTKLIERNTQIPTKKSQVFSTYQD    456
tr|A0A422N1T5|A0A422N1T5_TRYRA      AVQAAVLTGES-EVGGRVVLVDVIPLSLGIETVGGVMTKLIERNTQIPTKKSQVFSTYQD    456
tr|F9WRV9|F9WRV9_TRYVY              AVQAAVLTGES-EVGGRVVLVDVIPLSLGIETVGGIMTKLIERNTQIPTKKSQVFSTHAD    460
tr|F9W3P7|F9W3P7_TRYCI              AVQAAVLTGES-EVGGRVVLVDVIPLSLGIETVGGVMTKLIERNTQIPTKKSQIFSTHAD    459
tr|Q384Q5|Q384Q5_TRYB2              AVQAAVLTGES-EVGGRVVLVDVIPLSLGIETVGGVMTKLIERNTQIPTKKSQVFSTHAD    459
tr|A0A1Y2AI48|A0A1Y2AI48_9FUNG      AVQGGVLSGNS-DELNNVLLLDVNPLTLGIETTGGVMTKLIARNTQIPTKKSQIFSTAAD    457
tr|A0A1Y1X8R1|A0A1Y1X8R1_9FUNG      AVQGGVLSGEDKENLSNVLLLDVNPLTLGIETTGGVMTKLIGRNTQIPTKKSQIFSTAAD    463
sp|P11021|BIP_HUMAN                 AVQAGVLSGD--QDTGDLVLLDVCPLTLGIETVGGVMTKLIPRNTVVPTKKSQIFSTASD    456
tr|A0A0G4EPG2|A0A0G4EPG2_VITBC      AVQAGILSGEGGQ---ELLLLDVTPLTLGIETVGGVMTKLINRNTVIPTKKSQVFSTYQD    453
tr|M1BLB0|M1BLB0_SOLTU              AVQGGILSGEGGDETKDILLLDVAPLTLGIETVGGVMTKLIPRNTVIPTKKSQVFTTYQD    465
sp|Q9LKR3|BIP1_ARATH                AVQGGILSGEGGDETKDILLLDVAPLTLGIETVGGVMTKLIPRNTVIPTKKSQVFTTYQD    464
tr|I1KPN3|I1KPN3_SOYBN              AVQGSILSGEGGEETKDILLLDVAPLTLGIETVGGVMTKLIPRNTVIPTKKSQVFTTYQD    464
                                    ***..:: *         ::: **  *:*****.**:**::: *** :*:**** *:*  *

tr|A0A0L1KCY6|A0A0L1KCY6_9EUGL      NQPSVTIVVYEGERAMSKDNRELGKFDLGGITPAPRGSPQIEVTFDVDENGIMEVSAKDN    512
tr|A0A0S4JK39|A0A0S4JK39_BODSA      NQPGVLIQVFEGERQMTKDNRMLGKFELTGIRRAAKGVPQIEVTFDVDEGSILQVSATDK    526
tr|A0A7G2CL24|A0A7G2CL24_9TRYP      NQPGVLIQVFEGERQMTKDNRLLGKFELSGIPPAPRGVPQIEVSFDVDENSILQVSAVDK    522
tr|A0A0S4IPK6|A0A0S4IPK6_BODSA      NQPGVLIQVFEGERQMTKDNRLLGKFELSGIPPAPRGTPQIEVTFDVDENSILQVSATDK    519
tr|A0A0N0DZK2|A0A0N0DZK2_LEPPY      NQPGVLIQVFEGERQMTKDNRLLGKFELSGIPPAPRGVPQIEVSFDVDENSILQVTASDK    522
tr|A4I3J9|A4I3J9_LEIIN              NQPSVLIQVFEGERGMTKDNRLLGKFDLSGIPPAPRGVPQIEVAFDVDENSILQVTASDK    522
tr|Q4Q8E6|Q4Q8E6_LEIMA              NQPSVLIQVFEGERGMTKDNRLLGKFDLSGIPPAPRGVPQIEVAFDVDENSILQVTASDK    522
tr|A4HGG7|A4HGG7_LEIBR              NQPGVLIQVFEGERQMTKDNRLLGKFELSGIPPAPRGVPQIEVAFDVDENSILQVTASDK    522
tr|S9U467|S9U467_9TRYP              NQPGVLIQVFEGERQMTKDNRLLGKFELSGIPPAPRGVPQIEVSFDVDENSILQVGAVDK    591
tr|S9W3A0|S9W3A0_9TRYP              NQPGVLIQVFEGERQMTKDNRLLGKFELSGIPPAPRGVPQIEVSFDVDENSILQVGAVDK    521
```

```
tr|A0A1X0NYK3|A0A1X0NYK3_9TRYP      NQPGVLIQVFEGERQMTKDNRLLGKFELSGIPPAARGVPQIEVTFDVDENSILQVSAVDK    519
tr|Q4D620|Q4D620_TRYCC              NQPGVLIQVYEGERQMTKDNRLLGKFELSGIPPAPRGVPQIEVTFDVDENSILQVSAVDK    514
tr|K2NLB6|K2NLB6_TRYCR              NQPGVLIQVYEGERQMTKDNRLLGKFELSGIPPAPRGVPQIEVTFDVDENSILQVSAVDK    514
tr|A0A422P6Q7|A0A422P6Q7_9TRYP      NQPGVLIQVFEGERQMTKDNRLLGKFELSGIPPAARGVPQIEVTFDVDENSILQVSAVDK    516
tr|A0A422N1T5|A0A422N1T5_TRYRA      NQPGVLIQVFEGERQMTKDNRLLGKFELSGIPPAARGVPQIEVTFDVDENSILQVSAVDK    516
tr|F9WRV9|F9WRV9_TRYVY              NQPGVLIQVYEGERQLTKDNRLLGKFELSGIPPAPRGVPQIEVTFDVDENSILQVSAVDK    520
tr|F9W3P7|F9W3P7_TRYCI              NQPGVLIQVYEGERQLTKDNRLLGKFELTGIPPAARGVPQIEVTFDVDENSILQVSAMDK    519
tr|Q384Q5|Q384Q5_TRYB2              NQPGVLIQVYEGERQLTKDNRLLGKFELSGIPPAARGVPQIEVTFDVDENSILQVSAMDK    519
tr|A0A1Y2AI48|A0A1Y2AI48_9FUNG      NQPTVLIQVFEGERPLTKDNNLLGKFDLNGIPPAPRGTPQIEVTFEISVDGLLRVAAHDK    517
tr|A0A1Y1X8R1|A0A1Y1X8R1_9FUNG      NQSIVLIQVYEGERSMTKDNNLLGKFELKDIPPAPRGVPQIEVTFEIDVNGILKVSAEDK    523
sp|P11021|BIP_HUMAN                 NQPTVTIKVYEGERPLTKDNHLLGTFDLTGIPPAPRGVPQIEVTFEIDVNGILRVTAEDK    516
tr|A0A0G4EPG2|A0A0G4EPG2_VITBC      NQPAVMIQVFEGERPMTKDNHLLGKFELSGIPPAPRGVPQIEVTFEIDSNGILNVSAEDK    513
tr|M1BLB0|M1BLB0_SOLTU              QQTTVTISVFEGERSMVKDCRLLGKFDLTGIPPAPRGTPQIEVTFEVDANGILNVKAEDK    525
sp|Q9LKR3|BIP1_ARATH                QQTTVSIQVFEGERSLTKDCRLLGKFDLNGIPPAPRGTPQIEVTFEVDANGILNVKAEDK    524
tr|I1KPN3|I1KPN3_SOYBN              QQTTVSIQVFEGERSLTKDCRLLGKFDLSGIPPAPRGTPQIEVTFEVDANGILNVKAEDK    524
                                    :*  * * *:**** : ** . **.*:* .*  * :* *****:*::. ..::.* * *:

tr|A0A0L1KCY6|A0A0L1KCY6_9EUGL      STNKEEKITITNDKGRLSQEEIDRMVREAEEHGEEDRIKKETIEARNHLESIAYSLKNAV    572
tr|A0A0S4JK39|A0A0S4JK39_BODSA      SSNTKEEITIRNDKGRLSQREIDEMVREAELFEEEDRKLRGKVEARTALESAVYSLKGEL    586
tr|A0A7G2CL24|A0A7G2CL24_9TRYP      SSGKREEIRISNDKGRLTDEEIDAMIREAAQFEEEDRKVRERVEARNSLEGIAYSLRNQV    582
tr|A0A0S4IPK6|A0A0S4IPK6_BODSA      SSNKREEITITNDKGRLSEEEIERMVREAAEFEEEDRKLREKVEARNSLESVAYSLKNQI    579
tr|A0A0N0DZK2|A0A0N0DZK2_LEPPY      SSGKREEITITNDKGRLSEAEIEAMVEEAAQFAEEDRKLRERVEAKNSLESIAYSLRNQV    582
tr|A4I3J9|A4I3J9_LEIIN              SSGKREEITITNDKGRLSEEEIERMVREAAEFEDEDRKVRERVEAKNSLESIAYSLRNQI    582
tr|Q4Q8E6|Q4Q8E6_LEIMA              SSGKREEITITNDKGRLSEEEIERMVREAAEFEDEDRKVRERVEAKNSLESIAYSLRNQI    582
tr|A4HGG7|A4HGG7_LEIBR              SSGKREEITITNDKGRLSEEEIERMVREAEEFEDEDRKVRERVEAKNSLESIAYSLRNQI    582
tr|S9U467|S9U467_9TRYP              SSGKKEEITITNDKGRLSEEDIERMVREAAENEESDRKVRERVEAKNSLESIAYSLRNQV    651
tr|S9W3A0|S9W3A0_9TRYP              SSGKKEEITITNDKGRLSEEDIERMVREAAENEESDRKVRERVEAKNSLESIAYSLRNQV    581
tr|A0A1X0NYK3|A0A1X0NYK3_9TRYP      SSGKKEEITITNDKGRLSEEEIERMVREAAEFEEEDRKVRERVDARNSLEGIAYSLRTQV    579
tr|Q4D620|Q4D620_TRYCC              SSGKKEEITITNDKGRLSEEEIERMVREAAEFEDEDRKVRERVDARNSLESVAYSLRNQV    574
tr|K2NLB6|K2NLB6_TRYCR              SSGKKEEITITNDKGRLSEEEIERMVREAAEFEDEDRKVRERVDARNSLESTAYSLRNQV    574
tr|A0A422P6Q7|A0A422P6Q7_9TRYP      SSGKKEEITITNDKGRLSEEEIERMVREAAEFEDEDRKVRERVEARNSLESIAYSLRNQV    576
tr|A0A422N1T5|A0A422N1T5_TRYRA      SSGKKEEITITNDKGRLSEEEIERMVREAAEFEDEDRKVRERVEARNSLESIAYSLRNQV    576
tr|F9WRV9|F9WRV9_TRYVY              SSGKKEEITITNDKGRLSEEEIERMVREAAEFENEDRKVRERVDARNTLESVTYSLRSQV    580
tr|F9W3P7|F9W3P7_TRYCI              SSGKKEEITITNDKGRLSEEEIERMVREAAEFEDEDRKVRERVEARNSLESAAYSLRNQV    579
tr|Q384Q5|Q384Q5_TRYB2              SSGKKEEITITNDKGRLSEEEIERMVREAAEFEDEDRKVRERVDARNSLESVAYSLRNQV    579
tr|A0A1Y2AI48|A0A1Y2AI48_9FUNG      GTGKSESITITNDKGRLTEEEIERMVREAEQFAEEDQLLKEKIEAKNGLENYIYQIKQQV    577
tr|A0A1Y1X8R1|A0A1Y1X8R1_9FUNG      GTGKSESITITNDKGRLSEEEIERMVAEAEQFAEEDKLLKEKIEAKNTLENYLYTIKNQV    583
sp|P11021|BIP_HUMAN                 GTGNKNKITITNDQNRLTPEEIERMVNDAEKFAEEDKKLKERIDTRNELESYAYSLKNQI    576
tr|A0A0G4EPG2|A0A0G4EPG2_VITBC      GTGKSEKITITNDKGRLSQEDIEKMIKEAEDYAEEDKKVRERVDAKNSFDNYLHSMKQSV    573
tr|M1BLB0|M1BLB0_SOLTU              ASGKSEKITITNDKGRLSQEEIERMVKEAEEFAEEDKKVKERVDARNSLETYVYNMRNQI    585
sp|Q9LKR3|BIP1_ARATH                ASGKSEKITITNEKGRLSQEEIDRMVKEAEEFAEEDKKVKEKIDARNALETYVYNMKNQV    584
tr|I1KPN3|I1KPN3_SOYBN              GTGKSEKITITNEKGRLSQEEIDRMVREAEEFAEEDKKVKERIDARNSLETYVYNMKNQV    584
                                    .:.. :.* * *::.**:  :*: *: :*      :.*:  :  ::::. ::   : ::  :

tr|A0A0L1KCY6|A0A0L1KCY6_9EUGL      NDEEKLGNKLDDSEKETIKEAVRETIEWMDENPT-ADKSDYEERLEKLQSVTNPIIQKVN    631
tr|A0A0S4JK39|A0A0S4JK39_BODSA      GDVTKLVGL-AGDERAALEVAVREAMSFLDENPN-AELEEYAEARERLQSVAGPIMRRVQ    644
tr|A0A7G2CL24|A0A7G2CL24_9TRYP      NDKEKLGGKLSDEDKKAIEGAVKEALDFVDENPN-AEKEDYKTAQEKLQSVTNPIVQKVY    641
tr|A0A0S4IPK6|A0A0S4IPK6_BODSA      NDKDKLKDKLDADDKQAVEDAVKEAINFLDENPN-AEKEEYDEAKEKLQSVTNPIIQKAY    638
tr|A0A0N0DZK2|A0A0N0DZK2_LEPPY      NDKEKLGDKLDADDKKAIEAAVRVALDFVDENPN-ADKEEFEEAREQLQKVTNPIIQKVY    641
tr|A4I3J9|A4I3J9_LEIIN              NDKDKLGDKLAADDKKAIEEAVKDALDFVDENPN-ADREEFEAARTKLQSVTNPIIQKVY    641
tr|Q4Q8E6|Q4Q8E6_LEIMA              NDKDKLGDKLDADDKKAIEEAVKDALDFVDENPN-ADREEFEAARTKLQSVTNPIIQKVY    641
tr|A4HGG7|A4HGG7_LEIBR              NDKDKLGEKLDADDKKAVEEAVKDALAFVDENPN-ADREDFEGAREKLQSVTNPIIQKVY    641
tr|S9U467|S9U467_9TRYP              NDKEKLGGKIDAEDKKAVEDAVKEAIDFVDENPS-AEKEEYDAAREKLQSVTNPIIQKVY    710
tr|S9W3A0|S9W3A0_9TRYP              NDKEKLGGKIDAEDKKAVEDAVKEAIDFVDENPS-AEKEEYDAAREKLQSVTNPIIQKVY    640
tr|A0A1X0NYK3|A0A1X0NYK3_9TRYP      NDKDKLGGKLSADDKSAVEAAVKEAIQFLDENPN-AEKEEYDEAREKLQSVTNPIIQKAY    638
tr|Q4D620|Q4D620_TRYCC              NDKEKLGGKLSADDKSAVEAAVKEAMQFLDDNPN-ADKEEYDEARDKLQSVTNPIIQKVY    633
tr|K2NLB6|K2NLB6_TRYCR              NDKEKLGGKLSAEDKSAVEAAVKEAMQFLDDNPN-ADKEEYDEARDKLQSVTNPIIQKVY    633
tr|A0A422P6Q7|A0A422P6Q7_9TRYP      NDKEKLGGKLSAEDKSAVEDAVKEAIHFLDENPN-ADKEEYDEAREKLQSVTNPIIQKAY    635
tr|A0A422N1T5|A0A422N1T5_TRYRA      NDKEKLGGKLSAEDKSAVEAAVKEAIHFLDENPN-ADKEEYDEAREKLQGVTNPIIQKAY    635
tr|F9WRV9|F9WRV9_TRYVY              NDKDKLGGKLSADEKSTVEAAVKEAIRFLDENPN-AEKEEYDAAREKLQGVTNPIIQKAY    639
tr|F9W3P7|F9W3P7_TRYCI              NDKDKLGGKLSADDKAAVEAAVKEAIRFLDENPN-AEKEEYKTALDTMQSVTNPIVQKAY    638
tr|Q384Q5|Q384Q5_TRYB2              NDKDKLGGKLDPNDKAAVETAVAEAIRFLDENPN-AEKEEYKTALETLQSVTNPIIQKTY    638
tr|A0A1Y2AI48|A0A1Y2AI48_9FUNG      TDEDKLGGKIDEDDKKAILDAIKEANEWVDANIATATKEDIDEQKSELEAIVNPITSKLY    637
tr|A0A1Y1X8R1|A0A1Y1X8R1_9FUNG      NDEEGLGSKISEDDKKKILDAVEEKITWLDENGNTAVKEDFEEQKSAVESIVNGITSKLY    643
sp|P11021|BIP_HUMAN                 GDKEKLGGKLSSEDKETMEKAVEEKIEWLESHQD-ADIEDFKAKKKELEEIVQPIISKLY    635
tr|A0A0G4EPG2|A0A0G4EPG2_VITBC      EDKDKLANKLDEDQKQQILDAVKDGEEWMQEHPE-EDAEEIKAKHKEIEEICNPIISAVY    632
tr|M1BLB0|M1BLB0_SOLTU              NDKDKLADKLESDEKEKIETATKEALEWLDDNQN-AEKEDYEEKLKEVEAVCNPIITAVY    644
sp|Q9LKR3|BIP1_ARATH                NDKDKLADKLEGDEKEKIEAATKEALEWLDENQN-SEKEEYDEKLKEVEAVCNPIITAVY    643
tr|I1KPN3|I1KPN3_SOYBN              SDKDKLADKLESDEKEKIETAVKEALEWLDDNQS-VEKEDYEEKLKEVEAVCNPIISAVY    643
                                    *   *    .::  :  *         ::: :    .:       :: :    *

tr|A0A0L1KCY6|A0A0L1KCY6_9EUGL      QESGSAGGYQSE-----TMEDL-------------------    648
tr|A0A0S4JK39|A0A0S4JK39_BODSA      EATAGGAGDEPE-----SVDDL-------------------    661
tr|A0A7G2CL24|A0A7G2CL24_9TRYP      QATGGAPDAEAE-----PMDDL-------------------    658
tr|A0A0S4IPK6|A0A0S4IPK6_BODSA      RGSGGAGGEEGGADATESMDDL-------------------    660
tr|A0A0N0DZK2|A0A0N0DZK2_LEPPY      QAAGGAAGEEEP----DAMDDL-------------------    659
tr|A4I3J9|A4I3J9_LEIIN              QGAAGS-GAEEA----DAMDDL-------------------    658
tr|Q4Q8E6|Q4Q8E6_LEIMA              QGTAGS-GAEEA----DAMDDL-------------------    658
tr|A4HGG7|A4HGG7_LEIBR              QAGGAP-GSEPT----DAMDDL-------------------    658
tr|S9U467|S9U467_9TRYP              AAAGGSPDSEAA----EPMDDL-------------------    728
tr|S9W3A0|S9W3A0_9TRYP              AAAGGSPDSEAA----EPMDDL-------------------    658
tr|A0A1X0NYK3|A0A1X0NYK3_9TRYP      QAGGGAADEE-------KXQSRWPCLRIWVTIKQISARQRQ    672
tr|Q4D620|Q4D620_TRYCC              QSGGGADGDE-------RPEPMDDL----------------    651
tr|K2NLB6|K2NLB6_TRYCR              QSGGGADGDE-------RPEPMDDL----------------    651
tr|A0A422P6Q7|A0A422P6Q7_9TRYP      QAGGASGEEG-------QPEPMDDL----------------    653
tr|A0A422N1T5|A0A422N1T5_TRYRA      QAGGASGEDE-------QPEPMDDL----------------    653
tr|F9WRV9|F9WRV9_TRYVY              QAG--GEKPQ-------PMDDL-------------------    652
tr|F9W3P7|F9W3P7_TRYCI              QSAGAGDKPQ-------PMDDL-------------------    653
```

```
tr|Q384Q5|Q384Q5_TRYB2              QSAGGGDKPQ-------PMDDL-------------------    653
tr|A0A1Y2AI48|A0A1Y2AI48_9FUNG      GGSAGG---DGE------DVPDHDEL---------------    654
tr|A0A1Y1X8R1|A0A1Y1X8R1_9FUNG      GQQKGDATMEDE------DIPDHDEL---------------    663
sp|P11021|BIP_HUMAN                 GSAGPPPTGEEDTAE-------KDEL---------------    654
tr|A0A0G4EPG2|A0A0G4EPG2_VITBC      GAAGGAGGPTGEE-----DYEAHDEL---------------    653
tr|M1BLB0|M1BLB0_SOLTU              QRSGGAPGGASEDSN--EDDDSHDEL---------------    668
sp|Q9LKR3|BIP1_ARATH                QRSGGAPGGAGGESSTEEEDESHDEL---------------    669
tr|I1KPN3|I1KPN3_SOYBN              QRSGGAPGGAGGEGED--EDDSHDEL---------------    667
```

### 10. HYOU1-like HSP70 chaperones

*******************************

The animal Hypoxia upregulated protein 1 (HYOU1) is a representing member of this chaperone family that is found in almost all eukaryotes studied. All examined kinetoplastids also possess at least a single gene of this group. Although the KDEL signal can be missing in extreme cases, most members of this family present a very well conserved, ancestral retention signal at their C-termini. Note that HYOU1-like chaperones are rather different from BIP family members, despite the shared HSP70 domain core. They seem to have diverged already at the stem of Eukaryota, therefore they should be treated a separate family.

```
tr|A0A0C9MAJ6|A0A0C9MAJ6_9FUNG      ------------------------------------------------------------    0
sp|Q9Y4L1|HYOU1_HUMAN               -----------------------MADKVRRQRPRRRVCWALVAVL----LADLLALSDTL    33
sp|Q63617|HYOU1_RAT                 -----------------------MAATVRRQRPRRLLCWALVAVL----LADLLALSDTL    33
tr|A0A067CBX2|A0A067CBX2_SAPPC      ---------------------------------MRL---SHLLPL----LLAALVATVSA    20
tr|D8RVG4|D8RVG4_SELML              ---------------------------------MRNLVLPALLL------LFSTLAIVQA    21
tr|A0A1P8B8R9|A0A1P8B8R9_ARATH      --------MWLLLYNCRGIAALYSRCLVYDWNMGKIFSWLVVLLS----LISLVPVPSES    48
tr|A0A176WKM4|A0A176WKM4_MARPO      -----------MAPEER---------GRRRWRSMASL--PVVLI------VLCLLRPANG    32
tr|A0A3R7K087|A0A3R7K087_9TRYP      --MNRWRLPGGLLAGRRGEASGRCKAAACCATRAPPLLTAAFFLLTLAS--IVACTAGAY    56
tr|Q4E175|Q4E175_TRYCC              MKMEGEQWMGGVAGVCRGKASHFLKATSCSTTRSPLLLFCIILLLTGTP--FMSFLVEAH    58
tr|Q4E0G9|Q4E0G9_TRYCC              MKMEGEQWMGGVAGVRRGKASHFLKATSCSTTRSLLLLFCIILLLTGTP--FMSLLVEAH    58
tr|A0A6A5CDJ2|A0A6A5CDJ2_NAEFO      ---------------------MF-GFTTPRSSGVGTLISVALFVIIL--SYILVEPSHQG    36
tr|A0A0S4IUZ2|A0A0S4IUZ2_BODSA      ---------------------MQAHKRSPRKRSLPLAM----LMLTVTAVLC--IGCTSG    33
tr|A0A0L1KXM1|A0A0L1KXM1_9EUGL      ---------------------------MSRSLISPVALTL--LSMAIVHELCPPICMVQA    31
tr|A0A0S4JJU2|A0A0S4JJU2_BODSA      ------------------------------------------MLKLLLCA-CILVVVANA    17
tr|A0A0N0VI62|A0A0N0VI62_LEPPY      -----------------------------------MACKNLLLAALMVAVFAILSSTSSA    25
tr|A4HND7|A4HND7_LEIBR              ---------------------------------MKLKYRVVFLAAVLACLFE--PLASLA    25
tr|A4IC10|A4IC10_LEIIN              ---------------------------------MKLKCKVAFLAYVLACLFI--AQASLA    25
tr|E9AFU8|E9AFU8_LEIMA              ---------------------------------MKPKCNVVLLAYALACFFI--TQASLA    25
tr|A0A7G2C9U8|A0A7G2C9U8_9TRYP      ------------------------------------------------------------    0
tr|S9TX72|S9TX72_9TRYP              ----------------------------MKA------LATFLLLA----FAVCFCGVASG    22
tr|S9UBL2|S9UBL2_9TRYP              ----------------------------MKA------LATFLLLA----FAVCFCGVASG    22
tr|Q38E16|Q38E16_TRYB2              ----------------------------MRVLANEIRRMLVAFLL----TAAFGYDVTSA    28
tr|F9W5M9|F9W5M9_TRYCI              ---------------------------------MHLGVGLVLLLV----TATFACDVTFG    23
tr|A0A1X0P4V9|A0A1X0P4V9_9TRYP      -------------------------------------MRIIFFFV----AVF-YLCTISY    18
tr|Q4CS27|Q4CS27_TRYCC              -------------------------------------MQIVSFVI----LAILLSYGASA    19
tr|Q4E4F4|Q4E4F4_TRYCC              -------------------------------------MQIVSFVI----LAILLSYGASA    19
tr|A0A422P1B1|A0A422P1B1_TRYRA      -------------------------------------MRIVFLVI----LTLLLSDGVSA    19
tr|A0A422N0X0|A0A422N0X0_9TRYP      -------------------------------------MRLITLVV----LTLLSSYGVSA    19


tr|A0A0C9MAJ6|A0A0C9MAJ6_9FUNG      --MSIDYGTEWFKVGLIK--P-GMP----LDVALNKDSKRKTQSVVTIR-----------    40
sp|Q9Y4L1|HYOU1_HUMAN               AVMSVDLGSESMKVAIVK--P-GVP----MEIVLNKESRRKTPVIVTLK-----------    75
sp|Q63617|HYOU1_RAT                 AVMSVDLGSESMKVAIVK--P-GVP----MEIVLNKESRRKTPVTVTLK-----------    75
tr|A0A067CBX2|A0A067CBX2_SAPPC      SVAGVDFGGEFFKIALVK--P-GRP----FEIVTNVHSKRKTETIVSFN-----------    62
tr|D8RVG4|D8RVG4_SELML              AVFSVDLGAEWMKVAVVDVKPGQSP----IGVALNEMSKRKSSSVVAFS-----------    66
tr|A0A1P8B8R9|A0A1P8B8R9_ARATH      AVLSVDLGSEWVKVAVVNLKRGQSP----ISVAINEMSKRKSPALVAFQ-----------    93
tr|A0A176WKM4|A0A176WKM4_MARPO      AVFSIDLGSEWMKVAVVNLKPGQAP----IAIAINEMSKRKSPALVAFS-----------    77
tr|A0A3R7K087|A0A3R7K087_9TRYP      HVLAVDLGVEWAKAATLG---GGSGASSNPAVVLNDQANRKSPQCIAFRFLPYDTDDVLQ    113
tr|Q4E175|Q4E175_TRYCC              HILAVDLGVDWAKAATLG---GGSGASLIPTIVLNDQANRKSPQCIAFRFLPYDTDDILQ    115
tr|Q4E0G9|Q4E0G9_TRYCC              HILAVDLGVDWAKAATLG---GGSGASLIPTIVLNDQANRKSPQCIAFRFLPYDTDDILQ    115
tr|A0A6A5CDJ2|A0A6A5CDJ2_NAEFO      PIVGIDLGSRWLKVGIAK---SGSP----IDLVLNEQSKRKTSNIIGFR-----------    78
tr|A0A0S4IUZ2|A0A0S4IUZ2_BODSA      TIVAIDMGSEFLKIAAPK----DTS----IEVCLNEQSHRKSDTWIGFR-----------    74
tr|A0A0L1KXM1|A0A0L1KXM1_9EUGL      QVVGIDLGSEFIKIASVR---RSDG----IDIVLNEETRRKTPHYVGFR-----------    73
tr|A0A0S4JJU2|A0A0S4JJU2_BODSA      NVMGIDFGSEFVKVTSPH---GNAS----IDIVLNEQTRRKSNNFVGFR-----------    59
tr|A0A0N0VI62|A0A0N0VI62_LEPPY      HVIGVDFGSEYIKVSGPH---GDKG----LDIVLNELSMRKTENFIGFR-----------    67
tr|A4HND7|A4HND7_LEIBR              HVIGVDLGSEYIKVAGPH---GDKG----IDIVLNEQSRRKTDNFIGFH-----------    67
tr|A4IC10|A4IC10_LEIIN              HVIGVDLGSEYIKVAGPH---GDKG----VDIVLNEQSRRKTDNFIGFR-----------    67
tr|E9AFU8|E9AFU8_LEIMA              HVIGVDLGSEYIKVAGPH---GDKG----VDIVLNEQSRRKTDNFIGFR-----------    67
tr|A0A7G2C9U8|A0A7G2C9U8_9TRYP      ------------------------------------------------------------    0
tr|S9TX72|S9TX72_9TRYP              HVLGVDFGSEFIKIAAPH---GDQV----VDIVLNELSRRKTENFIGFR-----------    64
tr|S9UBL2|S9UBL2_9TRYP              HVLGVDFGSEFIKIAAPH---GDQV----VDIVLNELSRRKTENFIGFR-----------    64
tr|Q38E16|Q38E16_TRYB2              NVIGVDFGSDYIEVAGPI---NGVN----VDIILNEQSHRKTDNYIGFR-----------    70
tr|F9W5M9|F9W5M9_TRYCI              SVMGIDFGSDYIEIAGPI---NGVN----VNIVLNEQSHRKTDNYIGFR-----------    65
tr|A0A1X0P4V9|A0A1X0P4V9_9TRYP      ANVGIDFGSDYIEVAGPH---TGNN----VDIVLNEQSHRKTDSYIGFR-----------    60
tr|Q4CS27|Q4CS27_TRYCC              NVLGIDFGSEYIEVAGPH---NGNN----VDIVLNEQSHRKTDNYIGFK-----------    61
tr|Q4E4F4|Q4E4F4_TRYCC              NVLGIDFGSEYIEVAGPH---NGNN----VDIVLNEQSHRKTDNYIGFK-----------    61
tr|A0A422P1B1|A0A422P1B1_TRYRA      NVLGIDFGSDYIEVAGPH---NGNN----VDIVLNEQSHRKTDNYIGIK-----------    61
tr|A0A422N0X0|A0A422N0X0_9TRYP      NVLGIDFGSDYIEVAGPH---NGNN----VDIVLNEQSHRKTDNYIGFK-----------    61
```

```
tr|A0A0C9MAJ6|A0A0C9MAJ6_9FUNG      HDERIYGGDAISLAGRFPHLTYSNLKSIIAKKYDDPLTQEF----------------R--    82
sp|Q9Y4L1|HYOU1_HUMAN               ENERFFGDSAASMAIKNPKATLRYFQHLLGKQADNPHVALY----------------Q--    117
sp|Q63617|HYOU1_RAT                 ENERFLGDSAAGMAIKNPKATLRYFQHLLGKQADNPHVALY----------------R--    117
tr|A0A067CBX2|A0A067CBX2_SAPPC      GDERVYGADAMNIEVRRPQVAYSQIRRFLGATVDHPLVSSL---------------TE--    105
tr|D8RVG4|D8RVG4_SELML              GGNRLLAEEAMGIAARFPERVYSRVRDMVGKPSES--VKRI---------------AS--    107
tr|A0A1P8B8R9|A0A1P8B8R9_ARATH      SGDRLLGEEAAGITARYPNKVYSQLRDMVGKPFKH--VKDF---------------ID--    134
tr|A0A176WKM4|A0A176WKM4_MARPO      NGDRLLAEEAAGIVARYPERVYSRVRDMIGKPFES--VKQT---------------LD--    118
tr|A0A3R7K087|A0A3R7K087_9TRYP      KVERIFSEQALALEPRFPEHVVCGPSLLAGRGVWRGFVAGSDTGAAA-----QAALAPED    168
tr|Q4E175|Q4E175_TRYCC              RVERFFSEQAQSLEPRFPDQVVCGPSLLAGRGVSRVVTDGNNTGGVTQGSNNKDALSLDE    175
tr|Q4E0G9|Q4E0G9_TRYCC              RVERFFSEQAQSLEPRFPDQVVCGPSLLAGRGVSRVVTDGNNTGGVTQGSNNKDTLSLDE    175
tr|A0A6A5CDJ2|A0A6A5CDJ2_NAEFO      GKDRYIGEAGYTMVARFPDKMLRFLNFALGKSYDLELNEVS---------------RRYN    123
tr|A0A0S4IUZ2|A0A0S4IUZ2_BODSA      GEERFFGADAKSLSARFPDAMIPAVPRLVGVPHDAAFQQGW---------------LN--    117
tr|A0A0L1KXM1|A0A0L1KXM1_9EUGL      GKDSYIGEDAKNLVGRFPDMIFTLLNRLIGLSPTESLRQWY---------------HS--    116
tr|A0A0S4JJU2|A0A0S4JJU2_BODSA      GDDRFFGDEAKNLAPRFPDNMFSMINQLIGFPFESNKTAQF---------------QQ--    102
tr|A0A0N0VI62|A0A0N0VI62_LEPPY      NGDIYIGDTAKGLAARFPLCTASAINQLVSIRKDSELHAVF---------------QD--    110
tr|A4HND7|A4HND7_LEIBR              RSDLYIGDTAKSLAARFPLCTASAVNQLIGIRKDSHLLSFF---------------FD--    110
tr|A4IC10|A4IC10_LEIIN              RSDLYIGDTAKSLAARFPLCTASAVNQLVRIRKDSSLLPFF---------------RD--    110
tr|E9AFU8|E9AFU8_LEIMA              RSDLYIGDTAKSLAARFPLCTASAVNQLVRIRKDSSLLPFF---------------SD--    110
tr|A0A7G2C9U8|A0A7G2C9U8_9TRYP      ----------------------------MGLKKDSPEWKEF---------------ES--    15
tr|S9TX72|S9TX72_9TRYP              NGERYIGNEAKTLAARFPLKIASAINQLIGVQDKTEELKDF---------------EQ--    107
tr|S9UBL2|S9UBL2_9TRYP              NGERYIGNEAKTLAARFPLKIASAINQLIGVQDKTEELKDF---------------EQ--    107
tr|Q38E16|Q38E16_TRYB2              NGERSIGAQAKSLAARFPTNMIAMINHLVGITYNSSDFANF---------------KK--    113
tr|F9W5M9|F9W5M9_TRYCI              NGERSIGAQAKSLAARFPQNMLTMINHLIGITRNSSEFSNF---------------KQ--    108
tr|A0A1X0P4V9|A0A1X0P4V9_9TRYP      NGDRYIGDQAKALAARFPLNMVTMMNKLIGVTYNSTDFENF---------------KK--    103
tr|Q4CS27|Q4CS27_TRYCC              NGERYIGDQAKALAARFPLNMVTMINQLIGILCDSTEFAML---------------KD--    104
tr|Q4E4F4|Q4E4F4_TRYCC              NGERYIGDQAKALAARFPLNMVTMINQLIGILCDSTEFTML---------------KD--    104
tr|A0A422P1B1|A0A422P1B1_TRYRA      NGERYIGEQAKALAARFPLNMVTMINRLIGILCNSTEFANF---------------QN--    104
tr|A0A422N0X0|A0A422N0X0_9TRYP      NGERYIGDQAKALAARFPLNMVTMINRMIGISYNSTEFSDF---------------RN--    104


tr|A0A0C9MAJ6|A0A0C9MAJ6_9FUNG      RRYINNMI-V-DNERDMPVFIH------NETTQLSIEELIAYQFQNAKHQASVT--AGEK    132
sp|Q9Y4L1|HYOU1_HUMAN               ARFPEHELTF-DPQRQTVHFQI------SSQLQFSPEEVLGMVLNYSRSLAEDF--AEQP    168
sp|Q63617|HYOU1_RAT                 SRFPEHELNV-DPQRQTVRFQI------SPQLQFSPEEVLGMVLNYSRSLAEDF--AEQP    168
tr|A0A067CBX2|A0A067CBX2_SAPPC      NEYFPYTLTK-NLTRGSVALQH------SSEHTFHAEELAAMVFGHAKQITNDF--AEGP    156
tr|D8RVG4|D8RVG4_SELML              ASYLPYDFVE-ESPQVTSIRVD-------SQELYRSEELLGMILGYCRGLAEAN--VKAT    157
tr|A0A1P8B8R9|A0A1P8B8R9_ARATH      SVYLPFDIVE-DSRGAVGIKID------DGSTVYSVEELLAMILGYASNLAEFH--AKIP    185
tr|A0A176WKM4|A0A176WKM4_MARPO      ASYLPFDVIK-DDRGAVSIRTH------DKQSVFRSEELLAMVLSYGRDLAEAH--AKGV    169
tr|A0A3R7K087|A0A3R7K087_9TRYP      AASLTYAVVP-HASRDKLAVRIAGGRDKQSVLEFSAEELVGMFLTYLRRIAERGLDG-EP    226
tr|Q4E175|Q4E175_TRYCC              LASLTFSVVP-HSSRDKVAVRILGGKNK-SVLEFSTEELIGMFFAYLKRIAERGLNG-EP    232
tr|Q4E0G9|Q4E0G9_TRYCC              VASLTFSVVP-HSSRDKVTVRILGGKNK-SVLEFSTEELIGMFFAYLKRIAERGLNG-EP    232
tr|A0A6A5CDJ2|A0A6A5CDJ2_NAEFO      ELKVPVSL-VKNEERGTVDVKF------SNDATYSPEELLSMIFLYIKQLADD--GKTAA    174
tr|A0A0S4IUZ2|A0A0S4IUZ2_BODSA      KMRYTYHTASSGTTRGTVNVTFPASEG-VEERQYTADELMGMMLGYAKTQAEREISG--P    174
tr|A0A0L1KXM1|A0A0L1KXM1_9EUGL      DMLFTNALLP-IPERESMAFSVQRGAK-AVNIRYAIETLLGMTFEYVRSITASYVGEGCS    174
tr|A0A0S4JJU2|A0A0S4JJU2_BODSA      -LLTTFALKEGEERKGVNVVCR----Q-DPECNYSAELLIAMYFQYIKLITGK--DAKIK    154
tr|A0A0N0VI62|A0A0N0VI62_LEPPY      -FQYEYHVNF--NNHGSATVNI----C-NVEEPFTAEELFSLILSYCKAAAEH--DDVVT    160
tr|A4HND7|A4HND7_LEIBR              -LHYEYHIGF--NNHGSAIVSI----C-DNKDPFTAEELYSMVLSYCRTAAVN--DEVVD    160
tr|A4IC10|A4IC10_LEIIN              -LQYEYHVSF--NNHGSATVSI----C-DTKDPFTAEELYSMVLSYCKTTAVK--DDVVD    160
tr|E9AFU8|E9AFU8_LEIMA              -LQYEYHVGF--NNHGSATVSI----C-DTKDPFTAEELYSMVLSYCKTTAVK--DDVVD    160
tr|A0A7G2C9U8|A0A7G2C9U8_9TRYP      -LLLEYSIEA--NERGTPVVPF----G-EEHAPFSSEELFAFFLDYFYRIGVN--DGVVD    65
tr|S9TX72|S9TX72_9TRYP              -LLYEYKFGF--SPRGTSQVLI----D-GVKEPYTTEELYAMMFTYFKHIAVN--DDIID    157
tr|S9UBL2|S9UBL2_9TRYP              -LLYEYKFGF--SPRGTSQVLI----D-GVKEPYTTEELYAMMFTYFKHIAVN--DDIID    157
tr|Q38E16|Q38E16_TRYB2              -LQCEFDPHP--EERGTVGFRF----E-DNNDTYTAEEIYAMMLNYCRSISEK--AGVPN    163
tr|F9W5M9|F9W5M9_TRYCI              -LQCEFDPLA--EERDTVGFNF----T-DTGDTYTVEEIYAMMLNYCRSTSRN--AGVPS    158
tr|A0A1X0P4V9|A0A1X0P4V9_9TRYP      -LEYEFETRH--EEDNTVGFSF----G-GRDGNYTVEELYAMMLHYCQNIAQK--DGVND    153
tr|Q4CS27|Q4CS27_TRYCC              -LEFEFETRP--EERNTIGFCF----S--QDGNYTAEELYAMVLQYCQSISEK--DGVVD    153
tr|Q4E4F4|Q4E4F4_TRYCC              -LEFEFETRP--EERNTIGFCF----S--QDGNYTAEELYAMVLQYCQSISEK--DGVVD    153
tr|A0A422P1B1|A0A422P1B1_TRYRA      -MMLEFETRS--DERSTVGFSF----A--QDGNYTVEELYAMMLQYCQIISEK--DGIVN    153
tr|A0A422N0X0|A0A422N0X0_9TRYP      -LNFEFETRP--EERNTVGFSF----P--QDGSYTVEELYAMMLRYCQSISEK--DGVVD    153
                                                                    : : .  :

tr|A0A0C9MAJ6|A0A0C9MAJ6_9FUNG      VKDCVITVTPFANQFERQAILDAAELAGLNVLTLMHDETAVALNYAVNR--E-IG----K    185
sp|Q9Y4L1|HYOU1_HUMAN               IKDAVITVPVFFNQAERRAVLQAARMAGLKVLQLINDNTATALSYGVFRRKD-IN----T    223
sp|Q63617|HYOU1_RAT                 IKDAVITVPAFFNQAERRAVLQAARMAGLKVLQLINDNTATALSYGVFRRKD-IN----S    223
tr|A0A067CBX2|A0A067CBX2_SAPPC      VKDWVITVPMYFAEPQRQAIIDAAEIAGIRVLSLIDENTAAALHHGVDYEAP-EP----N    211
tr|D8RVG4|D8RVG4_SELML              VKDAVITVPPYFGQSERQAVIAAAQAAGINVLSLMNEHAAAALQYGIDKDF---S----T    210
tr|A0A1P8B8R9|A0A1P8B8R9_ARATH      VKDMVVSVPPYFGQAERRGLIQASQLAGVNVLSLVNEHSGAALQYGIDKDF---A----N    238
tr|A0A176WKM4|A0A176WKM4_MARPO      IKDAIITVPPYLGQAERQGILDAAQIAGITVLGLINEHSGAALQYGIDKDF---G----N    222
tr|A0A3R7K087|A0A3R7K087_9TRYP      LRHLVVAVSTHASLAQRQAVVDAAAVAGLRTVRLVHGTTAAAVQLAYLNADQFFAAAREN    286
tr|Q4E175|Q4E175_TRYCC              LRHLVVTVSAHASLAQRQTFVDAAAVAGLRAVRLVHGTTAAAVQLAHLNIEQFFSSAREK    292
tr|Q4E0G9|Q4E0G9_TRYCC              LRHLVVTVSAHASLAQRQTFVDAAAVAGLRAVRLVHGTTAAAVQLAHLNIEQFFSSAREQ    292
tr|A0A6A5CDJ2|A0A6A5CDJ2_NAEFO      VTDAIVSIPHFFTRAQRQAILDSASIANIKILALMHDHTATALQYGIKSAK--IIKELTK    232
tr|A0A0S4IUZ2|A0A0S4IUZ2_BODSA      VRDAILVVPRGYTSRQRQAMVDAAAITGLRVLSFIHPTTAAALQLGLQNRG--LG----D    228
tr|A0A0L1KXM1|A0A0L1KXM1_9EUGL      KQDAVLTVPHFFDMHQRRALFQSAALGNTNVIATIHTTTAIALQYGVQNRG--FG----S    228
tr|A0A0S4JJU2|A0A0S4JJU2_BODSA      PQEAVFTVPADWTMNQRQALVDAAGLTDLKVLSLMHSTTATALQYGMQKRG--FG----N    208
tr|A0A0N0VI62|A0A0N0VI62_LEPPY      PTGVVVTIPFHTSPVERRSILDAARFSGLKVLGLMHSTTAAAFYYGVRHRG--FG----N    214
tr|A4HND7|A4HND7_LEIBR              PNGIVVTIPFHTSPAARRAILDAARLSGLNVLGLMHSTTAAAFYYGVRHRG--FG----N    214
tr|A4IC10|A4IC10_LEIIN              PKGVVVTIPFHTSPAERRAILDAARLSGLSVLGLMHSTTAAAFYYGVRHRG--FG----N    214
tr|E9AFU8|E9AFU8_LEIMA              PKGVVVTIPFHTSPAERRAILNAARLSGLSVLGLMHSTTAAAFYYGVRHRG--FG----N    214
tr|A0A7G2C9U8|A0A7G2C9U8_9TRYP      PKNVVITVPFHATISERRSILLATHLADVSVLGFMHSTTAAAFYYGIRRRG--FD-----    118
tr|S9TX72|S9TX72_9TRYP              PKSVVLTLPFHSTLSERRSILEAAHFSDTRVLAYMHSTTAAAFYYGVRRRG--FE----G    211
tr|S9UBL2|S9UBL2_9TRYP              PKSVVLTLPFHSTLSERRSILEAAHFSDTRVLAYMHSTTAAAFYYGVRRRG--FE----G    211
tr|Q38E16|Q38E16_TRYB2              PQNVVITLPFHSSLGRRQTILEAARLVHINTLGLLHSTTATALYYGVRRRG--FG----N    217
tr|F9W5M9|F9W5M9_TRYCI              PESMVITLPYNSSFGKRQSVLEAARLVNLNVLGLMHSTTASALYYGIRRRG--FG----N    212
tr|A0A1X0P4V9|A0A1X0P4V9_9TRYP      PKNFVVTVPFHSSLGKRQIILEAARMVGLNVMGLMHSTTAAALYYGVRRRG--FE----N    207
tr|Q4CS27|Q4CS27_TRYCC              PKSVVITIPFHSSMGKRQAILEAARLVGMNVLGLMHSTTAAAFYYGIRRRG--LG----N    207
tr|Q4E4F4|Q4E4F4_TRYCC              PKSVVITIPFHSSMGKRQAILEAARLVGMNVLGLMHSTTAAAFYYGIRRRG--LG----N    207
tr|A0A422P1B1|A0A422P1B1_TRYRA      PKSVVITIPFHSSVGRRQTILEAARLVGANVLGLMHSTTAAALYYGIRRRG--LG----N    207
```

```
tr|A0A422N0X0|A0A422N0X0_9TRYP      PKSVVITIPFHSSLGRRQAILEAARLVGADVLGLMHSTTAAALYYGIRRRG--LG----N    207
                                             :. :     *: .. ::          :  :.  :. *.  .

tr|A0A0C9MAJ6|A0A0C9MAJ6_9FUNG      SAENHIFYDMGAGSTVASIVTFSN-VETKDG-----KISRSAPQLEVRG-VGFDRTLGGH    238
sp|Q9Y4L1|HYOU1_HUMAN               TAQNIMFYDMGSGSTVCTIVTYQM-VKTKEA-----G---MQPQLQIRG-VGFDRTLGGL    273
sp|Q63617|HYOU1_RAT                 TAQNIMFYDMGSGSTVCTIVTYQT-VKTKEA-----G---TQPQLQIRG-VGFDRTLGGL    273
tr|A0A067CBX2|A0A067CBX2_SAPPC      TPERIMLYNMGSTSLQVSIVEYKTRVVPDGF-----KKNKTIVEFDVLA-KAWDETLGGA    265
tr|D8RVG4|D8RVG4_SELML              EPRYVLFYDMGANSAFAAVVLFSS-YSAKEY-----GKNVSHNYFELKG-IRWDAEIGGQ    263
tr|A0A1P8B8R9|A0A1P8B8R9_ARATH      GSRHVIFYDMGSSSTYAALVYYSA-YSEKEY-----GKTVSVNQFQVKD-VRWDLGLGGQ    291
tr|A0A176WKM4|A0A176WKM4_MARPO      ETKHVVFYDMGANSLYAAVVYYSA-YSGKER-----GKTTSFNQFQVKG-IRWDATLGGQ    275
tr|A0A3R7K087|A0A3R7K087_9TRYP      ASKYVMVYDMGSRGTEVAVAYAFAQSLQ-------------QPGTITLLA-AVVHDTLGGR    332
tr|Q4E175|Q4E175_TRYCC              GPKYIMVYDMGGRRTEVAVYEFAPSRQ-------------RLGTITLRT-AIVNNTLGGR    338
tr|Q4E0G9|Q4E0G9_TRYCC              GPKYIMVYDMGGRRTEVAVYEFAPSRQ-------------RLGTITLRT-AIVNNTLGGR    338
tr|A0A6A5CDJ2|A0A6A5CDJ2_NAEFO      PRHVALFYDIGSVSTTVSIAEYTQSK-----------EKTSLGNIKILG-FASDENLGGS    280
tr|A0A0S4IUZ2|A0A0S4IUZ2_BODSA      EPQHVLIYDMGSTKTEAAVYRFDPIATPAAAAKGGKKSTNSFGTISLVGSIASDMTLGGR    288
tr|A0A0L1KXM1|A0A0L1KXM1_9EUGL      MTKNVLILDMGASKTEVGIYTFFTPAAD-GA------KRSESLGTLTTRA-IVTDPFFGGR    280
tr|A0A0S4JJU2|A0A0S4JJU2_BODSA      ETVNVVIYDMGSTKTEVGVYRYSPPEERKDG--KKVKVAVSLGHIETLH-IEVDSTLGGR    265
tr|A0A0N0VI62|A0A0N0VI62_LEPPY      NTVKLLVFDLGATHTEVGVYEISPAPR-RP------PLENAFGTLRTLG-VVEDRSLGGR    266
tr|A4HND7|A4HND7_LEIBR              NSLKMVVFDLGSTHTEVGVYEFLPAAK-RA------PFSSAFGVLRTLG-VVEDRSLGGR    266
tr|A4IC10|A4IC10_LEIIN              NTLRVVVFDLGSTHTEVGVYEFLPAAQ-KA------PLSSAFGTLRTLG-VVEDRSLGGR    266
tr|E9AFU8|E9AFU8_LEIMA              NTLKVVVFDLGSTHTEVGVYEYLPPAP-KA------PLSSAFGTLRTLG-VVEDRSLGGR    266
tr|A0A7G2C9U8|A0A7G2C9U8_9TRYP      KEVNIIVFDVGATHTEAGVYSLSPPGP-NA------TFGDKLGTLKTIK-VLSDKTLGGR    170
tr|S9TX72|S9TX72_9TRYP              RTVNLVIFDVGSTHSEIGVYTFSPPKN-NSA------FSEKLGTLSTRT-VLVDKTLGGR    263
tr|S9UBL2|S9UBL2_9TRYP              RTVNLVIFDVGSTHSEIGVYTFSPPKN-NSA------FSEKLGTLSTRT-VLVDKTLGGR    263
tr|Q38E16|Q38E16_TRYB2              RTVHLLVYDIGSTHTEVGVYKFSPPVQ-EQG--KRTKNVESFGTLTTMG-IVSDATLGGR    273
tr|F9W5M9|F9W5M9_TRYCI              RTVNVLIYDIGSTHTEVGVFKFSPPVQ-QPG--KRVRNVDSFGTLTTLA-IVADPTLGGR    268
tr|A0A1X0P4V9|A0A1X0P4V9_9TRYP      KTINLLIYDIGSTHTEVGIYKFSPPVQ-EGG--KKIKSADSFGTLTTMS-VVDDNFLGGR    263
tr|Q4CS27|Q4CS27_TRYCC              KTMNLLVYDIGSTHTEVGIYKFSPPVA-QSG--KKIKNADSFGTLTTMA-VVDDTFLGGR    263
tr|Q4E4F4|Q4E4F4_TRYCC              KTMNLLVYDIGSTHTEVGIYKFSPPVA-QSG--KKIKNADSFGTLTTMA-VVDDTFLGGR    263
tr|A0A422P1B1|A0A422P1B1_TRYRA      KTINLLVYDIGSTHTEVGIYKFSPPLV-QPG--KKIRNADSFGTLTTMA-VVDDTTLGGR    263
tr|A0A422N0X0|A0A422N0X0_9TRYP      KTINLLVYDIGSTHTEVGIYKFSPPVA-QPG--KKIRNADSFGTLTTMA-VVDDTMLGGR    263
                                             :. ::*.           :                          :        .  :**

tr|A0A0C9MAJ6|A0A0C9MAJ6_9FUNG      ELDVRLQQLLVAGFMKAHGG-----RVSTDIKTSSGAMTRLMKEANRVKQILSANTETMA    293
sp|Q9Y4L1|HYOU1_HUMAN               EMELRLRERLAGLFNEQRKG-----QRAKDVRENPRAMAKLLREANRLKTVLSANADHMA    328
sp|Q63617|HYOU1_RAT                 EMELRLREHLAKLFNEQRKG-----QKAKDVRENPRAMAKLLREANRLKTVLSANADHMA    328
tr|A0A067CBX2|A0A067CBX2_SAPPC      QFDLRLAEKFANEFNAKLTN-------GDDVRKIPRAMAKLRAAARKTKIVLSANEAIPV    318
tr|D8RVG4|D8RVG4_SELML              NLELRLVEHFAKEFEKKT---------GVDVRAFPKAVAKLKKQAKRAKEILSANTEASF    314
tr|A0A1P8B8R9|A0A1P8B8R9_ARATH      SMEMRLVEHFADEFNKQLGN-------GVDVRKFPKAMAKLKKQVKRTKEILSANTAAPI    344
tr|A0A176WKM4|A0A176WKM4_MARPO      TMEARLVDYFAKEFNAQVGM-------FFDVRKHPKAMAKLKKQVKRTKEILSANTEAPL    328
tr|A0A3R7K087|A0A3R7K087_9TRYP      AFDKCIARYIERELFPNARPKAIEPVLAASAPAARKAAASLLRAVKSARERLSVNQEAPV    392
tr|Q4E175|Q4E175_TRYCC              AFDRCIARYIEREHFPKARPKAIEPVLGVSSPAARKAAVSLMRAVQNARERLSVNQEAPV    398
tr|Q4E0G9|Q4E0G9_TRYCC              AFDRCIARYIEREHFPKARPKAIEPVLGVSKPAARKAAVSLMRAVQNARERLSVNQEAPV    398
tr|A0A6A5CDJ2|A0A6A5CDJ2_NAEFO      HFDNVLADFFAEKFIDKHKS---------DPRKEVRPLTRLVEESQKIKHILSANNDAYL    331
tr|A0A0S4IUZ2|A0A0S4IUZ2_BODSA      AVDACLAEFIENAYMTTAPTAGKQRVLTGTTPQSRKAVMSLLRAANKAKEMLSANREAPV    348
tr|A0A0L1KXM1|A0A0L1KXM1_9EUGL      IFDVAIAKMIAEDFHKK---TGIAVLGDTGSLDGRKGIAVLMRSANRAKEVLSSNKECPV    337
tr|A0A0S4JJU2|A0A0S4JJU2_BODSA      TFDACIARFLEEEIVKT---MKIPRIIGGTSLQQHKAMFSLIRAANGIKETLSANQAAPV    322
tr|A0A0N0VI62|A0A0N0VI62_LEPPY      AFDLCVAGIIENEAREK---LNIGSVLGGKTPAQLKSQFSLLRAANKVRETLSVNSVTPY    323
tr|A4HND7|A4HND7_LEIBR              AFDLCVARVIEKEARTK---LGIEPVLGGLSAVQLKSQFSLLRAANKVRETLSVNSNTPY    323
tr|A4IC10|A4IC10_LEIIN              AFDLCVARVIEAEARAK---LSIGPVLGGTTSAQLKSQFSLLRAANKVRETLSVNSNTPY    323
tr|E9AFU8|E9AFU8_LEIMA              AFDLCVARVIEAEARAK---LSIGPVLGGTTSTQLKSQFSLLRAANKVRETFSVNSNTPY    323
tr|A0A7G2C9U8|A0A7G2C9U8_9TRYP      AFDLCIAREMENEAVEK---LNISRVIGKKTAAELKSQYSLLRAANKVREVLSANTQTPY    227
tr|S9TX72|S9TX72_9TRYP              AFDLCIARVMEKEAVEK---MGIKPVIGGKTQAQLKSQYSLLRAANQVREILSVNSVTPF    320
tr|S9UBL2|S9UBL2_9TRYP              AFDLCIARVMEKEAVEK---MGIKPVIGGKTQAQLKSQYSLLRAANQVREILSVNSVTPF    320
tr|Q38E16|Q38E16_TRYB2              ALDSCIAKGIEAEAISK---MKISPVLGGSTVSQRKAQFSLFRAAKRAREVLSVNSKTPV    330
tr|F9W5M9|F9W5M9_TRYCI              TLDLCIAKGIEAEAMSK---IKISKVLGGTTIAQKKAQFSLFRAAKHAREVLSANSATPV    325
tr|A0A1X0P4V9|A0A1X0P4V9_9TRYP      AFDLCISRLLEKEAMAK---MKIGKVIGGKTVAERKSQFSLLRAAKKAREVLSANSKTPV    320
tr|Q4CS27|Q4CS27_TRYCC              AFDLCIAKIFEAEAMNK---MKISKVIGGKTIPERKSQFSLLRAAKKSREILSANSKTPV    320
tr|Q4E4F4|Q4E4F4_TRYCC              AFDLCIAKIFEAEAMNK---MKISKVIGGKTIPERKSQFSLLRAAKKSREILSANSKTPV    320
tr|A0A422P1B1|A0A422P1B1_TRYRA      AFDLCIAKLLETEAMNK---MKIPKVIGGKSIAERKSQFSLLRAAKKAREILSANSKTPV    320
tr|A0A422N0X0|A0A422N0X0_9TRYP      AFDLCIAKMLEVEAMNK---MKIPKVIGGKSIPERKSQFSLLRAAKKAREILSANSKTPV    320
                                    .:  :   :                           *        .  :  :* *

tr|A0A0C9MAJ6|A0A0C9MAJ6_9FUNG      SIEGLHE--GIDFKLKVTRAELEDICKDLIARVRNPLQTALEAA---NMTVDDIQSVVLV    348
sp|Q9Y4L1|HYOU1_HUMAN               QIEGLMD--DVDFKAKVTRVEFEELCADLFERVPGPVQQALQSA---EMSLDEIEQVILV    383
sp|Q63617|HYOU1_RAT                 QIEGLMD--DVDFKAKVTRVEFEELCADLFDRVPGPVQQALQSA---EMSLDQIEQVILV    383
tr|A0A067CBX2|A0A067CBX2_SAPPC      IVPSLHA--DLDYKGHASRAELEEIAADLFARVLDPVKSALDQA---GLTVGDLSAVEII    373
tr|D8RVG4|D8RVG4_SELML              FVDSLYD--DQDFKSHITRQAFEELCSDLWERAVIPLQKVLEDV---GMTSEQLYAVELL    369
tr|A0A1P8B8R9|A0A1P8B8R9_ARATH      SVESLHD--DRDFRSTITREKFEELCKDLWERSLTPLKDVLKHS---GLKIDDISAVELI    399
tr|A0A176WKM4|A0A176WKM4_MARPO      SVEALHD--DRDFRSTITRKKFEELCSDLWERALIPLKDVLADA---GLKVEDVNAVELI    383
tr|A0A3R7K087|A0A3R7K087_9TRYP      VVQGVQEG-GGDFTATISRARFAQECAHLFDEAARLRDAAIAQTRGVVPSLRDLARFEVI    451
tr|Q4E175|Q4E175_TRYCC              VVQGVQEGDSGDFTTIISRAVFEQECAHLFDEAVRLRDEAIAQTKGIVSSVRELTRFEMI    458
tr|Q4E0G9|Q4E0G9_TRYCC              VVQGVQEGDSGDFTTIISRAVFEQECAHLFDEAVRLRDEAIAQTKGIVSSVRELTRFEVI    458
tr|A0A6A5CDJ2|A0A6A5CDJ2_NAEFO      SIENLYN--DRDLSIKITREEFEKLSEQLLNRLTKPITAALSVA---NLTLDDIDAFEMS    386
tr|A0A0S4IUZ2|A0A0S4IUZ2_BODSA      TVEGIAQ--DSDFSLRISRATLERECGGIWDRAVRVRDAALKSA---NVTLAQLSRFEVV    403
tr|A0A0L1KXM1|A0A0L1KXM1_9EUGL      TVEGITK--GRDHSTIVRRADFEAHCASLFAKVSTLVKSAIDQS---GLDLSEIHAVEVH    392
tr|A0A0S4JJU2|A0A0S4JJU2_BODSA      TVEGVAP--DRDFSGKVTREQFETECSALFERAVEIAKKAIDKS---GLALSDLSSFELM    377
tr|A0A0N0VI62|A0A0N0VI62_LEPPY      TVEGIVP--DRDFHSSMTRATFESGCSDIFDRVKDLAVNVTKSV---NISLSDLTAFEMM    378
tr|A4HND7|A4HND7_LEIBR              TVEGIAP--DRDFHSSMTRATFESECDPLFQRVKGLAANVASVV---NISFEEINSFEMM    378
tr|A4IC10|A4IC10_LEIIN              TVEGIAP--DRDYHSSMTRATFEAECASLFQRVKDLATSVTKVV---NISFKDLNSFEMM    378
tr|E9AFU8|E9AFU8_LEIMA              TVEGIAP--DRDYHSSMTRATFESECAPLFQRVKDLAMSVTDVV---NISFKDLNSFEMM    378
tr|A0A7G2C9U8|A0A7G2C9U8_9TRYP      TVEGISR--DKDFSSTFTREQFERACGGLFDRVKALATEVIKES---GIPVKKLDAFEMM    282
tr|S9TX72|S9TX72_9TRYP              TVEGIAP--DRDFDSHFSREQFEAECGHLFTRVTKLASDALAAS---GLSKSEIDAFEMM    375
tr|S9UBL2|S9UBL2_9TRYP              TVEGIAP--DRDFDSHFSREQFEAECGHLFTRVTKLASDALAAS---GLSKSEIDAFEMM    375
tr|Q38E16|Q38E16_TRYB2              TVEGIAP--DRDFSTEVSRKEFETNCSELLKRFPRVAQEAVTKS---SLTLKDIDAFEMM    385
tr|F9W5M9|F9W5M9_TRYCI              TVEGIAP--ERDFNTVISRKDFEANCSELFKRFPVVAREALNKS---SLSLGDIDAFEMM    380
tr|A0A1X0P4V9|A0A1X0P4V9_9TRYP      TVEGIVP--DRDFHTEITRKDFEEKCSELFERVPKVAEEALKKS---GLDLKDIDAFEMM    375
tr|Q4CS27|Q4CS27_TRYCC              TIEGIVP--ERDFSTEISRKDLEEKCSHLFERVPKLAEEALSKA---GLSLSDIDAFEMM    375
```

```
tr|Q4E4F4|Q4E4F4_TRYCC              TIEGIVP--ERDFSTEISRKDLEEKCSHLFERVPKLAEEALSKA---GLSLSDIDAFEMM    375
tr|A0A422P1B1|A0A422P1B1_TRYRA      TVEGIVP--DRDFNTQISRKDLEEKCSYLFERVPKVAEEAVSKS---GLSLKDIDAFEMM    375
tr|A0A422N0X0|A0A422N0X0_9TRYP      TVEGIVP--DRNFNTQISRKDLEVKCAYLFERVPKVVEEALSKS---GLSLKEIDAFEMM    375
                                    :  :      :         *  :   .  :  .    .                .:  . :

tr|A0A0C9MAJ6|A0A0C9MAJ6_9FUNG      GGGVRVPSVQKQLVDLVGAQ---KIAKNVNADEAAVLGAAFRGASLSNQFRLSKQISIKD    405
sp|Q9Y4L1|HYOU1_HUMAN               GGATRVPRVQEVLLKAVGKE---ELGKNINADEAAAMGAVYQAAALSKAFKVKP-FVVRD    439
sp|Q63617|HYOU1_RAT                 GGPTRVPKVQEVLLKPVGKE---ELGKNINADEAAAMGAVYQAAALSKAFKVKP-FVVRD    439
tr|A0A067CBX2|A0A067CBX2_SAPPC      GGGVRIPKIQALLQAFVQ-R---DLGKRLNGDEAMALGAAFNAANLSTSFRV-RHVGMTD    428
tr|D8RVG4|D8RVG4_SELML              GGGTRVPKLQQVLAQALGKK---PLERHLDADEAITLGAALYAANISDGIKLNRKIGMFD    426
tr|A0A1P8B8R9|A0A1P8B8R9_ARATH      GGATRVPKLQSTIQEFIGKQ---QLDKHLDADEAIVLGSALHAANLSDGIKLKRRLGIVD    456
tr|A0A176WKM4|A0A176WKM4_MARPO      GGATRVPKLQAVLSDFLGKK---NLDRHLDADEAVVLGASLQAANLSDGFKLNRKLGMID    440
tr|A0A3R7K087|A0A3R7K087_9TRYP      GGATRMPKLLERLSDGYGR----AVDRTLNSDEATVVGAAYVGAARAG-IPVR-GFRVVE    505
tr|Q4E175|Q4E175_TRYCC              GGATRMPKLLERLSDGYGR----AVDRTLNSDEAAVVGASYMGAARAN-IPIR-GFAVVE    512
tr|Q4E0G9|Q4E0G9_TRYCC              GGATRMPKLLERLSDGYGR----AVDRTLNSDEAAVVGASYMGAARAN-IPIR-GFAVVE    512
tr|A0A6A5CDJ2|A0A6A5CDJ2_NAEFO      GGSSRIPAVQDRISKFVGSK--LMVGYSLNADEAIAMGSSFYGAMLSPSFKVT-AFKIFD    443
tr|A0A0S4IUZ2|A0A0S4IUZ2_BODSA      GGGLRVPIIIEKLSEGYNKAGSGVVDRTLNSDEAMVVGAGYYGAALSGHFRIK-GFRLVE    462
tr|A0A0L1KXM1|A0A0L1KXM1_9EUGL      GGAIRIPKLLDVLAEELGQ----PIQRTLNGDEAAVLGAAFHAARVSGTFGVK-GFAIQE    447
tr|A0A0S4JJU2|A0A0S4JJU2_BODSA      GGGVRPPKIVSDLSAFLGR----PVDRTLNGDEAAAFGAGYYGARLSGYFRVR-SFSIRD    432
tr|A0A0N0VI62|A0A0N0VI62_LEPPY      GGISRTPKIIADLSAFLGR----DVDRTMNMDEAAAMGAGYYAAKLSPLYHAK-SLKLDE    433
tr|A4HND7|A4HND7_LEIBR              GGLSRTPRIIADLSKVIGR----GVDRTMNMDEAAAIGAGYYAAKLSPLYHAK-SLKLDE    433
tr|A4IC10|A4IC10_LEIIN              GGLSRTPKIIADLSEAIGR----DVDRTMNMDEAAAIGAGYYAAKLSPLYRAK-SLKLDE    433
tr|E9AFU8|E9AFU8_LEIMA              GGLSRTPKIIADLSEVIGR----DVDRTMNMDEAAAIGAGYYAAKLSPLYRAK-SLKLDE    433
tr|A0A7G2C9U8|A0A7G2C9U8_9TRYP      GGVSRTPKIISDISAILKR----DVDRTMNMDEAAAIGSAYFAVKKSVHYKSK-SFRVDE    337
tr|S9TX72|S9TX72_9TRYP              GGASRTPKIITDVSVFLGK----DVDRTMNMDEAAAIGAAYYAVKISPHYRSK-SFKVQE    430
tr|S9UBL2|S9UBL2_9TRYP              GGASRTPKIITDVSVFLGK----DVDRTMNMDEAAAIGAAYYAVKISPHYRSK-SFKVQE    430
tr|Q38E16|Q38E16_TRYB2              GGASRTPKIISDLSTFWGK----EVNRTLNSDEAAAMGAAYYALRLSPHYRPH-SFRIIE    440
tr|F9W5M9|F9W5M9_TRYCI              GGTSRTPKIINDLSAFWGR----EVNRTLNSDEAAAMGAAYYALRLSPHYRPH-SFRIIE    435
tr|A0A1X0P4V9|A0A1X0P4V9_9TRYP      GGTSRTPKILADLSTFLGR----EVDRTLNSDESAAIGAAYYALRLSPFYRVR-SFRVVE    430
tr|Q4CS27|Q4CS27_TRYCC              GGTSRTPKIISDLSAMLGR----EVDRTLNSDEAAAVGAAYYALRLSPFYRVR-SFRVEE    430
tr|Q4E4F4|Q4E4F4_TRYCC              GGTSRTPKIISDLSAMLGR----EVDRTLNSDEAAAVGAAYYALRLSPFYRVR-SFRVEE    430
tr|A0A422P1B1|A0A422P1B1_TRYRA      GGTSRTPKIIAELSALLGR----EVDRTLNSDEAAAMGAAYHALRLSSFYRVR-SFRLEE    430
tr|A0A422N0X0|A0A422N0X0_9TRYP      GGTSRTPKIISDLSAMLGR----EVDRTLNSDEAAALGAAYYALRLSSFYRVR-SFRVEE    430
                                    **   * * :    :                 :   :: **: ..*:    .    :             . : :

tr|A0A0C9MAJ6|A0A0C9MAJ6_9FUNG      VTLFPIEVTYKPENKG-K---EAKTAVD---------TTLFNKFDAIGTRKIMTFNRV-N    451
sp|Q9Y4L1|HYOU1_HUMAN               AVVYPILVEFTREVEE-EPGIHSLKHNK---------RVLFSRMGPYPQRKVITFNRYSH    489
sp|Q63617|HYOU1_RAT                 AVIYPILVEFTREVEE-EPGLRSLKHNK---------RVLFSRMGPYPQRKVITFNRYSH    489
tr|A0A067CBX2|A0A067CBX2_SAPPC      IASFPVGVRLVDLASH------AADDDHPEAKHWVKRAALFGERQKLNVKKSVSFSH-AD    481
tr|D8RVG4|D8RVG4_SELML              GASYGVVLKVNDDGP------------YEL---------VVPRLRRIPVKLARAVKQQ-E    464
tr|A0A1P8B8R9|A0A1P8B8R9_ARATH      GSPYGFLVELEGPNVK------KDESTKQQ---------LVPRMKKLPSKMFRSFVLD-K    500
tr|A0A176WKM4|A0A176WKM4_MARPO      GASYSILMELDGASPG------ALENVDNV---------LVHRLKKIPSKAYRALKNQKS    485
tr|A0A3R7K087|A0A3R7K087_9TRYP      PLMNDVYFSLTPP--LQGGGG-----------AGTRHLVFAKARTLVPAVQSLRFKNRTA    552
tr|Q4E175|Q4E175_TRYCC              PLMNDVWFSLTPS--LQDPRDAAAPP-----RDSTRHLIFPKGSTILPAVRSLQLKNRTA    565
tr|Q4E0G9|Q4E0G9_TRYCC              PLMNDVWFSLTPL--LQDPRDAAAPP-----RNSTRHLIFPKGSTILPAVRSLQLKNRTA    565
tr|A0A6A5CDJ2|A0A6A5CDJ2_NAEFO      TVPHKVDFLLSGND--KE-------------------INLFNHKDEIDAKKTITVPRT-E    481
tr|A0A0S4IUZ2|A0A0S4IUZ2_BODSA      RLGFPVWITVALNSSSNTEATGTAEA-----PSSQQRKLLFPSEARLPTRLRVRVNATVS    517
tr|A0A0L1KXM1|A0A0L1KXM1_9EUGL      TIPYNITFSISPK--ASDKGLGTTED-----APVKSRGLFAN--ASWPAKKSVSVNRT-E    497
tr|A0A0S4JJU2|A0A0S4JJU2_BODSA      FFPQSIHFKIVSA--DGT--------------SSAIRPLFVK--SPFGARKSITVNRT-E    473
tr|A0A0N0VI62|A0A0N0VI62_LEPPY      TIPYSVEFEVHPR--LNKK------------KPLTRRPLFGTDGVLLGDAVSLTFNRT-D    478
tr|A4HND7|A4HND7_LEIBR              TIPYRIDFEVDPP--LSKE------------KPAARRPLFSVADFLLGDSVSLTFNRT-E    478
tr|A4IC10|A4IC10_LEIIN              TIPYGVDFEVEPP--LSKE------------KPAVRRPLFGADGFLLGDSVSLTFNRT-E    478
tr|E9AFU8|E9AFU8_LEIMA              TIPYRVDFEVEPP--LSKE------------KPAVRRPVFSADGFLLGDSVSITFNRT-E    478
tr|A0A7G2C9U8|A0A7G2C9U8_9TRYP      RVPYPVYFSLST-----NT------------SNSSRRLLLSS-DELIGQSFSITFEAD-T    378
tr|S9TX72|S9TX72_9TRYP              RVPFPLYFSVYPA--FHSN------------KPSARRMLLHS-DAFIGDAVSITFNRT-D    474
tr|S9UBL2|S9UBL2_9TRYP              RVPFPLYFSVYPA--FHSN------------KPSARRMLLHS-DAFIGDAVSITFNRT-D    474
tr|Q38E16|Q38E16_TRYB2              RVPYTFLFAVSPE--VKNS------------SNSKR--MLAN-NPIIGSRWSITVNRT-D    482
tr|F9W5M9|F9W5M9_TRYCI              RIPSTLSFVVNPR--CYEL-----------------------------------------    452
tr|A0A1X0P4V9|A0A1X0P4V9_9TRYP      HIPFKFFFTITPS--LNNS------------SVSKR--VLVE-QPIIGSRKSITLNRT-D    472
tr|Q4CS27|Q4CS27_TRYCC              HIPFVFFFSITSS--SKNS------------SQSRR--LLAE-NPALGMRQSITLNRT-E    472
tr|Q4E4F4|Q4E4F4_TRYCC              HIPFVFFFSITSS--SKNS------------SQSRR--LLAE-NPALGMRQSITLNRT-E    472
tr|A0A422P1B1|A0A422P1B1_TRYRA      HIPFVFYFRITPF--VKAS------------APSLR--LLAE-NPVLGMRRSVTLNRT-E    472
tr|A0A422N0X0|A0A422N0X0_9TRYP      HIPFVFSFCVTPF--LKTS------------SPSPR--LLVE-NPALGMRRSITLNRT-E    472
                                                  . .

tr|A0A0C9MAJ6|A0A0C9MAJ6_9FUNG      DFEFDLSYGKNAAADVDG--------IAKVKVTGLTEAMKKHKDDIKAS-----------    492
sp|Q9Y4L1|HYOU1_HUMAN               DFNFHINYGDLGFLGPEDLRVFGSQNLTTVKLKGVGDSFKKYPDYE-S------------    536
sp|Q63617|HYOU1_RAT                 DFNFHINYGDLGFLGPEDLRVFGSQNLTTVKLKGVGESFKKYPDYE-S------------    536
tr|A0A067CBX2|A0A067CBX2_SAPPC      DVSCTFRYDKPSALPLGVS-----SIISRYNITGIEAFAKKMEDKKLG------------    524
tr|D8RVG4|D8RVG4_SELML              DYELSLHYDPEGELPLGII----DREIAAFKISGVTDSVAKYSSYNLS------------    508
tr|A0A1P8B8R9|A0A1P8B8R9_ARATH      DFDVSLAYESEGILPPGTT----SPVFAQYSVSGLADASEKYSSRNLS------------    544
tr|A0A176WKM4|A0A176WKM4_MARPO      DFKVALKYD-ATLLPPGVI----SPEVATYQITGVADAANKYASYNVS------------    528
tr|A0A3R7K087|A0A3R7K087_9TRYP      DFTLTLEDDGGRF-------------ARATSIHGVGESISAA--------------RGRR    585
tr|Q4E175|Q4E175_TRYCC              DFTLTLEDESGHF-------------TRATFIRGVNESIMAAQLLPQSFRASDGAGKKRR    612
tr|Q4E0G9|Q4E0G9_TRYCC              DFTLTLEDESGHF-------------TRATFIRGVNESIIAAQLLPQSLRASDGAGKKRK    612
tr|A0A6A5CDJ2|A0A6A5CDJ2_NAEFO      DFTITLKYASEYL-PSGYST--EHAAFATYEITGVTKAMSNWTFEDE---------R---    526
tr|A0A0S4IUZ2|A0A0S4IUZ2_BODSA      DFIVTLDETTNSS----SSA--TIVSKSNTLVSGVLGALQQVGYGDVDAAD-----R---    563
tr|A0A0L1KXM1|A0A0L1KXM1_9EUGL      NFEMTFSDQNG--------------PLQVLKVSGLHDSLEKLGHFSPER-D-----P---    534
tr|A0A0S4JJU2|A0A0S4JJU2_BODSA      DFTIALLSTVDGE----SF-----TPLMSVNVSGVKSALEGLNYFNPTIVH-----E---    516
tr|A0A0N0VI62|A0A0N0VI62_LEPPY      DFSVDLFAGTD------AK-----LPIATIEVTDVKKALTNLGALSPTVQH-----P---    519
tr|A4HND7|A4HND7_LEIBR              DFSINFYTEGD------AT-----RPFVSVTVTGVKQALTRMNSLPSVVKH-----A---    519
tr|A4IC10|A4IC10_LEIIN              DFSLNFYIDDD------AA-----RPFVRVTVTGVKQALTDMKALSPAVEH-----A---    519
tr|E9AFU8|E9AFU8_LEIMA              DFSLNFYSD-D------AA-----TPFVCITVTGVKQALTDMNALSPVVKH-----A---    518
tr|A0A7G2C9U8|A0A7G2C9U8_9TRYP      NFELDFYDSDS------AP-----QPFLKETFSSVES---LLKLYEPSVQH-----E---    416
tr|S9TX72|S9TX72_9TRYP              DFTVRLYQSET------TE-----EPFAEVRVEGVKKKLDVMNFFEPLIAH-----E---    515
tr|S9UBL2|S9UBL2_9TRYP              DFTVRLYQSET------TE-----EPFAEVRVEGVKKKLDVMNFFEPLIAH-----E---    515
tr|Q38E16|Q38E16_TRYB2              DFSIQLFDSDA------R--------IGNVNITGVKGALERLGFFQPKLNH-----T---    520
tr|F9W5M9|F9W5M9_TRYCI              ------------------------------------------------------------    452
```

```
tr|A0A1X0P4V9|A0A1X0P4V9_9TRYP      DFSIQLFDSQD------F--------VGEISVTGVKEALKLLQEKKSDFDH-----P---    510
tr|Q4CS27|Q4CS27_TRYCC              DFTVEIFESNN------C--------VVKIQVMGVKTTLERLGFFAPTIVH-----P---    510
tr|Q4E4F4|Q4E4F4_TRYCC              DFTVEIFESNN------C--------VVKIQVIGVKTTLERLGFFAPTIVH-----P---    510
tr|A0A422P1B1|A0A422P1B1_TRYRA      DFVIELFESNN------F--------VGKITVSGVKTALDQLGYFEPAIVH-----P---    510
tr|A0A422N0X0|A0A422N0X0_9TRYP      DFAIELFESNT------I--------VGKITVIGVKAALEQLGYFTPTILH-----P---    510


tr|A0A0C9MAJ6|A0A0C9MAJ6_9FUNG      ----EIPPKVRVSIELSNSGLVSVPEATLQI------------GKLTFK---EKVKSFFG    533
sp|Q9Y4L1|HYOU1_HUMAN               -------KGIKAHFNLDESGVLSLDRVESVFETLVEDSAEEESTLTKLG---NTISSLFG    586
sp|Q63617|HYOU1_RAT                 -------KGIKAHFNLDESGVLSLDRVESVFETLVEDSPEEESTLTKLG---NTISSLFG    586
tr|A0A067CBX2|A0A067CBX2_SAPPC      -----EP-KVTLSFNLDASGLASIVKAEATLEEEIEVPAPTKKAAKKADD--SSSSSAD-    575
tr|D8RVG4|D8RVG4_SELML              -----APIKSVLHFALSRSGVLSLDRAETVVEFTELVEVPVNVSTNATTLESTSNASV-    562
tr|A0A1P8B8R9|A0A1P8B8R9_ARATH      -----APIKANLHFSLSRSGILSLDRGDAVIEITEWVDVPKKNVTIDSNTTTSTGNATD-    598
tr|A0A176WKM4|A0A176WKM4_MARPO      -----APLKTSLHFSLSRSGVISLDKAETVVEISEWYEVPVANATTDVNATVEAKSTEEG    583
tr|A0A3R7K087|A0A3R7K087_9TRYP      GGMSLEHTEVVVEVTVAESGIPFVSNAYLRATYVKRTAAARKGQTGSANTTRNEFFAA--    643
tr|Q4E175|Q4E175_TRYCC              ERLSLEHTEVMLEVTATESGIPFVSNAYLHATYLKVTVAKEPTD-----TVQNESNAG--    665
tr|Q4E0G9|Q4E0G9_TRYCC              ERLSLEHTEVMLEVTATESGIPFVSNAYLHATYLKVTVAKEPTD-----TVQNESNAG--    665
tr|A0A6A5CDJ2|A0A6A5CDJ2_NAEFO      ----GKKKKVKATFKVNKFGLVELDSVNAILEETITVR--VESNNT-----------T--    567
tr|A0A0S4IUZ2|A0A0S4IUZ2_BODSA      ----NITHTVWVDIRASEAAMVYIEKAEIEIHYFVNASRKVRKLVT-----------P--    606
tr|A0A0L1KXM1|A0A0L1KXM1_9EUGL      ----QNSCSVRIEAKLDGNGVLSASSVKL--KYNEWKVKSSESDES-----------S--    575
tr|A0A0S4JJU2|A0A0S4JJU2_BODSA      ----NNTHIVRVELKLSENGVVVVEDSEVRVRLAANVTKKLKTRKN-----------S--    559
tr|A0A0N0VI62|A0A0N0VI62_LEPPY      ----NNSHMIRMQVIMNETGLVQLEHTEALVRYAAQVTEKTKENVT-----------D--    562
tr|A4HND7|A4HND7_LEIBR              ----NNSHMIRLQIVLNETGLVQVEYAEAVVRYAEEVTQNTRENVT-----------G--    562
tr|A4IC10|A4IC10_LEIIN              ----NNSHMVRLQIVLNETGLVQVEHAEVVVRYAEEVTQKVTENVT-----------D--    562
tr|E9AFU8|E9AFU8_LEIMA              ----NNSHMVRLQIVLNETGLVEVEHAEVVVRYAEEVTQKVTENVT-----------D--    561
tr|A0A7G2C9U8|A0A7G2C9U8_9TRYP      ----NNTHMVRIQMRVNSTGLPQIEEADIIFRRVENATKHGKSGNE-----------T--    459
tr|S9TX72|S9TX72_9TRYP              ----NNSHLVRLQLLINETGLPMVDASDVVYRYVTNVTERHSVTDA-----------A--    558
tr|S9UBL2|S9UBL2_9TRYP              ----NNSHLVRLQLLINETGLPMVDASDVVYRYVTNVTERHSVTDA-----------A--    558
tr|Q38E16|Q38E16_TRYB2              ----NNSHIIRIQVRFNESGLLEVEEAGVFYRYAVNVSSSTKTTGV-----------Q--    563
tr|F9W5M9|F9W5M9_TRYCI              ------------------------------------------------------------    452
tr|A0A1X0P4V9|A0A1X0P4V9_9TRYP      ----NNTHIVRIQLLLNESGLFEVEEADVIIRYAVNVSVKVKVNTT-----------D--    553
tr|Q4CS27|Q4CS27_TRYCC              ----NNSHIVRVQLRINDSGLFEVDEADVIYRYAANVSSKIKLNTT-----------T--    553
tr|Q4E4F4|Q4E4F4_TRYCC              ----NNSHIVRVQLRINDSGLFEVDEADVIYRYAANVSSKIKLNTT-----------T--    553
tr|A0A422P1B1|A0A422P1B1_TRYRA      ----NNTHIVRVQLRLNDSGLFEVEEAEVIYRYAANVSRKIRVNAT-----------S--    553
tr|A0A422N0X0|A0A422N0X0_9TRYP      ----NNTHVVRVQLRLNDSGLFEVEEADAIYRYAANVSRKIKINAT-----------S--    553


tr|A0A0C9MAJ6|A0A0C9MAJ6_9FUNG      GKDSKDAEADKKNEDANQKNETAKT-----------------------------------    558
sp|Q9Y4L1|HYOU1_HUMAN               GGTTPDAKENGTDT--VQEEEESPAEGSKDEP--------------GE------------    618
sp|Q63617|HYOU1_RAT                 GGTSSDAKENGTDA--VQEEEESPAEGSKDEP--------------AE------------    618
tr|A0A067CBX2|A0A067CBX2_SAPPC      -----DDKD---------------------------------------------------    579
tr|D8RVG4|D8RVG4_SELML              ------------------------------------------------------------    562
tr|A0A1P8B8R9|A0A1P8B8R9_ARATH      ----ENSQENKEDL---Q------T-----------------------------------    610
tr|A0A176WKM4|A0A176WKM4_MARPO      AGSPEEAKDEEVKL---SSEEVKQT-----------------------------------    605
tr|A0A3R7K087|A0A3R7K087_9TRYP      ----SAAEENATDTEASEEAQKHPTPQRPGEQAEEAGAGPTLPPPPPQPPQKQQQPAGGA    699
tr|Q4E175|Q4E175_TRYCC              ----SAAGENATDAEAPNGTEKSSKPQQPDEYP-------------GEEQEEQQQRDLGA    708
tr|Q4E0G9|Q4E0G9_TRYCC              ----SAAGENATDAEAPNGTEKSSKLQQPDEYP-------------GEEQEGQQQRDLGG    708
tr|A0A6A5CDJ2|A0A6A5CDJ2_NAEFO      ----T----------------------------------------SNE------------    571
tr|A0A0S4IUZ2|A0A0S4IUZ2_BODSA      ----PAEEKNASDGDASASPATEKTT-----------------PESGEEQPSTAEDAGGD    645
tr|A0A0L1KXM1|A0A0L1KXM1_9EUGL      ------------------------------------------------------------    575
tr|A0A0S4JJU2|A0A0S4JJU2_BODSA      ----SEEIPDQEK-----------TN-----------------DDSNE------------    575
tr|A0A0N0VI62|A0A0N0VI62_LEPPY      ----ST---------------------------------------TGE------------    567
tr|A4HND7|A4HND7_LEIBR              ----EQ---------------------------------------SSE------------    567
tr|A4IC10|A4IC10_LEIIN              ----KA---------------------------------------SGE------------    567
tr|E9AFU8|E9AFU8_LEIMA              ----KA---------------------------------------SGE------------    566
tr|A0A7G2C9U8|A0A7G2C9U8_9TRYP      ------------------------------------------------------------    459
tr|S9TX72|S9TX72_9TRYP              ----DEEE------------------------------------AHNE------------    566
tr|S9UBL2|S9UBL2_9TRYP              ----DEED------------------------------------AHNE------------    566
tr|Q38E16|Q38E16_TRYB2              ----ADS-----------------------------------------------------    566
tr|F9W5M9|F9W5M9_TRYCI              ------------------------------------------------------------    452
tr|A0A1X0P4V9|A0A1X0P4V9_9TRYP      ----ELPNT--T------------------------------------------------    559
tr|Q4CS27|Q4CS27_TRYCC              ----KDLNT--SD-----------IT-----------------NTSNI------------    567
tr|Q4E4F4|Q4E4F4_TRYCC              ----NDLNT--SD-----------IT-----------------NTSNI------------    567
tr|A0A422P1B1|A0A422P1B1_TRYRA      ----NDLND--TD-----------N-----------------------------------    561
tr|A0A422N0X0|A0A422N0X0_9TRYP      ----N---D--TD-----------N-----------------------------------    558


tr|A0A0C9MAJ6|A0A0C9MAJ6_9FUNG      ---------------------------------DG--------------TADSKNKTAAE    571
sp|Q9Y4L1|HYOU1_HUMAN               ----QVELKEEA----------------EAPVEDGSQPPPPEPKG----DATPEGEKATE    654
sp|Q63617|HYOU1_RAT                 ----QGELKEEA----------------EAPMEDTSQPPPSEPKG----DAAREGETPDE    654
tr|A0A067CBX2|A0A067CBX2_SAPPC      --------------------------------DDKTT--------------------PEN    587
tr|D8RVG4|D8RVG4_SELML              ------------------------------------------------------AE-NVT    567
tr|A0A1P8B8R9|A0A1P8B8R9_ARATH      --------------------------------DAENS--------------TASNTTAEE    624
tr|A0A176WKM4|A0A176WKM4_MARPO      --------------------------------QADSSPEAGD-VS----GVSSEGEPSKD    628
tr|A0A3R7K087|A0A3R7K087_9TRYP      G-----AKWEEAPEAAVEAEAEEEVEEEEEAEVAQSNPEE-GQAGERGDNGTGEGAVPGV    753
tr|Q4E175|Q4E175_TRYCC              E-----EKWDERQEKAAEEEGEEERKENHNVS---------DQASEKD-SNASEATAPRA    753
tr|Q4E0G9|Q4E0G9_TRYCC              E-----EKWDERQEKAVEEEGEEERKENYNVS---------DQSSEKE-SNTSEATAPRA    753
tr|A0A6A5CDJ2|A0A6A5CDJ2_NAEFO      -----------T----------------------------------------------KT    574
tr|A0A0S4IUZ2|A0A0S4IUZ2_BODSA      SASDEGEISDEAPDATTTS--------------SSSKPSASDDITEGAKKASKDSTAPKK    691
tr|A0A0L1KXM1|A0A0L1KXM1_9EUGL      ------------------------------------------------------------    575
tr|A0A0S4JJU2|A0A0S4JJU2_BODSA      -----------T-------------------------------------------AQATE    581
tr|A0A0N0VI62|A0A0N0VI62_LEPPY      -----------V------------------------------------------------    568
tr|A4HND7|A4HND7_LEIBR              -----------T------------------------------------------------    568
tr|A4IC10|A4IC10_LEIIN              -----------R------------------------------------------------    568
tr|E9AFU8|E9AFU8_LEIMA              -----------R------------------------------------------------    567
tr|A0A7G2C9U8|A0A7G2C9U8_9TRYP      ------------------------------------------------------------    459
tr|S9TX72|S9TX72_9TRYP              -----------S------------------------------------------------    567
tr|S9UBL2|S9UBL2_9TRYP              -----------S------------------------------------------------    567
```

```
tr|Q38E16|Q38E16_TRYB2                 ------------------------------------------------------------   566
tr|F9W5M9|F9W5M9_TRYCI                 ------------------------------------------------------------   452
tr|A0A1X0P4V9|A0A1X0P4V9_9TRYP         ------------------------------------------------------------   559
tr|Q4CS27|Q4CS27_TRYCC                 -----------T------------------------------------------------   568
tr|Q4E4F4|Q4E4F4_TRYCC                 -----------T------------------------------------------------   568
tr|A0A422P1B1|A0A422P1B1_TRYRA         -----------T------------------------------------------------   562
tr|A0A422N0X0|A0A422N0X0_9TRYP         -----------T------------------------------------------------   559

tr|A0A0C9MAJ6|A0A0C9MAJ6_9FUNG         EAK------ESVSIAKIPLTIEFIPTGLAP------------------------------   595
sp|Q9Y4L1|HYOU1_HUMAN                  KEN-----GDKSEAQKPSEKAEAGPEGVAPAPEGEKKQKPARKRRMVEEIGVELVVLDLP   709
sp|Q63617|HYOU1_RAT                    KES-----GDKSEAQKPNEKGQAGPEGVPPAPEEEKKQKPARKQKMVEEIGVELAVLDLP   709
tr|A0A067CBX2|A0A067CBX2_SAPPC         DEV-----PEMIKKTKVHRVTLTVVRSDAA-------------------HRP-EAGMAIL   622
tr|D8RVG4|D8RVG4_SELML                 QAP-----VETIMEKKLKKKTIRIPSKLEV-------------------LRD---T---G   597
tr|A0A1P8B8R9|A0A1P8B8R9_ARATH         PAV-----ASLGTEKKLKKRTFRIPLKVVE-------------------KTV-GPG---A   656
tr|A0A176WKM4|A0A176WKM4_MARPO         SPV-----AEPVVKKKLRRRTIRIPLKIKD-------------------LSK-GPN---M   660
tr|A0A3R7K087|A0A3R7K087_9TRYP         GSVSTAKREEEVMVVKRVVRGVPLRFA--P-----------------------APFSAGV   788
tr|Q4E175|Q4E175_TRYCC                 -----EDTSAEKEIVERVVRSFPLRLV--S-----------------------VPLSVGV   783
tr|Q4E0G9|Q4E0G9_TRYCC                 -----EDKSAEKEIVERVVRSFPLSLV--S-----------------------VPLSVGV   783
tr|A0A6A5CDJ2|A0A6A5CDJ2_NAEFO         EPV---Y--------KKDTRVTKVALDVKF-----------------------IDISVIK   600
tr|A0A0S4IUZ2|A0A0S4IUZ2_BODSA         KPT---YVTVTEVHVEARSRVFPLTLTTTF-----------------------AS--PAV   723
tr|A0A0L1KXM1|A0A0L1KXM1_9EUGL         -ST---EVS--EEISEPIKHDISLPMEWEY-----------------------IG--RRP   604
tr|A0A0S4JJU2|A0A0S4JJU2_BODSA         VSA---ELP--SVQIQMKKRSTSIKPTIEL-----------------------SQ--PSM   611
tr|A0A0N0VI62|A0A0N0VI62_LEPPY         KEV---DKT--HTVIRMRHKTAVVPAALIW-----------------------KN--PDA   598
tr|A4HND7|A4HND7_LEIBR                 KPV---EKS--VKVVKMRHQVVDLTATLAW-----------------------KN--PTE   598
tr|A4IC10|A4IC10_LEIIN                 KSV---EKS--VKVIKMRIRAADVTATLAW-----------------------KN--PTG   598
tr|E9AFU8|E9AFU8_LEIMA                 KSV---EKS--VKVIKMRSRAADVSATLAW-----------------------KN--PTG   597
tr|A0A7G2C9U8|A0A7G2C9U8_9TRYP         --------A--EPVLETRRHSFVLTSKPTW-----------------------YD--PAE   484
tr|S9TX72|S9TX72_9TRYP                 APH---VKY--VYKLKTRKRELHVPATLHW-----------------------FH--PRE   597
tr|S9UBL2|S9UBL2_9TRYP                 APH---VKY--VYKLKTRKRELHVPATLHW-----------------------FH--PRE   597
tr|Q38E16|Q38E16_TRYB2                 NSS---NES--GYVVEMKRKSARLSARVTF-----------------------TN--PEP   596
tr|F9W5M9|F9W5M9_TRYCI                 ------------------------------------------------------------   452
tr|A0A1X0P4V9|A0A1X0P4V9_9TRYP         ------EEV--KQVVKMKRETMSLSTKVRF-----------------------VN--PSL   586
tr|Q4CS27|Q4CS27_TRYCC                 NTS---YRT--VHNIKMKRTSATVATTVSF-----------------------TN--PSP   598
tr|Q4E4F4|Q4E4F4_TRYCC                 NTS---YRT--VHNIKMKRTSATVATTVSF-----------------------TN--PSP   598
tr|A0A422P1B1|A0A422P1B1_TRYRA         NTS---FQR--VYNIHMKRKSLLISTTVSF-----------------------TN--PSP   592
tr|A0A422N0X0|A0A422N0X0_9TRYP         NTS---YQK--VYSIIMKRKSLPISTAVSF-----------------------TN--PSP   589

tr|A0A0C9MAJ6|A0A0C9MAJ6_9FUNG         -LSEKDKAIAKKRIAELDALDSRKKLREESRNALETFVYKVQ--DFLYNDI--VALVATE   650
sp|Q9Y4L1|HYOU1_HUMAN                  DLPEDKLAQSVQKLQDLTLRDLEKQEREKAANSLEAFIFETQ--DKLYQPE--YQEVSTE   765
sp|Q63617|HYOU1_RAT                    DLPEDELAHSVQKLEDLTLRDLEKQEREKAANSLEAFIFETQ--DKLYQPE--YQEVSTE   765
tr|A0A067CBX2|A0A067CBX2_SAPPC         PMTFRDKKESIAMLAEMDKVDNIRKANAEAKNRLEAFVYESR--EYLSSHDDVLEKVTTT   680
tr|D8RVG4|D8RVG4_SELML                 AYSEDQVAQFRSRLDRLKELDDEKRQTEEAKNNLESYIYTTK--EMLDTIKD-LSKVSTE   654
tr|A0A1P8B8R9|A0A1P8B8R9_ARATH         PFSKESLAEAKIKLEALDKKDRERRRTAELKNNLESYIYATK--EKLETPE--FEKISTQ   712
tr|A0A176WKM4|A0A176WKM4_MARPO         RLSEKDIKEAVARMEKLHLKDAEKRKTAEAKNSLEAYIYATK--DKLESLEG-IETVSTE   717
tr|A0A3R7K087|A0A3R7K087_9TRYP         NLNKTETMAARDRLRAFQRIDDERLMRSAIRNDLEALILHYKSLDAWDKAT-----VG-S   842
tr|Q4E175|Q4E175_TRYCC                 NLNKLEAAAARDRLYAFQRVDDERLMRSALRNDIETLILHYKSLDAWDKAV------E-N   836
tr|Q4E0G9|Q4E0G9_TRYCC                 NLNKLEAAAARDRLYAFQRVDDERLMRSALRNDIETLILHYKSLDAWDKAV------E-N   836
tr|A0A6A5CDJ2|A0A6A5CDJ2_NAEFO         PLTPEQLKASREKVKNVEDFEALKRRISAARNNLESALYS-TREKLLDNPD-YKGFFT-T   657
tr|A0A0S4IUZ2|A0A0S4IUZ2_BODSA         PMSKNESILSRDLLRRWRRSDELHLQKSAARNDLEAYLIWALSDGFVYNDE-ALKVVGSQ   782
tr|A0A0L1KXM1|A0A0L1KXM1_9EUGL         L-STTEFESEKLLLKEQEAYEASKRAIAQARNDLETLIIGYEGDALEAL-Q-ESVGIS-I   660
tr|A0A0S4JJU2|A0A0S4JJU2_BODSA         G-TEEVLE-AKSILSGLEKRDKIKRDTATSKNNLETFIQWIRLDGLFENNE-ATSLMS-T   667
tr|A0A0N0VI62|A0A0N0VI62_LEPPY         L-TLQESSASEKKLADIWKAEHAKHVRATAKNNLETYIFWVKYDGVGGNET-LAAAVG-D   655
tr|A4HND7|A4HND7_LEIBR                 L-TSEEMESSQAKLKTIWHAEHVKHLRATAKNNLETYIFWTKYEGVADNAE-LTATAG-A   655
tr|A4IC10|A4IC10_LEIIN                 L-TSEEMETSQTKLKTIWQAEHVKHLRATAKNNLEAYIFWAKNEGVSDNAA-LTAAAG-A   655
tr|E9AFU8|E9AFU8_LEIMA                 L-TPEEMQTSQAKLEAIWQAEHVKHLRATAKNNLEAYIFWAKNDGVSDNAA-LTAAAG-A   654
tr|A0A7G2C9U8|A0A7G2C9U8_9TRYP         P-SKEELEKSKEKISLIETSERIKYERAHAKNDLETFIFWAKREGVLENSD-VTSKIT-E   541
tr|S9TX72|S9TX72_9TRYP                 L-TEEEAAHIAERLFAVDEHERVKAERAHARNDLESFIFWAKREGILENTT-IVKKTD-A   654
tr|S9UBL2|S9UBL2_9TRYP                 L-TEEEAAHIAERLFAVDEHERVKAERAHARNDLESFIFWAKREGILENTT-IVKKTD-A   654
tr|Q38E16|Q38E16_TRYB2                 L-SYELLNNSRVKIFQLLEKERKKHEAATAKNNLETYLFWAKTEGILENTT-ALEPYT-P   653
tr|F9W5M9|F9W5M9_TRYCI                 ------------------------------------------------------------   452
tr|A0A1X0P4V9|A0A1X0P4V9_9TRYP         L-SEENFQKSRKKILDILDKERRKHEAATAKNNLETYVFWVKREGVLENTS-FLSLLS-E   643
tr|Q4CS27|Q4CS27_TRYCC                 L-SEEAFRRSKRRIAEILDKERKKHEAAVAKNNLEAYVFWAKSEGILENET-ALGVIS-T   655
tr|Q4E4F4|Q4E4F4_TRYCC                 L-SEEAFRRSKKRIAEILDKERKKHEAAVAKNNLEAYVFWAKSEGILENET-ALGVIS-T   655
tr|A0A422P1B1|A0A422P1B1_TRYRA         L-SGEVYQHLSEKNSSDT------------------------------------------   609
tr|A0A422N0X0|A0A422N0X0_9TRYP         L-SLEAFQHSREKISLILEKEQMKHEAAVAKNNLEAYVFWAKSEGILGNST-ALGLLS-A   646

tr|A0A0C9MAJ6|A0A0C9MAJ6_9FUNG         SDIEKLRERLSEVSDWIYDEG-----EHADTPVFTSKLKELQLIEQPIQHRVREYNERDN   705
sp|Q9Y4L1|HYOU1_HUMAN                  EQREEISGKLSAASTWLEDEG-----VGATTVMLKEKLAELRKLCQGLFFRVEERKKWPE   820
sp|Q63617|HYOU1_RAT                    EQREEISGKLSATSTWLEDEG-----FGATTVMLKDKLAELRKLCQGLFFRVEERRKWPE   820
tr|A0A067CBX2|A0A067CBX2_SAPPC         EQRETLSAAVDATEEWLYEDG-----DDLDAKAYNERLSGMRKQLDDMVFRVSELTALPE   735
tr|D8RVG4|D8RVG4_SELML                 KQRQEFLARLDEAGEWLYSDG-----EAATASEFKKRLGELKAIGDPIFFRLEQLTARPA   709
tr|A0A1P8B8R9|A0A1P8B8R9_ARATH         EERKAFVEKLDEVQDWLYMDG-----EDANATEFEKRLDSLKAIGSPISFRSEELTARPV   767
tr|A0A176WKM4|A0A176WKM4_MARPO         EQRDSLRAELNEAEDWLYTDG-----EDATAAEFKKKLDGLKKSGDAIFSRLDELIARPA   772
tr|A0A3R7K087|A0A3R7K087_9TRYP         AGEGNWREVVADVARWLDDAS-----DNVALAELQRQQRRMKELKTGDAATG--------   889
tr|Q4E175|Q4E175_TRYCC                 EGEANWREVVKEVEQWLDDTS-----DNVEISELQRQQQRMNKLQIGGATTG--------   883
tr|Q4E0G9|Q4E0G9_TRYCC                 EGEANWREVVKEVEQWLDDTS-----DNVEISELQRQQQRMNKLQIGGATTG--------   883
tr|A0A6A5CDJ2|A0A6A5CDJ2_NAEFO         KERDEINTVIDELEVWLGENDEEGVVEPSRVDLFTSKLKLLTDLTDPIQDRWREHKGRKE   717
tr|A0A0S4IUZ2|A0A0S4IUZ2_BODSA         EEMDTTAEKLRELQNWLEDA--G-TEATVPTEDLQAKLRQAKLIVRTALREAPTVEETKA   839
tr|A0A0L1KXM1|A0A0L1KXM1_9EUGL         EQTNVLLATISEVKEWLYDG--DGVSDSASVDSFRSRQQEILTIVEGIRYANINGSDDVD   718
tr|A0A0S4JJU2|A0A0S4JJU2_BODSA         EEKQTIEETTEQVKEWYEDG--EGSYDGCTKAEFDEKLALLKNVTATIREKIQEKENERK   725
tr|A0A0N0VI62|A0A0N0VI62_LEPPY         AAVHHVLDEVAKIQDWLEDG--PGSSDHCSADEYDARHAELERLVYELTKEPESESSHD-   712
tr|A4HND7|A4HND7_LEIBR                 TAIQRVMDEVTAVQEWLEGG--EGALDHCAASEYDSRLGNLRKMVSELTKKPEEATNIV-   712
tr|A4IC10|A4IC10_LEIIN                 AAVQRVIDEATAMQEWLEEG--EGASDNCAASQYESRLENLRGMVAELTKQPEEATKTV-   712
tr|E9AFU8|E9AFU8_LEIMA                 AAVQRVIDEATAVQEWLEEG--EGASDHCAASQYESRLESLRGMVAELTKQPEEATSTV-   711
tr|A0A7G2C9U8|A0A7G2C9U8_9TRYP         KQLEEYAALLEKSLNWLEDG--EGSYDSCTGEMYENKLSELKETLQSYLPVEEPKTNST-   598
```

```
tr|S9TX72|S9TX72_9TRYP              KLLGEFEQQLRDALEWLEDG--DGSMDSCPTKEYKSLLSQLKEQASGIQKPLEPVPVPS-    711
tr|S9UBL2|S9UBL2_9TRYP              KLLGEFEQQLRDALEWLEDG--DGSMDSCPTKEYKSLLSQLKEQASGIQKPLEPVPVPS-    711
tr|Q38E16|Q38E16_TRYB2              EEVNSLKNALAEAQEWLEDG--DGSNESCSKEEYENKLATLKELVKNSGKNKNTT-NTT-    709
tr|F9W5M9|F9W5M9_TRYCI              ------------------------------------------------------------    452
tr|A0A1X0P4V9|A0A1X0P4V9_9TRYP      VEIDRLKGKLSEIQEWLEDG--EGSYETCTKEEYEKKLKELKDLVRQETQNNTETRNPQ-    700
tr|Q4CS27|Q4CS27_TRYCC              EKIEMLRKKLSEVQEWLENE--DCGSEPCSKEEYEKKMEEIKQIVRQEPDATNETEVVI-    712
tr|Q4E4F4|Q4E4F4_TRYCC              EKVEMLRAKLSEVQEWLEDE--DCGSEQCSKEEYEKKMEEIKQIVRQEPDANNENEVVI-    712
tr|A0A422P1B1|A0A422P1B1_TRYRA      ------------------------------------------------------------    609
tr|A0A422N0X0|A0A422N0X0_9TRYP      EEAEVLRAKLSEVQEWIEDG--ICSYEACSKDEFEKKLQEIKQLVAQKRQSSAEAEPLK-    703


tr|A0A0C9MAJ6|A0A0C9MAJ6_9FUNG      NINLVDSTIKLAREFVTNISQVADDLRYHTEEELDGLLTAAEKLEDWMAEKVAAQKKLAN    765
sp|Q9Y4L1|HYOU1_HUMAN               RLSALDNLLNHSSMFLKGARLIPEMDQIFTEVEMTTLEKVINETWAWKNATLAEQAKLPA    880
sp|Q63617|HYOU1_RAT                 RLSALDNLLNHSSIFLKGARLIPEMDQIFTDVEMTTLEKVINDTWTWKNATLAEQAKLPA    880
tr|A0A067CBX2|A0A067CBX2_SAPPC      AVAKAQQYAISTIDLM---AQWVDAKPQVTADERDDVLAKVSELKDWLKEQQAAQDAIAK    792
tr|D8RVG4|D8RVG4_SELML              AMEAARGSLVESEAAI---HEWKEKKPWISD---AEVRKEGQKLVDWIEAKESEQAKVAD    763
tr|A0A1P8B8R9|A0A1P8B8R9_ARATH      AIEYARKYLTELKEII---KEWETNKTWLPKEKIDEVSKEAEKVKSWLDKNVAEQEKTSL    824
tr|A0A176WKM4|A0A176WKM4_MARPO      AVSEARTYLKTITETL---EEWEESKPWIPVKSKDDVRKEVESLRQWLDDMESKQSKLSG    829
tr|A0A3R7K087|A0A3R7K087_9TRYP      ------------------------------------------------------------    889
tr|Q4E175|Q4E175_TRYCC              ------------------------------------------------------------    883
tr|Q4E0G9|Q4E0G9_TRYCC              ------------------------------------------------------------    883
tr|A0A6A5CDJ2|A0A6A5CDJ2_NAEFO      ALSYCKSVFNATNYAI---HYMRENQKHITEDELNELHQFNKETEMNITSILAKQDRSPL    774
tr|A0A0S4IUZ2|A0A0S4IUZ2_BODSA      P-----------------------------------------------------------    840
tr|A0A0L1KXM1|A0A0L1KXM1_9EUGL      D-----------------------------------------------------------    719
tr|A0A0S4JJU2|A0A0S4JJU2_BODSA      P-----------------------------------------------------------    726
tr|A0A0N0VI62|A0A0N0VI62_LEPPY      ------------------------------------------------------------    712
tr|A4HND7|A4HND7_LEIBR              ------------------------------------------------------------    712
tr|A4IC10|A4IC10_LEIIN              ------------------------------------------------------------    712
tr|E9AFU8|E9AFU8_LEIMA              ------------------------------------------------------------    711
tr|A0A7G2C9U8|A0A7G2C9U8_9TRYP      ------------------------------------------------------------    598
tr|S9TX72|S9TX72_9TRYP              ------------------------------------------------------------    711
tr|S9UBL2|S9UBL2_9TRYP              ------------------------------------------------------------    711
tr|Q38E16|Q38E16_TRYB2              ------------------------------------------------------------    709
tr|F9W5M9|F9W5M9_TRYCI              ------------------------------------------------------------    452
tr|A0A1X0P4V9|A0A1X0P4V9_9TRYP      ------------------------------------------------------------    700
tr|Q4CS27|Q4CS27_TRYCC              ------------------------------------------------------------    712
tr|Q4E4F4|Q4E4F4_TRYCC              ------------------------------------------------------------    712
tr|A0A422P1B1|A0A422P1B1_TRYRA      ------------------------------------------------------------    609
tr|A0A422N0X0|A0A422N0X0_9TRYP      ------------------------------------------------------------    703


tr|A0A0C9MAJ6|A0A0C9MAJ6_9FUNG      TDDPVLVTSHVVDRAQTVKEHLVKLMSKKKPKVPKKTPEVPK--NDT-E--KEQNDSNED    820
sp|Q9Y4L1|HYOU1_HUMAN               TEKPVLLSKDIEAKMMALDREVQYLLNKAKFTKPRPRPKDKN--GTRAE--PPLNASASD    936
sp|Q63617|HYOU1_RAT                 TEKPVLLSKDIEAKMMALDREVQYLLNKAKFTKPRPRPKDKN--GTRTE--PPLNASAGD    936
tr|A0A067CBX2|A0A067CBX2_SAPPC      HDEPVFTSAHVLKKISGIKKVVATLAKKPKPTP-KPTEEAKPAEGDEAKKDDEPTTSSDE    851
tr|D8RVG4|D8RVG4_SELML              HEQPAFSSTEVYARIEKFRVLVARTGAQKAPKP-PKIEEVVK--NDAGSEEPKVGEENPG    820
tr|A0A1P8B8R9|A0A1P8B8R9_ARATH      WSKPVFTSTEVYAKVFTLQDKVTKVNKIPKPKP--KIEKVTK--TENTTKEEEQSKSSDE    880
tr|A0A176WKM4|A0A176WKM4_MARPO      HVEPAFTSDLVYTEVSRLRGKVLLLSKLQKPKP--KVEKPTE--NIEAETKPSESSQVDD    885
tr|A0A3R7K087|A0A3R7K087_9TRYP      ------------------------------------------------------------    889
tr|Q4E175|Q4E175_TRYCC              ------------------------------------------------------------    883
tr|Q4E0G9|Q4E0G9_TRYCC              ------------------------------------------------------------    883
tr|A0A6A5CDJ2|A0A6A5CDJ2_NAEFO      YEAPTVRAKDIKKQCGKVSERAFLLSKKPVPKPEKPATNET---NETT--IPSTNEETKT    829
tr|A0A0S4IUZ2|A0A0S4IUZ2_BODSA      ---------------------------KAPSVTTESAE-TAA--EGEA--VDEEIEEGDD    868
tr|A0A0L1KXM1|A0A0L1KXM1_9EUGL      --------------------------------PTNEDS--VA--GEGD----------EN    733
tr|A0A0S4JJU2|A0A0S4JJU2_BODSA      ---------------------------KPAPVPKKQPKKRKP--ETPQ--EP--AEV-PE    752
tr|A0A0N0VI62|A0A0N0VI62_LEPPY      ----------------------------------ASNN--G---------------VGGA    721
tr|A4HND7|A4HND7_LEIBR              ----------------------------------TE------------------------    714
tr|A4IC10|A4IC10_LEIIN              ----------------------------------TT------------------------    714
tr|E9AFU8|E9AFU8_LEIMA              ----------------------------------TT------------------------    713
tr|A0A7G2C9U8|A0A7G2C9U8_9TRYP      ----------------------------------TQEVNSTV--GEED--VD--FEMGTD    618
tr|S9TX72|S9TX72_9TRYP              ----------------------------------ENDA----------------------    715
tr|S9UBL2|S9UBL2_9TRYP              ----------------------------------ENDA----------------------    715
tr|Q38E16|Q38E16_TRYB2              ----------------------------------SGSD----------------------    713
tr|F9W5M9|F9W5M9_TRYCI              ------------------------------------------------------------    452
tr|A0A1X0P4V9|A0A1X0P4V9_9TRYP      ----------------------------------ESDD----------------------    704
tr|Q4CS27|Q4CS27_TRYCC              ----------------------------------ESSD----------------------    716
tr|Q4E4F4|Q4E4F4_TRYCC              ----------------------------------ESSD----------------------    716
tr|A0A422P1B1|A0A422P1B1_TRYRA      ------------------------------------------------------------    609
tr|A0A422N0X0|A0A422N0X0_9TRYP      ----------------------------------EATG----------------------    707


tr|A0A0C9MAJ6|A0A0C9MAJ6_9FUNG      S---EKKQQEEATQS---E--IPAAT-------SEEHEH---------------------    844
sp|Q9Y4L1|HYOU1_HUMAN               Q---GEKVIPPAGQT---EDAEPISEPEKVETGSEPGDTEPLELGGPGAEPEQKEQSTGQ    990
sp|Q63617|HYOU1_RAT                 Q---EEKVIPPTGQT---EEAKAILEPDKEGLGTEAADSEPLELGGPGAESEQAEQTAGQ    990
tr|A0A067CBX2|A0A067CBX2_SAPPC      E----------PK--------TP------EPTEEE-------------------------    862
tr|D8RVG4|D8RVG4_SELML              ASSGGGDSNSGSTQEGEREIESPDQAQAFEDESST-------------------------    855
tr|A0A1P8B8R9|A0A1P8B8R9_ARATH      A----------AKEEESHDEL---------------------------------------    891
tr|A0A176WKM4|A0A176WKM4_MARPO      T----------AKQEEKIEETVPQGEGAEMNQESK-------------------------    910
tr|A0A3R7K087|A0A3R7K087_9TRYP      ------------------------------------------------------------    889
tr|Q4E175|Q4E175_TRYCC              ------------------------------------------------------------    883
tr|Q4E0G9|Q4E0G9_TRYCC              ------------------------------------------------------------    883
tr|A0A6A5CDJ2|A0A6A5CDJ2_NAEFO      E---ENTANSQEQQS-----TNE-------------------------------------    844
tr|A0A0S4IUZ2|A0A0S4IUZ2_BODSA      D---GA--------D-----S---------------------------------------    873
tr|A0A0L1KXM1|A0A0L1KXM1_9EUGL      A---PN--------D-----P---------------------------------------    738
tr|A0A0S4JJU2|A0A0S4JJU2_BODSA      E---PT--------E-----ANP-------------------------------------    759
tr|A0A0N0VI62|A0A0N0VI62_LEPPY      A---SN--------E-----GD--------------------------------------    727
tr|A4HND7|A4HND7_LEIBR              --------------S-----SD--------------------------------------    717
tr|A4IC10|A4IC10_LEIIN              --------------D-----SA--------------------------------------    717
```

```
tr|E9AFU8|E9AFU8_LEIMA               --------------D-----SA--------------------------------------    716
tr|A0A7G2C9U8|A0A7G2C9U8_9TRYP       E---PA--------E-----PD--------------------------------------    624
tr|S9TX72|S9TX72_9TRYP               --------------A-----AT--------------------------------------    718
tr|S9UBL2|S9UBL2_9TRYP               --------------A-----AT--------------------------------------    718
tr|Q38E16|Q38E16_TRYB2               --------------D-----A---------------------------------------    715
tr|F9W5M9|F9W5M9_TRYCI               ------------------------------------------------------------    452
tr|A0A1X0P4V9|A0A1X0P4V9_9TRYP       --------------D-----A---------------------------------------    706
tr|Q4CS27|Q4CS27_TRYCC               --------------D---------------------------------------------    717
tr|Q4E4F4|Q4E4F4_TRYCC               --------------D---------------------------------------------    717
tr|A0A422P1B1|A0A422P1B1_TRYRA       ------------------------------------------------------------    609
tr|A0A422N0X0|A0A422N0X0_9TRYP       --------------D---------------------------------------------    708
```

```
tr|A0A0C9MAJ6|A0A0C9MAJ6_9FUNG       ------DEL    847
sp|Q9Y4L1|HYOU1_HUMAN                KRPLKNDEL    999
sp|Q63617|HYOU1_RAT                  KRPLKNDEL    999
tr|A0A067CBX2|A0A067CBX2_SAPPC       --PTHSDEL    869
tr|D8RVG4|D8RVG4_SELML               --TPLRDEL    862
tr|A0A1P8B8R9|A0A1P8B8R9_ARATH       ---------    891
tr|A0A176WKM4|A0A176WKM4_MARPO       --PTEHDEL    917
tr|A0A3R7K087|A0A3R7K087_9TRYP       ---------    889
tr|Q4E175|Q4E175_TRYCC               ---------    883
tr|Q4E0G9|Q4E0G9_TRYCC               ---------    883
tr|A0A6A5CDJ2|A0A6A5CDJ2_NAEFO       --QKQKDEL    851
tr|A0A0S4IUZ2|A0A0S4IUZ2_BODSA       --GASRDEL    880
tr|A0A0L1KXM1|A0A0L1KXM1_9EUGL       ------EEL    741
tr|A0A0S4JJU2|A0A0S4JJU2_BODSA       --EVPQEEL    766
tr|A0A0N0VI62|A0A0N0VI62_LEPPY       --DGAEGDL    734
tr|A4HND7|A4HND7_LEIBR               --DYNEDDL    724
tr|A4IC10|A4IC10_LEIIN               --DDNEGDL    724
tr|E9AFU8|E9AFU8_LEIMA               --DDNEDDL    723
tr|A0A7G2C9U8|A0A7G2C9U8_9TRYP       --ETLEGDL    631
tr|S9TX72|S9TX72_9TRYP               --SEGEEDL    725
tr|S9UBL2|S9UBL2_9TRYP               --SEGEEDL    725
tr|Q38E16|Q38E16_TRYB2               ----VKGDL    720
tr|F9W5M9|F9W5M9_TRYCI               ---------    452
tr|A0A1X0P4V9|A0A1X0P4V9_9TRYP       ----VKGDL    711
tr|Q4CS27|Q4CS27_TRYCC               ----DRGDL    722
tr|Q4E4F4|Q4E4F4_TRYCC               ----DRGDL    722
tr|A0A422P1B1|A0A422P1B1_TRYRA       ---------    609
tr|A0A422N0X0|A0A422N0X0_9TRYP       ----DRGDL    713
```

### 11. Shewanella-like phosphatases

The Shewanella-like phosphatases are an unusual family of enzymes with both prokaryotic and eukaryotic members. Although not present in animals, it might be an important enzymes for several organisms, e.g. fungi, and might not only act as a protein phosphatase. The KDEL-containing branch we identified in kinetoplastids seems to be ancient, as other eugleniids also seem to possess shewanella-like phosphatases of the essentially same architecture.

```
tr|C5L0A1|C5L0A1_PERM5               ------------------------------------------------------------    0
tr|A0A0S4JBT9|A0A0S4JBT9_BODSA       ------------------------------------------------------------    0
tr|Q583S8|Q583S8_TRYB2               ------------------------------------------------------------    0
tr|A0A1X0NU01|A0A1X0NU01_9TRYP       ------------------------------------------------------------    0
tr|A0A3R7MVL3|A0A3R7MVL3_9TRYP       ------------------------------------------------------------    0
tr|Q4DFK9|Q4DFK9_TRYCC               ------------------------------------------------------------    0
tr|Q4DQ38|Q4DQ38_TRYCC               ------------------------------------------------------------    0
tr|A0A0N0VHG5|A0A0N0VHG5_LEPPY       ------------------------------------------------------------    0
tr|A4HCJ2|A4HCJ2_LEIBR               ------------------------------------------------------------    0
tr|A4I008|A4I008_LEIIN               ------------------------------------------------------------    0
tr|Q4QBJ8|Q4QBJ8_LEIMA               ------------------------------------------------------------    0
sp|Q944L7|SLP2_ARATH                 ------------------------------------------------------------    0
tr|A0A813HX77|A0A813HX77_POLGL       MSFAFMAVLGLSQDVVFSVEQAIEQDMVVNANFAFSHNMGHKGLEDAHSFADIFSWLRLG    60
tr|A0A0S4JMN0|A0A0S4JMN0_BODSA       ------------------------------------------------------------    0
tr|Q4CWC2|Q4CWC2_TRYCC               ------------------------------------------------------------    0
tr|A0A1Y2EN62|A0A1Y2EN62_9FUNG       ------------------------------------------------------------    0
tr|A0A0C9MUE8|A0A0C9MUE8_9FUNG       ------------------------------------------------------------    0
```

```
tr|C5L0A1|C5L0A1_PERM5               ------------------------------------------------------------    0
tr|A0A0S4JBT9|A0A0S4JBT9_BODSA       --------------------------------------------MVVFFLMLPQQERNKI    16
tr|Q583S8|Q583S8_TRYB2               ------------------------------------------------------------    0
tr|A0A1X0NU01|A0A1X0NU01_9TRYP       ------------------------------------------------------------    0
tr|A0A3R7MVL3|A0A3R7MVL3_9TRYP       ------------------------------------------------------------    0
tr|Q4DFK9|Q4DFK9_TRYCC               ------------------------------------------------------------    0
tr|Q4DQ38|Q4DQ38_TRYCC               ------------------------------------------------------------    0
```

```
tr|A0A0N0VHG5|A0A0N0VHG5_LEPPY      -----------------------------------------------------------M    1
tr|A4HCJ2|A4HCJ2_LEIBR              -----------------------------------------------------------M    1
tr|A4I008|A4I008_LEIIN              -----------------------------------------------------------M    1
tr|Q4QBJ8|Q4QBJ8_LEIMA              -----------------------------------------------------------M    1
sp|Q944L7|SLP2_ARATH                ------------------------------------------------------------    0
tr|A0A813HX77|A0A813HX77_POLGL      FLPLVIQPSWSYSESRDADSASVFDLAPMRASVSNFLVRQIQQDGIPFFLFHPSTFRETS    120
tr|A0A0S4JMN0|A0A0S4JMN0_BODSA      ------------------------------------------------------------    0
tr|Q4CWC2|Q4CWC2_TRYCC              ------------------------------------------------------------    0
tr|A0A1Y2EN62|A0A1Y2EN62_9FUNG      ------------------------------------------------------------    0
tr|A0A0C9MUE8|A0A0C9MUE8_9FUNG      ------------------------------------------------------------    0


tr|C5L0A1|C5L0A1_PERM5              ------------------------------------------------------------    0
tr|A0A0S4JBT9|A0A0S4JBT9_BODSA      ----------------------------NNNPSYNMRL-SLTIVTLCL--VA--------    37
tr|Q583S8|Q583S8_TRYB2              -------------------------------------MVGKYLLISLVLVAH--------    15
tr|A0A1X0NU01|A0A1X0NU01_9TRYP      -----------------------------------MRLVTPF--FVIILLVY--------    15
tr|A0A3R7MVL3|A0A3R7MVL3_9TRYP      -----------------------------------MKLAALLLLQLCLIIVT--------    17
tr|Q4DFK9|Q4DFK9_TRYCC              -----------------------------------MRLVEFLLPLLVILLGF--------    17
tr|Q4DQ38|Q4DQ38_TRYCC              -----------------------------------MRLVEFLLPLLVILLGF--------    17
tr|A0A0N0VHG5|A0A0N0VHG5_LEPPY      ----------------------------RRCAVNYYFTFFALVTSICSVLVA--------    25
tr|A4HCJ2|A4HCJ2_LEIBR              ----------------------------RNYVLRAVRCYSALLVSFSLLLVS--------    25
tr|A4I008|A4I008_LEIIN              ----------------------------KVCALRIGRCYGALLLLFYLLLIS--------    25
tr|Q4QBJ8|Q4QBJ8_LEIMA              ----------------------------KVCDLQIGRCYGALLLLFYVLLIS--------    25
sp|Q944L7|SLP2_ARATH                -------------------------MSSRENPSG------ICKSIPKLI-----------    18
tr|A0A813HX77|A0A813HX77_POLGL      SSDERGYIRFFTHTRGISFLASYVFIKFGARPAI-LAAFLIQKHVFNVFNVFRSRASLSE    179
tr|A0A0S4JMN0|A0A0S4JMN0_BODSA      -----------------------------------MNASLFATSIFVVLTLL--------    17
tr|Q4CWC2|Q4CWC2_TRYCC              -------------MRGV----------L-------------------LLLLF--------    10
tr|A0A1Y2EN62|A0A1Y2EN62_9FUNG      -------------MSDI----------LSANPSLTTSTSVFQSK-STSTVVP--------    28
tr|A0A0C9MUE8|A0A0C9MUE8_9FUNG      ----------------M----------V---------------K-FTSFLSA--------    10


tr|C5L0A1|C5L0A1_PERM5              ----------MTFHRI---------AVVAALLVVSVATSSSRPLKRIVAVADVHGDRRNL    41
tr|A0A0S4JBT9|A0A0S4JBT9_BODSA      -TTLFVDA------------------------------RDAAGAPRRIIAVGDVHGDLAQL    67
tr|Q583S8|Q583S8_TRYB2              -LGIRCCANV--------------------DANPLSRNVIPIEIHRIVAVGDVHGDAERF    54
tr|A0A1X0NU01|A0A1X0NU01_9TRYP      -IPYFCEGES--------------------RSNLETHDTIPIEIHRIIAVGDVHGDLDHF    54
tr|A0A3R7MVL3|A0A3R7MVL3_9TRYP      -TVDVCYGES--------------------RANTHIGTTVPLEIHRIIAVGDVHGDADNF    56
tr|Q4DFK9|Q4DFK9_TRYCC              -PVVFCYGEG--------------------TASGGDVHAIPIEIHRIIAVGDVHGDTENF    56
tr|Q4DQ38|Q4DQ38_TRYCC              -PVVFCYGES--------------------TASGGDVHAIPIEIHRIIAVGDVHGDTENF    56
tr|A0A0N0VHG5|A0A0N0VHG5_LEPPY      -TPAAT--------------------------------KELVEIHRIVTVGDVHGDAENF    53
tr|A4HCJ2|A4HCJ2_LEIBR              -FPVHCE-------------------------------RKLVEVHRIIAVGDVHGDADNF    53
tr|A4I008|A4I008_LEIIN              -CLARCE-------------------------------RKLVEVHRIIAVGDVHGDADNF    53
tr|Q4QBJ8|Q4QBJ8_LEIMA              -CFARCE-------------------------------RKLVEVHRIIAVGDVHGDADNF    53
sp|Q944L7|SLP2_ARATH                ------SSFVDTFVDYSVSGIFLPQDP---SSQNEILQTRFEKPERLVAIGDLHGDLEKS    69
tr|A0A813HX77|A0A813HX77_POLGL      CAPMGCNCQMLTSVELS-GGRWLRRWHPLLGVTFAVLAAGSSSPSRVVALADLHGDYDHA    238
tr|A0A0S4JMN0|A0A0S4JMN0_BODSA      ----NLASP---------------------HDGMASALFGYHPPPRLLAIGDLHGDITNA    52
tr|Q4CWC2|Q4CWC2_TRYCC              ----GAA------------------AYFVAFL-HVEAVNVRGTEPRLVAFGDIHGDVFRC    47
tr|A0A1Y2EN62|A0A1Y2EN62_9FUNG      ----GEQSKEDT-LDST------KKDSSTKEVKIEKRKKSEKVFKRVVAVGDIHGDYKKL    77
tr|A0A0C9MUE8|A0A0C9MUE8_9FUNG      ----GC----------------------LLFSSLVSAAISVEQDRRIVALGDLHGDLANT    44
                                                                                 *:::..*:***  .

tr|C5L0A1|C5L0A1_PERM5              MQALENGRVLVHKHGPEQE-------------------GVEWHPEAS--DPAEPVMEGTQ    80
tr|A0A0S4JBT9|A0A0S4JBT9_BODSA      RAVLEMANVIND-------------------------EAQWIDRD------------NTV    90
tr|Q583S8|Q583S8_TRYB2              RQILEMSGVISLRSNSSKQ----V--------VW----KPRWGTKEGNFFREYGTRLRTT    98
tr|A0A1X0NU01|A0A1X0NU01_9TRYP      LKILSMADVIAFPNNDTND----V--------VW----KPKWENKEIELHRTHRTKLRTT    98
tr|A0A3R7MVL3|A0A3R7MVL3_9TRYP      RQILFMTGIISYSSNTDKN----V--------IW----KPMWDEKEIELRKSHRTKLRTT    100
tr|Q4DFK9|Q4DFK9_TRYCC              RQILSMAGIVSYSSNADKQ----V--------LW----KPMWDEKEIELHKRHHTKLRST    100
tr|Q4DQ38|Q4DQ38_TRYCC              RQILSMAGIVSYSSNADKQ----L--------LW----KPMWDEKEIELHKRHHTKLRST    100
tr|A0A0N0VHG5|A0A0N0VHG5_LEPPY      LKVLRLADVVEEGTGGVVD----V--------LTN---PPQWKFSQ---APNASVTSWTT    95
tr|A4HCJ2|A4HCJ2_LEIBR              LKILRIANLIEDSASGASD----V--------LDS---PPRWKYSS---SQISDTTVRTT    95
tr|A4I008|A4I008_LEIIN              LKILRIANLIEDGVTGASG----V--------LDN---PPRWKYSS---SRPNGTAVRTT    95
tr|Q4QBJ8|Q4QBJ8_LEIMA              LKILRIANLIEDSVTGASG----V--------LDN---PPRWKYSS---SRTNGTSVRTT    95
sp|Q944L7|SLP2_ARATH                REAFKIAGLIDSS--------------------------DRWTGG------------STM    91
tr|A0A813HX77|A0A813HX77_POLGL      LAILRAAGLADENSEIGVSEADELLVAEAARKPWSRFRGVQWLGG------------NAT    286
tr|A0A0S4JMN0|A0A0S4JMN0_BODSA      KEILKFAQIVDCE--------------------------GKWIAG------------HDT    74
tr|Q4CWC2|Q4CWC2_TRYCC              RQILQLANITDTE--------------------------DRWIAG------------SSI    69
tr|A0A1Y2EN62|A0A1Y2EN62_9FUNG      IKVLYTAKIIDNK--------------------------SNWIAK------------DTI    99
tr|A0A0C9MUE8|A0A0C9MUE8_9FUNG      LEILRFSNIIDQD--------------------------NHWIGG------------DTI    66
                                    :        :                               *

tr|C5L0A1|C5L0A1_PERM5              VVQLGDLVDRGPLGLQCYRLMQDLYVAE----GANEVVRVLGNHEVLNLLGMAGRYVTDE    136
tr|A0A0S4JBT9|A0A0S4JBT9_BODSA      VVQVGDLLDRGPHDKGVLDFMMQLQAEAPT--HGGKVVSILGNHELMNIMNQ-VHYVHPE    147
tr|Q583S8|Q583S8_TRYB2              LIQTGDLIDRGEEDLEVLEMAVSLFNEVRTNYTDDKVVLLMGNHELLNLQGH-FHYVHSK    157
tr|A0A1X0NU01|A0A1X0NU01_9TRYP      LVQMGDLIDRGANDLGVLEMVFSLFDQVKKNHTSDNLVLLLGNHELLNLQGQ-FYYVEPE    157
tr|A0A3R7MVL3|A0A3R7MVL3_9TRYP      LIQMGDLIDRGEDDLAVLEMAASLLDQVKSNHTSDNFVLLLGNHELLNLQQQ-FHYVHPE    159
tr|Q4DFK9|Q4DFK9_TRYCC              LIQMGDLIDRGEDDLGVLEMAFSLFEQVKSNHTSDNIVLLLGNHELLNLQEQ-FYYVHPE    159
tr|Q4DQ38|Q4DQ38_TRYCC              LIQMGDLIDRGEDDLGVLEMAFSLFEQVKSNHTSDNIVLLLGNHELLNLQEQ-FYYVHPE    159
tr|A0A0N0VHG5|A0A0N0VHG5_LEPPY      LVQMGDLVDRGEQDFESLNIAMALQEQTQNSLTSDRVILLIGNHELLNIQGH-YHYVNKR    154
tr|A4HCJ2|A4HCJ2_LEIBR              LVQVGDLIDRGEQDLETLNIAISLQEQTAQSGSQDKVVLLIGNHELLNLQGH-YHYVNEK    154
tr|A4I008|A4I008_LEIIN              LVQVGDLIDRGEQDLQALNIAISLQEQTAQSGTQDEVVLLIGNHELLNIQGH-YHYVNKN    154
tr|Q4QBJ8|Q4QBJ8_LEIMA              LVQVGDLIDRGEQDLEALNIAISLQEQTAQSGSQDEVVLLIGNHELLNIQGH-YHYVNKH    154
sp|Q944L7|SLP2_ARATH                VVQVGDVLDRGGEELKILYFLEKLKREAER--AGGKILTMNGNHEIMNIEGD-FRYVTKK    148
tr|A0A813HX77|A0A813HX77_POLGL      LVQTGDLVDRGPFARDLYALFSELRRQAPL--AGGQVINLIGNHEAMNVLGQ-LKYVTKE    343
tr|A0A0S4JMN0|A0A0S4JMN0_BODSA      VIQVGDVVDRGPHGHEIIDYFTTLKEDARQ--SGGEFIQLLGNHELMNMQGN-MKFANAQ    131
tr|Q4CWC2|Q4CWC2_TRYCC              VVQLGDIADRGLHPHEIYDLFASLERQAMK--AGGEFIFLVGNHELMNILGI-FYYVHPD    126
tr|A0A1Y2EN62|A0A1Y2EN62_9FUNG      LIQTGDLIDRGNDTILIFDLMMKLKEQAKK--HGCILYLLLGNHELMNLQED-YRYVTRG    156
tr|A0A0C9MUE8|A0A0C9MUE8_9FUNG      FVQTGDVLDRGLDTIKLYDLLQNLRDEAPL--QGGLVIPLLGNHELMNLVGD-WRYVYPG    123
                                    .:* **: ***            *            .  : **** :*:      :.
```

```
tr|C5L0A1|C5L0A1_PERM5                  DVAEF------------------------------------------GGEEARRESWSP-    153
tr|A0A0S4JBT9|A0A0S4JBT9_BODSA          SIEYF------------------------------------------EGSVNRKRQFSA-    164
tr|Q583S8|Q583S8_TRYB2                  SMGGF------------------------------------------LTRALRKRAFEL-    174
tr|A0A1X0NU01|A0A1X0NU01_9TRYP          SMGGF------------------------------------------LTKTLRKRAFEK-    174
tr|A0A3R7MVL3|A0A3R7MVL3_9TRYP          SMGGF------------------------------------------LTKALRKRAFES-    176
tr|Q4DFK9|Q4DFK9_TRYCC                  TMGGF------------------------------------------LSKTLRKRAFEP-    176
tr|Q4DQ38|Q4DQ38_TRYCC                  TMGGF------------------------------------------LSKTLRKRAFEP-    176
tr|A0A0N0VHG5|A0A0N0VHG5_LEPPY          NYGGF------------------------------------------ISRPLRMEAMNA-    171
tr|A4HCJ2|A4HCJ2_LEIBR                  NHGGF------------------------------------------MSKALRAEGMKV-    171
tr|A4I008|A4I008_LEIIN                  NYGGF------------------------------------------LSKALRAEGMKA-    171
tr|Q4QBJ8|Q4QBJ8_LEIMA                  NYGGF------------------------------------------LSKALRAEGMKA-    171
sp|Q944L7|SLP2_ARATH                    GLEEFQIWADWYCLGNKMKTLCSGLDKPKDPYEGIPMSFPRMRADCFEGIRARIAALRP-    207
tr|A0A813HX77|A0A813HX77_POLGL          DTDEF------------------------------------------GSHAEREQAFAA-    360
tr|A0A0S4JMN0|A0A0S4JMN0_BODSA          TVTAF------------------------------------------GGMESLKQAFDPI    149
tr|Q4CWC2|Q4CWC2_TRYCC                  VMSAF------------------------------------------GGEEAYAAAFGP-    143
tr|A0A1Y2EN62|A0A1Y2EN62_9FUNG          DVISF------------------------------------------GGMANRRKEFSL-    173
tr|A0A0C9MUE8|A0A0C9MUE8_9FUNG          EPETF------------------------------------------GGLEARRKAFEK-    140
                                                 *

tr|C5L0A1|C5L0A1_PERM5                  GGEIWTILKDHYELVHVYGGHRFVHGGVMPALTDRSIDELNEQASRMIKN----------    203
tr|A0A0S4JBT9|A0A0S4JBT9_BODSA          EGVYGQWIRQN-PLLHVEGSTLFVHAGLSPQFAAKGVDALNKEAKEALSQ----------    213
tr|Q583S8|Q583S8_TRYB2                  DGTFGGFILENFTVAYAVADTLFVHAGIDEHVVSDGIERLNREAKQAIRT----------    224
tr|A0A1X0NU01|A0A1X0NU01_9TRYP          NGEFGSFILENFTVLHLEDSTVFVHAGLNEQVASMGVEAINKVTKQAVQD----------    224
tr|A0A3R7MVL3|A0A3R7MVL3_9TRYP          NGIFGRFLLYNFTVAHFDAETLFVHAGLNEHFASMGVDRLNKETMQAVRE----------    226
tr|Q4DFK9|Q4DFK9_TRYCC                  SGTFGRFLLDKFNVLYFDAETVFVHAGIDQRFASVGVEMLNKKTMQAIRE----------    226
tr|Q4DQ38|Q4DQ38_TRYCC                  SGTFGRFLLDKFNVLYFDAETVFVHAGIDQHFASVGVEMLNKKTMQAIRE----------    226
tr|A0A0N0VHG5|A0A0N0VHG5_LEPPY          DGAFGKYIIDNFRTAYLDENTLFVHAGIEADMLLPDLNSLNSEVREALRK----------    221
tr|A4HCJ2|A4HCJ2_LEIBR                  TGVFGKYIVDNFKVAHIDEGVLFVHGGIETGMNIKDVDALNEDVRSALRQ----------    221
tr|A4I008|A4I008_LEIIN                  TGAFGKYIVDNFKAAHMDEGVLFVHAGIETSMNIKDVEALNADIREALRQ----------    221
tr|Q4QBJ8|Q4QBJ8_LEIMA                  TGAFGKYIVDNFKAAHMDEGVLFIHAGIETSMNIKDVEALNADVREALRQ----------    221
sp|Q944L7|SLP2_ARATH                    DGPIAKRFLTKNQTVAVVGDSVFVHGGLLAEHIEYGLERINEEVRGWING---------F    258
tr|A0A813HX77|A0A813HX77_POLGL          GGWVGRQILEEFKALAVVAETLFVHAGLLPEHAAMGVESLDLQVRTDLAK-----AAASR    415
tr|A0A0S4JMN0|A0A0S4JMN0_BODSA          TGAYGTYLTKTCDAAVLRNRTLFVHAGVLPQYAHLGVAELNRQVRNAILA----------    199
tr|Q4CWC2|Q4CWC2_TRYCC                  EGPYGLYILQ-HPVTVVREGVVFAHAGITPEYAAKGVEGINAELMNGFRGERDLLLEGDA    202
tr|A0A1Y2EN62|A0A1Y2EN62_9FUNG          DGKYGKLLRSEMNATMIIDDTLFVHAGLTTVFAKYGVDQMNDHIHYILKT----YPADQL    229
tr|A0A0C9MUE8|A0A0C9MUE8_9FUNG          DGFIGEYLKL-LNITTKVGSTVFCHGGITPHFSRLGINWINDKTHETIVT----YMESQG    195
                                        *        :                 * *.*:            .:  ::   .

tr|C5L0A1|C5L0A1_PERM5                  -GALKN----PLL-LSESSPLWSRVYALGTD-EEACPPLLNVLRHY----------GVAR    246
tr|A0A0S4JBT9|A0A0S4JBT9_BODSA          -KSLRHLDRSALF--GVHGPIWTRLLITDAM-NGRCKDVEESLSLL---------PGIER    260
tr|Q583S8|Q583S8_TRYB2                  -KNFGH----ILL--GSTGPLWSRKMFLDAS-NGRCADTKKALASL----------GVKR    266
tr|A0A1X0NU01|A0A1X0NU01_9TRYP          -KDYRH----PLL--GSSGPLWTRQMIMDAM-NGHCTNIQKMLLAI----------GAER    266
tr|A0A3R7MVL3|A0A3R7MVL3_9TRYP          -KDYQN----PLL--GAFGPLWTRRMLMDAT-NGHCTGIKKMISSI----------GAKR    268
tr|Q4DFK9|Q4DFK9_TRYCC                  -KDYGN----PLL--GTSGPLWTRKMITDAA-NGRCTGIQKMLSFI----------GAKR    268
tr|Q4DQ38|Q4DQ38_TRYCC                  -KDYGN----PLL--GTSGPLWTRKMITDAA-NGRCTGIQKMLSFI----------GAKR    268
tr|A0A0N0VHG5|A0A0N0VHG5_LEPPY          -RDFRH----AYL--RSNGPLWTRKMISDSM-MGDCDEVNKILQRF----------NVSR    263
tr|A4HCJ2|A4HCJ2_LEIBR                  -NIFRH----SFL--RSSGPLWTRKMIMESM-SDECADVEAALKQL----------NASR    263
tr|A4I008|A4I008_LEIIN                  -GIFRH----SFL--GSSGPLWTRKMIIESM-SEGCSDVRAALKQL----------NATR    263
tr|Q4QBJ8|Q4QBJ8_LEIMA                  -GIFRH----SFL--GSSGPLWTRKMIIESM-SDECSDVRAALKQL----------NATR    263
sp|Q944L7|SLP2_ARATH                    KGGRYA----PAYCRGGNSVVWLRKFSEEMAHKCDCAALEHALST---------IPGVKR    305
tr|A0A813HX77|A0A813HX77_POLGL          SGGHGQ----PLL--KSNGPLWVRQFAKGQD-SKVCPLLAATLKLVG----------AKR    458
tr|A0A0S4JMN0|A0A0S4JMN0_BODSA          -KNISD----PIL--LDDGPLWTRQIIYPAQ-KGNCSNLYESLKILSEYETKHGRKPVDR    251
tr|Q4CWC2|Q4CWC2_TRYCC                  KGDGVH----PLS--NSSSPLWSRAVLDEAK-RGNCSLLMESLRLLSEHELASGRPPVRV    255
tr|A0A1Y2EN62|A0A1Y2EN62_9FUNG          ---FYA----PIF--SNNGPFWTRFLSMGQE-VAVCEELNMVLEMMN----------VKR    269
tr|A0A0C9MUE8|A0A0C9MUE8_9FUNG          HRGKSH----GIF--GGVGPTWYRGYALEEE-DEICSVLDEALELLE----------ANR    238
                                                          .  * *            *        :

tr|C5L0A1|C5L0A1_PERM5                  MVVGHTPSEDGRMKVRCGGRAILADVALSRWMGR----------YPHHGHPAALEMTLVN    296
tr|A0A0S4JBT9|A0A0S4JBT9_BODSA          MVVGHTPQRSGRVETYCDDKLIAVDVGLSKWMY---------------GNLAALEITQKV    305
tr|Q583S8|Q583S8_TRYB2                  VVVGHTPQRSGRVETFCGGSVIAIDVGMSRWMY---------------GNIAALEITVTS    311
tr|A0A1X0NU01|A0A1X0NU01_9TRYP          IVVGHTPQRSGHVETYCDDSVIAVDVGLSKWMY---------------GNLAALEILVKT    311
tr|A0A3R7MVL3|A0A3R7MVL3_9TRYP          IVVGHTPQRSGHVELFCDDSVIAVDVGLSRWMY---------------GNLAALELMVTT    313
tr|Q4DFK9|Q4DFK9_TRYCC                  IVVGHTPQRSGHVEVFCNDSVIAIDVGLSRWMY---------------GNLAALELMVTS    313
tr|Q4DQ38|Q4DQ38_TRYCC                  IVVGHTPQRSGHVEVFCNDSVIAIDVGLSRWMY---------------GNLAALELMVTS    313
tr|A0A0N0VHG5|A0A0N0VHG5_LEPPY          IVVGHTPQDSGQIEQYCNKRVIAADVGMSRWMY---------------NNVAALEMVFFK    308
tr|A4HCJ2|A4HCJ2_LEIBR                  VVVGHTPQESGHIGQYCGGQVLAIDVGISRWMY---------------DRAVALELVFLK    308
tr|A4I008|A4I008_LEIIN                  VVVGHTPQESGHIGQHCDGQVFAIDVGMSRWMY---------------DKVAALELVFSK    308
tr|Q4QBJ8|Q4QBJ8_LEIMA                  IVVGHTPQESGHIGQHCDGQVLAIDVGMSRWMY---------------DKVAALELVFSK    308
sp|Q944L7|SLP2_ARATH                    MIMGHTIQDAG-INGVCNDKAIRIDVGMSKGCAD--------------GLPEVLEIRRDS    350
tr|A0A813HX77|A0A813HX77_POLGL          MVIGHTQVDEGEIRQRCGNRLLLADTIISRQGYPECWEPDSWQRERCKGSLSYVELRGDE    518
tr|A0A0S4JMN0|A0A0S4JMN0_BODSA          MVIGHTIMQDGAIHQFCDGKLIAIDIAISQYMA------------SGGRQ-GILELRYDF    298
tr|Q4CWC2|Q4CWC2_TRYCC                  MVGGHTVQEGGVMAVECNGSLVGADVGLSRFFS------------SYGGYVAYVEFLLDD    303
tr|A0A1Y2EN62|A0A1Y2EN62_9FUNG          MVVGHTVQENRKINTRCGNKFLLIDVGMSEFYG------------GAFAY---LEFLNDK    314
tr|A0A0C9MUE8|A0A0C9MUE8_9FUNG          MVVGHTVQHDGAIRTRCNGKVILIDIGISRAYN------------SGKGA---LEIRGNV    283
                                        :: ***   :   *.   .  *  :*.                           :*:

tr|C5L0A1|C5L0A1_PERM5                  ATHLEKIEA-------------------HY-GPENDTGAGQHQLLWTNTD-GLIEVGKDE    335
tr|A0A0S4JBT9|A0A0S4JBT9_BODSA          D--GSVELREIIPHNVATPPAAPPAIETEGDNHDAKQGSGIEGGVSDPILLQELMEAIQE    363
tr|Q583S8|Q583S8_TRYB2                  YSSGDGVEEEVVLH--------------EVLASDARRNGTLEDFLNDTLFLEELQHAVEE    357
tr|A0A1X0NU01|A0A1X0NU01_9TRYP          HKIGSGAKKSVEIR--------------EIIASDTRRSQTLDEALDDSLVLEELQHAVEE    357
tr|A0A3R7MVL3|A0A3R7MVL3_9TRYP          YKSDSGIWRDVVMR--------------EVTTSDSRRFRTLDESLTDPLLLEELNHAVEE    359
tr|Q4DFK9|Q4DFK9_TRYCC                  YKCDLGICRDVVMR--------------EITNSDARRSQTIDESLKDSLLLEELMHAVEE    359
tr|Q4DQ38|Q4DQ38_TRYCC                  YKCDLGICRDVVMR--------------EITNSDARRSQTIDESLKDSLLLEELMHAVEE    359
tr|A0A0N0VHG5|A0A0N0VHG5_LEPPY          YLDTDLQQ------------------------------------ISTEFIIRELREGVSA    332
tr|A4HCJ2|A4HCJ2_LEIBR                  YTDSLTGS------------------------------------SSSSFVVSELREGVSG    332
tr|A4I008|A4I008_LEIIN                  YKDNITGT------------------------------------GSASFVVRELREGVSG    332
tr|Q4QBJ8|Q4QBJ8_LEIMA                  YMDKITGT------------------------------------GSASFVVRELREGVSG    332
sp|Q944L7|SLP2_ARATH                    GVRI-----------VTSNPLYKENLYSHV-APDSKT--GLGLLV--PVP---KQVEVKA    391
```

```
tr|A0A813HX77|A0A813HX77_POLGL     AYAMRAL-G---IDAIESAPE--------------Q------RLL--PIEGESGE-----    547
tr|A0A0S4JMN0|A0A0S4JMN0_BODSA     ATH-----TDI-VTPYIQYPSQPFR--QHA--------------L--PAVMFPKRIPVPP    334
tr|Q4CWC2|Q4CWC2_TRYCC             SDP---------SR-RIAVPQYPFG--RAV-RPPTS------PLL--KDPMRPPRRMPDN    342
tr|A0A1Y2EN62|A0A1Y2EN62_9FUNG     NEIWAVY-SDT-NRERID------------------------------------------    330
tr|A0A0C9MUE8|A0A0C9MUE8_9FUNG     --VNALY-ADK-RVKLASPPPYKS----RV--------------L--AS----KKHDHDE    315

tr|C5L0A1|C5L0A1_PERM5             FADSDEDE--------------------------L-------------------------    344
tr|A0A0S4JBT9|A0A0S4JBT9_BODSA     MKVKQQAEKAALASKEQQE----VVARDEL------------------------------    389
tr|Q583S8|Q583S8_TRYB2             YNQRQT-----------------TKGKNDVLDD-L-------------------------    374
tr|A0A1X0NU01|A0A1X0NU01_9TRYP     FRHRQN-----------------TRDQEETLGD-L-------------------------    374
tr|A0A3R7MVL3|A0A3R7MVL3_9TRYP     YYQSEN-----------------TVPTEDLVGD-L-------------------------    376
tr|Q4DFK9|Q4DFK9_TRYCC             FYQSPK-----------------DNPPEDTDYE-L-------------------------    376
tr|Q4DQ38|Q4DQ38_TRYCC             FYQSPK-----------------DNPPEDTDYE-L-------------------------    376
tr|A0A0N0VHG5|A0A0N0VHG5_LEPPY     FSMTSDGRNDGEQLSPKSS----FIV-DDDNGD-L-------------------------    361
tr|A4HCJ2|A4HCJ2_LEIBR             FCYTCVEARDDTDSPRGGG----SVE-KDTYDD-L-------------------------    361
tr|A4I008|A4I008_LEIIN             FCHTCVEERSGLDTTGGGD----SDEENDIFDD-L-------------------------    362
tr|Q4QBJ8|Q4QBJ8_LEIMA             FCHTCLDERSGLDTTVGGD----SD-ENDIFDD-L-------------------------    361
sp|Q944L7|SLP2_ARATH               ------------------------------------------------------------    391
tr|A0A813HX77|A0A813HX77_POLGL     -APSGTLGK--LASWLQGDWPAWTNVEL--------------------------------    572
tr|A0A0S4JMN0|A0A0S4JMN0_BODSA     FEAKEKGGV-----QGA-----ALTIRT--YEDSMTKSELVLLC-----IAFFFVAGLVA    377
tr|Q4CWC2|Q4CWC2_TRYCC             WQPSSVRGK-----EGEGKWRRAVKRRTKRGDDGLLHVSVQTLCFFISVLAFFLVCVVRL    397
tr|A0A1Y2EN62|A0A1Y2EN62_9FUNG     ------------------------------------------------------------    330
tr|A0A0C9MUE8|A0A0C9MUE8_9FUNG     L-----------------------------------------------------------    316

tr|C5L0A1|C5L0A1_PERM5             ----------------------------   344
tr|A0A0S4JBT9|A0A0S4JBT9_BODSA     ----------------------------   389
tr|Q583S8|Q583S8_TRYB2             ----------------------------   374
tr|A0A1X0NU01|A0A1X0NU01_9TRYP     ----------------------------   374
tr|A0A3R7MVL3|A0A3R7MVL3_9TRYP     ----------------------------   376
tr|Q4DFK9|Q4DFK9_TRYCC             ----------------------------   376
tr|Q4DQ38|Q4DQ38_TRYCC             ----------------------------   376
tr|A0A0N0VHG5|A0A0N0VHG5_LEPPY     ----------------------------   361
tr|A4HCJ2|A4HCJ2_LEIBR             ----------------------------   361
tr|A4I008|A4I008_LEIIN             ----------------------------   362
tr|Q4QBJ8|Q4QBJ8_LEIMA             ----------------------------   361
sp|Q944L7|SLP2_ARATH               ----------------------------   391
tr|A0A813HX77|A0A813HX77_POLGL     ----------------------------   572
tr|A0A0S4JMN0|A0A0S4JMN0_BODSA     SQRVKDGGKGLFSLSWLGGRRRKLEHAA   405
tr|Q4CWC2|Q4CWC2_TRYCC             RRR--RAANGRHRSWWKL----------   413
tr|A0A1Y2EN62|A0A1Y2EN62_9FUNG     ----------------------------   330
tr|A0A0C9MUE8|A0A0C9MUE8_9FUNG     ----------------------------   316
```

### 12. Calreticulin chaperones

Calreticulins are essential chaperones for secreted and transmembrane protein folding, and are found in an ER-resident, KDEL-motif restreined form in most eukaryotes. Kinetoplastids are no exception.

```
tr|S9W2Z8|S9W2Z8_9TRYP             ---MSRTTTFLVLLAALCALCVVQAEIYYHEKFDNLD----EWVTSSH-S---KYGKVRF    49
tr|A4HJP8|A4HJP8_LEIBR             --MTQRTMLAAVVGVLVLCVCLVQAEIFFHEEFNTLD----GWVQSEHKD---DYGKVEL    51
tr|A4I765|A4I765_LEIIN             --MAQRAMLAAVVGVLVLCVYVVQAEIFFHEEFNTMD----GWVQSEHTS---DYGKVAL    51
tr|Q4Q601|Q4Q601_LEIMA             --MAQRAMLAAIVGVLVLCVYVVQAEIFFHEEFNTMD----GWVQSEHTS---DYGKVAL    51
tr|A0A0S4KMN1|A0A0S4KMN1_BODSA     ----MSRVIALVAA-AALLCAAAQAKVYFHEEFNSLD----GWVVSEAKE---GLGKAVL    48
tr|A0A1X0NIG9|A0A1X0NIG9_9TRYP     ----MRAIVLLCALLGLSMLVSVNGKILFHEDFSSLD----KWVQSKERD---DYGKVEL    49
tr|F9WHI5|F9WHI5_TRYCI             ----MRRGTIVCALLGLVAAATVQGTIYFHEKFTSIN----HWTPSKARS---DYGKVEL    49
tr|Q57YU3|Q57YU3_TRYB2             MLMCMRPVAVACVFVALATVATVHGAIHFHEKFSSID----HWTASKARS---DYGKVEL    53
tr|Q584K4|Q584K4_TRYB2             MLMCMRPVAVACVFVALATVATVHGAIHFHEKFSSID----HWTASKARS---DYGKVEL    53
tr|Q4CPZ0|Q4CPZ0_TRYCC             ----MRAAIFFCALLGLATLSAVHGTVYFHEEFKSME----HWTTSKHRD---DFGKVEI    49
tr|Q4DDX3|Q4DDX3_TRYCC             ----MRAAIFFCALLSLATLSAVHGTVYFHEEFKSME----HWTTSKHRD---DFGKVEI    49
tr|A0A3R7NR70|A0A3R7NR70_9TRYP     ----MRAALLVCALLGLAALSAVEATVYFHEEFSSLE----RWTPSKHRD---DLGKVEL    49
tr|A0A3R7KPX3|A0A3R7KPX3_TRYRA     ----MPAAILVCALLALATLAAVDATVYFHEEFKSLD----HWTPSKYRD---DLGKVEL    49
tr|A0A367JDX0|A0A367JDX0_RHIAZ     ----MKIPTI---TAILGLAALASAEVFFHETFSDGEGWKDRWTPSTYRE---DLGKLEV    50
tr|S2IZR0|S2IZR0_MUCC1             ----MKIPTI---AAVLGLATLVSAEVFLHETFSDGEGWKDRWTASEHRE---DLGKLEV    50
sp|O04151|CALR1_ARATH              MAKLNPKFIS---LILFALVVIVSAEVIFEEKFE--DGWEKRWVKSDWKKDDNTAGEWKH    55
sp|Q9ZNY3|CALR_EUGGR               --M--RKELW---LGLLLSSQAVLSTIYYKETFE--PDWETRWTHSTAKS---DYGKFKL    48
tr|Q6PE26|Q6PE26_DANRE             MRI--TAAVC---FISALA-FIAHADVYFKEQFLDGDGWKSRWVESKHKS---DYGQWKL    51
sp|P27797|CALR_HUMAN               MLL--SVPLL---LGLLGL-AVAEPAVYFKEQFLDGDGWTSRWIESKHKS---DFGKFVL    51
                                                              .   :  .* *       *  *   .         *:

tr|S9W2Z8|S9W2Z8_9TRYP             SSGEKPVDR-EMQRGLQLAEESSFYAVSKKLPTRITNDGKPLVVSFQLLHRGRFTCGGGY    108
tr|A4HJP8|A4HJP8_LEIBR             SAGALHVDA-AKEQGMKLTEDSKFYAISKELPTPVSNDGKPLVISFSVKNEQNLKCGGAY    110
tr|A4I765|A4I765_LEIIN             SVGAIHVDA-EKEQGLKLMEDAKFYAVSKKLPKAVSNDGKSIVVSFSVKNEQKLTCGGTY    110
tr|Q4Q601|Q4Q601_LEIMA             SVGAIHVDA-EKEQGLKLMEDAKFYAVSKKLPKAVSNDGKPIVVSFSVKNEQKLTCGGTY    110
tr|A0A0S4KMN1|A0A0S4KMN1_BODSA     SSGKFFGDE-KINQGLKTSEDAHFYAIAKTLPEAVTNEGKDLVVSISVKHEQGLDCGGGY    107
tr|A0A1X0NIG9|A0A1X0NIG9_9TRYP     SAGKNEPEGSSNKKGLRLTEDARFYAISTEFPTPISNENKELVFSFVVKHEQDLKCGGAY    109
tr|F9WHI5|F9WHI5_TRYCI             SAGKFFADA-EKSKGLRLTEDARFYALSTPLPTPISNEKKDFVVSFSVKHEQGLRCGGGY    108
tr|Q57YU3|Q57YU3_TRYB2             SAGKFYADA-EKSKGLRLTEDARFYALSTPLPTPITNEKKDFVVSFSVKHEQDLRCGGGY    112
tr|Q584K4|Q584K4_TRYB2             SAGKFYADA-EKSKGLRLTEDARFYALSTPLPTPITNEKKDFVVSFSVKHEQDLRCGGGY    112
tr|Q4CPZ0|Q4CPZ0_TRYCC             SAGKFYADA-EKSKGLRLTEDARFYALSTAFPTPINNEKKSLVVSFSVKHEQDLKCGGGY    108
tr|Q4DDX3|Q4DDX3_TRYCC             SAGKFYADA-EKSKGLRLTEDARFYALSTAFPTPITNEKKSLVVSFSVKHEQDLKCGGGY    108
```

```
tr|A0A3R7NR70|A0A3R7NR70_9TRYP     SAGDFYADA-EKSKGLRLTEDARFYAVSTALPTPITNDGKDLVVSFSVKHGQELKCGGGY    108
tr|A0A3R7KPX3|A0A3R7KPX3_TRYRA     SAGDFYADA-EKSKGLRLMDDARFYAFSTALPTPITNDGKDFVVSFSVKHGQDLKCGGGY    108
tr|A0A367JDX0|A0A367JDX0_RHIAZ     SPGKWYVDE-KANAGLRTTEDYRFYAVSSK-TKPFTNKDKNLVIQFDVKNEQDIDCGGSY    108
tr|S2IZR0|S2IZR0_MUCC1             SPGKWFADE-AYNAGLRTTEDYRFYATSTKIPKPFSNKDKDLVIQFDVKNEQDIDCGGSY    109
sp|O04151|CALR1_ARATH              TAGNWSGDA--NDKGIQTSEDYRFYAISAEF-PEFSNKDKTLVFQFSVKHEQKLDCGGGY    112
sp|Q9ZNY3|CALR_EUGGR               TSGKFYGDK-AKDAGIQTSQDAKFYAISSPIASSFSNEGKDLVLQFSVKHEQDIDCGGGY    107
tr|Q6PE26|Q6PE26_DANRE             TSGKFYGDA-ELDKGLQTSQDARFYALSSRF-DSFSNEGKTLVIQFTVKHEQKIDCGGGY    109
sp|P27797|CALR_HUMAN               SSGKFYGDE-EKDKGLQTSQDARFYALSASF-EPFSNKGQTLVVQFTVKHEQNIDCGGGY    109
                                   : *      :        . *::  ::  *** :          ..*. : :*..: : :   : *** *

tr|S9W2Z8|S9W2Z8_9TRYP             IKLLP-AMDQKDFHGETEYGLMFGPDICGG-ERRVHIIFNYKGQNYLWNKRELFV---VD    163
tr|A4HJP8|A4HJP8_LEIBR             LKFFS-ELNQKDLHSESPYWLMFGPDVCGF-QNRLHFIFNYNDENHLWRSFWRLTKELNE    168
tr|A4I765|A4I765_LEIIN             LKFFS-ELDQKDLHGESAYWLMFGPDTCGS-STRLQFILSYNGTNHLWKKLWRPK---TD    165
tr|Q4Q601|Q4Q601_LEIMA             LKFFS-ELDQKDLHGESAYWLMFGPDICGS-NKRLQFILSYNGTNHLWKKLSTPK---TD    165
tr|A0A0S4KMN1|A0A0S4KMN1_BODSA     AKFLP-ATELKGFNGDTEYLLMFGPDQCGY-NKRVHVIFNYKGTNLLWKKEPKYP---DD    162
tr|A0A1X0NIG9|A0A1X0NIG9_9TRYP     LKYLP-AMDPTKFNGESKYWLMFGPDRCGS-KNKVHIILHYNGVNHEWSKNFRYP---ED    164
tr|F9WHI5|F9WHI5_TRYCI             IKLMP-NMDPAEFKGETKYWLMFGPDRCGY-NNKIHIIINYNNTNMEWKKHPRYP---DD    163
tr|Q57YU3|Q57YU3_TRYB2             IKLLP-QMDPAELKGETKYWLMFGPDRCGY-DKKIHIIISYNGANREWKKRPSYP---DD    167
tr|Q584K4|Q584K4_TRYB2             IKLLP-QMDPAELKGETKYWLMFGPDRCGY-DKKIHIIISYNGANREWKKRPSYP---DD    167
tr|Q4CPZ0|Q4CPZ0_TRYCC             IKLLP-SMDPEKFHGETKYWLMFGPDRCGS-QNRVHIILHYNGENREWSKRIRFP---ED    163
tr|Q4DDX3|Q4DDX3_TRYCC             IKLLP-SMDPEKFHGETKYWLMFGPDRCGS-QNRVHIILHYNGENREWSKRIRFP---ED    163
tr|A0A3R7NR70|A0A3R7NR70_9TRYP     IKLLP-AMDAEKFNGETKYWLMFGPDRCGA-KSKVQIILHRNGENHEWSKEVRFP---VD    163
tr|A0A3R7KPX3|A0A3R7KPX3_TRYRA     IKLLP-AMDAEKFNGETKYWLMFGPDRCGA-KNKVHIILHRNGENHEWSKNIHFP---DD    163
tr|A0A367JDX0|A0A367JDX0_RHIAZ     LKFFGGEFDPKTFNGDTSYNIMFGPDICGP-KSIVHAIFNYNGTNYDFKKTISAP---KD    164
tr|S2IZR0|S2IZR0_MUCC1             LKIFG-DLDPKAFNGDSEYNIMFGPDICGP-KAMVHAIFNYNGTNHDLKKSISAP---KD    164
sp|O04151|CALR1_ARATH              MKLLSDDVDQTKFGGDTPYSIMFGPDICGYSTKKVHAILTYNGTNHLIKKEVPCE---TD    169
sp|Q9ZNY3|CALR_EUGGR               LKLLPS-VDAAKFTGDTPYHIMFGPDICGA-TKKIHFILTYKGKNLLWKKEPRCE---TD    162
tr|Q6PE26|Q6PE26_DANRE             VKVFPAEMDQTEMHGESQYYIMFGPDICGYSTKKVHVIFNYKGQNHLIKKDIKCK---DD    166
sp|P27797|CALR_HUMAN               VKLFPNSLDQTDMHGDSEYNIMFGPDICGPGTKKVHVIFNYKGKNVLINKDIRCK---DD    166
                                   * :      :   : .:: * :***** **      :: *:  :. *      .        :

tr|S9W2Z8|S9W2Z8_9TRYP             ELTHAYTLAIYPNNTYALYLDANLTVSGSIAEEWSMLPPEMIDDPSDKKPAHWDDEPAMI    223
tr|A4HJP8|A4HJP8_LEIBR             KATHVYTVQISPNNTYQLYVDGKYIQEGSLVDEWEMLPPKTIADPEEKKPSNWIEDS-MM    227
tr|A4I765|A4I765_LEIIN             KATHVYTVEIAPNNTYQLYVDGMHIQEGSFEEEWDMLPPKTIPDPTDEKPADWVDDM-MM    224
tr|Q4Q601|Q4Q601_LEIMA             TATHVYTVEIAPNNTYQLYVDGMHIQEGSLEEEWDMLPPKTISDPTDKKPADWVDDM-MM    224
tr|A0A0S4KMN1|A0A0S4KMN1_BODSA     KLTHVYTLRVTPENKYELYIDQELKESGALEADWEFLQPKEIDDATDSKPADWVDAS-QI    221
tr|A0A1X0NIG9|A0A1X0NIG9_9TRYP     SITHAYTLRIAPDNSYSFYLDGQERMKGTLEEDWSLLPAKEIADPSDVKPADWVDEP-TI    223
tr|F9WHI5|F9WHI5_TRYCI             KLTHAYTLHIAPDNSYKFYLDGSPTEKGNLEDDWDFLPPKEIEDPEDKKPADWVDAK-TI    222
tr|Q57YU3|Q57YU3_TRYB2             KLTHVYTLHITPSNSYEFFLDGVSKEKGTLEADWDFLPEKEIDDPEDKKPADWVDVP-TI    226
tr|Q584K4|Q584K4_TRYB2             RLTHVYTLHITPSNSYEFFLDGVSKEKGTLEADWDFLPEKEIDDPEDKKPADWVDVP-TI    226
tr|Q4CPZ0|Q4CPZ0_TRYCC             KLTHVYTLHIAADNSYEFFLDGESKAKGQLEEDWSLLLPREIVDETDKKPEDWVDEE-TM    222
tr|Q4DDX3|Q4DDX3_TRYCC             KLTHVYTLHIAADNSYEFFLDGESKAKGQLDGDWTLLPPREIVDETDKKPEDWVDEE-TM    222
tr|A0A3R7NR70|A0A3R7NR70_9TRYP     KLTHVYTLHIAADNTYEFFLDGELRAKGELEADWHLLPPREIVDPDDTKPADWVDAA-TI    222
tr|A0A3R7KPX3|A0A3R7KPX3_TRYRA     KLTHVYTLRIATDNTYEFFLDGESKAKGALEDDWSLLPPREITDETDKKPDDWVDEK-TM    222
tr|A0A367JDX0|A0A367JDX0_RHIAZ     TLTHTYTLIVKPDQTYEILVDSDKKASGSLLEDWDFLPPKQIKDPNAVKPADWVDDA-LI    223
tr|S2IZR0|S2IZR0_MUCC1             TLTHTYTLVVKPDQTYEILIDGKSKADGSLLEDWDFLPPKTIKDPNASKPEDWVEEA-MI    223
sp|O04151|CALR1_ARATH              QLTHVYTFVLRPDATYSILIDNVEKQTGSLYSDWDLLPAKKIKDPSAKKPEDWDDKE-YI    228
sp|Q9ZNY3|CALR_EUGGR               TLSHTYTAVIKADRTYEVLVDQVKKESGTLEEDWEILKPKTIPDPEDKKPADWVDEP-DM    221
tr|Q6PE26|Q6PE26_DANRE             ELTHLYTLILRPDQTYEVKIDNEKVESGSLEEDWDFLPPKKIKDPEAKKPDDWDDRA-KI    225
sp|P27797|CALR_HUMAN               EFTHLYTLIVRPDNTYEVKIDNSQVESGSLEDDWDFLPPKKIKDPDASKPEDWDERA-KI    225
                                   :* **  :  . .* . :*        * :  :* :*  . * * ** .* :  :

tr|S9W2Z8|S9W2Z8_9TRYP             PDPDDTKPEDWDKEPATILDPDATKPNDWNDAEDGEWEPPIITNPNYKGDWFPRKIKNPK    283
tr|A4HJP8|A4HJP8_LEIBR             DDPYDTKPEDWDDELPTIADDTAVKPDDWVDEEDGEWEAPRIPNPKYRGAWVPRRIDNPN    287
tr|A4I765|A4I765_LEIIN             DDPSDTKPEHWDDEPATITDSEAVKPVDWDDAEDGVWEAPKIPNPNYRGAWTPRRIHNPD    284
tr|Q4Q601|Q4Q601_LEIMA             DDPSDTKPEDWEDEPATITDSEAVKPVDWDDSEDGVWEAPKVPNPNYRGAWKPRRIHNPD    284
tr|A0A0S4KMN1|A0A0S4KMN1_BODSA     DDPEDVKPADWESEPATVVDPEAKKPDDWDDEEDGAWEAPQIPNPKAKGEWRARRIPNPA    281
tr|A0A1X0NIG9|A0A1X0NIG9_9TRYP     VDPNDVKPADWDNEPAMIPDTEAKRPDDWDDAEDGEWEAPMVPNPNAKGEWHPKRIPNPA    283
tr|F9WHI5|F9WHI5_TRYCI             DDPEDKKPDDWDNEPETIVDPDAKKPDDWNDAEDGTWEAPMIPNPKSKGPWSPRKITNPA    282
tr|Q57YU3|Q57YU3_TRYB2             DDPEDKKPEDWDSEPEKIVDPEAKKPEDWNDAEDGAWEAPMIPNPKSKGPWAPRKIPNPA    286
tr|Q584K4|Q584K4_TRYB2             DDPEDKKPEDWDSEPEKIVDPEAKKPEDWNDAEDGAWEAPMISNPKSKGPWAPRKIPNPA    286
tr|Q4CPZ0|Q4CPZ0_TRYCC             DDPEDKKPEDWDNEPAMIPDADAKKPDDWDDAEDGPWEAPMIPNPKSKGTWKPRQIPNPA    282
tr|Q4DDX3|Q4DDX3_TRYCC             DDPEDKKPEDWDNEPAMIPDADAKKPDDWDDAEDGPWEAPMIPNPKSKGTWKPRQIPNPA    282
tr|A0A3R7NR70|A0A3R7NR70_9TRYP     DDPADTKPADWDDEPAMIPDAAAKKPDDWDDAEDGEWEAPMVPNPKAKGAWTPRKIPNPA    282
tr|A0A3R7KPX3|A0A3R7KPX3_TRYRA     DDPEDKKPADWDEEPEMIPDEEAKKPEDWDDAEDGAWEAPMIPNPKAKGAWKPRQIPNPA    282
tr|A0A367JDX0|A0A367JDX0_RHIAZ     DDPNDVKPEGYDDIPEFIPDPNAKKPEDWDDDMDGEWEAPSIANPDYKGPWTPKKIPNPA    283
tr|S2IZR0|S2IZR0_MUCC1             VDETDVKPANYDDIPEFIPDPEAKKPEDWDDDMDGEWEAPSIANPEYQGEWSPKKIPNPL    283
sp|O04151|CALR1_ARATH              PDPEDTKPAGYDDIPKEIPDTDAKKPEDWDDEEDGEWTAPTIPNPEYNGEWKPKKIKNPA    288
sp|Q9ZNY3|CALR_EUGGR               VDPEDKKPEDWDKEPAQIPDPDATQPDDWDEEEDGKWEAPMISNPKYKGEWKAKKIPNPA    281
tr|Q6PE26|Q6PE26_DANRE             DDPEDTKPEDWDK-PENIPDPDAKKPDDWDEDMDGEWEPAMIPNPEYKGEWKPKQIDNPS    284
sp|P27797|CALR_HUMAN               DDPTDSKPEDWDK-PEHIPDPDAKKPEDWDEEMDGEWEPPVIQNPEYKGEWKPRQIDNPD    284
                                   *  * **  ::.      : *  * :* ** :  ** *      : **. .* *   ::* **

tr|S9W2Z8|S9W2Z8_9TRYP             YKGQWKPRQIKNPNFYPDPDLYRMKKPLEYVGIDVWTVEAGSIYTHIMIGDNVEEVLDVL    343
tr|A4HJP8|A4HJP8_LEIBR             YKGVWSPQQIPNPDYKEDPNLYKSPAPLKYVGIDVWQVESGSIFDNIIIGDDLQEVLKVV    347
tr|A4I765|A4I765_LEIIN             YKGKWAARQIPNPAYKEDPNLYRAPAPLQYVGIDVWQVEGGSIFDDIIIGDDITEVLGVV    344
tr|Q4Q601|Q4Q601_LEIMA             YKGEWAARQIPNPVYKKDPNLYKVPAPLQYVGIDVWQVEGGSIFDDIIIGDDITEVLRVV    344
tr|A0A0S4KMN1|A0A0S4KMN1_BODSA     YKGPWKARQVANPKYEEDAKLYLLRKPLAAVGIDVWQVKAGSIFDNIIVGDDLAEVNKLI    341
tr|A0A1X0NIG9|A0A1X0NIG9_9TRYP     YKGEWAPRMIPNPEYKEDKELYKVREPLAHVGIDVWQVESGSIFRDIIIGDDVNEVLEMV    343
tr|F9WHI5|F9WHI5_TRYCI             YKGPWAPRKIANPAYKSDPNLYMIPEPLTHVGIDVWQVESGSIYKDIVVGDDLKEVLDIV    342
tr|Q57YU3|Q57YU3_TRYB2             YKGPWAPRRIPNPAYKNDEELYKIPEPLTHVGIDVWQVESGSIFKDIIIGDDVKEVLDIV    346
tr|Q584K4|Q584K4_TRYB2             YKGPWAPRRIPNPAYKNDEELYKIPEPLTHVGIDVWQVESGSIFKDIIIGDDVKEVLDIV    346
tr|Q4CPZ0|Q4CPZ0_TRYCC             YKGVWEPRKIPNPDFVEDSELHKVPEPLTHVGIDVWQVESGSIFKDIVIGDDLKEVLDLV    342
tr|Q4DDX3|Q4DDX3_TRYCC             YKGVWEPRKIPNPDFVEDSELHKVPEPLTHVGIDVWQVESGSIFKDIMIGDDLKEVLELV    342
tr|A0A3R7NR70|A0A3R7NR70_9TRYP     YKGAWEPRKIPNPAYKPDDSLYKIPEPLAHVGIELWQVESGSIFKDIVLGDDVAEVLDLV    342
tr|A0A3R7KPX3|A0A3R7KPX3_TRYRA     YKGEWKPRVIPNPEYKPDDGLYKIPEPLAHVGIDIWQVESGSIFKNIVLGDDVAEVLELV    342
tr|A0A367JDX0|A0A367JDX0_RHIAZ     YKGEWEHPLIDNPDYKVDNEIYAYD--FANVGIDVWQVKSGTIFDNILITDDIEEAEKVR    341
tr|S2IZR0|S2IZR0_MUCC1             YKGEWVHPEIDNPEYKVDNEIYAYD--FANVGIDVWQVKSGTIFDNILITDDVEEAKKIR    341
```

```
sp|O04151|CALR1_ARATH              YKGKWKAPMIDNPEFKDDPELYVFP-KLKYVGVELWQVKSGSLFDNVLVSDDPEYAKKLA    347
sp|Q9ZNY3|CALR_EUGGR               YKGVWKPRDIPNPEYEADDKVHIFD-EIAAVGFDLWQVKSGTIFDNIIVTDSLAEAKAFY    340
tr|Q6PE26|Q6PE26_DANRE             YKGTWVHPEIDNPEYAADDAIYKFD-SIGVLGLDLWQVKSGTIFDNFLITDDVEEAEKFG    343
sp|P27797|CALR_HUMAN               YKGTWIHPEIDNPEYSPDPSIYAYD-NFGVLGLDLWQVKSGTIFDNFLITNDEAYAEEFG    343
                                   *** *    : ** :  *  ::     :  :*.::* *:.*::: ..:: :.   .  .

tr|S9W2Z8|S9W2Z8_9TRYP             KTTFVNINKLEARLEKAEREKMEREREEKALEEAEKDETAA-------------------    384
tr|A4HJP8|A4HJP8_LEIBR             ESTYGAMAEDEMNLIEAEAEERQRKERKQE---EENSLTEQEKENSV-----------TE    393
tr|A4I765|A4I765_LEIIN             KSTYGAMAEKERDLIQAEEKKEATKEPAEA---AAEK-------PNV-----------GE    383
tr|Q4Q601|Q4Q601_LEIMA             KSTYGAMAEKERDLIQAEAKKEAMKEPAEA---AAEK-------PNM-----------DE    383
tr|A0A0S4KMN1|A0A0S4KMN1_BODSA     DATWGATKDAEKAAEDAAAAATKAADE--AAKADESKEE---------------------    378
tr|A0A1X0NIG9|A0A1X0NIG9_9TRYP     KKTVEEERKKEEAEKE---KKEADAKKE-----EEKKEGEKK------------------    377
tr|F9WHI5|F9WHI5_TRYCI             QSTYGDMKKAEEEAFDAFEKKEEEERKSKEAKEEEKKEEEKKEEEKK-----------E-    390
tr|Q57YU3|Q57YU3_TRYB2             KSTYDGMKKAEEDALAAFEKKKEQDKKEENKKETDGEEDKKKK-----------------    389
tr|Q584K4|Q584K4_TRYB2             KSTYDGMKKAEEDALAAFEKKEEQDEKEEDKKETDGEEDKKKK-----------------    389
tr|Q4CPZ0|Q4CPZ0_TRYCC             EKTYGGLKKAEADALKVMEDMEKEKRKKEEEEEREKEKAEE-------------------    383
tr|Q4DDX3|Q4DDX3_TRYCC             EKTYGSLKKAEADALKVMEDMEKEKRKKEEEEEKEKEKAEE-------------------    383
tr|A0A3R7NR70|A0A3R7NR70_9TRYP     KKTYGSLKQAEAEALNAFEGKQKEAEK--QKEAEEQKEAEAA---EN-------------    384
tr|A0A3R7KPX3|A0A3R7KPX3_TRYRA     KETYGSLKQAEAEALKAFETKQEAKKAEEAKEAEEAKEAEDEKEAED-----------EK    391
tr|A0A367JDX0|A0A367JDX0_RHIAZ     NETKDLHEKEEDAKKAHEEKVDAENKAKAEEAEKDAK-----------------------    378
tr|S2IZR0|S2IZR0_MUCC1             EETQVLRSPEEGAQAAYNEKIQAEAKAKAEAEGAAPPDEGAEKIDLEKFEPAVKFDEVPP    401
sp|O04151|CALR1_ARATH              EETWGKHKDAEKAAFD---EAEKKRE---EEESKDAPAESDAEEEAE--------DDDNE    393
sp|Q9ZNY3|CALR_EUGGR               DQTNGATKDAEKKAFD---SAEADKRKKEEDERKKQEEEEKKTAE-E--------DEDDD    388
tr|Q6PE26|Q6PE26_DANRE             TDTWGATKGPEKKMKD---QQEEEERKKREEEEKSKKDDNEEDEEDE--------DEDEP    392
sp|P27797|CALR_HUMAN               NETWGVTKAAEKQMKD---KQDEEQRLKEEEEDKKRKEEEEAEDKED--------DEDKD    392
                                   *        *

tr|S9W2Z8|S9W2Z8_9TRYP             --------------------AAEDDEDLGDE-DEENTDL    402
tr|A4HJP8|A4HJP8_LEIBR             HVEEETE---------APVALNGANDNEGNQ-DSNNEDL    422
tr|A4I765|A4I765_LEIIN             HADHTP--------------------DEGD--SEDKEDL    400
tr|Q4Q601|Q4Q601_LEIMA             HADHTP--------------------DEDK--SEGEEDL    400
tr|A0A0S4KMN1|A0A0S4KMN1_BODSA     --------------------------------------L    379
tr|A0A1X0NIG9|A0A1X0NIG9_9TRYP     -----------------------------EA-DEDKGDL    386
tr|F9WHI5|F9WHI5_TRYCI             ------------------EEKKEEEKKEEGK-KEEKADL    410
tr|Q57YU3|Q57YU3_TRYB2             ---------------------------------EDKSDL    395
tr|Q584K4|Q584K4_TRYB2             ---------------------------------EDKSDL    395
tr|Q4CPZ0|Q4CPZ0_TRYCC             --------------------EKDEEELEEKG-DGDKEDL    401
tr|Q4DDX3|Q4DDX3_TRYCC             --------------------EKDEEELEEKG-DGDKEDL    401
tr|A0A3R7NR70|A0A3R7NR70_9TRYP     --EVEAE---------AAAMEDEEAEKKADA-DGDKEDL    411
tr|A0A3R7KPX3|A0A3R7KPX3_TRYRA     EAEDEKE---------SEEVKESEEAKDAGA-DGDKEDL    420
tr|A0A367JDX0|A0A367JDX0_RHIAZ     -KDQEPETPKEEEILEALEDAVEEAVEADAAKKPARDEL    416
tr|S2IZR0|S2IZR0_MUCC1             AAAEALEKAKEEEILEAIKEEAEKKVEEEDAKKPVKDEL    440
sp|O04151|CALR1_ARATH              -GDDSDNESKSEETKEA------EETKEAEETDAAHDEL    425
sp|Q9ZNY3|CALR_EUGGR               --DEEEEED-------D-----------------KKDEL    401
tr|Q6PE26|Q6PE26_DANRE             EEDDHTEEP-------P------EEEEEGEDDALPKDEL    418
sp|P27797|CALR_HUMAN               E-DEEDEED-------K------EEDEEEDVPGQAKDEL    417
                                                                    *
```

### 13. Endoplasmin-like HSP90 chaperones

Endoplasmins represent a nearly ubiquitious ER-resident clade of the greater HSP90 family. These essential chaperones contain KDEL-like motifs in most eukaryotic lineages examined.

```
tr|A0A0L1KBS8|A0A0L1KBS8_9EUGL     ---------------------------------MRFWSIALVLA-IASVSSSPT------    20
tr|A0A0L1KHT0|A0A0L1KHT0_9EUGL     ---------------------------------------MSRISVYSLLVFGILLLSG--    19
tr|A0A1X0P079|A0A1X0P079_9TRYP     ---------------------------------MSKRTLFLQGLLIVLVVLGVTAVED--    25
tr|Q4DW89|Q4DW89_TRYCC             ----------------------------------MARHPIIQAILIALIVLGVAVTGV--    24
tr|A0A422PWD8|A0A422PWD8_9TRYP     -------------------------------MLVRRW--ILKALLIAVVALSAVASEE--    25
tr|A0A422NRM6|A0A422NRM6_TRYRA     ---------------------------------MRGWHAILQILLVAVVALSVVATED--    25
tr|A0A0S4KLS2|A0A0S4KLS2_BODSA     ----------------------------------MRLSLIFRIACVALLAFA---IV---    20
tr|A0A0N0VF86|A0A0N0VF86_LEPPY     ----------------------------------MRRSPLLRVALVALIFLGTISAT---    23
tr|A4HH83|A4HH83_LEIBR             ----------------------------------MANSNVHRVVLVALLLLGSVT-V---    22
tr|E9ADS8|E9ADS8_LEIMA             ----------------------------------MANSSLLRVVLVALLLLGSVT-V---    22
sp|Q9NGD0|ENPL_LEIIN               ----------------------------------MANSSLLRVVLVALLLLGSVT-V---    22
tr|A4I4C9|A4I4C9_LEIIN             ----------------------------------MANSSLLRVVLVALLLLGSVT-V---    22
sp|Q8T7E0|ENPL_LEIDO               ----------------------------------MANSSLLRVVLVALLLLGSVT-V---    22
tr|S9U4L8|S9U4L8_9TRYP             -----------------------------MARLNKTVKVVLQIALVVCLLLGATVAK---    28
tr|A0A7G2C3S2|A0A7G2C3S2_9TRYP     MAYSSSLSLFCSCSLPIIIIIAYYYNDWLTHPMGKVTQVVLKVVLIALLLVGATIAK---    57
sp|P14625|ENPL_HUMAN               ----------------------------------MRALWVL-GLCCVLLTFGSVRA----    21
tr|Q7T3L3|Q7T3L3_DANRE             ----------------------------------MRRLWII-GLLCALLAFASVKA----    21
sp|Q9STX5|ENPL_ARATH               ------------------------MRKRTLVSVLFL-FSLL-FLL----PDQGRKLHANA    30
tr|A0A061FKM2|A0A061FKM2_THECC     ------------------------MRKWAIPSALILLCL-L-SLL----SDQGRKVHANA    30
tr|Q5Z9N8|Q5Z9N8_ORYSJ             ------------------------MRKWALSSALLLLLLLL-TTL----PDPAKKLQVNA    31
tr|W2ZJ09|W2ZJ09_PHYPR             ------------------------MKGG-CRSLLLLLLMAV-TLCYSSLSSSSAR--AEV    32
tr|W2ZJ74|W2ZJ74_PHYPR             ------------------------MKGG-CRSLLLLLLMAV-TLCYSSLSSSSAR--AEV    32
tr|F0Y644|F0Y644_AURAN             --------------------------------MKLSILKLV-A----------LVFGAVA    17
tr|D8LHY7|D8LHY7_ECTSI             ----------------------------------MRRPRFV-ALCAAAALQSSTPWGGSA    25


tr|A0A0L1KBS8|A0A0L1KBS8_9EUGL     ----------------M--------------------------------AVEENVAESGG    32
tr|A0A0L1KHT0|A0A0L1KHT0_9EUGL     --------------GVE--------------------------------KTFLGAEKGET    33
```

```
tr|A0A1X0P079|A0A1X0P079_9TRYP      --------------AAK--------------------------------SDGGAQDKGKP    39
tr|Q4DW89|Q4DW89_TRYCC              --------------TVK--------------------------------DD-GSVEKGRP    37
tr|A0A422PWD8|A0A422PWD8_9TRYP      --------------AAQ--------------------------------TD-ATVEKGKP    38
tr|A0A422NRM6|A0A422NRM6_TRYRA      --------------AVQ--------------------------------SD-VVVEKGKP    38
tr|A0A0S4KLS2|A0A0S4KLS2_BODSA      -------------------------------------------------GEESASGKGSP    31
tr|A0A0N0VF86|A0A0N0VF86_LEPPY      ---------------------------------------------------SAGDGRGTP    32
tr|A4HH83|A4HH83_LEIBR              ---------------------------------------------------SAGDARGSP    31
tr|E9ADS8|E9ADS8_LEIMA              ---------------------------------------------------SAGDGRGTP    31
sp|Q9NGD0|ENPL_LEIIN                ---------------------------------------------------SAGDGRGTP    31
tr|A4I4C9|A4I4C9_LEIIN              ---------------------------------------------------SAGDGRGTP    31
sp|Q8T7E0|ENPL_LEIDO                ---------------------------------------------------SAGDGRGTP    31
tr|S9U4L8|S9U4L8_9TRYP              -------------------------------------------------ETTEQVAKGEP    39
tr|A0A7G2C3S2|A0A7G2C3S2_9TRYP      -------------------------------------------------ENTEAIQKGSP    68
sp|P14625|ENPL_HUMAN                ------DDEVDVDGTVEEDLGKSREGSRTDDEVVQREEEAIQLDGLNASQIRELREKSEK    75
tr|Q7T3L3|Q7T3L3_DANRE              ------DDDVDIDGTVEEDLGKSRDGSRTDDEVVQREEEAIQLDGLNTSQLKEIRDKAEK    75
sp|Q9STX5|ENPL_ARATH                EES---SDDVTDPPKVEEKIGGH-GGLSTDSDVVHRESESMS--------KKTLRSNAEK    78
tr|A0A061FKM2|A0A061FKM2_THECC      D------EGAVDPPKVEEKIGAVPNGLQTDSDVVKRESESIS--------SRSLRSNAEK    76
tr|Q5Z9N8|Q5Z9N8_ORYSJ              DDS---TDELVDPPKVEEKIGGVPHGLSTDSEVVQREAESIS--------RKTLRSSAEK    80
tr|W2ZJ09|W2ZJ09_PHYPR              STGSDEVLGRST-PS-----------L----------LVDDL--------NDEPLTGGET    62
tr|W2ZJ74|W2ZJ74_PHYPR              STGSDEVLG-ST-PS-----------L----------LVDDL--------NDEPLTGGET    61
tr|F0Y644|F0Y644_AURAN              --------------S-----------L----------GTVAY--------AEDAGAAPET    34
tr|D8LHY7|D8LHY7_ECTSI              SSGLARAEEAVAPAS-----------T----------MDAAF--------EEQLKDGAEA    56


tr|A0A0L1KBS8|A0A0L1KBS8_9EUGL      HSYQAEVSKMLEILIHSLYTNRSIFLREIISNASDALDKIKYMYLTNPKSPKNKDGEEPT    92
tr|A0A0L1KHT0|A0A0L1KHT0_9EUGL      HKYKAEVGKMLDILINSLYTNHSIFLREIISNASDALDKVRFLYLTKPKNPKNDAGKEPT    93
tr|A0A1X0P079|A0A1X0P079_9TRYP      VAFQAEVSKMLDILINSLYTNRAVFLRELISNGSDALDKIRMLYLTAPKEPVNKDGVAPT    99
tr|Q4DW89|Q4DW89_TRYCC              ISFQAEVSKMLDILINSLYTNRAVFLRELISNGSDALDKIRMLYLTAPKEPKNKDGEVPA    97
tr|A0A422PWD8|A0A422PWD8_9TRYP      ISFQAEVSKMLDILINSLYTNRAVFLRELISNGSDALDKIRMLYLTAPKEPKNKDGEVPA    98
tr|A0A422NRM6|A0A422NRM6_TRYRA      ISFQAEVSKMLDILINSLYTNRAVFLRELISNGSDALDKIRMLYLTAPKEPKNKDGEVPA    98
tr|A0A0S4KLS2|A0A0S4KLS2_BODSA      ITFQAEVSKMLDILINSLYTNRNIFLREIISNASDALDKIRFFYLTTPREPTNANGEAPT    91
tr|A0A0N0VF86|A0A0N0VF86_LEPPY      ITFQAEVSKMLDILVNSLYTNRAIFLRELISNGSDALDKIRVLYLTAPKEPLNADGVSPT    92
tr|A4HH83|A4HH83_LEIBR              ITFQAEVSKMLDILVNSLYTNHAVFLRELISNGSDALDKIRVLYLTSPKEPLTKDGETPT    91
tr|E9ADS8|E9ADS8_LEIMA              IAFQAEVSKMLDILVNSLYTNRAVFLRELISNGSDALDKIRVLYLTSPKEPLTKDGEAPT    91
sp|Q9NGD0|ENPL_LEIIN                IAFQAEVSKMLDILVNSLYTNRAVFLRELISNGSDALDKIRVLYLTSPKEPLTKDGEAPT    91
tr|A4I4C9|A4I4C9_LEIIN              IAFQAEVSKMLDILVNSLYTNRAVFLRELISNGSDALDKIRVLYLTSPKEPLTKDGEAPT    91
sp|Q8T7E0|ENPL_LEIDO                IAFQAEVSKMLDILVNSLYTNRAVFLRELISNGSDALDKIRVLYLTSPKEPLTKDGEAPT    91
tr|S9U4L8|S9U4L8_9TRYP              TTFTAEVSKMLDILINSLYTNRAIFLRELISNGSDALDKVRVLYLTDPKNPVNKDGEPVT    99
tr|A0A7G2C3S2|A0A7G2C3S2_9TRYP      TTFTAEVSKMLDILINSLYTNRAIFLRELISNGSDALDKVRVTYLTEPKDPRNDAGEAPT    128
sp|P14625|ENPL_HUMAN                FAFQAEVNRMMKLIINSLYKNKEIFLRELISNASDALDKIRLISLTDENA--LSG--NEE    131
tr|Q7T3L3|Q7T3L3_DANRE              HAFQAEVNRMMKLIINSLYKNKEIFLRELISNASDALDKIRLLSLTNEDA--LAG--NEE    131
sp|Q9STX5|ENPL_ARATH                FEFQAEVSRLMDIIINSLYSNKDIFLRELISNASDALDKIRFLALTDKDV--LGEGDTAK    136
tr|A0A061FKM2|A0A061FKM2_THECC      FEFQAEVSRLMDIIINSLYSNKDIFLRELISNASDALDKIRFLSLTDKEV--LGEGDTAK    134
tr|Q5Z9N8|Q5Z9N8_ORYSJ              FEFQAEVSRLMDIIINSLYSNKDIFLRELISNASDALDKIRFLALTDKEV--LGEGDTAK    138
tr|W2ZJ09|W2ZJ09_PHYPR              FAFQAEVSRLMDILINSLYRTKEIFLRELISNASDALDKIRFLALSNNEL--LGK--LRD    118
tr|W2ZJ74|W2ZJ74_PHYPR              FAFQAEVSRLMDILINSLYRTKEIFLRELISNASDALDKIRFLALSNNEL--LGK--LRD    117
tr|F0Y644|F0Y644_AURAN              FEFQAEVNRLMDIIINSLYKNKEIFLREVISNGSDALDKIRFLAVSDAAA--LDT--KKE    90
tr|D8LHY7|D8LHY7_ECTSI              FEFQAEVNRLMDIIINSLYKNKDIFLRELISNASDALDKIRFLSVSDPDK--LGT--AKD    112
                                    : ***.:::.::::*** .: :****:***.******::    ::

tr|A0A0L1KBS8|A0A0L1KBS8_9EUGL      FDIHLKVDHEKRTLTLRDGGVGMTKDELSAHLGSLGASGTKKFVEKLDDTESKTQDDVNT    152
tr|A0A0L1KHT0|A0A0L1KHT0_9EUGL      MDVRILIDHETNELRLIDGGVGMTKSELSANLGSLGASGTKSFWENINESKNA-GEELSN    152
tr|A0A1X0P079|A0A1X0P079_9TRYP      LDMHVSIDPSNKTLTLRDGGIGMTREELENHLGSLGTSGTKRFMEKLQES------KDAS    153
tr|Q4DW89|Q4DW89_TRYCC              LEMRVIIDNERKTLTLRDGGIGMTKAELEEHLGSLGTSGTKRFMEKLKET------KDDS    151
tr|A0A422PWD8|A0A422PWD8_9TRYP      LEMRVTVDAERKTLTLRDGGIGMTREELEEHLGSLGTSGTKRFLERLKET------KDAS    152
tr|A0A422NRM6|A0A422NRM6_TRYRA      LEMRVTVDTERKTLTLRDGGIGMTRAELEEHLGSLGTSGTKRFMDRLKET------KDAS    152
tr|A0A0S4KLS2|A0A0S4KLS2_BODSA      MDIRIVVDREKRLFIMRDGGVGMTKEELAANLGSLGSSGTKRFLEKMKDS------SDAN    145
tr|A0A0N0VF86|A0A0N0VF86_LEPPY      MDIRISFNKEKSELVIRDGGVGMNKDDLAGHLGSLGTSGTKRFLEKLQEGG--TAGDQNN    150
tr|A4HH83|A4HH83_LEIBR              MDLRISFDNENHELILRDGGIGMTKEELTQHLGSLGSSGTKHFLEKLQEGSGAVGGDQSN    151
tr|E9ADS8|E9ADS8_LEIMA              MDLRISFDKEKSELILRDGGVGMTKEELAKHLGSLGTSGTKHFLEKLQEGVGAGGGDQNN    151
sp|Q9NGD0|ENPL_LEIIN                MDLRISFDKEKSELILRDGGVGMTKEELAKHLGSLGTSGTKHFLEKLQEGVGAGGGDQNN    151
tr|A4I4C9|A4I4C9_LEIIN              MDLRISFDKEKSELILRDGGVGMTKEELAKHLGSLGTSGTKHFLEKLQEGVGAVGGDQNN    151
sp|Q8T7E0|ENPL_LEIDO                MDLRISFDKEKSELILRDGGVGMTKEELAKHLGSLGTSGTKHFLEKLQEGVGAVGGDQNN    151
tr|S9U4L8|S9U4L8_9TRYP              MDIHISIDKEQKQLIIRDGGIGMTKEDLSKNLGSLGTSGTKRFLEGMKSG--ATSGDSSN    157
tr|A0A7G2C3S2|A0A7G2C3S2_9TRYP      MDIHISVSPETKELIIRDGGVGMTKEELGTHLGSLGTSGTKRFLENLKAS----STDNNN    184
sp|P14625|ENPL_HUMAN                LTVKIKCDKEKNLLHVTDTGVGMTREELVKNLGTIAKSGTSEFLNKMTEAQED-GQSTSE    190
tr|Q7T3L3|Q7T3L3_DANRE              LTIKIKSDKEKNMLHITDTGIGMTKEELVKNLGTIAKSGTSEFLNKMTEVQDD-SQSTSE    190
sp|Q9STX5|ENPL_ARATH                LEIQIKLDKAKKILSIRDRGIGMTKEDLIKNLGTIAKSGTSAFVEKMQ------SSGDLN    190
tr|A0A061FKM2|A0A061FKM2_THECC      LEIQIKLDKEKKILTLRDRGIGMTKEDLIKHLGTIAKSGTSAFVEKMQ------SSGDLN    188
tr|Q5Z9N8|Q5Z9N8_ORYSJ              LEIQIKLDKEKKILSIRDRGIGMTKEDLIKNLGTIAKSGTSAFVEKMQ------TGGDLN    192
tr|W2ZJ09|W2ZJ09_PHYPR              LEIRISFDKDAGTLTIRDTGVGMTKDDLVNNLGTVAKSGTANFVEAMQA---G--ADDSS    173
tr|W2ZJ74|W2ZJ74_PHYPR              LEIRISFDKDAGTLTIRDTGVGMTKDDLVNNLGTVAKSGTANFVEAMQA---G--ADDSS    172
tr|F0Y644|F0Y644_AURAN              LEIRISFDKDARTLTIQDSGVGMTKADLVANLGTVAKSGTTNFVEAMSG---D-ASGDLS    146
tr|D8LHY7|D8LHY7_ECTSI              LEIRISADKDARTLTIRDAGVGMTKADLISNLGTVARSGTTNFVEALT------ETGDLG    166
                                    : :::  .          : : * *:**.: :*  :**::. ***  * : :

tr|A0A0L1KBS8|A0A0L1KBS8_9EUGL      LIGQFGVGFYSVFLVADTIKVISKSDNSDEQWIWESNGDGKFQLYEDPNGNTLGRGTELV    212
tr|A0A0L1KHT0|A0A0L1KHT0_9EUGL      LIGQFGVGFYSVFLVADEVRVASKNDAEEKQWMWESKGDGKYFLYEDPRGNTLGRGTELV    212
tr|A0A1X0P079|A0A1X0P079_9TRYP      LIGQFGVGFYSAFLVADRVRVASKSDDSDKQWVWESAGDGQYYIYEDERGNTLGRGTEIT    213
tr|Q4DW89|Q4DW89_TRYCC              LIGQFGVGFYSAFLVADRVRVASKSDDSDVQWVWESAGDGQYYIYEDERGNTLGRGTEIT    211
tr|A0A422PWD8|A0A422PWD8_9TRYP      LIGQFGVGFYSAFLVAERVRVASKSDDNDTQWVWESAGDGQYFLYEDERGNTLGRGTEVT    212
tr|A0A422NRM6|A0A422NRM6_TRYRA      LIGQFGVGFYSAFLVAERVRVASKSDDSDTQWVWESTGDGQYYVYEDERGNTLGRGTEIT    212
tr|A0A0S4KLS2|A0A0S4KLS2_BODSA      FIGQFGVGFYSVFLVADKVRVASKHDDSEKQWVWESTGDGTFFLYEDERGNTLGRGSELT    205
tr|A0A0N0VF86|A0A0N0VF86_LEPPY      LIGQFGVGFYSVFLVGDRVRVASKSDDGDEQYVWESSGNGQYFLYPDPRGNTLGRGTEIT    210
tr|A4HH83|A4HH83_LEIBR              LIGQFGVGFYSVFLVGNRVRVASKSDDSDEQYVWESKGDGEYFLYPDPRGNTLGRGTEIT    211
tr|E9ADS8|E9ADS8_LEIMA              LIGQFGVGFYSVFLVGDRVRVASKSDDSDEQYVWESKGDGQYFLYPDPRGNTLGRGTEIT    211
sp|Q9NGD0|ENPL_LEIIN                LIGQFGVGFYSVFLVGDRVRVASKSDDSDEQYVWESKGDGQYFLYPDPRGNTLGRGTEIT    211
tr|A4I4C9|A4I4C9_LEIIN              LIGQFGVGFYSVFLVGDRVRVASKSDDSDEQYVWESKGDGQYFLYPDPRGNTLGRGTEIT    211
```

```
sp|Q8T7E0|ENPL_LEIDO                LIGQFGVGFYSVFLVGDRVRVASKSDDSDEQYVWESKGDGQYFLYPDPRGNTLGRGTEIT    211
tr|S9U4L8|S9U4L8_9TRYP              LIGQFGVGFYSVFLVAKRVIVASKHDSSDVQYVWESHGDGRFFLYEDPRGNTLGRGTELR    217
tr|A0A7G2C3S2|A0A7G2C3S2_9TRYP      LIGQFGVGFYSVFLVGKNVRVASKSDADSKQYVWESSGNGQFFLYEDPRGNTLGRGTEIR    244
sp|P14625|ENPL_HUMAN                LIGQFGVGFYSAFLVADKVIVTSKHNN-DTQHIWESDSN-EFSVIADPRGNTLGRGTTIT    248
tr|Q7T3L3|Q7T3L3_DANRE              LIGQFGVGFYSAFLVADKVIVTSKHNN-DTQHMWESDSN-QFSVIEDPRGDTLGRGTTIT    248
sp|Q9STX5|ENPL_ARATH                LIGQFGVGFYSAYLVADYIEVISKHND-DSQYVWESKANGKFAVSEDTWNEPLGRGTEIR    249
tr|A0A061FKM2|A0A061FKM2_THECC      LIGQFGVGFYSVYLVADYVEVISKHND-DKQYVWESKADGAFAISEDTWNEPLGRGTEIR    247
tr|Q5Z9N8|Q5Z9N8_ORYSJ              LIGQFGVGFYSVYLVADYVEVISKHND-DKQHVWESKADGSFAISEDTWNEPLGRGTEIR    251
tr|W2ZJ09|W2ZJ09_PHYPR              LIGQFGVGFYSVYLVADRVRVVSKNNN-DDQYIWESDANASFTITKDPRGDTLGRGTEIT    232
tr|W2ZJ74|W2ZJ74_PHYPR              LIGQFGVGFYSVYLVADRVRVVSKNNN-DDQYIWESDANASFTITKDPRGDTLGRGTEIT    231
tr|F0Y644|F0Y644_AURAN              LIGQFGVGFYSVYLVADRVRVRSKHND-DDQHVWESAADSSFTVWQDPLGDTLGRGTEIK    205
tr|D8LHY7|D8LHY7_ECTSI              LIGQFGVGFYSVYLVADKVQVISKSND-DDQYVWESTADSTFTVSKDPRGNTLGRGTEIV    225
                                    :**********.:**.. : * ** :  . * :*** .:  : :  *  .: ****: :

tr|A0A0L1KBS8|A0A0L1KBS8_9EUGL      IYLRKDADEYMEEFKLKEIIHRYSQFVHFPIYLWNVKQKEKPKAKDENEEDSEDIVID--    270
tr|A0A0L1KHT0|A0A0L1KHT0_9EUGL      LKLKKEAEEYLQEEKLKKIIHYYSEFINFPIYVGYATKDEREKQ----------------    256
tr|A0A1X0P079|A0A1X0P079_9TRYP      LEMKPDALEFLNSDNVRRIVHQYSEFVHFPIYMEKG------------------------    249
tr|Q4DW89|Q4DW89_TRYCC              LEMKPDALEFLSTDNVRDIVHQYSEFVHFPIYMQKG------------------------    247
tr|A0A422PWD8|A0A422PWD8_9TRYP      LEMKPDALEFLNIDGVRKIVHQYSEFVHFPIYMQKG------------------------    248
tr|A0A422NRM6|A0A422NRM6_TRYRA      LEMKPDALEFLSADNVRHIVHQYSEFVHFPIYMQKE------------------------    248
tr|A0A0S4KLS2|A0A0S4KLS2_BODSA      LELKKDADEYLDIDKVKEAIHKYSEFIHFPIYIQTTKTEKVKKAAEAESTEAK----E--    259
tr|A0A0N0VF86|A0A0N0VF86_LEPPY      IELKPDAEQFLSAETIKKTVHQYSEFINFPIYVEEEVTVEAAKKED--------ADTE--    260
tr|A4HH83|A4HH83_LEIBR              IELKPEDQEFLSAETIKKTIHQYSEFINFPIYVQEEVEVKPKTEIRTSEPGAEVGFMK--    269
tr|E9ADS8|E9ADS8_LEIMA              IELKPDAEQFLSAETIKKTIHQYSEFINFPIYVQEEVEVASTA--ATPEPAAEEGSLD--    267
sp|Q9NGD0|ENPL_LEIIN                IELKPDAEQFLSAETIKKTIHQYSEFINFPIYVQEEVEVASTA--ATPEPAAEEGSLD--    267
tr|A4I4C9|A4I4C9_LEIIN              IELKPDAEQFLSAETIKKTIHQYSEFINFPIYVQEEVEVASTA--ATPESAAEERSLD--    267
sp|Q8T7E0|ENPL_LEIDO                IELKPDAEQFLSAETIKKTIHQYSEFINFPIYVQEEVEVASTA--ATPESAAEERSLD--    267
tr|S9U4L8|S9U4L8_9TRYP              IELKPDSIEYLNMDRIKETIHQYSEFIDFPIHVQEQMAEDEVVKELQ----------A--    265
tr|A0A7G2C3S2|A0A7G2C3S2_9TRYP      IELKSDAVEYLKTDRVKKTIHQYSEFIDFPIYLEEAEKDNDQVQKEI----------E--    292
sp|P14625|ENPL_HUMAN                LVLKEEASDYLELDTIKNLVKKYSQFINFPIYVWSSKTETVEEPMEEEEAAKEEKEESD-    307
tr|Q7T3L3|Q7T3L3_DANRE              LVMKEEASDYLELETIKNLVKKYSQFINFPIYVWSSKTETVEEPIEDEAEAEKEEATED-    307
sp|Q9STX5|ENPL_ARATH                LHLRDEAGEYLEESKLKELVKRYSEFINFPISLWASKEVETEVPVEEDESADEETETTST    309
tr|A0A061FKM2|A0A061FKM2_THECC      LHLRDEAQEYLEEGKLKELVKKYSEFINFPIYIWASKEVDVEVPAEEDESSDEEETSDST    307
tr|Q5Z9N8|Q5Z9N8_ORYSJ              LHLRDEAKEYVEEDKLKDLVKKYSEFINFPIYLWATKEVDVEVPADEDESSESSEEEESS    311
tr|W2ZJ09|W2ZJ09_PHYPR              LFLKPDATEFQDQDKLKNLVGHYSEFITFPIYVNTTSTETYEVEEEPADVEEVVEEETDE    292
tr|W2ZJ74|W2ZJ74_PHYPR              LFLKPDATEFQDQDKLKNLVGHYSEFITFPIYVNTTSTETYEVEEEPADVEEVVEEETDE    291
tr|F0Y644|F0Y644_AURAN              LFLKEDAGEFLDQDRLEELVLRYSEFITFPIYLYKSKTETVEVDDDEDEDEEDLDEEEEE    265
tr|D8LHY7|D8LHY7_ECTSI              LHLKEDAGEFLSESTLKNLIHRYSEFITFPIYQLVEKEEEIEVEDDEEEEEDDGDEDSGD    285
                                    : :: :  :: .   :.  :  **:*: ***

tr|A0A0L1KBS8|A0A0L1KBS8_9EUGL      --------EENEDEKQEEVVSEEKNEPEVEQWVWEHVNEHEPIWIRDPAEVKEDDYVSFY    322
tr|A0A0L1KHT0|A0A0L1KHT0_9EUGL      -------------------------DEKYQGIDWELVNVIKPIWTRNPQTVERKEYVEFY    291
tr|A0A1X0P079|A0A1X0P079_9TRYP      -------------------------------DKWEVVNENKPIWTRKPSEVTEEEYVKFY    278
tr|Q4DW89|Q4DW89_TRYCC              -------------------------------EKWEVVNENKPIWTRKPSEVTEEEYHKFY    276
tr|A0A422PWD8|A0A422PWD8_9TRYP      -------------------------------DKWEVVNENKPIWTRKPAEVAADEYHKFY    277
tr|A0A422NRM6|A0A422NRM6_TRYRA      -------------------------------DKWEVVNENKPIWTRKPSEVTPDEYHKFY    277
tr|A0A0S4KLS2|A0A0S4KLS2_BODSA      --------DGDDE---APAEEKKVEEEEVVTHDWELINENKPIWTRKAVEITDAEYNSFF    308
tr|A0A0N0VF86|A0A0N0VF86_LEPPY      --------DKALD---EDEVEDEVEAVPVTERKWTLVNENRPIWTRPIGNVTEAEYNKFY    309
tr|A4HH83|A4HH83_LEIBR              --------EVVVE---EDNN--GEEKVPRMEKRWTLINENRPIWTRQIGNVTEEEYIKFY    316
tr|E9ADS8|E9ADS8_LEIMA              --------EGAVE---EDSDKEGGTQGVAKERRWVLVNENRPIWTRPIGNVTEEEYHKFY    316
sp|Q9NGD0|ENPL_LEIIN                --------EGAVE---EDPDKEGDTQGVVKERRWTLVNENRPIWTRPIGNVTEEEYHTFY    316
tr|A4I4C9|A4I4C9_LEIIN              --------EGAVE---EDPDKEGDTQGVVKEKRWTLVNENRPIWTRPIGNVTEEEYHKFY    316
sp|Q8T7E0|ENPL_LEIDO                --------EGAVE---EDPDKEGDTQGVVKEKRWTLVNENRPIWTRPIGNVTEEEYHKFY    316
tr|S9U4L8|S9U4L8_9TRYP              --------QGMVP---E----EVEAASKEPMYRWTLVNKNRPIWTRSFDEVKEADYDSFY    310
tr|A0A7G2C3S2|A0A7G2C3S2_9TRYP      --------YG------E------DVENTTGNKTWTLVNKNKPIWTRPTSEVTEEEYDSFY    332
sp|P14625|ENPL_HUMAN                --------DEAAVEEEEEEKKPKTKKVEKTVWDWELMNDIKPIWQRPSKEVEEDEYKAFY    359
tr|Q7T3L3|Q7T3L3_DANRE              --------EAEVEEEEEDKDKPKTKKVEKTVWDWELMNDIKPIWQRPAKEVEEDEYTAFY    359
sp|Q9STX5|ENPL_ARATH                E-EEK----EEDAEEEDGEKKQKTKKVKETVYEWELLNDVKAIWLRSPKEVTEEEYTKFY    364
tr|A0A061FKM2|A0A061FKM2_THECC      S-EEGEDEDAEKSEDEGTEKKPKTKKVKETSNEWELLNDVKAIWLRSPKEVTEEEYTKFY    366
tr|Q5Z9N8|Q5Z9N8_ORYSJ              P-ESTEEE----ETEESEEKKPKTKTVKETTTEWELLNDVKAIWLRSPKEVTEEEYTKFY    366
tr|W2ZJ09|W2ZJ09_PHYPR              E-KPSEEDEELEAVEEEDVKTPKTRAETRTVWKWERVNEVKAIWTRSKDEISDEEYESFY    351
tr|W2ZJ74|W2ZJ74_PHYPR              E-KPSEEDEELEAVEEEDVKTPKTRAETRTVWKWERVNEVKAIWTRSKDEISDEEYESFY    350
tr|F0Y644|F0Y644_AURAN              ESEEDLEDDEYE-----EEEDDEPRTTTVTNWDWHQVNSESAIWSRDASDVDDEEYRSFY    320
tr|D8LHY7|D8LHY7_ECTSI              E--DGDEEDEFE-----EV-EVDVKYKTVKTLDWERVNANVAVWARDKDEITDEEYHNFY    337
                                                                        *  :*   :* *     :   :*  *:

tr|A0A0L1KBS8|A0A0L1KBS8_9EUGL      KSMSHE--YEDPMHHYHFASKGDDVGFKSILYVPGRAPHDGYNSK-KGASHDNIRLYVRR    379
tr|A0A0L1KHT0|A0A0L1KHT0_9EUGL      RGFNKE--FDEPMYYSHFVASGGEIDFKALLYVPSTLPTNAFNVHD-ESSYKNMRLYVRR    348
tr|A0A1X0P079|A0A1X0P079_9TRYP      KALTHD--YRSPMYYSHFTVE-GEVEFSSVLFIPEEATQDIFINNE--NTRDNIKLYVRR    333
tr|Q4DW89|Q4DW89_TRYCC              KSLTHD--YRNPMYYSHFNVE-GEVEFSSVLFIPQEASQDIFVNNE--DTRDNIKLYVRR    331
tr|A0A422PWD8|A0A422PWD8_9TRYP      KALTHD--YRDPMYYSHFVVE-GEVEFSSVLFIPQEASQDIFVNNE--NTRDNIKLYVRR    332
tr|A0A422NRM6|A0A422NRM6_TRYRA      KTLTRD--YRDPMYYSHFMVE-GEVEFSSVLFIPQEASQDIFVNNE--NTRDNIKLYVRR    332
tr|A0A0S4KLS2|A0A0S4KLS2_BODSA      KSLTKD--YDDPMFYTHFSAE-GEVEFRSILFIPSHSNTNVF---DTSVVQANIRLYVRR    362
tr|A0A0N0VF86|A0A0N0VF86_LEPPY      KAFSGD--YRDPLYFNHFRVE-GEVEFDSILFVPAAVDPATF-SDDNTVPNTNIKLYVRR    365
tr|A4HH83|A4HH83_LEIBR              KAFSGD--YRDPLYFSHFKVE-GEVDFDSILFIPSTVDPASF-SDDKANPSTNIKLYVRR    372
tr|E9ADS8|E9ADS8_LEIMA              KAFSGD--YRDPLYFSHFKVE-GEVDFDSILFVPTTVDPASF-SDDNAAPNTNIKLYVRR    372
sp|Q9NGD0|ENPL_LEIIN                KAFSGD--YRDPLYFNHFKVE-GEVDFDSILFVPTTVDPASF-SDDNSVPNTNIKLYVRR    372
tr|A4I4C9|A4I4C9_LEIIN              KAFSGD--YRDPLYFNHFKVE-GEVDFDSILFVPTTVDPASF-SDDNAVPNTNIKLYVRR    372
sp|Q8T7E0|ENPL_LEIDO                KAFSGD--YRDPLYFNHFKVE-GEVDFDSILFVPTTVDPASF-SDDNAVPNTNIKLYVRR    372
tr|S9U4L8|S9U4L8_9TRYP              KALTHD--YREPLYYSHFKVE-GEVEFDSILFIPTEASEEEL---DSTQRTKNIKLYVRR    364
tr|A0A7G2C3S2|A0A7G2C3S2_9TRYP      KAVSSD--WRTPMYYSHFHVE-GEVEFDSILYIPEIINREEQATNDASMLNRNIKLYVRR    389
sp|P14625|ENPL_HUMAN                KSFSKE--SDDPMAYIHFTAE-GEVTFKSILFVPTSAPRGLFDEYG-SKKSDYIKLYVRR    415
tr|Q7T3L3|Q7T3L3_DANRE              KTFSRD--TDEPLSHIHFTAE-GEVTFKSILFVPASAPRGLFDEYG-TKKNDFIKLFVRR    415
sp|Q9STX5|ENPL_ARATH                HSLSKDFTDEKPMAWSHFNAE-GDVEFKAVLYVPPKAPHDLYESYY-NSNKANLKLYVRR    422
tr|A0A061FKM2|A0A061FKM2_THECC      QSLAKDFGDEKPLAWSHFTAE-GDVEFKAVLFVPPKAPHDLYESYY-NTNKANLKLYVRR    424
tr|Q5Z9N8|Q5Z9N8_ORYSJ              HSLAKDFGDDKPLSWSHFTAE-GDVEFKALLFVPPKAPHDLYESYY-NSNKSNLKLYVRR    424
tr|W2ZJ09|W2ZJ09_PHYPR              HSLQKTD-ITDPLTWIHFQAE-GEIEFKSILYVPGQAPRDLYTRFE-NK-KADIKLYVRK    407
tr|W2ZJ74|W2ZJ74_PHYPR              HSLQKTD-ITDPLTWIHFQAE-GEIEFKSILYVPGQAPRDLYTRFE-NK-KADIKLYVRK    406
```

```
tr|F0Y644|F0Y644_AURAN             KTLSK-D-TADPTTWIHFKAE-GEVEFKSILFVPGQVPFDMYDSYH-TK-SAQLRLYVRK    375
tr|D8LHY7|D8LHY7_ECTSI             KALSG-D-TSDAATWIHFKAE-GEVEFKSILFAPSKAPHNMYDHYY-DDSRGGLKLYVRK    393
                                   : .              **  . .:: * ::*: *              ::*:**:

tr|A0A0L1KBS8|A0A0L1KBS8_9EUGL     IFITDEFKDLLPRYLSFVRGVVDSDDLPLNVSREILQESKIIKLIQKKLVRKCIDMFVEM    439
tr|A0A0L1KHT0|A0A0L1KHT0_9EUGL     VFITDEFKDLFPRYLNFVQGVVDSDDLPLNVSRELLQENRVLKTIKKQLVRKALGMFIEI    408
tr|A0A1X0P079|A0A1X0P079_9TRYP     IFITDEFRELLPRYLSFVRGVVDSNDLPLNVSREVLQESRILRVIKKKLVRKALGMISEI    393
tr|Q4DW89|Q4DW89_TRYCC             IFITDEFRQLLPRYLSFVRGIVDSNDLPLNVSREVLQESRILRVIKKKLVRKALSMISEI    391
tr|A0A422PWD8|A0A422PWD8_9TRYP     IFITDEFRELLPRYLNFVRGVVDSNDLPLNVSREVLQESRILRVIKKKLVRKALSMIAEI    392
tr|A0A422NRM6|A0A422NRM6_TRYRA     IFITDEFRELLPRYLSFVRGVVDSNDLPLNVSREVLQESRILRVIKKKLVRKALSMIAEI    392
tr|A0A0S4KLS2|A0A0S4KLS2_BODSA     VFITDDFRDLLPRYLNFIKGVVDSDDLPLNVSREVLQESRILRVIKKKLVRKALAMIADI    422
tr|A0A0N0VF86|A0A0N0VF86_LEPPY     VFITDEFRDLLPRYLNFVKGIVDSNDLPLNVSREVLQESRILRVIKKKLVRKALTMFSDI    425
tr|A4HH83|A4HH83_LEIBR             VFITDEFRDLLPRYLNFVKGIVDSNDLPLNVSREVLQESRILRVIKKKLVRKALNMFANI    432
tr|E9ADS8|E9ADS8_LEIMA             VFITDEFRDLLPRYLNFVKGIVDSNDLPLNVSREVLQESRILRVIKKKLVRKTLSMFADI    432
sp|Q9NGD0|ENPL_LEIIN               VFITDEFRDLLPRYLNFVKGIVDSNDLPLNVSREVLQESRILRVIKKKLVRKTLSMFADI    432
tr|A4I4C9|A4I4C9_LEIIN             VFITDEFRDLLPRYLNFVKGIVDSNDLPLNVSREVLQESRILRVIKKKLVRKTLSMFADI    432
sp|Q8T7E0|ENPL_LEIDO               VFITDEFRDLLPRYLNFVKGIVDSNDLPLNVSREVLQESRILRVIKKKLVRKTLSMFADI    432
tr|S9U4L8|S9U4L8_9TRYP             VFITDDFKDLLPRYLSFVKGVVDSNDLPLNVSRELLQESRVLQVIKKKLVRKALAMFADI    424
tr|A0A7G2C3S2|A0A7G2C3S2_9TRYP     VFITDEFKDLLPHYLNFVKGVVDSNDLPLNVSREVLQESRVLRVIKKKLVRKVLSMIADI    449
sp|P14625|ENPL_HUMAN               VFITDDFHDMMPKYLNFVKGVVDSDDLPLNVSRETLQQHKLLKVIRKKLVRKTLDMIKKI    475
tr|Q7T3L3|Q7T3L3_DANRE             VFITDDFHDMMPKYLNFIKGVVDSDDLPLNVSRETLQQHKLLKVIRKKLVRKTLDMIKKI    475
sp|Q9STX5|ENPL_ARATH               VFISDEFDELLPKYLSFLKGLVDSDTLPLNVSREMLQQHSSLKTIKKKLIRKALDMIRKL    482
tr|A0A061FKM2|A0A061FKM2_THECC     VFISDEFDELLPKYLNFLKGLVDSDTLPLNVSREMLQAHSSLKTIKKKLIRKALDMIRKI    484
tr|Q5Z9N8|Q5Z9N8_ORYSJ             VFISDEFDELLPKYLSFLKGLVDSDTLPLNVSREMLQQHSSLKTIKKKLIRKALDMIRKL    484
tr|W2ZJ09|W2ZJ09_PHYPR             VLITDDFDDFLPRYLNFIAGVVDSDDLPINVSRETLQENKILRVIRKKLVRKVLEMLRKL    467
tr|W2ZJ74|W2ZJ74_PHYPR             VLITDDFDDFLPRYLNFIAGVVDSDDLPINVSRETLQENKILRVIRKKLVRKVLEMLRKL    466
tr|F0Y644|F0Y644_AURAN             VLITDEFEDLVPRYLNFLRGVVDSDDLPLNVSRETLQQHKVLKVMGKKLVRKALEMLRKL    435
tr|D8LHY7|D8LHY7_ECTSI             VLITDEFDELLPKYLSFIRGVVDSDDLPLNVSREMLQEHKVLKVMAKKLVRKTLEMLRKL    453
                                   ::*:*:* ::.*:**.*: *:***: **:***** **       :: : ::*:** : *: .:

tr|A0A0L1KBS8|A0A0L1KBS8_9EUGL     AEDESGIKQKRKLRKK----------------------KREDGGEEFSYSKFWREFGKS    476
tr|A0A0L1KHT0|A0A0L1KHT0_9EUGL     SKEDDEVWKEIDATE---------------------DKELKEKKFSELKYPAFWDEYGKS    447
tr|A0A1X0P079|A0A1X0P079_9TRYP     AETDARIKDEMKDKPK-------DE-------NKT-EDKGSIKHPKEPLYPKFWAQFGKH    438
tr|Q4DW89|Q4DW89_TRYCC             AEKDARLKEGLEKEKSGEDGEAKDE-------NTTEKKS-DDDKGKEPLYPKFWAQFGKH    443
tr|A0A422PWD8|A0A422PWD8_9TRYP     AENDSRLKERLAKEKAEQKAT------------DAKDGDGDGKAKEEPMYPKFWAQFGKH    440
tr|A0A422NRM6|A0A422NRM6_TRYRA     GENDARLKERLEKEKTAAVEEQADK-------NATETKDDDDKSKKEPMYPKFWAQFGKH    445
tr|A0A0S4KLS2|A0A0S4KLS2_BODSA     AASDKKLEAAKDDDEAAEAEEK------------KDDVTAGNKQLKASTYPKFWEEYGKN    470
tr|A0A0N0VF86|A0A0N0VF86_LEPPY     AEQDEAIAKGT----QSES----------------PAP-SGHTHLTQPTYAKFWELFGKH    464
tr|A4HH83|A4HH83_LEIBR             AAQDVEIANKD----HAED----------------PAP-SGHTHLKKPTYAKFWDLFGKH    471
tr|E9ADS8|E9ADS8_LEIMA             AAQDEAIADGK----QVEN----------------PAL-SGHTHLKKPAYTKFWELYGKH    471
sp|Q9NGD0|ENPL_LEIIN               AAQDEAIANGK----QVES----------------PAP-SGHTHLKKPAYTKFWELYGKH    471
tr|A4I4C9|A4I4C9_LEIIN             AAQDEAIANGK----QVEN----------------PAP-SGHTHLKKPAYTKFWELYGKH    471
sp|Q8T7E0|ENPL_LEIDO               AAQDEAIANGK----QVEN----------------PAP-SGHTHLKKPAYTKFWELYGKH    471
tr|S9U4L8|S9U4L8_9TRYP             QAQDDRIAVLKKEGREVET----------------DAPNKGVRVLTKPTYKTFWELFGRH    468
tr|A0A7G2C3S2|A0A7G2C3S2_9TRYP     QVQDDNLRKLKEENKTFE-----------------PAKGTGVRQLKVETYPKFWETYGRF    492
sp|P14625|ENPL_HUMAN               ADDK-------------------------------------------YNDTFWKEFGTN    491
tr|Q7T3L3|Q7T3L3_DANRE             AEEQ-------------------------------------------YNDKFWKEFGTN    491
sp|Q9STX5|ENPL_ARATH               AEEDPDEIHDDEKKDVEK--------------------SGE-NDEKKGQYTKFWNEFGKS    521
tr|A0A061FKM2|A0A061FKM2_THECC     AEEDPDESSGKDEKEVEK--------------------SGTDDDEKKGQYAKFWNEFGKS    524
tr|Q5Z9N8|Q5Z9N8_ORYSJ             AEEDPDEYSNKDKTDEEK--------------------SA--MEEKKGQYAKFWNEFGKS    522
tr|W2ZJ09|W2ZJ09_PHYPR             AEKDEDDDEEDEEDDEVVDTSADESNDKKETEGDAEKEEEEEEEDGNTAYNKFWEEFGKN    527
tr|W2ZJ74|W2ZJ74_PHYPR             AEKDEDDDEEDEEDDEVVDTSADESNDKKETEGDAEKEEEEEEEDGNTAYNKFWEEFGKN    526
tr|F0Y644|F0Y644_AURAN             SQLKEKGDDDEDEDDEES------------------EDADESEDDDEDPYIGFWSKFGKN    477
tr|D8LHY7|D8LHY7_ECTSI             ATEE----------------------------------VEDEEGEEEHPYIQFWDEFGKS    479
                                   .                                                 **  :*

tr|A0A0L1KBS8|A0A0L1KBS8_9EUGL     IRLGIIEDSSNRQRLARLVRYPSSTTEFKATESSTSLNEYISRMKPRQRDIFYVVGESTS    536
tr|A0A0L1KHT0|A0A0L1KHT0_9EUGL     LRLGLVEDGGNRMRITKLLRYRSSMYPFHYN-KLVSLDDYVSRMKEKQNHIFFVVGEDIA    506
tr|A0A1X0P079|A0A1X0P079_9TRYP     IRLGIILEDANNRGRLAKLLRYASSKS---NG-TLISLQEYTDRMKPDQKNIYYLTGDSVD    494
tr|Q4DW89|Q4DW89_TRYCC             IRLGILEDANNRGRLAKLLRYTSTKS---NG-TLVSLQEYTDRMKPEQKHIYFLTGDSVK    499
tr|A0A422PWD8|A0A422PWD8_9TRYP     IRLGILEDANNRGRLAKLLRYTSSKS---NG-TLVSLQDYTDRMKPGQKSIYFLTGESVS    496
tr|A0A422NRM6|A0A422NRM6_TRYRA     IRLGIIDDANNRGRLAKLLRYTSSTS---NG-TLVSLQEYTDRMKPGQKNIYFLTGDSVS    501
tr|A0A0S4KLS2|A0A0S4KLS2_BODSA     LRLGMIEDGSNRARLTKLLRYKSSKS---DN-KLISLQDYVDRMPESQKDIYYVSAESIE    526
tr|A0A0N0VF86|A0A0N0VF86_LEPPY     LRLGVMLDSNNRNRLTKLFRYKSSKS---DG-AYISLQTYVDRMKKGQKGIYYISGDTIE    520
tr|A4HH83|A4HH83_LEIBR             LRLGAITDSNNRNRLMKLFRYKSSAS---ET-EYISLQAYVDRMKKGQKGIYYLSGDSVD    527
tr|E9ADS8|E9ADS8_LEIMA             LRLGVMLDSNNRNRLTKLFRYKSSRS---ES-EYISLQTYVDRMKKGQKGIYYLSGDSVA    527
sp|Q9NGD0|ENPL_LEIIN               LRLGVMLDSNNRNRLTKLFRYKSSRS---ES-EYISLQTYVDRMKKGQKGIYYLSGDSVA    527
tr|A4I4C9|A4I4C9_LEIIN             LRLGVMLDSNNRNRLTKLFRYKSSRS---ES-EYISLQTYVDRMKKGQKGIYYLSGDSVA    527
sp|Q8T7E0|ENPL_LEIDO               LRLGVMLDSNNRNRLTKLFRYKSSRS---ES-EYISLQTYVDRMKKGQKGIYYLSGDSVA    527
tr|S9U4L8|S9U4L8_9TRYP             LRYGVVSDPNNRGRLLKLLRYRSSMS---RG-KYVTLKEYTDRMKKGQRGIYYLTGDSIE    524
tr|A0A7G2C3S2|A0A7G2C3S2_9TRYP     LRYGLIVDPSNRGRITKLLRYKTSRS---ED-RYVGLQDYIDHMQKGQKAIYYLSGDSIA    548
sp|P14625|ENPL_HUMAN               IKLGVIEDHSNRTRLAKLLRFQSSHH---PT-DITSLDQYVERMKEKQDKIYFMAGSSRK    547
tr|Q7T3L3|Q7T3L3_DANRE             IKLGVIEDHSNRTRLAKLLRFQTSHS---DT-VLSSLEQYVERMKEKQDKIYFMAGTSRK    547
sp|Q9STX5|ENPL_ARATH               VKLGIIEDAANRNRLAKLLRFETTKS---DG-KLTSLDQYIKRMKKSQKDIFYITGSSKE    577
tr|A0A061FKM2|A0A061FKM2_THECC     IKLGIIEDATNRNRLAKLLRFESTKS---DG-KLTSLDQYISRMRAGQKDIFYITGNSKE    580
tr|Q5Z9N8|Q5Z9N8_ORYSJ             VKLGIIEDATNRNRLAKLLRFESTKS---EG-KLASLDEYISRMKPGQKDIFYITGSSKE    578
tr|W2ZJ09|W2ZJ09_PHYPR             IKLGVMDDAANRAKLVKLLRFVTSES---DG-KWTSLEQYVDRMKDWQDSIYYIAAENAD    583
tr|W2ZJ74|W2ZJ74_PHYPR             IKLGVMDDAANRAKLVKLLRFVTSES---DG-KWTSLEQYVDRMKDWQDSIYYIAAENAD    582
tr|F0Y644|F0Y644_AURAN             IKLGIIEDSANRSKLTKLLRYKSNKS---GE-GYVSLEEYVENMKDWQKAIYYIAGESVA    533
tr|D8LHY7|D8LHY7_ECTSI             IKLGVMEDNANKSKLVKLLRFKTNKS---DG-KWVSLEDYVAGMPEWQSSIFYIAGESTE    535
                                   :: * : *  *: :: :*.*: :.            *. *    *    *   *::: .

tr|A0A0L1KBS8|A0A0L1KBS8_9EUGL     KLSKLAVVEDARNHGIEVLLMDDAIDEYVVASLTQYGGKNLVNLSKEGVQLEELTDY-EK    595
tr|A0A0L1KHT0|A0A0L1KHT0_9EUGL     KLSHLPVVEDAMNRGVEVLYMVDAIDEYVTSSVTDYSGYRLVNLALNMAQPEPPTEL-EE    565
tr|A0A1X0P079|A0A1X0P079_9TRYP     KMKQSPHMEEALQRDVEVIFMTDAIDEYVVGQIHDFANKKLINLAKEGAQFDEQTDK-EK    553
tr|Q4DW89|Q4DW89_TRYCC             KMRQSPHIEEALERDVEVLFMTDAIDEYVVSQVQDFGNKRLINLAKDNARLDEPTER-DK    558
tr|A0A422PWD8|A0A422PWD8_9TRYP     KMKQTPHIEEALQRDVEVLFMTDAIDEYVVSQIQDFANKRLVNLAKDSVQFEEQTEK-EK    555
tr|A0A422NRM6|A0A422NRM6_TRYRA     KMKQSPHIEEALQRGVEVLFMTDAIDEYVASQIHDFASKRLINLAKESAQFEELTER-EK    560
```

```
tr|A0A0S4KLS2|A0A0S4KLS2_BODSA      KIKQLPVLEDATNRNLEVLFMTDAIDEYVVGHVTDFAGKKLVNLAKEGVKFEDESKR-EK    585
tr|A0A0N0VF86|A0A0N0VF86_LEPPY      RIQKSPMLEDAVNHDVEVIFMTDAIDEYVVAQVTDFAGKKLINLAKEGVQFEETDAR-QR    579
tr|A4HH83|A4HH83_LEIBR              RIKKSPVLEDALNHGFEVIFMTDPIDEYAVSHLTDFAGNKLINLAKEGVAFEETDSR-QR    586
tr|E9ADS8|E9ADS8_LEIMA              RIKKSPVLEDAVNHDVEVIFMTDAIDEYVVSQLTDFAGKKLINLAKEGVQFEESDAR-QR    586
sp|Q9NGD0|ENPL_LEIIN                RIKKSPVLEDAVNHDVEVIFMTDAIDEYVVSQLTDFAGKKLINLAKEGVQLEESDAR-QR    586
tr|A4I4C9|A4I4C9_LEIIN              RIKKSPVLEDAVNHDVEVIFMTDAIDEYVVSQLTDFAGKKLINLAKEGVQFEESDAR-QR    586
sp|Q8T7E0|ENPL_LEIDO                RIKKSPVLEDAVNHDVEVIFMTDAIDEYVVSQLTDFAGKKLINLAKEGVQFEESDAR-QR    586
tr|S9U4L8|S9U4L8_9TRYP              LIERSPLLQDAKKRGIEVLYMTDTIDEYVVNNVPDFAGKKLINIAAADADYEEDDDR-QR    583
tr|A0A7G2C3S2|A0A7G2C3S2_9TRYP      KIKTSPLLQDALKRNVEVIYMTDAVDEYVVAQVPDFSGKKLVNLAKEGLQFDEENSPEAL    608
sp|P14625|ENPL_HUMAN                EAESSPFVERLLKKGYEVIYLTEPVDEYCIQALPEFDGKRFQNVAKEGVKFDESEKTKES    607
tr|Q7T3L3|Q7T3L3_DANRE              EAESSPFVEKLLKKGYEVVYLTEPVDEYCIQALPEFDGKRFQNVAKEGVKFDESDKAKEK    607
sp|Q9STX5|ENPL_ARATH                QLEKSPFLERLIKKGYEVIFFTDPVDEYLMQYLMDYEDKKFQNVSKEGLKVGKDSKDK--    635
tr|A0A061FKM2|A0A061FKM2_THECC      QLEKSPFLERLKKKNYEVIFFTDPVDEYLMQYLMDYDGKQFQNVSKEGLKIGKDSKNK--    638
tr|Q5Z9N8|Q5Z9N8_ORYSJ              QLEKSPFLERLTKKNYEVIYFTDPVDEYLMQYLMDYEDKKFQNVSKEGLKLGKDSKLK--    636
tr|W2ZJ09|W2ZJ09_PHYPR              ACEKSPFMEKMRAKGLEVLYFVDALDEYMVSHISEFDGKKLVSITKEGIKFGDEDESLTQ    643
tr|W2ZJ74|W2ZJ74_PHYPR              ACEKSPFMEKMRAKGLEVLYFVDALDEYMVSHISEFDGKKLVSITKEGIKFGDEDESLTQ    642
tr|F0Y644|F0Y644_AURAN              AVEESPFLEKCKAKDLEVLYLVDPIDEYAIQHVTEFDGKKLQSVTKEGLKFGDENEDVEK    593
tr|D8LHY7|D8LHY7_ECTSI              AVEKSPFLEKLNKKGLEVLYLTEPIDEMTMGSITDFEDKKMQSVTKEGLSFGDEDVADVK    595
                                             ::       :. **: : : :**    : :: . .: .::

tr|A0A0L1KBS8|A0A0L1KBS8_9EUGL      EIEVLREKKYATFIQWFSKIIEKATHRKINKIVLGKRKTEQPLVLSSPSDGMTPNLARIL    655
tr|A0A0L1KHT0|A0A0L1KHT0_9EUGL      SLKEKREEKYKPLIEWFGKLLEDHKV-HSDKIVLCHRHTSRPAIIASPKIGTTANMARIN    624
tr|A0A1X0P079|A0A1X0P079_9TRYP      AIEKKRIEKYRPLTDRLARMFKRSNV---RKVILTKRQTSEPFILSSQEGELTARMVNIM    610
tr|Q4DW89|Q4DW89_TRYCC              AIEKERKKKYEPLTERLLTLFGKSQV---RKVILTRRQSSEAFILSTQENDMTPRMVNVM    615
tr|A0A422PWD8|A0A422PWD8_9TRYP      AIEKKRIEKYQPLTERLANLFKKNDV---RKVILTKRQSSEAFILSTQENDLTPRMVNVM    612
tr|A0A422NRM6|A0A422NRM6_TRYRA      AIEKKRIERYEPLTSRLATLFKKSGV---RKVILTKRQSSEAFILSTQESDLTPRMVSVM    617
tr|A0A0S4KLS2|A0A0S4KLS2_BODSA      AIDAKRKEKYEPVLKYFKDLLGE-QV---TKVVLTKRKTSEPIILSSRQHDVTARMANII    641
tr|A0A0N0VF86|A0A0N0VF86_LEPPY      VIDKKRKEKYEALFTRLRSLFGYAEV---RKVILTKRLTNEAFILSSGENQITARLANIM    636
tr|A4HH83|A4HH83_LEIBR              VVDKKRREKYNSLFTHLRGIFGYSEV---RKVILTRRMTNEAFIVSSGDNQITARLASIM    643
tr|E9ADS8|E9ADS8_LEIMA              VADRKRKEKYDSFFTHLRVLFGYSEV---RKVILTKRMTNEAFIVSSGENQITARLASIM    643
sp|Q9NGD0|ENPL_LEIIN                VADRKRKEKYDSFFTHLRALFGYSEV---RKVILTKRMTNEAFLVSSGENQITARLASIM    643
tr|A4I4C9|A4I4C9_LEIIN              VVDRKRKEKYDSFFTHLRALFGYSEV---RKVILTKRMTNEAFIVSSSENQITARLASIM    643
sp|Q8T7E0|ENPL_LEIDO                VVDRKRKEKYDSFFTHLRALFGYSEV---RKVILTKRMTNEAFIVSSSENQITARLASIM    643
tr|S9U4L8|S9U4L8_9TRYP              ALRQKREERFEPLFSLLRQLFGEKEV---KKIVLTKRHTDLPFLLTNDENHMSIRMQKLM    640
tr|A0A7G2C3S2|A0A7G2C3S2_9TRYP      NLEKKRNEKYAPLLDRLKRLFGK-QV---RKLVLTKRYTTEPFLLIADENHLTARMANIL    664
sp|P14625|ENPL_HUMAN                --REAVEKEFEPLLNWMKDKALKDKI---EKAVVSQRLTESPCALVASQYGWSGNMERIM    662
tr|Q7T3L3|Q7T3L3_DANRE              --REALEKEFEPLTTWMKDKALKEQI---EKAVLSQRLTNSPCALVASQYGWSGNMERIM    662
sp|Q9STX5|ENPL_ARATH                ----ELKEAFKELTKWWKGNLASENV---DDVKISNRLADTPCVVVTSKFGWSANMERIM    688
tr|A0A061FKM2|A0A061FKM2_THECC      ----ELKESFKELTKWWKGALTSENV---DDVKITNRLDNTPCVVVTSKFGWSANMERIM    691
tr|Q5Z9N8|Q5Z9N8_ORYSJ              ----DLKESFKELTDWWKKALDTESV---DSVKISNRLSDTPCVVVTSKYGWSANMEKIM    689
tr|W2ZJ09|W2ZJ09_PHYPR              KREQLYADKYVALTTALKT-LYGDKI---SRVTMSQRVVDSPAVMVTSQWGYSANMQRIM    699
tr|W2ZJ74|W2ZJ74_PHYPR              KREQLYADKYVALTTALKT-LYGDKI---SRVTMSQRVVDSPAVMVTSQWGYSANMQRIM    698
tr|F0Y644|F0Y644_AURAN              KRMQLYKETFKPLTDYLKA-LYGDAV---VKVSVSRRVETTPTVIVTSQYGNSANMERIM    649
tr|D8LHY7|D8LHY7_ECTSI              KREKYYKKMFTPLAEHLKD-MFKGKI---SKVSVSQRVEGTPAIIVSAAYGYSANMERIM    651
                                             . :  .                     : .*     :       : .:  :

tr|A0A0L1KBS8|A0A0L1KBS8_9EUGL      KGQTFGDTRM--YVKD--AEKIVEVNYLHPAVEEIFNRFQEDENDEVSMNAALLLYETAC    711
tr|A0A0L1KHT0|A0A0L1KHT0_9EUGL      KGQTLGDH----GHKS--ADKIVEINHLSPIMDEIYRRVIVDENDEVAKNAAILVFETAC    678
tr|A0A1X0P079|A0A1X0P079_9TRYP      NQQAISSM----QQMH--HTRVLEINYRHPLVQDLLKRFEADPKDQVAVDVAWVLFGTAN    664
tr|Q4DW89|Q4DW89_TRYCC              NQQAISSL----HTMR--YSRVLELNHRHPLVRDLLTRFEADANDQTAIDVAWVLFGTAN    669
tr|A0A422PWD8|A0A422PWD8_9TRYP      NQQAISST----QTMR--YSRVLEINHRHPLVLDLLTRFKADPKDQTAIDVAWVLFGTAN    666
tr|A0A422NRM6|A0A422NRM6_TRYRA      NQQAISSS----QMMR--YSRVLEINHRHPLVRDLLTRFESDPKDQTAIDVAWVLFGTAN    671
tr|A0A0S4KLS2|A0A0S4KLS2_BODSA      RGQALGDAKQ-NEAQT--AKRVMEINHLHPLIEEIFKRVKADDKDKVAEDVALVLFDTAN    698
tr|A0A0N0VF86|A0A0N0VF86_LEPPY      RGQSMALM---DQQTT--AERVLEVNYRHPLVDEMFKRFAVDEDDAVAMDVAWVLYDTAN    691
tr|A4HH83|A4HH83_LEIBR              RGQSMSLA---DQKVT--AERVLEVNYRHPLVDEMFKRFTVNEDDEVATDIAWVLYDTAN    698
tr|E9ADS8|E9ADS8_LEIMA              RGQSMSLA---NQQMT--AERVLEVNYRHPLVDEMFKRFTVDENDEVATDIAWVLYDTAN    698
sp|Q9NGD0|ENPL_LEIIN                RGQSMSLA---NQQMT--AERVLEVNYRHPLVDEMFKRFTVDEDDEVATDIAWVLYDTAN    698
tr|A4I4C9|A4I4C9_LEIIN              RGQSMSLA---NQQMT--AERVLEVNYRHPLVDEMFKRFTVDEDDEVATDIAWVLYDTAN    698
sp|Q8T7E0|ENPL_LEIDO                RGQSMSLA---NQQLT--AERVLEVNYRHPLVDEMFKRFTVDEDDEVATDIAWVLYDTAN    698
tr|S9U4L8|S9U4L8_9TRYP              RSQMTGST---SSMLR--AQRVLEVNYRHPLVDELYRRFIVDPNDNIAEDIAWVLFDTAN    695
tr|A0A7G2C3S2|A0A7G2C3S2_9TRYP      KSQSLGRGATEDVPLT--AERVVEVNWRHPLVDEVYKRFLVDENDLVAEDIAWTLFDTAN    722
sp|P14625|ENPL_HUMAN                KAQAYQTGKDISTNYYASQKKTFEINPRHPLIRDMLRRIKEDEDDKTVLDLAVVLFETAT    722
tr|Q7T3L3|Q7T3L3_DANRE              KAQAYQTGKDISTNYYASQKKTLEINPRHPLIKEMLRRVNEDAEDKTAADLAVVLFETAT    722
sp|Q9STX5|ENPL_ARATH                QSQTLSDANKQAY-MR--GKRVLEINPRHPIIKELKDRIASDPEDESVKETAQLMYQTAL    745
tr|A0A061FKM2|A0A061FKM2_THECC      QSQTLSDASKQAY-MR--GKRVLEINPRHPIIKELRERVVKDPEDEGVKQTAQLIYQTAL    748
tr|Q5Z9N8|Q5Z9N8_ORYSJ              QSQTLSDASKQAY-MR--GKRVLEINPRHPIIKELRDKVAQDSESESLKQTAKLVYQTAL    746
tr|W2ZJ09|W2ZJ09_PHYPR              KAQTFGNGDKNSP-MYGTGSAILELNPRHPIVSKLNELMVSDPEKEETKDLAWLLYDTAL    758
tr|W2ZJ74|W2ZJ74_PHYPR              KAQTFGNGDKNSP-MYGTGSAILELNPRHPIVSKLNELMVSDPEKEETKDLAWLLYDTAL    757
tr|F0Y644|F0Y644_AURAN              RAQAFSEKQAMGQ-MS--SQKTMEINPRHPIIAKLKELVAAEKDDEYTSDLSWLLLDNAL    706
tr|D8LHY7|D8LHY7_ECTSI              KAQTLADSKQMGL-LG--GHRSMEINPRHPIVHELNKKIEEDPDSEETKDLSWLLYETAL    708
                                    . *                        .*:*   * : .:   .  : ..   : :  :  .*

tr|A0A0L1KBS8|A0A0L1KBS8_9EUGL      HQSAFDIDEVHNYAARMYAMIAQGLDVENHNTRLTEQASDYEAPQEDL------------    759
tr|A0A0L1KHT0|A0A0L1KHT0_9EUGL      LESGFEVEDLQSFSDRVLKVIYQGLDMGLDQAMLEEDLSQYQVESNSNADQHVEL-----    733
tr|A0A1X0P079|A0A1X0P079_9TRYP      LQAEFPITDQAMFAKRMNRLLRGRVGISTDDTLLPPDDDEYDISDVKPDTAATDENVLLP    724
tr|Q4DW89|Q4DW89_TRYCC              LQAGFPVSNQAMYAKRVNRLLRGRVGLAADDTILPPDDDEYDISDVKPDTTGTDEGLLLP    729
tr|A0A422PWD8|A0A422PWD8_9TRYP      LQGGFPVSNQAMYAKRVSRLLRGRVGLAADDTMLPPDDNEYDVSDVEPDTAGTDEGLLLP    726
tr|A0A422NRM6|A0A422NRM6_TRYRA      LQGDFPVSNQAMYAKRVNRLLRGRVGLAADDTMLPPDDNEYDVSDIKPSTSGTDEGLLLP    731
tr|A0A0S4KLS2|A0A0S4KLS2_BODSA      LQNGFDIEDTLAFSRRMSRLLRQSVDIPADAAMLTEDVSEYEIEDNEDAEDDE-------    751
tr|A0A0N0VF86|A0A0N0VF86_LEPPY      LQAEFPVADVAAYSRRINRLLRSSVDLAADETLLPPDDAEYAVSTTEAAEEEE-------    744
tr|A4HH83|A4HH83_LEIBR              LQAEFPVADVAAYAKRINRLLRSSVDLNADDSLLPPDDDEYTVFDTETEEEKP-------    751
tr|E9ADS8|E9ADS8_LEIMA              LQAEFPVADVAAYSKRINRLLRSSVDLSADDSLLPPDDAEYTVSDTEAEEEEE-------    751
sp|Q9NGD0|ENPL_LEIIN                LQAEFPVADVAAYSKRINRLLRSSVDLSADDSLLPPDDAEYTVSDTEAEEEEE-------    751
tr|A4I4C9|A4I4C9_LEIIN              LQAEFPVADVAAYSKRINRLLRSSVDLSADDSLLPPDDAEYTVSDTETEEEEE-------    751
sp|Q8T7E0|ENPL_LEIDO                LQAEFPVADVAAYSKRINRLLRSSVDLSADDSLLPPDDAEYTVSDTETEEEEE-------    751
tr|S9U4L8|S9U4L8_9TRYP              MQAEFPVLDVAAYSQRVVRLLATSVDLDPDQPLMAEDNGEAEEEETEDL-----------    744
tr|A0A7G2C3S2|A0A7G2C3S2_9TRYP      LQGDFPVVDTQAYAQRVTRLLSSSVDLDPTMGLLPPDVDDDGLGDL--------------    767
sp|P14625|ENPL_HUMAN                LRSGYLLPDTKAYGDRIERMLRLSLNIDPDAKVEEEPEEEP----EETAEDTTEDTE-QD    777
```

```
tr|Q7T3L3|Q7T3L3_DANRE              LRSGYQLQDTKAYGERIERMLRLSMNVDLDAQVEEEPEEEP----EEQTEEAEDEEE-VQ    777
sp|Q9STX5|ENPL_ARATH                IESGFILTDPKDFAARIYNSVKSGLNISPDAVADEEIEAAE----EPETSEATETK----    797
tr|A0A061FKM2|A0A061FKM2_THECC      MESGFSLPDPKDFASRIYSSVKSSLNISPDATIEDDDDVEE----T-ETE--TETE----    797
tr|Q5Z9N8|Q5Z9N8_ORYSJ              MESGFNLPDPKDFASSIYRSVQKSLDLSPDAAVEEEEEVEE----A-EVE--EKES----    795
tr|W2ZJ09|W2ZJ09_PHYPR              INSGFDMTDTTQFSTRVHRIMKSSMGIDSLELEPEI-EVPE----EEQEVDEEEEA--ED    811
tr|W2ZJ74|W2ZJ74_PHYPR              INSGFDMTDTTQFSTRVHRIMKSSMGIDSLELEPEI-EVPE----EEQEVDEEEEA--ED    810
tr|F0Y644|F0Y644_AURAN              MQSGFEASDVAAFSERALRILKSGLSVESMDLLPEI-EVPD----EPEEEEEEEDA--VD    759
tr|D8LHY7|D8LHY7_ECTSI              TASGFQVEDTEAFANRVQRAMAKTLNLSSMDLLEEM-EIPD----EEEEEEGEGDEQFQD    763
                                             :   :   :.        :   :.:

tr|A0A0L1KBS8|A0A0L1KBS8_9EUGL      ------------------------------------------------------------    759
tr|A0A0L1KHT0|A0A0L1KHT0_9EUGL      ------------------------------------------------------------    733
tr|A0A1X0P079|A0A1X0P079_9TRYP      IDKETDEIPEAEKEGSTEGTKTAA-------EG--------------STEEKEKEKEKSK    763
tr|Q4DW89|Q4DW89_TRYCC              VDNDGDESSDKEDAEPTAAEM--------------------------------------K    751
tr|A0A422PWD8|A0A422PWD8_9TRYP      VDKEEGEATAAAAEEPTAAEAAEEPTAAAAAEEPTAAAAAEEPTAAAAAEEPKAAAAESK    786
tr|A0A422NRM6|A0A422NRM6_TRYRA      VDKDGEETSSEAAAAE-------Q-----------------------------PAAAESK    755
tr|A0A0S4KLS2|A0A0S4KLS2_BODSA      -------APKADADDDKEEL----------------------------------------    764
tr|A0A0N0VF86|A0A0N0VF86_LEPPY      -------GEAAAENANAAET----------------------------------------    757
tr|A4HH83|A4HH83_LEIBR              -------TNDAEYTVSDTATEEEKPTNDA----------------------------DAS    776
tr|E9ADS8|E9ADS8_LEIMA              -------QP----------------KVDA----------------------------NAD    760
sp|Q9NGD0|ENPL_LEIIN                -------QP----------------KVDA----------------------------NAD    760
tr|A4I4C9|A4I4C9_LEIIN              -------QP----------------KVDT----------------------------NAH    760
sp|Q8T7E0|ENPL_LEIDO                -------QP----------------KVDT----------------------------NAH    760
tr|S9U4L8|S9U4L8_9TRYP              ------------------------------------------------------------    744
tr|A0A7G2C3S2|A0A7G2C3S2_9TRYP      ------------------------------------------------------------    767
sp|P14625|ENPL_HUMAN                EDEEMDVGT-------DEEEETAKEST---------------------------------    797
tr|Q7T3L3|Q7T3L3_DANRE              ADEA----E-------EESEA---------------------------------------    787
sp|Q9STX5|ENPL_ARATH                ---SDDLAG--------GLNIEAEPVEQQE------------------------------    816
tr|A0A061FKM2|A0A061FKM2_THECC      ---TKEGAG--------SSKGEAEPANNDAD-----------------------------    817
tr|Q5Z9N8|Q5Z9N8_ORYSJ              ---S-------------NIKEEAEPSSYDKD-----------------------------    810
tr|W2ZJ09|W2ZJ09_PHYPR              LDESEAQAE--------ATEEADEPVVDGKD-----------------------------    834
tr|W2ZJ74|W2ZJ74_PHYPR              LDESEAQAE--------ATEEADEPVVDGKD-----------------------------    833
tr|F0Y644|F0Y644_AURAN              LDDFEDAGE--------GE-----------------------------------------    770
tr|D8LHY7|D8LHY7_ECTSI              LDEL--------------------------------------------------------    767

tr|A0A0L1KBS8|A0A0L1KBS8_9EUGL      -----------    759
tr|A0A0L1KHT0|A0A0L1KHT0_9EUGL      -----------    733
tr|A0A1X0P079|A0A1X0P079_9TRYP      EDAAAQDAGDL    774
tr|Q4DW89|Q4DW89_TRYCC              PTKTEDDAGDL    762
tr|A0A422PWD8|A0A422PWD8_9TRYP      PAAKDDGADDL    797
tr|A0A422NRM6|A0A422NRM6_TRYRA      PATKDDDVGDL    766
tr|A0A0S4KLS2|A0A0S4KLS2_BODSA      -----------    764
tr|A0A0N0VF86|A0A0N0VF86_LEPPY      ---DDDDEADL    765
tr|A4HH83|A4HH83_LEIBR              ETTGADSDGDL    787
tr|E9ADS8|E9ADS8_LEIMA              EKAEAVDEGDL    771
sp|Q9NGD0|ENPL_LEIIN                EEAEAVGEDDL    771
tr|A4I4C9|A4I4C9_LEIIN              EEAETDGEGDL    771
sp|Q8T7E0|ENPL_LEIDO                EEAETDGEGDL    771
tr|S9U4L8|S9U4L8_9TRYP              -----------    744
tr|A0A7G2C3S2|A0A7G2C3S2_9TRYP      -----------    767
sp|P14625|ENPL_HUMAN                -----AEKDEL    803
tr|Q7T3L3|Q7T3L3_DANRE              -----TSKDEL    793
sp|Q9STX5|ENPL_ARATH                ----ENTKDEL    823
tr|A0A061FKM2|A0A061FKM2_THECC      ----TDLKDEL    824
tr|Q5Z9N8|Q5Z9N8_ORYSJ              ----EL-----    812
tr|W2ZJ09|W2ZJ09_PHYPR              ----E-----L    836
tr|W2ZJ74|W2ZJ74_PHYPR              ----E-----L    835
tr|F0Y644|F0Y644_AURAN              ----------L    771
tr|D8LHY7|D8LHY7_ECTSI              -----------    767
```

### 14. Protein disulphide isomerases (Thioredoxins)

Protein disulfide isomerases (often consisting of multiple thioredoxin domains, arranged in tandem) regulate the oxidative crosslinking of cysteines and their correct pairing, that is absolutely necessary for luminal and extracellular protein domain functionality. Intracellular forms of thioredoxins also exist (without signal peptides) alongside with secreted ones. Although not all secreted thioredoxins have ER retention signals, those that do are often very highly conserved across many eukaryotic lineages, including kinetoplastids.

```
tr|W6KQG4|W6KQG4_9TRYP              ------------------MRSVLLF--FAVLIAL-------------------------T    15
tr|A0A0S4KL52|A0A0S4KL52_BODSA      ------------------MKAF-PLIALLAVAII-------------------------A    16
tr|A0A1X0NMX9|A0A1X0NMX9_9TRYP      -------------------MRLLLLIAF-LLSLL-------------------------R    15
```

```
tr|Q4E3F7|Q4E3F7_TRYCC              ------------------MIHFIFFVAL-FFCSL-------------------------R    16
tr|A0A3R7L858|A0A3R7L858_TRYRA      ------------------MMRFLLLATL-LLCSL-------------------------R    16
tr|A0A422Q717|A0A422Q717_9TRYP      ------------------MIRFLVFVTL-LLCSL-------------------------R    16
tr|A0A7G2CFK3|A0A7G2CFK3_9TRYP      ------------------MAKL-TAALLASLLLA-------------------------A    16
tr|S9V6G7|S9V6G7_9TRYP              ------------------MSKFFTLCILAVLLLS-------------------------T    17
tr|S9VLI6|S9VLI6_9TRYP              ------------------MSKFFTLCILAVLLLS-------------------------T    17
tr|A4HQL6|A4HQL6_LEIBR              ------------------MKLLYVVFFVCT-LLL-------------------------C    16
tr|A7YBW7|A7YBW7_LEIAM              ------------------MQRSFLAFVLCA-LLF-------------------------C    16
tr|Q4Q059|Q4Q059_LEIMA              ------------------MQRSFLVFVLCA-LLF-------------------------C    16
tr|Q8I8E1|Q8I8E1_LEIMA              ------------------MQRSFLVFVLCA-LLF-------------------------C    16
tr|A4ICD5|A4ICD5_LEIIN              ------------------MQRSFLAFVVCA-ILF-------------------------C    16
tr|B3VA16|B3VA16_LEIDO              ------------------MQRSFLAFVVCA-ILF-------------------------C    16
tr|A0A0M9G4E8|A0A0M9G4E8_LEPPY      ------------------MKQSFLLLAVCA-LFL-------------------------C    16
tr|A0A0N0P805|A0A0N0P805_LEPSE      ------------------MKQPLFLLALCA-LFL-------------------------C    16
tr|A0A7E6F1P9|A0A7E6F1P9_OCTVU      ---------------MALFSS-RLNYLLLLLALISQVIDFRCYGEEADEADEATDDDNEP    44
tr|E3NBU5|E3NBU5_CAERE              -------------------------MSWIQAALVASFLAF-------------------A    16
tr|A0A4Z2C989|A0A4Z2C989_9TELE      ---------------------------MLKLMLLAALTGF-------------------T    14
tr|W5KW40|W5KW40_ASTMX              ------------------MAN-SVSGLLSGALFVCLWGGS-------------------V    22
tr|A0A0G4EK32|A0A0G4EK32_VITBC      ---------------MKLVIS-VLAICLAAPAVV------------------------RA    20
tr|A0A058Z7G2|A0A058Z7G2_FONAL      ---------MAPA----TRSAFGLA-LMALLALT------------------------AI    22
tr|G4MPX2|G4MPX2_MAGO7              ---------------MHTVRSFALGLLATAAVVS------------------------AS    21
tr|A0A8J2IGP4|A0A8J2IGP4_FUSEQ      ---------MQHKKIACSFMA--------ALAAY------------------------AS    19
tr|A0A8I3AHZ3|A0A8I3AHZ3_9PEZI      MGRSTPPPPQASGTRACLLASYRASSPSPSLIVH------------------------HI    36


tr|W6KQG4|W6KQG4_9TRYP              RASADVVELTSKNFDEVIAKD-GITFIKFFTTWCGYCKKAAPEFANASDILKD---VATL    71
tr|A0A0S4KL52|A0A0S4KL52_BODSA      VVADDAIDAKDADFDEVVSRN-ALTLVKFYAPWCGHCKRIAPEWDKAATALVG---KAGL    72
tr|A0A1X0NMX9|A0A1X0NMX9_9TRYP      ADASDVVAATVDDFDSVIKKG-EIALVKFYAPWCGHCQKLAPEWEKAAKEVPG---EAIM    71
tr|Q4E3F7|Q4E3F7_TRYCC              AEGSEVVEATDKDFDDVISSG-EIALVKFYAPWCGHCQKLAPEWEKAAKEIPS---GAVM    72
tr|A0A3R7L858|A0A3R7L858_TRYRA      VEGSEVVAATDKNFDSVIARS-KVALVKFYAPWCGHCQKLAPEWEKAAKEIPS---EAIM    72
tr|A0A422Q717|A0A422Q717_9TRYP      VEGSEVVAATDKDFDSVLARG-EVALVKFYAPWCGHCQKLAPEWEKAAKEIPS---EAIM    72
tr|A0A7G2CFK3|A0A7G2CFK3_9TRYP      GCQGKVASLNEADFETGIKKT-EYTLVKFFAPWCGHCKKLAPEFEKASDALGD---KATL    72
tr|S9V6G7|S9V6G7_9TRYP              FACAEVITGTEKNLDDIIAKD-KVSIVKFYAPWCGHCKSLAPEFEKAADALKE---QASL    73
tr|S9VLI6|S9VLI6_9TRYP              FACAEVITGTEKNLDDIIAKD-KVSIVKFYAPWCGHCKSLAPEFEKAADALKE---QASL    73
tr|A4HQL6|A4HQL6_LEIBR              LTSAEVQVATQDNFDNVVSG--DLTLVKFYAPWCGHCKTLAPEFVKAAEMLTG---IATL    71
tr|A7YBW7|A7YBW7_LEIAM              VASAEVQVATKDNFDKIVSG--DLTLVKFYAPWCGHCKTLAPEFIKAAEMLAG---VATL    71
tr|Q4Q059|Q4Q059_LEIMA              VASAEVQVATKDNFDKVVSG--DLTLVKFYAPWCGHCKTLAPEFVKAADMLAG---IATL    71
tr|Q8I8E1|Q8I8E1_LEIMA              VASAEVQVATKDNFDKVVIG--DLTLVKFYAPWCGHCKTLAPEFVKAADMLAG---IATL    71
tr|A4ICD5|A4ICD5_LEIIN              VASAEVQVATKDNFDKIVSG--DLTLVKFYAPWCGHCKTLAPEFVKAADMLAG---IATL    71
tr|B3VA16|B3VA16_LEIDO              VASAEVQVATKDNFDKIVSG--DLTLVKFYAPWCGHCKTLAPEFIKAADMLAG---IATL    71
tr|A0A0M9G4E8|A0A0M9G4E8_LEPPY      VASAEVQVATKSNFDKIIGG--DLTLVKFYAPWCGHCKTLAPEFEKASVTLKG---VATL    71
tr|A0A0N0P805|A0A0N0P805_LEPSE      VASAEVQVATKSNFDEAISG--DLTLVKFYAPWCGHCKTLAPEFEKAAEALKG---VATL    71
tr|A0A7E6F1P9|A0A7E6F1P9_OCTVU      IEEDDVLVLNNNNFDDVIYSR-DTVLVEFYAPWCGHCKRLAPEYEKAAQKLAVNDPPIPL    103
tr|E3NBU5|E3NBU5_CAERE              SAGGAVLEYTDGNFDDLIQTH-DIALVKFYAPWCGHCKKIAPEYEKAAPKLASNDPPVAL    75
tr|A0A4Z2C989|A0A4Z2C989_9TELE      QAASDVLEFTDNDFESKIGDH-EIILVEFFAPWCGHCKRLAPEYEKAATALKG---VVPL    70
tr|W5KW40|W5KW40_ASTMX              SARSDVVELTDVDFDYVAAEH-ETMLVKFYAPWCGHCKKLAPEFETAAKRLKG---VVTL    78
tr|A0A0G4EK32|A0A0G4EK32_VITBC      EDEEAVVTLGTDNFTSFIEQN-PLSLVEFYAPWCGHCKRLAPEYEKAAKKLKESG-GPPL    78
tr|A0A058Z7G2|A0A058Z7G2_FONAL      AAASDVVDLNKDNFDSVVNAEGATSMVEFFAPWCGHCKQLAPKYEEAATKLKGI---ATI    79
tr|G4MPX2|G4MPX2_MAGO7              DAPSDVVQLKEDTFDAFIKEN-DLVLAEFFAPWCGHCKALAPHYEKAATSLKEKN--IKL    78
tr|A0A8J2IGP4|A0A8J2IGP4_FUSEQ      AADSDVKQLTKDTFDEFVKSN-DLVLAEFFAPWCGHCKALAPEYEEAATTLKEKN--IRL    76
tr|A0A8I3AHZ3|A0A8I3AHZ3_9PEZI      KGSKCTASLKKDTFDDFVKTN-DLVLAEFFAPWCGHCKALAPEYEEAATSLKEKN--IKL    93
                                             .        :        : :*:: ***:*:  **.:  *:  :         :

tr|W6KQG4|W6KQG4_9TRYP              AEVDCDKERNICKRFEVNGYPKFLLFRKG-VNISTFSGQRVASEMVDYVKVQMASDTNSV    130
tr|A0A0S4KL52|A0A0S4KL52_BODSA      VRVDCTTETAVAQKFGIQGYPTIKVFREG-QLVGDYDAGRTSEAIVQYVISNSGPAVTVV    131
tr|A0A1X0NMX9|A0A1X0NMX9_9TRYP      VDVDCTKETKLAEKYSIKGFPTIILFRNG-EEVETYKGARQSDAIVKYIKANVGPAAVYP    130
tr|Q4E3F7|Q4E3F7_TRYCC              VDVDCTKESNLAQKYSIKGFPTIILFRDG-KEVEHYKGGRKSSDIVNYVKANLGTAVVHV    131
tr|A0A3R7L858|A0A3R7L858_TRYRA      VDVDCTKEKNLAKKYSIQGFPTIILFRDG-EEVEHYKGARKSTDLINYVKANVGPAVVYP    131
tr|A0A422Q717|A0A422Q717_9TRYP      VDVDCTKEKELAKKYSIQGFPTIILFREG-KEVEHYKGGRQSTDLVNYVKANVGPAVVYP    131
tr|A0A7G2CFK3|A0A7G2CFK3_9TRYP      AEVDCTQNEKLCGKFDVKGYPTLILFKNG-EKHLDFDGGRTAPEIISYVKSQYEPAVSVS    131
tr|S9V6G7|S9V6G7_9TRYP              VKVDCTVEKKLAEKYDIKGFPTLIVFRKG-EKVENYEGGRTAEEIISYVRSLSSPLFTPV    132
tr|S9VLI6|S9VLI6_9TRYP              VKVDCTVEKKLAEKYDIKGFPTLIVFRKG-EKVENYEGGRTAEEIISYVRSLSSPLFTPV    132
tr|A4HQL6|A4HQL6_LEIBR              AEVDCTTEKALAEKYEVKGFPTLYVFRNG-VKVKAYDGPRTADGIASYMKSHVGPSMKVV    130
tr|A7YBW7|A7YBW7_LEIAM              AEVDCTKEEALAEKYEVKGFPTLYLFRNG-EKVKDYDGPRTAAGIASYMKAQVGPSMKTI    130
tr|Q4Q059|Q4Q059_LEIMA              AEVDCTKEESLAEKYEIKGFPTLYIFRNG-EKVKIYDGPRTAAGIASYMKAHVGPSMKAI    130
tr|Q8I8E1|Q8I8E1_LEIMA              AEVDCTKEESLAEKYEIKGFPTLYIFRNG-EKVKIYDGPRTAAGIASYMKAHVGPSMKAI    130
tr|A4ICD5|A4ICD5_LEIIN              AEVDCTKEESLAEKYQIKGFPTLYIFRNG-EKVKDYDGPRTAAGIASYMKAHVGPSIKAI    130
tr|B3VA16|B3VA16_LEIDO              AEADCTKEESLAEKYQIKGFPTLYIFRNG-EKVKDYDGPRTAAGIASYMKAHVGPSIKAI    130
tr|A0A0M9G4E8|A0A0M9G4E8_LEPPY      AEVDCTKETELASKFDIKGFPTMMIFRNG-EKVEDYSGPRTAAGITAYIKAQVGPAVTAI    130
tr|A0A0N0P805|A0A0N0P805_LEPSE      VKVDCTKETELASKFGIKGFPTLIIFRNG-EKLEDYNGPRTAAGITAHMKAHVGPAVTTI    130
tr|A0A7E6F1P9|A0A7E6F1P9_OCTVU      AKIDATIATTSAMKYEVSGYPTLKIFRAG-KQ-FEYKGPREDYGIVNYMISQQGGASKLQ    161
tr|E3NBU5|E3NBU5_CAERE              VKVDCTTEKTVCDKFGVKGFPTLKIFRNG-VPAQDYDGPRDADGIVKFMRGQSGPSSKEL    134
tr|A0A4Z2C989|A0A4Z2C989_9TELE      AKVDCTSNSNICSKYQVSGYPTLKVFRDG-EESGAYDGPRTSDGIVSYFKKQVGPASVAL    129
tr|W5KW40|W5KW40_ASTMX              AKVDCTSSTEVCGRFGVTGYPTLKIFRNG-RESSSYDGPRSAVGIVDQMKKQAGPDSVPL    137
tr|A0A0G4EK32|A0A0G4EK32_VITBC      AKVDATQEKELAEKYEVRGYPTIKFFRNG--QAEEYDGGRTEPTIIEWIEKRTGPAVRVV    136
tr|A0A058Z7G2|A0A058Z7G2_FONAL      AKVDCTEQTELCQKHGIRGYPTLKVFRNKGDSVVDYSGAREVDALVSFMKKQVLPAVSTF    139
tr|G4MPX2|G4MPX2_MAGO7              IKVDCTEEQDLCQKHGVEGYPTLKVFRGL-DNVVPYKGQRQDDGIISYMVKQSLPAVSTI    137
tr|A0A8J2IGP4|A0A8J2IGP4_FUSEQ      AKIDCTEESDLCKDHGVEGYPTLKVFRGL-ENVTPYSGQRKAAGITSYMIKQSLPAVSLL    135
tr|A0A8I3AHZ3|A0A8I3AHZ3_9PEZI      AKIDCTEEADLCQTYGVEGYPTLKVFRGP-DNISPYSGQRKAAAITSYMIKQSLPAVSLL    152
                                    *.       .  . : *:*.: .*:           :.. *    :   .

tr|W6KQG4|W6KQG4_9TRYP              SSEEEFEKVKSENPRVCLIKTKSDETELAKLVNKLIPKYKATYRFVVINDEKVSPDDS--    188
tr|A0A0S4KL52|A0A0S4KL52_BODSA      SSADELEVLKKESPIVVVVFSAEEGTTSAVTHAEAAKTLRSKYRFVRVADATLFD-GA--    188
tr|A0A1X0NMX9|A0A1X0NMX9_9TRYP      TTAEEVEKLKEEHEILCVGLTSDAESPLAKELLSVAQRLRMKMKFAIVTKPEIVPGEK--    188
tr|Q4E3F7|Q4E3F7_TRYCC              ETAEELEKLREEHNAVCVGVTSDMESTLSKTLATSAEGLRMKMKFVVITDSNILPDEK--    189
tr|A0A3R7L858|A0A3R7L858_TRYRA      SSAEELAKLKEEHDAVCVGVTSDTESALSKALETSAKNLRMKLKFAVITDSKILPEEK--    189
tr|A0A422Q717|A0A422Q717_9TRYP      SSAEELAKLKEEHDAVCVGVASGTESALSTTLANSAQRLRMKLKFAVITDPEILPEEK--    189
tr|A0A7G2CFK3|A0A7G2CFK3_9TRYP      KTQADVDKLVKEGLPVVVIKTASETSQTFKTLEPIAKTRRLSNAFTLVSDAAVVKASP--    189
```

```
tr|S9V6G7|S9V6G7_9TRYP           KSQKELDELIALGLGVALVKTASEDGDVVTRLTPIAKDLSKVVAFAVGTDASIAADAP--    190
tr|S9VLI6|S9VLI6_9TRYP           KSQKELDELIALGLGVALVKTASEDGDVVTRLTPIAKDLSKVVAFAVGTDASIAADAP--    190
tr|A4HQL6|A4HQL6_LEIBR           AKAEELEDLMKENFPLCVVKTANADSEMASMMTKLADSLRSEMNFALVTDTAISPADA--    188
tr|A7YBW7|A7YBW7_LEIAM           AKAGELEDLKKEAFPVCVVKTASADSEMALMMTKVANFLRSQMNFVLVTDATISPDDG--    188
tr|Q4Q059|Q4Q059_LEIMA           STAEELEELKKETFPVCVVKTASTDSEMASMITKVADSLRSQMNFVLVTDAAISPNDA--    188
tr|Q8I8E1|Q8I8E1_LEIMA           STAEELEELKKETFPVCVVKTASTDSEMASMITKVADSLRSQMNFVLVTDAAISPNDA--    188
tr|A4ICD5|A4ICD5_LEIIN           SKAEELEELKKETFPLCVVKTASADSEMASMITKVADSLRSQMNFVLVTDAAISPNDA--    188
tr|B3VA16|B3VA16_LEIDO           SKAEELEELKKETFPVCVVKTASADSEMASMITKVADSLRTQMNFVFVTDAAISPNDA--    188
tr|A0A0M9G4E8|A0A0M9G4E8_LEPPY   EKAEQLDELKKEDLPVCVVKTASADSALATTMTKVANVFRTQVNFALVTDEAIAADAA--    188
tr|A0A0N0P805|A0A0N0P805_LEPSE   EKAEQLEEIKKGDLPVCIIKTADAKSALVETMTKVANSLRTQMNFVLITDAAISADDT--    188
tr|A0A7E6F1P9|A0A7E6F1P9_OCTVU   ASAKELQKYMKEDDITIVGFFNSLDAPLMSSYLDAANEVRDDYKFAHVLDKDVTKSYDLV    221
tr|E3NBU5|E3NBU5_CAERE           KTVAEFEKFTGGDENVVVGFFESE-SKLKDSFLKVADTERDRFAFAHTSNKDIIKKAG-Y    192
tr|A0A4Z2C989|A0A4Z2C989_9TELE   AGEEELQKFISEKDSSVVGFFADDKSTAQAEFLKAASALRDKYRFAHTNAEVLLKSQNVG    189
tr|W5KW40|W5KW40_ASTMX           HNEKDLETFVNHYDASVVGVFESSDSPQLVEFLKGANLMRESFRFAHTTDLQLGLKYNLT    197
tr|A0A0G4EK32|A0A0G4EK32_VITBC   TG-EELEKLKKDKDILMVAFVKDQEAELFKTYEKVADKNRDLALFIAVVSGDA-------    188
tr|A0A058Z7G2|A0A058Z7G2_FONAL   EKADELTKFIEDDRVVVVGFFEEGDA-KAANFKNVAEKLREKMSFASTTSADLAKAHGVT    198
tr|G4MPX2|G4MPX2_MAGO7           TT-DSLEEFKKTDKVVVVAYLSSDDKTSTETFTQAAEKLRDHYPFGSSTDAALAEAEGVK    196
tr|A0A8J2IGP4|A0A8J2IGP4_FUSEQ   TK-DTLEEFKTADKVVVVAYLNADDKSSNETFSKLAEGLRDTYLFGGVNDAAVAEAEGVK    194
tr|A0A8I3AHZ3|A0A8I3AHZ3_9PEZI   TK-DTLEEFKTADKVVLVAYIAADDKASNETFNKVAESLRDNYLFGGVNDAAVAEAEGVK    211
                                          .        :                          *

tr|W6KQG4|W6KQG4_9TRYP           MESITVYR------KGDDKE--VFDGNLS---SEALSTFLDKAPVMFIGTLSPESIPFYS    237
tr|A0A0S4KL52|A0A0S4KL52_BODSA   SDSIVIYK------QFDEGRVPYTGDVSK---VADLTAFLSTTSVRVFDEIGPENYKMYM    239
tr|A0A1X0NMX9|A0A1X0NMX9_9TRYP   AETVVVYR------KDDEKE--VYQGNME---ADNLKNFLETAVVPFAGEIGPANYLRYA    237
tr|Q4E3F7|Q4E3F7_TRYCC           PESIIVFR------KGGEKE--VFDGAME---TADLKSFLEVAFIPFMGEINPNTYLDYA    238
tr|A0A3R7L858|A0A3R7L858_TRYRA   LESIVVFR------KGGEKE--IYEGAME---VKELNSFLAVSFLPFAGEINPKTYMNYA    238
tr|A0A422Q717|A0A422Q717_9TRYP   PESIIVFR------KGGEKE--VYEGAME---AEELNSFLEVSVLPFAGEIGPRTYMNYA    238
tr|A0A7G2CFK3|A0A7G2CFK3_9TRYP   MESITFLA------NGESAVLEIKDGVS----EKDINKFLDLNSVPLFGTITPENFKNYM    239
tr|S9V6G7|S9V6G7_9TRYP           MESIVFYS------GGKVHK--YTDDLLSSDFEDKFTKFLNFAKVATFGEVTPSSFKNYL    242
tr|S9VLI6|S9VLI6_9TRYP           MESIVFYS------GGKVHK--YTDDLLSSDFEDKFTKFLNFAKVATFGEVTPSSFKNYL    242
tr|A4HQL6|A4HQL6_LEIBR           MESVTVYR------KGKERE--AYDSASPMT-DASMKNFLATAVLDFFGELGQMSFQKYM    239
tr|A7YBW7|A7YBW7_LEIAM           MESVTVYR------QNMERE--AYTGATPIT-MESVNSFLATATLDFFGELGQQSFQKYM    239
tr|Q4Q059|Q4Q059_LEIMA           MESVTVYR------KNAERE--AYTGATPMT-AESVKSFLTSAVLDYFGELGQESFQKYM    239
tr|Q8I8E1|Q8I8E1_LEIMA           MESVTVYR------KNAERE--AYTGATPMT-AESVKSFLTSAVLDYFGELGQESFQKYM    239
tr|A4ICD5|A4ICD5_LEIIN           MESVTVYR------KNAERE--AYTGAAPMT-AESVKRFLATAVLDYFGELGQESFQKYM    239
tr|B3VA16|B3VA16_LEIDO           MESVTVYR------KNAERE--AYTGAAPMT-AESVKRFLATAVLDYFGELGQESFQKYM    239
tr|A0A0M9G4E8|A0A0M9G4E8_LEPPY   MESVTVYR------HDKEHE--AYAGVLPVT-VESAKQFLSEAQLDFLGELGQETFQSYM    239
tr|A0A0N0P805|A0A0N0P805_LEPSE   MESVTVYR------HGTERE--AYSGELPMT-VESAKQFLTEAMIDSFGELGEGSFMNYM    239
tr|A0A7E6F1P9|A0A7E6F1P9_OCTVU   ANSIVVFVPERFQTEYEPKKHVLNKDDAQ---RSDILSFIEKNQFPLVGHYNFNNEKKYN    278
tr|E3NBU5|E3NBU5_CAERE           SDDIAVFVPKKLHNKFDSNEFKY-DGNYD---TDKIKNFLTHETVGLAGIRTQGNLFQFE    248
tr|A0A4Z2C989|A0A4Z2C989_9TELE   PEGIVLFRPPTLKNKFEDSSVKYSEEKYT---SNKIKRFIQDNVLGFCPHMTEDNKDQLM    246
tr|W5KW40|W5KW40_ASTMX           SEGILLFRPPRLASKFEESVVSH-TGPFS---VTALRRFIRDNIFGICPHLTKENIDQLR    253
tr|A0A0G4EK32|A0A0G4EK32_VITBC   KDEVIAYR------T-DEDPTTCT--GKS---ADEIEAFVKDESFPLFGAISGENYSKYV    236
tr|A0A058Z7G2|A0A058Z7G2_FONAL   APAVVLFK------KFDEGKNILQGSDFS---VDTVQKFVSDNSVALMEDLNQSNYQHFA    249
tr|G4MPX2|G4MPX2_MAGO7           APAIVVYK------TFDEGKAVFD-KKFD---VEEIEKFAKTAATPLIGEVGPETYSDYM    246
tr|A0A8J2IGP4|A0A8J2IGP4_FUSEQ   APALVVYK------AFDERKNTFT-EKFE---EEAISTFISTSATPLIGEVGPETYAGYM    244
tr|A0A8I3AHZ3|A0A8I3AHZ3_9PEZI   APAVVLYK------SFDEGKTTFS-DKFE---VEAIEKFAKTSATPLIGEVGPDTYSGYM    261
                                 :                                   *                 .

tr|W6KQG4|W6KQG4_9TRYP           -KLSH--EKIGILLMGSE-----TTDERIADLRLVAKKYRN---KITVISFEGESNE-VS    285
tr|A0A0S4KL52|A0A0S4KL52_BODSA   -ERG---LPIAWLFVKPSAD---NFEALKTQVSAVASAHQG---QLSVVWVDADKYGAMA    289
tr|A0A1X0NMX9|A0A1X0NMX9_9TRYP   -ELS---KPVGWVLLKPKEE---LSKELQPKLLEVGKKMRQ---HLAILWVDAEQYP-VW    286
tr|Q4E3F7|Q4E3F7_TRYCC           -GIS---GPVAWVLLKPSEE---ESKELKSKLLDVGKKMRR---LMVLLWVDAEQYG-VA    287
tr|A0A3R7L858|A0A3R7L858_TRYRA   -AIS---FPVGWVLLKPSGE---ESQELKPKLVDLGRRMRR---QVVLLWADADQYG-VA    287
tr|A0A422Q717|A0A422Q717_9TRYP   -GLS---TPVGWVLLKPSGE---ASQELKPKLVELGRKMRQ---HVVLLWVDAEQYG-VA    287
tr|A0A7G2CFK3|A0A7G2CFK3_9TRYP   -AAGEHGYAIGFLFTKDE-----KESPETKAAAAVAKQYRDN--KVILVTVNGALYGAFG    291
tr|S9V6G7|S9V6G7_9TRYP           -DISNMKFPLGYLFVTTL-----VCTE-AKAVKAIAEKMRP---RIIFASVDSGLYGGFG    292
tr|S9VLI6|S9VLI6_9TRYP           -DISNMKFPLGYLFVTTL-----VCTE-AKAVKAIAEKMRP---RIIFASVDSGLYGGFG    292
tr|A4HQL6|A4HQL6_LEIBR           -EAN-KEKPLGWVFIDKN-----TDPTLKKSLEAVAEKYRS---QVLMTYIDGDQYRPVS    289
tr|A7YBW7|A7YBW7_LEIAM           -EVN-KDKPLGWVFIDKN-----TDPALKGSLVAVAEKYRS---QVLLTYIDGDQYRPVS    289
tr|Q4Q059|Q4Q059_LEIMA           -EAN-KDKPLGWVFIDKN-----TDSALKGSLVAVAEKYRS---QVLLTYIDGDQYRPVS    289
tr|Q8I8E1|Q8I8E1_LEIMA           -EAN-KDKPLGWVFIDKN-----TDSALKGSLVAVAEKYRS---QVLLTYIDGDQYRPVS    289
tr|A4ICD5|A4ICD5_LEIIN           -EAN-KGKPLGWVFVDKN-----TDPALKGSLVAVAEKYRS---QVLLTFIDGDQYRPVS    289
tr|B3VA16|B3VA16_LEIDO           -EAN-KGKPLGWVFVDKN-----TDPALKGSLVAVAEKYRS---QVLLTFIDGDQYRPVS    289
tr|A0A0M9G4E8|A0A0M9G4E8_LEPPY   -EAN-KAKPLGWLFVDKD-----TAPALKESLVAVAEKFRS---QVLMSWVDGDKYRQVA    289
tr|A0A0N0P805|A0A0N0P805_LEPSE   -EAN-KVKPLGWFFLDKD-----TAPALRKSIAAVAEKYRS---QVLMSWINGDKFRQVS    289
tr|A0A7E6F1P9|A0A7E6F1P9_OCTVU   TKYP---LCLVFYTVDWSFDYRKATQIWRKKIAKIARNYKS----ITFAVANEDDHSNLM    331
tr|E3NBU5|E3NBU5_CAERE           -QKP---IVVVYYNVDYVKD-PKGSNYWRNRVLKVAQNYKR---KVQFAVSNKEEFSSEI    300
tr|A0A4Z2C989|A0A4Z2C989_9TELE   -GKD---LLVAYYDVDYERN-PKGSNYWRNRVMKVAKTFLDEGKKLNFAVANKARFMSVL    301
tr|W5KW40|W5KW40_ASTMX           -KRD---LLTAYYDLDYVHN-PKGSNYWRNRVMKVASQYSSRGL--LFSVANRKDFEDEL    306
tr|A0A0G4EK32|A0A0G4EK32_VITBC   -SRS---QDLVWACMSPD-DAK----KHGDAMRSAASKFRQ---SYSMVHLNTEEFGSHA    284
tr|A0A058Z7G2|A0A058Z7G2_FONAL   -SSP---LPMAVLFVTTDEHRA----TLGKGVEAIAKELKG---KMNFVYLDATKYGAFA    298
tr|G4MPX2|G4MPX2_MAGO7           -SAG---IPLAYIFAETAEERT----TLSEALKSIAEKHRG---AINFATIDAKAFGAHA    295
tr|A0A8J2IGP4|A0A8J2IGP4_FUSEQ   -SAG---IPLAYIFSETEEERK----ELGDALKPIAEKYKG---KINFATIDAKAFGAHA    293
tr|A0A8I3AHZ3|A0A8I3AHZ3_9PEZI   -EAG---LPLAYIFAETAEERE----ELSKALKPIAEKQRG---VINFATIDAKSFGAHA    310
                                                                     .        .   :

tr|W6KQG4|W6KQG4_9TRYP           -KNIGL--PEDTKYPAFVLTHD--NTAYPHPTDIQANSSTIDAFMQKYLNGEVKPIQKSQ    340
tr|A0A0S4KL52|A0A0S4KL52_BODSA   -ERLGV--RKGA-FPAFVVDRS--GEHFVLAEDKEISTAIVGDFVTAVLADTLKPTIRST    343
tr|A0A1X0NMX9|A0A1X0NMX9_9TRYP   -KNLGV--SEDAKYPAFVIAKG--DKHHVLPTSEPATVESIEEFIVKYAEGKTDAELKSQ    341
tr|Q4E3F7|Q4E3F7_TRYCC           -SSLGL--SDDAKYPAFVIARG--EDHFVHPSTEPVTAESIEKFIIEYSEKKLSPEIKSQ    342
tr|A0A3R7L858|A0A3R7L858_TRYRA   -SSLGL--PEDAKYPAFAIVQG--EDHFVHPSAEPVTAESIEKFIVGFSEGKITKEIKSQ    342
tr|A0A422Q717|A0A422Q717_9TRYP   -PSLDL--PEDAKYPAFVIARG--EDHFVHPSTEPVTAEGIEKFIVEVSEGKVAKELKSQ    342
tr|A0A7G2CFK3|A0A7G2CFK3_9TRYP   -DQLGA--PKE--YPSLVIDKS--RTKFLFPPAQKMTEEELKSFLDQFTNGKLAPHLKSE    344
tr|S9V6G7|S9V6G7_9TRYP           -EQLSL--PKDTK-TGFVIEWD--HKHYVFPLDKKIESKALEEFITAVLDKKISPTLKSE    346
tr|S9VLI6|S9VLI6_9TRYP           -EQLSL--PKDTK-TGFVIEWD--HKHYVFPLDKKIESKALEEFITAVLDKKISPTLKSE    346
tr|A4HQL6|A4HQL6_LEIBR           -RQLGI--PEGAEFPAFVIDHD--RRHHVMPVDIPVTVESITEFIEKYIKGETKQTLMSD    344
tr|A7YBW7|A7YBW7_LEIAM           -RQLGI--PEDAKFPAFVVDYE--HRHHVMDAATPVTSESVAAFVEKYIKGETQQTVMSD    344
```

```
tr|Q4Q059|Q4Q059_LEIMA              -RQLGI--PEDAKFPAFVVDFE--RRHHVMGTDTPVTSESVAAFVEKYVKGETKQTVMSD    344
tr|Q8I8E1|Q8I8E1_LEIMA              -RQLGI--PEDAKFPAFVVDFE--RRHHVMGTDTPVTSESVAAFVEKYVKGETKQTVMSD    344
tr|A4ICD5|A4ICD5_LEIIN              -RQLGI--PEDAKFPAFVVDFE--RRHHVMETYTPVTAESVAAFVEKYIKGETKQTLMSD    344
tr|B3VA16|B3VA16_LEIDO              -RQLGI--PEDAKFPAFVVDFE--RRHHVMETYTPVTAESVAAFVEKYIKGETKQTLMSD    344
tr|A0A0M9G4E8|A0A0M9G4E8_LEPPY      -FQLGM--PKDVKFPAFVLDFE--RHHHVMPTETPITAESVSEFVEKYSKGETAETLMSE    344
tr|A0A0N0P805|A0A0N0P805_LEPSE      -LQLGV--PKDVKFPVLVIDFE--RRHHLMPIDIPITAESVSEFVGKYIKGETTQTMMSE    344
tr|A0A7E6F1P9|A0A7E6F1P9_OCTVU      -KDFGFEDSGE-DMN-IGIMT--EEKRYTMEPMDEFESSHIESFLDDFVSGNLKPHIKSQ    386
tr|E3NBU5|E3NBU5_CAERE              -ETNGLGERKDSDKPIVAMLTN-EG-KFPMDQEFSV--ENLQQFVDEVLAGNSEPYMKSE    355
tr|A0A4Z2C989|A0A4Z2C989_9TELE      -SEFGLED-HSSAGPLVTIRTA-KGEKYAMTEEFLPDGKALERFLLSYFDGSLKPYLKSE    358
tr|W5KW40|W5KW40_ASTMX              EEDYGLGSTEGSEVPFVTIRTR-LGHKYSMREEFTRDGKSLERFLEDYYANSLKRYIKSE    365
tr|A0A0G4EK32|A0A0G4EK32_VITBC      -ENA-L---GVSEFPAVVVQKK--SGRYVYPTELAIEAEKLVSFLEDVSAGKVEKAIKSE    337
tr|A0A058Z7G2|A0A058Z7G2_FONAL      -EALNL---KQ-EFPAFAIQGQ-KGAKFPLSQDAAITAEAIGKLAKGMLDGSVTPTLKSE    352
tr|G4MPX2|G4MPX2_MAGO7              -GNLNL---KADKFPAFAIQETVKNQKFPFDQDKEITAEAISKFVEDFVAGKIEPSVKSE    351
tr|A0A8J2IGP4|A0A8J2IGP4_FUSEQ      -GNLNL---KTDKFPSFAIQEVVKNQKFPFDQEKEITHDSIAKFVEEFDAGKIEPSIKSE    349
tr|A0A8I3AHZ3|A0A8I3AHZ3_9PEZI      -GNLNL---QADKFPAFAIQETVKNQKFPFDQEKKITVEAITSFVDDFVAGKIEPSIKSE    366
                                                  . :       .             :  :             *

tr|W6KQG4|W6KQG4_9TRYP              PIPEEPTKDGLTILVGKTIDSY-IKKGKNLMIFFNAPWCGHCNKLHPIYDEFAKSHG---    396
tr|A0A0S4KL52|A0A0S4KL52_BODSA      EAPEEHTKDGLTTVVGSTFDDLVINSGKDVLIEFYAPWCGHCKKLQPTFEQVAKELK---    400
tr|A0A1X0NMX9|A0A1X0NMX9_9TRYP      PVPEVETVDGLTTIVGKTMNKY-LSSGKDILIEFFAPWCGHCKNFAGTYAKVAKEFE---    397
tr|Q4E3F7|Q4E3F7_TRYCC              PVPEIETVEGLTTVVGKTLDKY-LSSGKDMLIEFFAPWCGHCKNLAPIYAKVAKEFE---    398
tr|A0A3R7L858|A0A3R7L858_TRYRA      PIPEVETVEGLTTVVAKTLDKH-LSSGKDILIEFFAPWCGHCKNLAPTYAKLAKELE---    398
tr|A0A422Q717|A0A422Q717_9TRYP      PVPEVETVEGLTTVVAKTLDKH-LSSGKDMLIEFFAPWCGHCKNLAPTYAKLAKEFE---    398
tr|A0A7G2CFK3|A0A7G2CFK3_9TRYP      AIPEKETVDGLTTLVGKSFESH-LNKGKDMFVLFYAPWCGHCKKLHPDYEKMAKELE---    400
tr|S9V6G7|S9V6G7_9TRYP              EAPKTETTNGLTTLVGTTFAKY-VQEKKPMFVLFYAPWCGHCKKLHPVFDKLAEAFE---    402
tr|S9VLI6|S9VLI6_9TRYP              EAPKTETTNGLTTLVGTTFAKY-VQEKKPMFVLFYAPWCGHCKKLHPVFDKLAEAFE---    402
tr|A4HQL6|A4HQL6_LEIBR              EVPATETVNGLTTVVGQTISKY-TDGKQNVMLLFYAPWCGHCQKLHPDYEKMAENLQ---    400
tr|A7YBW7|A7YBW7_LEIAM              AIPAKETVNGLTTVVGHTFSKY-TDSTQNVMLLFYAPWCGHCQKLHPAYEKVAKSFE---    400
tr|Q4Q059|Q4Q059_LEIMA              AIPAKETVNGLTTVVGQTFAKY-TDGTQNVMLLFYAPWCGHCKKLHPVYDKVAKSFE---    400
tr|Q8I8E1|Q8I8E1_LEIMA              AIPAKETVNGLTTVVGQTFAKY-TDGTQNVMLLFYAPWCGHCKKLHPVYDKVAKSFE---    400
tr|A4ICD5|A4ICD5_LEIIN              AIPAKETVNGLTTVVGQTFAKY-TDGTQNVMLLFYAPWCGHCQKLHPVYEKVAKSLE---    400
tr|B3VA16|B3VA16_LEIDO              AIPAKETVNGLTTVVGQTFAKY-TDGTQNVMLLFYAPWCGHCQKLHPVYEKVAKSLE---    400
tr|A0A0M9G4E8|A0A0M9G4E8_LEPPY      SVPEVETVEGLTTIVGKTVDKY-TDGSKNIFVLFYAPWCGHCKKLHPEFEKLAKELE---    400
tr|A0A0N0P805|A0A0N0P805_LEPSE      SVPDVETVDGLTTIVGHTIAKY-TDGSKNAFVMFYAPWCGHCKKLHPDFEKLAKELE---    400
tr|A0A7E6F1P9|A0A7E6F1P9_OCTVU      PIPKKDK-GPVVTVVANNFDSVVLDKSKDVLIEFYAPWCGHCQKLEPIYKKLAKSLQ---    442
tr|E3NBU5|E3NBU5_CAERE              PIPEE-Q-GDVKVAVGKNFKQLIMDSDKDVLIEFYAPWCGHCKSLAPKYDELAEKLN---    410
tr|A0A4Z2C989|A0A4Z2C989_9TELE      PIPENND-GPVKVVVAENFDSIVNDDSKDVLIEFYAPWCGHCKSLEPKYKELGEKLA---    414
tr|W5KW40|W5KW40_ASTMX              PVPAKNN-GPVKVVVADTFEEIVNDPEKDVLIEFYAPWCGHCKKLEPKYTELGQELS---    421
tr|A0A0G4EK32|A0A0G4EK32_VITBC      AIPEKND-EAVKVVVAKNFEEMVLRDDKDVMLEVYAPWCGHCKKLEPVYKEFAEKLQKA-    395
tr|A0A058Z7G2|A0A058Z7G2_FONAL      PIPEKQE-GPITVVVGKTYDSIVNEKGKDVLVQFYAPWCGHCKNLIPIYDELAGDLK---    408
tr|G4MPX2|G4MPX2_MAGO7              PIPETND-GPVSVVVAHTYNDIVLDDTKDVLIEFYAPWCGHCKALAPKYEELGALYQKSE    410
tr|A0A8J2IGP4|A0A8J2IGP4_FUSEQ      PIPETQE-GPVTVVVAKSYNDIVLDDTKDVLIEFYAPWCGHCKALAPKYDDLASQYAASE    408
tr|A0A8I3AHZ3|A0A8I3AHZ3_9PEZI      PIPETQE-GPVTVVVAKNYEQIVLDDTKDVLIEFYAPWCGHCKALAPKYEELATLYGESE    425
                                    *         :   *. .   .       :  :: . *******: :    : ...

tr|W6KQG4|W6KQG4_9TRYP              SDN-LIISKIDGTANDFNR-TMFNVNGFPTIYFIPAGSQ-EAMFYNGKRTVEDLAAFVKK    453
tr|A0A0S4KL52|A0A0S4KL52_BODSA      DVDGIRIAQIDAGENDFNT-KLFTVSGFPTLYFVPANGS--PKLFEGSRSVMGILNFLKE    457
tr|A0A1X0NMX9|A0A1X0NMX9_9TRYP      SSD-VIIAAIDATANDVDR-SLFDVTGFPTLYFVPSGGK--PILYNGDRTFYDLYKFIRD    453
tr|Q4E3F7|Q4E3F7_TRYCC              SSD-VIIAAMDATANQMDN-SLFDVSGFPTIYFVPHGGK--PIMYDGGRTFYEIYKFVHE    454
tr|A0A3R7L858|A0A3R7L858_TRYRA      SSN-VIIAAMDATANHVNQ-SVFEVSGFPTIYFVPHGKK--PIIYEGERSFDDMYKFVRE    454
tr|A0A422Q717|A0A422Q717_9TRYP      SSD-VIIAAMDATANHVDQ-SVFAVSGFPTIYFVPHGKP--PISYEGDRTFNDMYEFVRE    454
tr|A0A7G2CFK3|A0A7G2CFK3_9TRYP      KED-VVIAKIDATANDVDP-SKYKVQGFPTIFFIPSGGA--PVSYNGDRSIDDMKKFIKE    456
tr|S9V6G7|S9V6G7_9TRYP              KED-VLIAKLDATENDVDR-ELFEVKGFPTLYFIKNG-A--GISYSGDRTLEDMTKFVRS    457
tr|S9VLI6|S9VLI6_9TRYP              KED-VLIAKLDATENDVDR-ELFEVKGFPTLYFIKNG-A--GISYSGDRTLEDMTKFVRS    457
tr|A4HQL6|A4HQL6_LEIBR              SEN-VMIAKMDATTNDFDR-EKFQVSGFPTIYFIPAGKP--PMMYEGGRSAKEMEEFVRS    456
tr|A7YBW7|A7YBW7_LEIAM              SEN-VIIAKMDATTNDFDR-KKFDVSGFPTIYFIPAGKP--PVAYEGDRTADDMMAFVKS    456
tr|Q4Q059|Q4Q059_LEIMA              SEN-VIIAKMDATTNDFDR-EKFEVSGFPTIYFIPAGKP--PIVYEGGRTADEIQVFVKS    456
tr|Q8I8E1|Q8I8E1_LEIMA              SEN-VIIAKMDATTNDFDR-EKFEVSGFPTIYFIPAGKP--PIVYEGGRTADEIQVFVKS    456
tr|A4ICD5|A4ICD5_LEIIN              SEN-VIIAKMDATTNDFDR-EKFEVSGFPTIYFIPAGKP--PIVYEGGRTADDIKAFVKS    456
tr|B3VA16|B3VA16_LEIDO              SEN-VIIAKMDATTNDFDR-EKFEVSGFPTIYFIPAGKP--PIVYEGGRTADDIKAFVKS    456
tr|A0A0M9G4E8|A0A0M9G4E8_LEPPY      SAD-VIIGKIDATANDFDR-AKFSVNGFPTLFFIPAGGK--TESYEGGRSLAEMKAFVVS    456
tr|A0A0N0P805|A0A0N0P805_LEPSE      AVN-VVIGKIDATTNDFDR-EKFVVNGFPTLYFIPAGGE--PLAYEGGRSTADMKAYVLS    456
tr|A0A7E6F1P9|A0A7E6F1P9_OCTVU      KNQNLVIAKMDATANDVPS--NFKTEGFPTIYFAPTNNKENPVKFTGGRELKDFVKFLEE    500
tr|E3NBU5|E3NBU5_CAERE              KE-DVIIAKMDATANDVPP--LFEVRGFPTLFWLPKNSKSNPIPYNGGREVKDFVNFISK    467
tr|A0A4Z2C989|A0A4Z2C989_9TELE      DDPNIVIAKMDATANDVPS--PYEVSGFPTLYFSPAGQKRNPKKYEGGREVSDFLSYLKR    472
tr|W5KW40|W5KW40_ASTMX              SDPNIVIAKMDATANDVPE--GYDVHGFPTIYFVPAGRKAEPKRYEGAREVKDFLNFLKR    479
tr|A0A0G4EK32|A0A0G4EK32_VITBC      SIDHIVVAKMDGTANESPV-EGFDWSGFPTIFWIPAGAK-GPSKYNGARTTEGLLKWVKE    453
tr|A0A058Z7G2|A0A058Z7G2_FONAL      SNSNIVIAKMDATANDLPLDAGFEIHGFPTIKLVTADGK--VVDFNGERTVEGFKKFIEA    466
tr|G4MPX2|G4MPX2_MAGO7              FKDKVVIAKVDATANDVPD----EIQGFPTIKLYAAGKKDSPATYSGSRTIEDLITFVKE    466
tr|A0A8J2IGP4|A0A8J2IGP4_FUSEQ      FKDKVVIAKVDATLNDVPD----EIQGFPTIKLYPAGGKDAPVTYQGSRTVEDLANFVKE    464
tr|A0A8I3AHZ3|A0A8I3AHZ3_9PEZI      FKDKVVIAKVDATLNDVPD----EIQGFPTIKLYPAGGKSEPVTYSGSRTIEDLIEFVRD    481
                                                : :. :*.  *.             ****:     .         : * *     :  ::

tr|W6KQG4|W6KQG4_9TRYP              HMKEPVEPKQED----------------------------LEGGDL    471
tr|A0A0S4KL52|A0A0S4KL52_BODSA      ASSVTFELPSEDR-------------------------------AEL    473
tr|A0A1X0NMX9|A0A1X0NMX9_9TRYP      HSSSFKDDKGASVPKE-----------EKK-----EEVRSDEDRGDL    484
tr|Q4E3F7|Q4E3F7_TRYCC              HSSTLKDVP---IPEE-----------VKR-----EE-EKNGDDDDL    481
tr|A0A3R7L858|A0A3R7L858_TRYRA      HSSALKDAP---IPEK-----------IKN-----ENKEKNGDEGDL    482
tr|A0A422Q717|A0A422Q717_9TRYP      HSTALKDTP---VSEE-----------VRN-----KEEDENGQEGDL    482
tr|A0A7G2CFK3|A0A7G2CFK3_9TRYP      HSSKKGGDAKKA---------------------------DAGDNADL    476
tr|S9V6G7|S9V6G7_9TRYP              HMEAPSSDDSV----------------------------------DL    470
tr|S9VLI6|S9VLI6_9TRYP              HMEAPSSDDSA----------------------------------DL    470
tr|A4HQL6|A4HQL6_LEIBR              HMTTSSG-----S----------------S------------DEGDL    470
tr|A7YBW7|A7YBW7_LEIAM              HLTASAG-----P-------------------------SDKSDEEDL    473
tr|Q4Q059|Q4Q059_LEIMA              HLTASAA-----P----------------S-----GGPSGNSEEEDL    477
tr|Q8I8E1|Q8I8E1_LEIMA              HLTASAA-----P----------------S-----GGPSGNSEEEDL    477
tr|A4ICD5|A4ICD5_LEIIN              HLTASAA-----P----------------S-----GGPSGNSDEEDL    477
tr|B3VA16|B3VA16_LEIDO              HLTASAA-----P----------------S-----GGPSGNSEEEDL    477
```

```
tr|A0A0M9G4E8|A0A0M9G4E8_LEPPY      HMASTPEATPSSP----------------S-----AAPTDEND-DDL    481
tr|A0A0N0P805|A0A0N0P805_LEPSE      HITEAPELTSS------------------A-----AAPTDDQD-KDL    479
tr|A0A7E6F1P9|A0A7E6F1P9_OCTVU      HSTVSLNTKHI----------------------------------EL    513
tr|E3NBU5|E3NBU5_CAERE              HSTDGLKGFNRDG------------KKK--------------KHTEL    488
tr|A0A4Z2C989|A0A4Z2C989_9TELE      EATNAPVVQEETK-----------KKKK--------------KRVEL    494
tr|W5KW40|W5KW40_ASTMX              EATSSLVLKGVKE-----------DL---------------------    494
tr|A0A0G4EK32|A0A0G4EK32_VITBC      KASKPVVHDEL------------------------------------    464
tr|A0A058Z7G2|A0A058Z7G2_FONAL      NGTAPKPAAAAAAAE-------EEKKEEKKDEKKDEEKKDHHAHGDL    506
tr|G4MPX2|G4MPX2_MAGO7              NGKYKAEVSVEAEKETPVAPAATESEEAKATEAAEKK---KDEHDEL    510
tr|A0A8J2IGP4|A0A8J2IGP4_FUSEQ      NGKYKAEISVKEEGTEEAAPAASEEKKEKKAEKAAEE---EDVHDEL    508
tr|A0A8I3AHZ3|A0A8I3AHZ3_9PEZI      NGKYKAEASKKEEVVAEESQAAPAAT---EGEAAKES---DESHDEL    522
```

### 15. Peptidyl-prolyl cis-trans isomerases (Cyclophilins)

Peptidyl-prolyl disulfide isomerases (PPDIs, also called cyclophilins) accelerate the cis-trans isomerization of the proline peptide bond, thereby accelerating protein folding. They exist both intra- and extracellularly. Interestingly, while many PPDIs are secreted into the endoplasmic reticulum (ER), only a smaller portion of them possesses KDEl-like signals. The rest are likely retained in the ER by indirect protein-protein interactions. The ER-localized PPDIs we found in kinetoplastids are more closely related to bacterial PPDI domains, even though the same PPDI lineage is also found in Dinoflagellates and their relatives. Therefore it is also possible that their KDEL signals evolved convergently, and are not immediately homologous to mammalian PPDIs.

```
tr|A0A7G2CH91|A0A7G2CH91_9TRYP      --------------MRLLTVVTFAALLLVSSTTVAGEDSYWVREMKRVILNAADDYMRYV    46
tr|A0A0M9G5W9|A0A0M9G5W9_LEPPY      -------MARSAV-----PTALVAALLLLLASAVSVQASYWSDEVTRVRTYAAQHYMDRT    48
tr|A0A0N1II12|A0A0N1II12_LEPSE      -----MKMARSAV-----LTVLVAT-LLLLASTISVQASYWSDEVTRVRTYSAQHYMDRT    49
tr|A4HA48|A4HA48_LEIBR              -------MNRFSTMRCVFAAVATLLLLLLTLCVVSVEASYWSNEVNRVRTYTAFNYMARM    53
tr|Q4QDB9|Q4QDB9_LEIMA              -------MARFSSLHSVLAAAATL--LLLALCVVSVEASYWSEEVNRVRTYAAVNYLERI    51
tr|A4HYB4|A4HYB4_LEIIN              -------MTRFSSLHSVLATAATL---LLTLCVLSVEASYWSEEVNRVRTYAAVNYLERI    50
tr|A0A3S5H764|A0A3S5H764_LEIDO      -------MTRFSSLHSVLATAATL---LLTLCVLSVEASYWSEEVNRVRTYAAVNYLERI    50
tr|Q4D932|Q4D932_TRYCC              -----MHRENYFSKMAFCLLGVLFLSCITSVQTVSGDAASHEERMNNYRKRVGRLFMEQK    55
sp|Q09734|MIP_TRYCR                 -----MHRENYFSKIAFCLLGVLFLSCITSVQTVSGDAASHEERMNNYRKRVGRLFMEQK    55
tr|A0A3S5IR72|A0A3S5IR72_TRYRA      MQLRPTKTGKFYSRATLCLMAALMLLSMSVVLTVSGETLSSEVRLNNYRKRVGRAFIKEK    60
tr|A0A3R7NGG2|A0A3R7NGG2_9TRYP      -------------------MAALMLLSLSSAPTVSGETLSPEDRINNYRKRVGRAFIKEK    41
tr|A0A812UP16|A0A812UP16_SYMMI      --------------MK----ASVLSL----LLLF--YGAVAAADMNKYYKRTGKKFLDEK    36
tr|A0A7S1LN17|A0A7S1LN17_NEODS      --------------MRAIATAA-LVA----LLLL--AATAAAGDMTKYYKRTGAKFLSTK    39
tr|A0A250XD25|A0A250XD25_9CHLO      --------------MRIFLLAAAFLC----AILH---------HATAGSNEKGLAFLEEN    33
tr|A0A0M0JR19|A0A0M0JR19_9EUKA      --------------MRSTLLV---------ATLV---------GLVAASNEVGTKFLEEN    28
tr|A0A7S1FII8|A0A7S1FII8_NOCSC      --------------MLSTFALT--TL----VALC---------RVAVASNEFGLSFLEGN    31
tr|R1B618|R1B618_EMIHU              --------------------MA--VL----AVTL---------FSLLASNDVGVKFLEEN    25
tr|R1D2M1|R1D2M1_EMIHU              ---------------------------------------------------VGVKFLEEN    9
                                                                                     .  ::

tr|A0A7G2CH91|A0A7G2CH91_9TRYP      ESWGNVTILPSGMGMNILQRGHGD-VAPGPNDSCLIRYTGMY-------RFPGVFEDTRQ    98
tr|A0A0M9G5W9|A0A0M9G5W9_LEPPY      AKDPNVSALPSGLLVTILRRGAGD-RAPAAEDVCEVHYTIHH-------RFPGIVDDTRD    100
tr|A0A0N1II12|A0A0N1II12_LEPSE      SKSPNVSALPSGMLVTVLGRGSGD-RAPAANDVCEIHYTIHH-------RYPGIVDSTRN    101
tr|A4HA48|A4HA48_LEIBR              AKQPNVSALPSGVLFTIERRGFGD-RAPAPEDKCEMHYTIHQ-------RFPDIVESTRH    105
tr|Q4QDB9|Q4QDB9_LEIMA              AKQPNVSALPSGLLFTIERRGFGD-RAPAAEDKCEMHYTIHY-------RFPGIVENTRH    103
tr|A4HYB4|A4HYB4_LEIIN              AKQPNVSALPSGLLFTIERRGFGD-RAPAAEDKCEMHYTIHY-------RFPGIVESTRH    102
tr|A0A3S5H764|A0A3S5H764_LEIDO      AKQPNVSALPSGLLFTIERRGFGD-RAPAAEDKCEMHYTIHY-------RFPGIVESTRH    102
tr|Q4D932|Q4D932_TRYCC              AAQPDAVKLPSGLVFQRIARGSGK-RAPAIDDKCEVHYTGRL-------RDGTVFDSSRE    107
sp|Q09734|MIP_TRYCR                 AAQPDAVKLPSGLVFQRIARGSGK-RAPAIDDKCEVHYTGRL-------RDGTVFDSSRE    107
tr|A0A3S5IR72|A0A3S5IR72_TRYRA      AAQAGAVTLPSGLVFRRIARGSGK-RAAAADDECEVHYTGKL-------RDGTVFDSSRE    112
tr|A0A3R7NGG2|A0A3R7NGG2_9TRYP      SAQPDAVTLPSGLVFQRIARGSGK-RAAAVDDECEVHYTGKL-------RDGSVFDSSRE    93
tr|A0A812UP16|A0A812UP16_SYMMI      AKEPNIITLKSGMLVEVLKQGPDDGKSPNKGDACEVTYKGTF-------KDGKGFDQ---    86
tr|A0A7S1LN17|A0A7S1LN17_NEODS      EAEEGVYKLPSGMLFKILEKGTGA-TSPRVADPCEVHYEGTL-------PNGNVFDSSFK    91
tr|A0A250XD25|A0A250XD25_9CHLO      KGKPGVITLPSGLQYKVLRQGEGA-YHPTADSSCDCHYKGTL-------IDGTVFDSSYD    85
tr|A0A0M0JR19|A0A0M0JR19_9EUKA      KSREGVITLPSGLQYKVLRSGDGD-SHPTPDSSCECHYEGRIAQKW---PSGETFDSSYK    84
tr|A0A7S1FII8|A0A7S1FII8_NOCSC      KNKEGVITLPSGLQYKVLRAGDGT-DHPTADSSCECHYEGRTAQSY---PDGTTFDSSYA    87
tr|R1B618|R1B618_EMIHU              AKKEGVISLPSGLQYKVLRAGDGD-SHPTASSSCECHYEGRTAQEYSKEPKGDTFDSSYA    84
tr|R1D2M1|R1D2M1_EMIHU              AKKEGVISLPSGLQYKVLRAGDGD-SHPTASSSCECHYEGRTAQEYSKEPKGDTFDSSYA    68
                                             .   * **:         * .      . *   *                   .:.

tr|A0A7G2CH91|A0A7G2CH91_9TRYP      YPFPVRRTPSQMIQGVGEALQYMREGDRWFLYVPYRLAYGREGNATLKIQPYMNLRYDLE    158
tr|A0A0M9G5W9|A0A0M9G5W9_LEPPY      KPYPVRRSPSQLLLGMAEAMQLMREGDRWFLYVPSKLAYGTEGWKERRVAGLANVRVDLE    160
tr|A0A0N1II12|A0A0N1II12_LEPSE      RPYPVRRSPSQLIPGMAEAMQLMREGDHWFLYVPSELAYGTKGWKERRVAGLSIVRVDLE    161
tr|A4HA48|A4HA48_LEIBR              NPYPVCRSPSQLIPGMAEAMQLMREGDRWHLYVPYQLAYGVEGCKEKKVAPMSSVRVEIE    165
tr|Q4QDB9|Q4QDB9_LEIMA              HPYPVRRSPSQLIPGMAEAMQLMREGDRWYLHVPYQLGYGKEGCKEKKVPSLSNLRVEME    163
tr|A4HYB4|A4HYB4_LEIIN              HPYPVRRSPSQLIPGMAEAMQLMREGDRWYLHVPYQLGYGKEGCKEKKVPSLSNLRVEIE    162
tr|A0A3S5H764|A0A3S5H764_LEIDO      HPYPVRRSPSQLIPGMAEAMQLMREGDRWYLHVPYQLGYGKEGCKEKKVPSLSNLRVEIE    162
tr|Q4D932|Q4D932_TRYCC              RGKPTTFRPNEVIKGWTEALQLMREGDRWRLFIPYDLAYGVTGGG-GMIPPYSPLEFDVE    166
sp|Q09734|MIP_TRYCR                 RGKPTTFRPNEVIKGWTEALQLMREGDRWRLFIPYDLAYGVTGGG-GMIPPYSPLEFDVE    166
tr|A0A3S5IR72|A0A3S5IR72_TRYRA      RGRPISFRPSGVIKGWAEALQLMREGDRWLLFIPHELGYGEAGAG-TSIPPFSPLEFDVE    171
tr|A0A3R7NGG2|A0A3R7NGG2_9TRYP      RGEPVSFRPRGVIKGWTEALQLMREGDRWRIFIPHELGYGAAGAG-PKIPPFSPLEFDVE    152
```

```
tr|A0A812UP16|A0A812UP16_SYMMI      --GTTSFAPNQVIKGWTEAMQFMGEGDKWKLYIPYDLAYGERGSP-PKIPPFTPLVFEIE    143
tr|A0A7S1LN17|A0A7S1LN17_NEODS      RGSPSSFAPNQVIKGWTEALQLMREGDKWEVYIPHELAYGARGAG-GVIPGYAALTFKMQ    150
tr|A0A250XD25|A0A250XD25_9CHLO      RGQPTAFAPNQVIKGWTEAMQLMVEGDKWEMYIPSDLGYGDGGSP-PKIGGGDTLIFQME    144
tr|A0A0M0JR19|A0A0M0JR19_9EUKA      RGSPTSFAPNQVIKGWTEAMQLMVEGDKWEMYIPSELGYGERGSP-PNIGGGDVLVFTME    143
tr|A0A7S1FII8|A0A7S1FII8_NOCSC      RGSPTSFAPNQVIKGWTEAMQLMVEGDKWEMYIPSELGYGDRGSP-PKIGGGDVLVFTME    146
tr|R1B618|R1B618_EMIHU              RGSPTSFAPNQVIKGWTEAMQLMVEGDKWEMYIPSELGYGDSGSP-PKIGGGDVLVFTME    143
tr|R1D2M1|R1D2M1_EMIHU              RGSPTSFAPNQVIKGWTEAMQLMVEGDKWEMYIPSELGYGDSGSP-PKIGGGDVLVFTME    127
                                             *  :: *  **:* * ***:* :.:*  *.**  *         :        :   ::

tr|A0A7G2CH91|A0A7G2CH91_9TRYP      VVSCDAPSGKKSAEID-AFLEPL--------LKR--PMPS-------------KT-----    189
tr|A0A0M9G5W9|A0A0M9G5W9_LEPPY      VLKCENPRGKTSEEID-AYLAPF--------LKT--PMPA-------------KS-----    191
tr|A0A0N1II12|A0A0N1II12_LEPSE      VFKCESPHGKTSEEID-AYLAPY--------LKT--PMPP-------------KS-----    192
tr|A4HA48|A4HA48_LEIBR              LYKCKSPSGKTSTEID-AYLAQF--------VKS--PMPV-------------KT-----    196
tr|Q4QDB9|Q4QDB9_LEIMA              LYKCESASGKTSAEID-AYLAKY--------MKT--RIPE-------------KA-----    194
tr|A4HYB4|A4HYB4_LEIIN              MYKCESASGKTSAEID-AYLAKY--------MKT--RIPE-------------KA-----    193
tr|A0A3S5H764|A0A3S5H764_LEIDO      LYKCESASGKTSAEID-AYLAKY--------MKT--RIPE-------------KA-----    193
tr|Q4D932|Q4D932_TRYCC              LISIKDGGKGRTAEEVDEILRKAEEDREDM------------------------------    196
sp|Q09734|MIP_TRYCR                 LISIKDGGKGRTAEEVDEILRKAEEDREDM------------------------------    196
tr|A0A3S5IR72|A0A3S5IR72_TRYRA      LLRIKGGGTGRTAEEVDKILREVEGDRGDM------------------------------    201
tr|A0A3R7NGG2|A0A3R7NGG2_9TRYP      LIRIKGGGTGRTAEEVDKILREAEEERGDM------------------------------    182
tr|A0A812UP16|A0A812UP16_SYMMI      IHKVKSGGKPVSEARD--MLKNAIATGAEL------------------------------    171
tr|A0A7S1LN17|A0A7S1LN17_NEODS      LLKVKSGGKPAAEADA--AIQEKLGK---AYADL--------------------------    179
tr|A0A250XD25|A0A250XD25_9CHLO      LIKIKGGRVPALKCVP--STKTDCSEKEQAYIDKVSTLDTTGIAKELVRLKAMKSKPMTP    202
tr|A0A0M0JR19|A0A0M0JR19_9EUKA      ILKIQGKKVPANKCDV--KTYAGCSEKEETYIKSKVEMTAEALGAEIKRLGGMASQKMSS    201
tr|A0A7S1FII8|A0A7S1FII8_NOCSC      ILKIKGNKVPASRCDV--KSLEGCSDKEKEYITKQSANGKEKVESELNRLSGMKGGSMKD    204
tr|R1B618|R1B618_EMIHU              IIKIKGEKVPASKCDV--KSLDGCTEQEAKYASSKKSLDGDGIAAEMKRLQGIAGQKMGP    201
tr|R1D2M1|R1D2M1_EMIHU              IIKIK-------------------------------------------------------    132
                                    :   .

tr|A0A7G2CH91|A0A7G2CH91_9TRYP      PKIDWEKM-----------------    197
tr|A0A0M9G5W9|A0A0M9G5W9_LEPPY      MPINYADL-----------------    199
tr|A0A0N1II12|A0A0N1II12_LEPSE      VLAYDTDM-----------------    200
tr|A4HA48|A4HA48_LEIBR              EPLDYADL-----------------    204
tr|Q4QDB9|Q4QDB9_LEIMA              APVDYTDL-----------------    202
tr|A4HYB4|A4HYB4_LEIIN              APVDYTDL-----------------    201
tr|A0A3S5H764|A0A3S5H764_LEIDO      APVDYTDL-----------------    201
tr|Q4D932|Q4D932_TRYCC              -------------------------    196
sp|Q09734|MIP_TRYCR                 -------------------------    196
tr|A0A3S5IR72|A0A3S5IR72_TRYRA      -------------------------    201
tr|A0A3R7NGG2|A0A3R7NGG2_9TRYP      -------------------------    182
tr|A0A812UP16|A0A812UP16_SYMMI      -------------------------    171
tr|A0A7S1LN17|A0A7S1LN17_NEODS      -------------------------    179
tr|A0A250XD25|A0A250XD25_9CHLO      ELLNWLERRIRILGLLDKTEL----    223
tr|A0A0M0JR19|A0A0M0JR19_9EUKA      TQSVWLSKRVGLLNKLKELKGKEEL    226
tr|A0A7S1FII8|A0A7S1FII8_NOCSC      DKLGWLNKRINLLQKLKDEL-----    224
tr|R1B618|R1B618_EMIHU              KQASWLAKRLSLLTKLKDEL-----    221
tr|R1D2M1|R1D2M1_EMIHU              -------------------------    132
```
